# Decoding pseudoexon activation by deep intronic variants using a high-throughput minigene assay in the *NF1* gene

**DOI:** 10.64898/2026.10.20.760351

**Authors:** Eliška Svobodová, Přemysl Souček, Kamila Réblová, Zdeněk Kubát, Marie Krátká, Tomáš Freiberger, Lucie Kajan Grodecká

## Abstract

Deep intronic variants can cause aberrant splicing, often manifested as pseudoexon inclusion, which represents a significant challenge in clinical diagnostics. Pseudoexon activations have been repeatedly described in the NF1 gene, making it a promising genomic context for studying underlying mechanisms. Here, we developed a specialized minigene assay by combining high-throughput mutagenesis and next-generation sequencing to evaluate the splicing effect of 501 deep intronic variants within NF1 intron 3. We identified and validated eleven pseudoexon-activating variants, including seven novel ones, which clustered into four activation-prone hotspots. Additionally, we detected a rare, out-of-frame wild-type pseudoexon whose inclusion was significantly enhanced by three specific mutations. Using this experimentally validated dataset, we benchmarked SpliceAI performance and ESRseq scores to refine variant prioritization. This integrated high-throughput framework offers a robust starting point for the systematic functional prioritization of deep intronic variation, providing a scalable model for variant annotation in other disease-associated genes.

**GRAPHICAL ABSTRACT:** 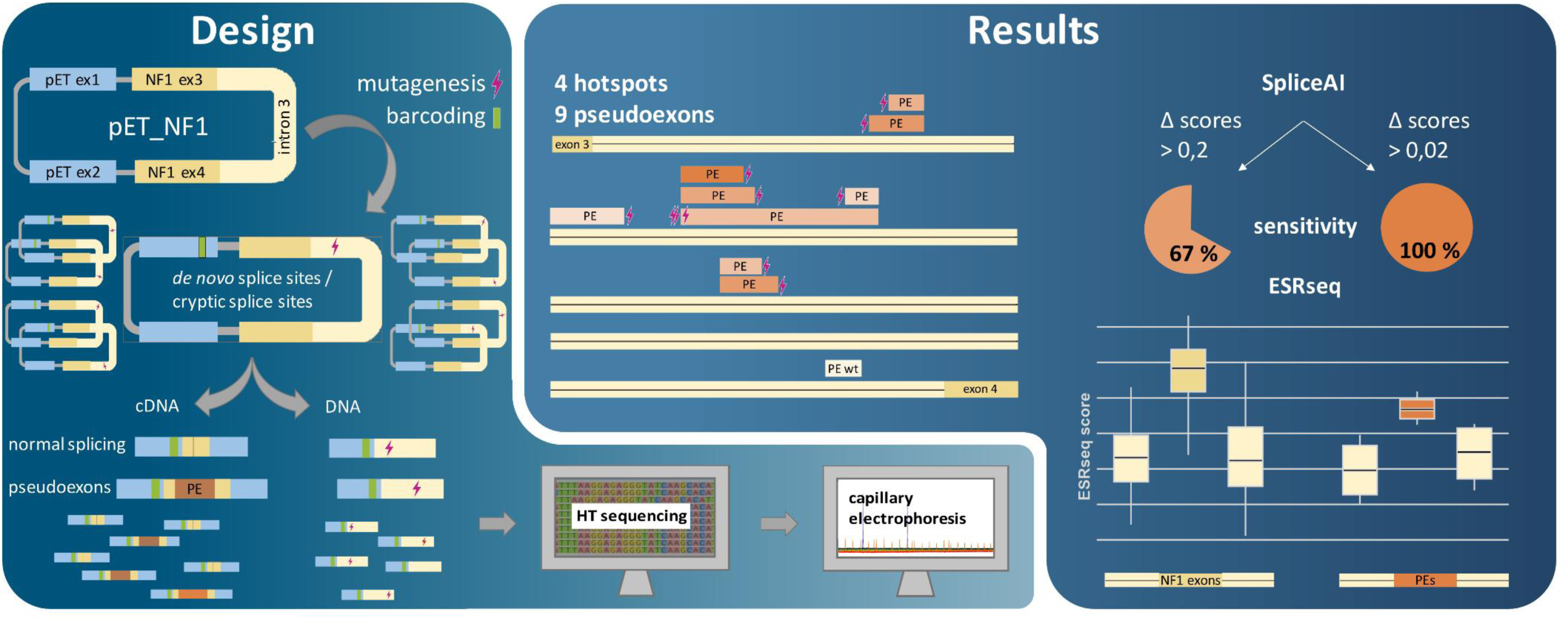

## INTRODUCTION

Deep intronic variants, located >100 bp from exon-intron boundary, were described in at least 77 different genes and associated with numerous disorders. Their contribution to monogenic diseases is emerging more frequently nowadays due to the application of genome sequencing approach in clinical studies (1–3). Introns generally contain a significant number of motifs with high similarity to authentic splice signals, known as cryptic splice sites. Given the great diversity in authentic splice site sequences(4), even a single nucleotide change can activate these normally inactive cryptic site. These deep intronic variants can be particularly challenging to detect and interpret, as they often escape conventional genetic screening and can have unpredictable effects on splicing.

Aberrant splicing can manifest in multiple forms; here we focus on pseudoexon (PE) inclusion caused by deep intronic variations. Even among experts, terminology is inconsistent, with “cryptic exons”, “poison exons” or “pseudoexons” often used interchangeably (5, 6). Common definition describes PEs as intronic sequences with the potential to be included into a final transcript but not normally recognized by the splicing machinery. They are flanked by at least one splice site, and range from 30-340 bp, (2) with a median length of 112 bp (7). However, this definition does not distinguish between predicted and experimentally validated events. In this study therefore, we define pseudoexons as originally intronic sequences with confirmed inclusion.

Occasionally, intronic sequence can be included even without an activating variant. This phenomenon is sometimes referred to as a wild-type pseudoexon (in this study abbreviated as “PEwt”) (5). The one characteristic that distinguishes it from an alternative exon, is the extreme rarity of its inclusion. If PEwts contain a premature termination codon (PTC), they are sometimes described as poison exons. However, without a known regulatory function these assumptions may be misleading (6).

Several mechanisms can activate PE inclusion (7). The most common is a deep intronic single-nucleotide variant creating a *de novo* splice site – more frequently donor site than acceptor – or strengthening pre-existing cryptic splice site (8). Less commonly, variants may create or disrupt splicing regulatory elements (SREs), altering the local regulatory landscape (9–12). The surrounding SRE composition is critical, as even small shifts can increase PE inclusion levels (7). Additional influences include structural variants (13). as well as smaller changes and SNVs that act as modifiers rather than primary drivers (14). Moreover, PEs can be activated from multiple sites with varying levels of inclusion depending on the affected regulatory element (15), thereby revealing activation-prone regions.

PEs are frequently studied based on *in-silico* predictions, but computational tools are often fallible and unable to provide accurate predictions of PE inclusion, largely due to the limited amount of available training data. SpliceAI, a widely used prediction tool, currently outperforms other algorithms, with an average precision of >90 % for variants activating PE inclusion (16). We evaluated its performance on a set of experimentally validated variants with varying impacts on PE inclusion.

Here, we developed a high-throughput functional minigene-based assay combined with next-generation sequencing to systematically evaluate deep intronic variants in NF1 intron 3. We analysed more than 500 single-nucleotide variants with potential splicing effects, enabling a comprehensive assessment of sequence determinants underlying PE inclusion. This approach allowed us to identify multiple regions predisposed to PE activation and provided insight into sequence features beyond canonical splice sites that contribute to aberrant splicing.

## MATERIAL AND METHODS

### Gene annotation

Sequence of the *NF1* gene was acquired from the Ensembl database using the transcript NM_001042492.3. For the purpose of this study, nucleotide numbering was simplified and is referenced relative to intron 3. Therefore, the first base of the intronic sequence is labelled as 1, corresponding to c.288+1 position in the transcript. In case of description of a splice site position, the number refers to G from GT dinucleotide in donors and AG in acceptors.

### Minigene construction

Expression vectors used in this study were derived from the Exontrap plasmid pET01 (MoBiTec), modified in our laboratory (17). A portion of the *NF1* gene sequence was inserted into this plasmid, including the entire intron 3 and its flanking exons 3 and 4 (with 192 bp of upstream intron 2 and 349 bp of downstream intron 4) using ApaI and XbaI restriction sites within the multiple cloning site. This minigene construct was named pET-NF1. For better manipulation during sequencing, nineteen AsiSI restriction sites were separately introduced into individual pET-NF1 plasmids, each shifted approximately 250 bp downstream from the previous one, creating a set of plasmids named pET-NF1_AsiSI1 to pET-NF1_AsiSI19 (Fig. 1). The sequences of these constructs were verified by Sanger sequencing and they subsequently served as default plasmids for further libraries preparation.

**Figure 1:**
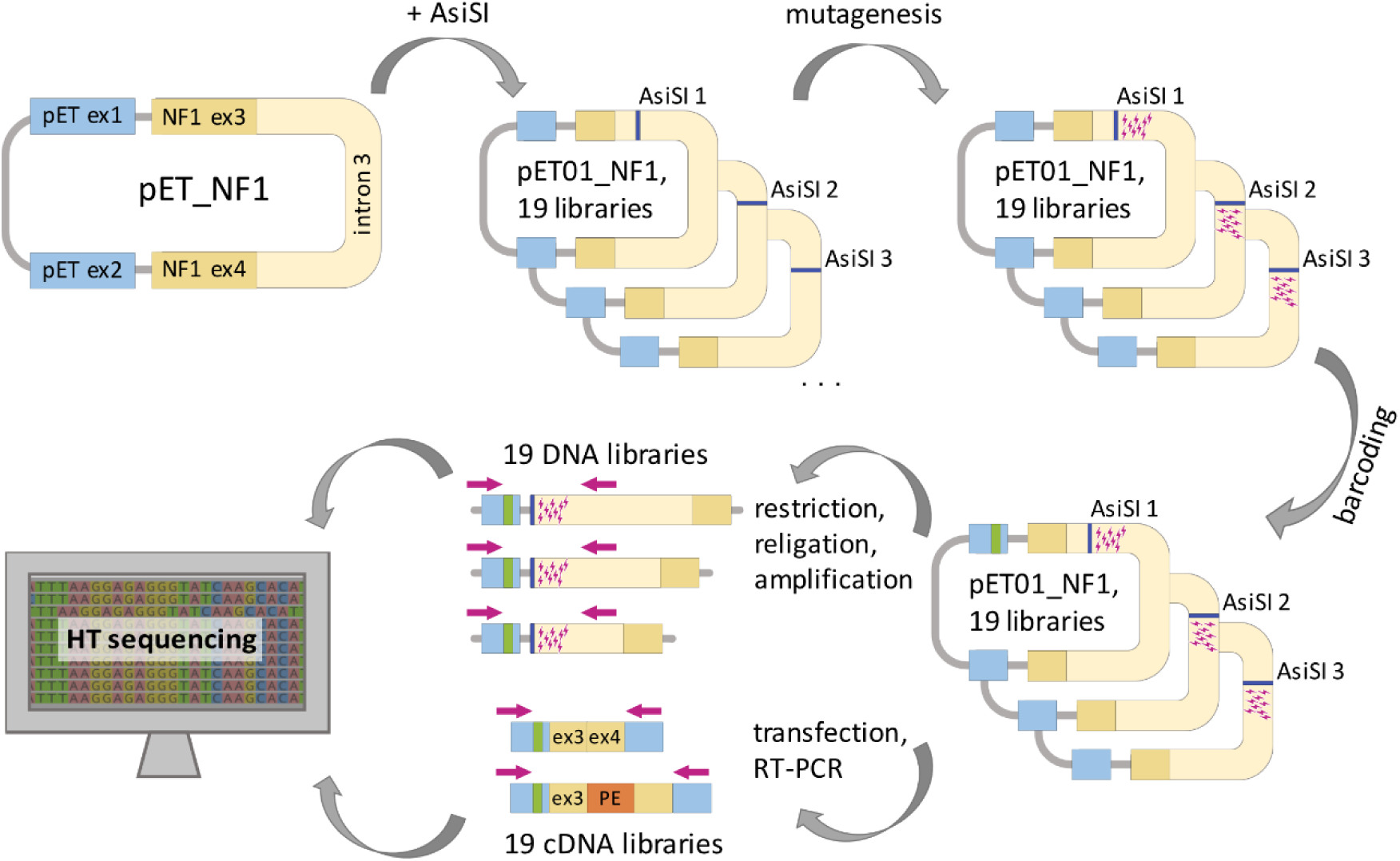
Schematic overview of the workflow using a modified pET01 plasmid containing the NF1 insert.

### Variant selection

Variants were divided into four groups: creating *de novo* donors/acceptors and strengthening cryptic donor/acceptor sites (Tab. S1). Splice site strength considered for PE inclusion was assessed based on MaxEntScan (ME) scores; thresholds were set at 5.66 (donor) 3.4 (acceptor), based on literature (8.66±3.00 for donor and 6.79±3.39 for acceptor splice sites) (18).

### Plasmid library preparation

Studied splicing variants were introduced into the expression vectors via site-directed PCR mutagenesis. Primers were designed using freely available QuickChange Primer Design tool from Agilent (https://www.agilent.com/store/primerDesignProgram.jsp). Premixed PrimeSTAR Max DNA Polymerase (TaKaRa, #R045A) was used for mutagenesis following the provided protocol. Subsequently, 1 µl of DpnI enzyme (Thermo Scientific) was added to restrict the original DNA strands.

DH5α strand of competent *Escherichia coli* (Thermo Scientific, #18265017) was chemically prepared for the transformation process using a heat-shock approach. If a sufficient number of colonies was reached, they were scraped and pooled into one homogenous glycerol stock and stored at -80 °C. Plasmid libraries were isolated from bacterial cultures after overnight incubation at 37°C in LB medium using the QIAprep Spin MiniPrep kit (Qiagen, ID.27106) according to the given protocol following all extra steps for maximal yield.

To enable variant-specific tracking at both the DNA and RNA levels, unique molecular barcodes were inserted into the first pET exon of the minigene. The most effective approach was again a site-directed PCR mutagenesis using primers GCGAAGTGNNNNNNCAGATCTCGAGGAC for forward and GTCCTCGAGATCTGNNNNNNCACTTCGC for reverse. For barcoding and all subsequent steps in NGS library preparation the QIAGEN Plasmid Mini Kit (Qiagen, ID.12125) was used for purification. Only a few colonies were verified by Sanger sequencing to confirm the efficiency of the process.

### Cell culture and transfection

HeLa cell lines (obtained from European Collection of Authenticated Cell Cultures (ECACC), #93021013) were used for all transfection experiments in this study. Cells were maintained in RPMI 1640 with L-glutamine (Sigma Aldrich), supplemented with 10% FBS (Sigma-Aldrich). They were cultured in humidified chamber at 37°C with 5% level of CO2.

Twenty-four hours before transfection, cells were seeded into 12-well plate aiming for 30% confluence. Transfection was subsequently performed using XtremeGene9™ DNA Transfection Reagent (Merck, XTG9-RO). Cells were incubated for another 24 hours and processed to RNA analysis. Transfections were conducted either with one unique plasmid per analysis or, in the case of NGS experiments, with the entire plasmid library. Experiments were mostly done in biological triplicates.

### RNA isolation and RT-PCR analysis

Total RNA from transfected HeLa cells was extracted using Quick-RNA MiniPrep (Zymoresearch, #R1055) following the manufacturer’s protocol. cDNA synthesis was performed using Transcriptor First Strand cDNA Synthesis Kit (Roche, #04897030001) or SuperScript™ II One-Step RT-PCR System (Thermo Scientific, #10928042) with random hexamers. Approximately 500 ng of extracted RNA was used per reaction.

In case of a standard minigene assay, the effect on splicing was evaluated using capillary electrophoresis fragment analysis. The PCR was performed using Taq DNA Polymerase (TopBio) and a set of primers: CAGCACCTTTGTGGTTCTCA tagged with FAM for forward and GCACTGATCCACGATG for reverse. Both primers align to pET exons of the vector eliminating the effect of a possible DNA contamination.

### NGS library preparation and data processing

To obtain a suitable length of fragments used for NGS-based isoform analysis, plasmids were digested by combination of AsiSI and PacI restriction enzymes (Thermo Scientific) following conditions calculated by the Double Digest Calculator [online tool](https://www.thermofisher.com/in/en/home/brands/thermo-scientific/molecular-biology/thermo-scientific-restriction-modifying-enzymes/restriction-enzymes-thermo-scientific/double-digest-calculator-thermo-scientific.html). The reaction was performed on 3 µg of plasmid DNA. The appropriate fragment was separated by agarose gel electrophoresis, cut and purified using QIAquick Gel Extraction Kit (Qiagen, ID.28704) according to the manufacturer’s instructions. Extracted fragment (50 ng) was re-ligated using 1 μl of T4 ligase (Thermo Scientific) in a 10 µl reaction volume. For plasmids pET-NF1_AsiSI17, 18 and 19, a combination of three restriction enzymes had to be used, allowing us to securely distinguish desired fragment. First combination was of AsiSI and SmaI followed by restriction with PacI.

The PCR amplification of the re-ligated plasmid libraries was performed in two rounds. The first round used minigene-specific set of primers – forward primer NF1_seq_F in combination with a different reverse primer for each plasmid (Tab. S2). In addition, forward primer contained an Illumina adapter sequence TCGTCGGCAGCGTCAGATGTGTATAAGAGACAG on its 5’ end and each reversed primer contained GTCTCGTGGGCTCGGAGATGTGTATAAGAGACAG sequence on its 3’ end. The second round used Nextera XT primers (Illumina) attaching indexes and sequencing adapters. Both reactions used hot start Q5 High Fidelity DNA Polymerase (New England Biolabs, #M0491L) for increased accuracy. After second PCR amplification the libraries were pooled, mixed with 1% PhiX and sequenced on an Illumina platform on 600 cycles cartridges (300 cycles on each strand) using a custom reverse primer complementary to exon 4 (TGCATGCTGGTTTCCTTCACGACA) for the DNA libraries.

Subsequently, cDNA libraries also underwent two rounds of PCR amplification, first with minigene-specific primers with Illumina adapters and second using Nextera XT primers. All Illumina NGS data was processed using custom scripts (code available at Zenodo, see Data availability section).

### Sequencing efficiency

Out of 604 designed variants we obtained 517, resulting in final yield of 85, 6%. The minimum sufficient read depth was set to 30 DNA reads per barcode. Variants not meeting this threshold were excluded from further analysis. Double variants were always in combination with another individually characterized variant and were evaluated manually: 15 (paired with another negative variant) showed no impact on splicing and were retained in the dataset, whereas 2 produced inconclusive results (paired with variant showing splice-altering potential) and were excluded to avoid possible false positive/negative interpretations.

Due to technical issues, we had to resequence some of the libraries. Specifically, library 14 was lacking all DNA data and, despite sufficient coverage, libraries 3 and 4 were missing more than 50% of designed variants. We prepared those two libraries anew. Due to cost-effectiveness, we engaged Nanopore sequencing. To assess Nanopore data reliability, we reanalysed one of the libraries, demonstrating substantial agreement between the two methods (Tab. S3).

### Nanopore sequencing and data processing

Plasmid libraries were linearized using the PstI restriction enzyme and purified by ethanol precipitation. Sequencing libraries were prepared using the Ligation Sequencing Kit V14 (Oxford Nanopore Technologies, SQK-LSK114) according to the manufacturer’s protocol, starting with 100 fmol of DNA per library for adapter ligation. For each run, two libraries were pooled and loaded onto a Flongle Flow Cell (FLO-FLG114). Sequencing was performed on a MinION Mk-1B device (Oxford Nanopore Technologies) for 24 hours.

Raw Nanopore sequencing data were basecalled using Dorado v0.9.1 (Oxford Nanopore Technologie) in duplex mode with the SUP (super-accuracy) model (dna_r10.4.1_e8.2_400bps_sup@v5.0.0), using the stereo duplex model (dna_r10.4.1_e8.2_5khz_stereo@v1.3) for duplex pairing. Reads were aligned to a custom plasmid reference sequence during basecalling. The resulting BAM file was sorted and indexed using SAMtools v1.21 (19). Quality metrics were assessed using NanoPlot with a maximum read length filter of 20, 000 bp (20). Variants were called with custom script, using SAM tools. For more details, see the Supplementary text 1.

### Evaluation of pseudoexon inclusion levels

To eliminate the effect of technical artefacts on assessment of PE inclusion levels, we defined a set of filters for our dataset. From any further analysis, we excluded ambiguous variants (e.g., insertions, deletions, or multiple variants) and variants that had less than 30% variant-containing reads at the DNA level. To avoid interference from low-read barcodes affecting the significance of the statistical analysis, we excluded all barcodes that did not reach at least 50 reads of standard wild-type splicing (exon 3 to exon 4). Simultaneously, we excluded any combination of a variant and PE that showed less than three reads containing that PE. Reads containing errors or unassigned positions (marked as NA) were also excluded from the analysis. The only two exceptions were reads [1018-NA] and [NA-1387] which were caused by inaccurate alignment at the splicing junction. We manually verified those reads in individual variants otherwise fulfilling set criteria and confirmed their positions as [1018–1135] or [1018–1154] and [1018–1387].

The differential usage of PEs between mutant and wild-type (WT) sequences was evaluated using a Generalized Linear Model (GLM) with a quasibinomial distribution. This model was chosen to account for the overdispersion typical of sequencing-based read counts. To stabilize the model estimates and handle zero-count observations, a Laplace correction (epsilon = 0.1) was applied to the read counts. P-values were adjusted for multiple testing using the Benjamini-Hochberg False Discovery Rate (FDR) procedure. For a variant to be considered significantly altering a PE inclusion, it had to meet two criteria: an FDR-adjusted p-value < 0.05 and a minimum absolute difference in percent usage of 0.5% compared to the WT reference.

### ESRseq score statistical analysis

For computations of ESRseq score of pseudoexnic and their flanking regions, we followed the suggested ranges for testing: -95 to -15 for upstream regions, and +7 to +95 for downstream regions (21), eliminating the respective splice site sequences. For exons and experimentally validated PEs, the entire sequence of the body was inspected from +2 to -4 bp. Several splice sites were used repeatedly by different PEs: to ensure that each sequence was included only once, we deliberately removed all duplicities and always chose the longer PE for analysis.

To create a negative control group of putative PEs, we selected regions in which no aberrant splicing events were observed, with minimum of 50 bp safe distance from even marginally used splice sites. To limit any overlap in the tested sequences, the negative group went through second selection process, where we filtered out all the splice sites distant less than 50 bp from another upstream splice site of the same kind (i.e. acceptor or donor).

Statistical significance of ESR score differences was evaluated using both two-tailed t-tests and Wilcoxon tests. A paired t-test and Wilcoxon signed rank test with continuity correction were employed to compare PEs and exons with their respective flanking intron regions. An unpaired Wilcoxon rank sum test with continuity correction and Welch’s t-test were used for comparisons between different sequence categories (e.g., experimentally validated vs. putative PEs), to account for unequal sample sizes and variances. To control the false discovery rate (FDR), all p-values were adjusted using the Benjamini-Hochberg (BH) procedure.

## RESULTS

### High-throughput minigene assay to asset splicing effect of intronic variants

To analyse the splicing effect of intronic variants on PE inclusion, we performed a minigene assay containing the whole intron 3 together with its flanking exons. We confirmed the effectivity of such construct in mimicking native splicing using both the wild type sequence and sequence containing either of two known PE-activating variants (22, 23). Specifically, both the variants (1137C>T and 2025T>G, numbered accordingly to their position in intron 3), create *de novo* donor splice sites. In both cases, we detected substantial PE activation (Fig. 2).

**Figure 2:**
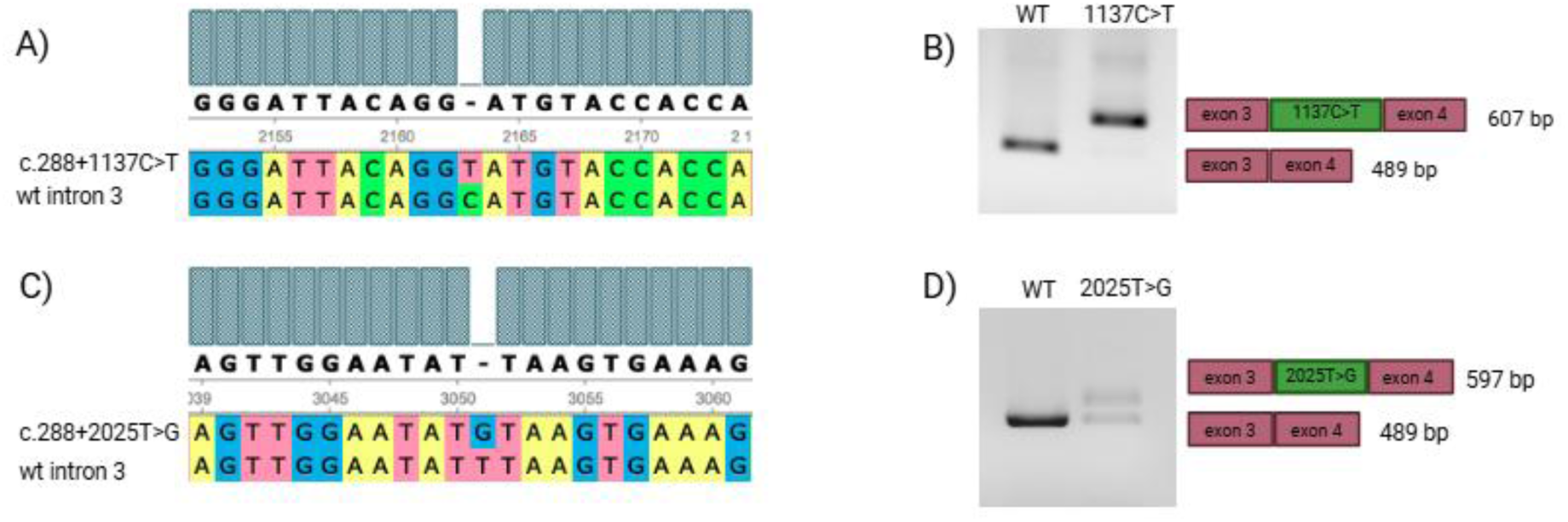
PE-activating variants used as positive controls. A) verification of mutagenesis 1137C>T by Sanger sequencing; B) gel electrophoresis and graphical depiction of PE inclusion after variant introduction; C) and D) stand for variant 2025T>G.

### Deep intronic variants activating the pseudoexon inclusion in intron 3

We next focused on analysing the splicing outcomes of all tested variants. In total, we identified 11 variants that activated PE inclusion in more than 0.5% of their transcripts (Tab. 1). Two of these corresponded to positive controls 1137C>T and 2025T>G, leaving nine variants specifically detected in this study (Fig. 3).

**Figure 3:**
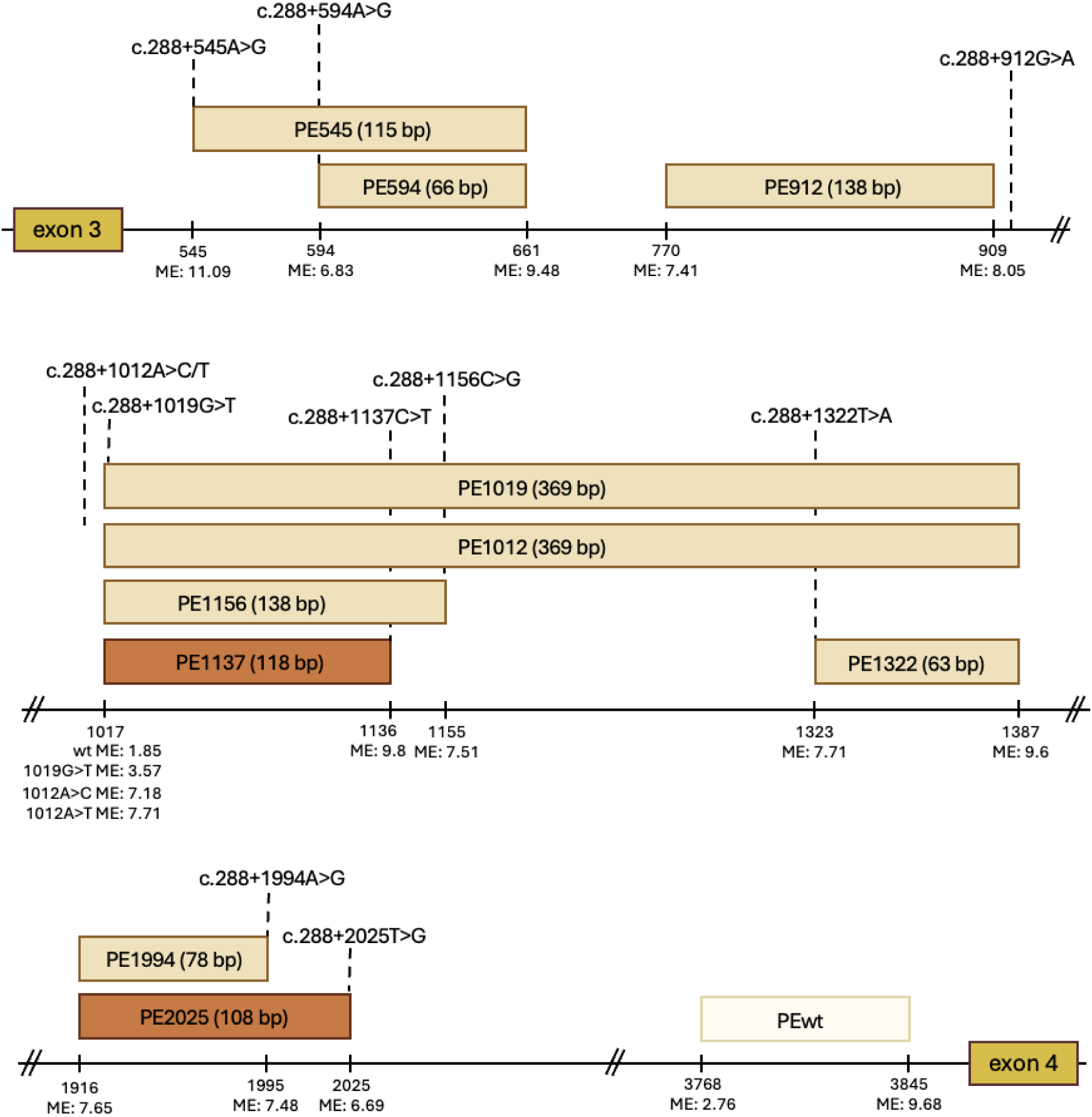
A schematic representation of activated PEs in intron 3 of the NF1 gene. Boxes indicate exons (red), positive control PEs (orange) and newly described PEs (ochre), black line stands for intronic sequence. MaxEnt (ME) scores for each used splice site are depicted. Intronic position of splice sites corresponds with the position of G nucleotide from GT and AG dinucleotide for donors and acceptors, respectively. Dotted line specifies activating variants.

**Table 1.** A list of variants activating the pseudoexon inclusion.

| Variant | Type | PE position | PE inclusion level (NGS) [%] | length [bp] | Presence of PTC | ME score | PE inclusion level (FA) [%] | Notes |
| --- | --- | --- | --- | --- | --- | --- | --- | --- |
| **1137C>T** | DD | [1018-1135] | 52.43 | 118 | yes | 9.8 | 73.1 | P1 |
| **545A>G** | DA | [546-661] | 21.56 | 115 | yes | 11.09 | 48.7 |  |
| **2025T>G** | DD | [1917-2024] | 17.13 | 108 | yes | 6.69 | 41.1 | P2 |
| **1156C>G** | DD | [1018-1154] | 8.90 | 138 | no | 7.51 | 13.9 |  |
| **594A>G** | DA | [595-661] | 4.75 | 66 | no | 6.83 | 19.4 |  |
| **1994A>G** | CD | [1917-1995] | 3.80 | 78 | yes | 7.48 | 21.7 |  |
| **1019G>T** | CA | [1018-1387] | 3.43 | 369 | yes | 3.57 | 4.5 |  |
| **1012A>C** | CA | [1018-1387] | 2.16 | 369 | yes | 7.18 | 1.3 |  |
| **1012A>T** | CA | [1018-1387] | 1.72 | 369 | yes | 7.71 | 3.0 |  |
| **1322T>A** | DA | [1324-1387] | 1.23 | 63 | yes | 7.71 | 3.0 |  |
| **912G>A** | CD | [771-909] | 0.70 | 138 | yes | 8.05 | 3.5 |  |

Note: Variants are sorted according to their PE inclusion levels, which were calculated as a mean value from biological triplicates. DD – *de novo* donor, CD – cryptic donor, DA – *de novo* acceptor, CA – cryptic acceptor. FA – fragment analysis. P1 and P2 – variants used as positive controls.

Apart from the variants selected as positive controls, the most prominent was variant 545A>G. This variant created a strong *de novo* acceptor with a ME score of 11.09, which led to 115 bp-long PE inclusion, using a cryptic donor in position 661 (ME: 9.48). This PE (referred to as PE545) was included in approximately 20% of transcripts, exceeding the level of one of the positive controls (2025, ∼ 17%). A complete list of all variants and their effects on splicing of intron 3 of the *NF1* gene is provided in Tab. S1.

### Low-level inclusion of a wild-type pseudoexon (PEwt)

While analysing our sequencing data, we noticed a recurrent occurrence of a 76 bp exonized sequence at low inclusion levels (average 0.76%). This PEwt is located at position c.288+3769_3844 and utilizes naturally occurring acceptor (ME score 2.76) and donor (ME score 9.68) splice site. The PEwt was activated independently on the introduced variants, and we confirmed its presence with fragment analysis (approximately 1% inclusion) in both mutant and wild-type constructs. To verify that this finding is not an artefact of plasmid-based expression, we performed fragment analysis on HeLa cells that had not been transfected with our plasmid. We obtained comparable results, with an inclusion level of 1%, what further supports the reliability of our minigene assay.

The only exception in PEwt inclusion was library 18, in which this splicing outcome was not present. Upon careful evaluation, we determined that the PEwt, as well as any other PEs, are unfortunately not seen in our sequencing data due to an unintended abolition of the cryptic acceptor splice site 3768 by AsiSI restriction site introduction. Therefore, we manually evaluated selected variants (devoid of any AsiSI restriction sites) using fragment analysis. Notably, variants that strengthened either splice site of this PEwt increased exon inclusion levels (Tab. 2).

**Table 2.** A list of evaluated variants increasing the inclusion level of PEwt.

| Variant | Type | ME score | PEwt inclusion level (FA) [%] | SD [%] |
| --- | --- | --- | --- | --- |
| wt | CA/CD | 2.76/9.68 | 0.99 | 0,07 |
| 3750G>C | CA | 4.03 | 17.0 | 1.33 |
| 3759G>T | CA | 5.13 | 4.6 | 0.58 |
| 3843T>A | CD | 11.08 | 33.0 | 0.44 |
| 3850G>T | CD | 10.65 | 54.6 | 2.5 |

Note: PE inclusion levels were calculated as a mean value from biological triplicates. FA – fragment analysis. CA – cryptic acceptor, CD – cryptic donor.

Following the changes in splicing outcome observed in library 18, we investigated the effect of AsiSI site introduction into the other libraries as well, and any eventual inconsistencies were addressed (see the Supplemental text 2).

### Splicing noise in pseudoexon inclusion

Apart from PEwt and variant-induced changes in splicing, we also observed low-level activation of additional PEs in the wild-type context. The presence of abnormal splicing events detected in only a few reads is commonly observed in RNA-seq data and is usually dismissed as background splicing or splicing noise (5, 24). In our settings, wild-type plasmids (containing AsiSI site) exhibited multiple “pseudoexons” activations, although in a very restricted manner. Particularly, 1018-NA was the only PE that was activated in more than three reads, totally in 0.03% of transcripts across most of the libraries (see Tab. S4). All other PEs were activated in 0.001% at maximum. Although we cannot exclude an effect of an undetected random DNA variant, we believe that this situation could be described as splicing noise, which in our hands did not exceed 0.03%.

### Efficiency of prediction tools for pseudoexon activation

We designed this project with reference to ME scores of individual splice sites, as specified in Methods. In concordance with previous studies, we did not observe a clear relationship between splice-site strength and the PE inclusion level (7, 8), not even among three variants affecting the same splice site (Fig. S1, Tab. S5). We detected only minor differences between activated *de novo* and cryptic splice sites, with slightly higher ME scores for *de novo* sites (Tab. S5).

In contrast, while most of the partner splice sites met the designed ME thresholds, one notable exception was the acceptor at position 1017 (ME score 1.85), substantially below the commonly applied threshold (ME>3) (25). Despite this, the acceptor splice site was repeatedly utilized by multiple activating variants, including the positive control 1137C>T. Notably, a stronger cryptic acceptor (956, ME 8.38) located 61 bp upstream was consistently ignored. This prompted us to examine auxiliary splicing elements in the surrounding region. ESRseq scores (21) indicated presence of a strong silencer element between the two cryptic splice sites, located within the PE body if the stronger site was used (Fig. S2). We hypothesize that this regulatory context may prevent recognition of the stronger acceptor by the splicing machinery. Nowadays, the most widely used *in-silico* prediction tool is SpliceAI (26). We evaluated its predictive accuracy using our validated dataset (n=501). Notably, in most PE-activating cases, Δscores for acceptor or donor gains were present not only at the mutated site but also for its corresponding counterpart.

When applying the recommended Δscore threshold (>0.2) for the mutated site (16, 25, 27), SpliceAI demonstrated high accuracy (98.2%), but with moderate precision (60%) due to nine false-positives (Tab. 3). Requiring a Δscore >0.2 at both splice sites increased precision to 73% but resulted in one additional missed PE activation. By raising the Δscore threshold to >0.5 (the value recommended for clinical use) we further improved precision to 86% but reduced sensitivity to 50% (compared to 75% and 66.7% for the lower-threshold models). Regarding potential clinical relevance, all the three variants leading to more than 10% PE inclusion with Illumina sequencing passed this score threshold. On the other hand, 1156C>G (8, 9 % inclusion) would be missed. Additionally, applying the thresholds recommended by the SpliceAI-10k calculator (28) further enhanced the performance even for low-level activating variants, yielding up to 100% sensitivity and over 96% accuracy (Tab. 3).

**Table 3.**
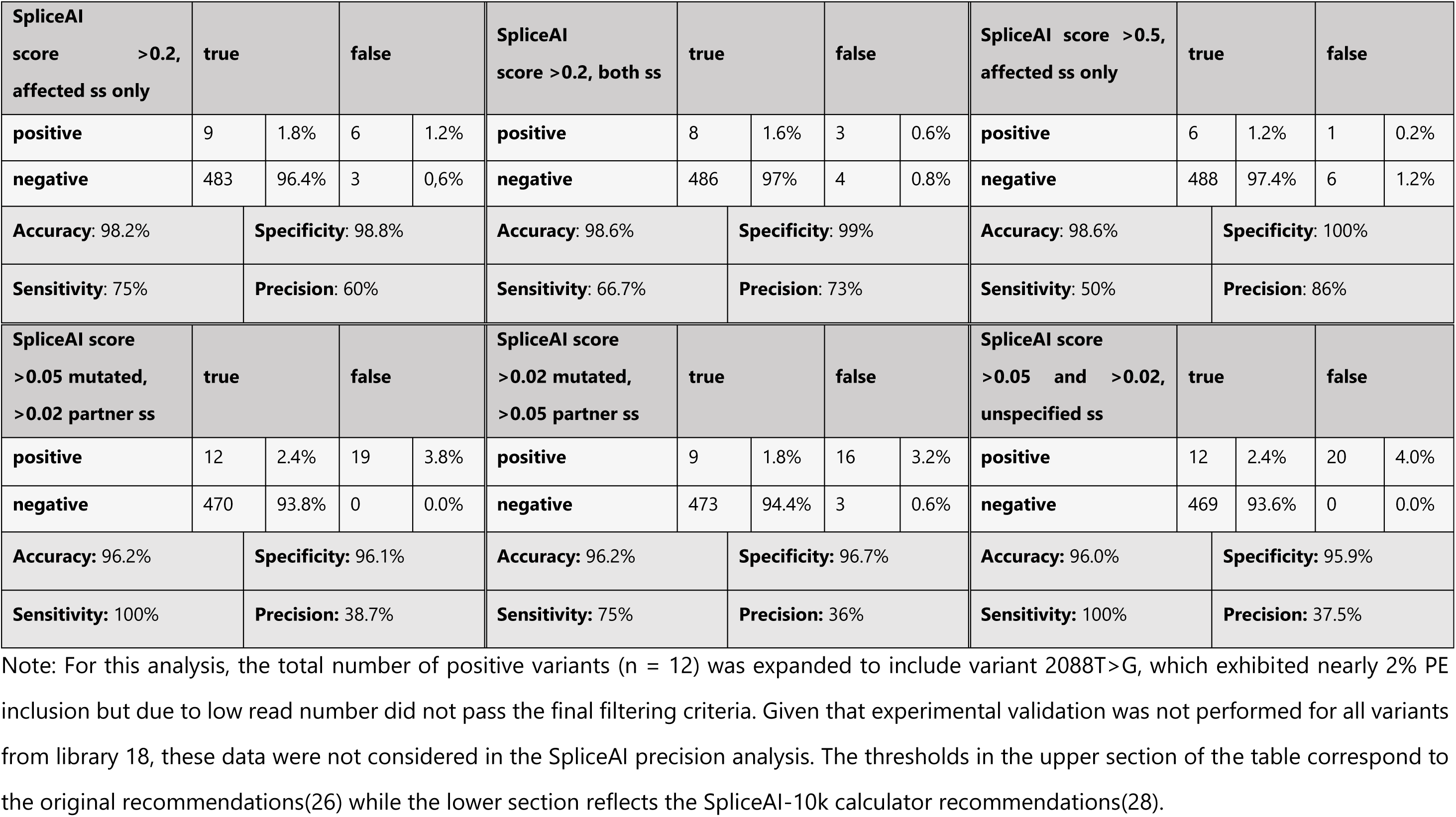
Performance of the SpliceAI prediction tool at different threshold values.

In addition, the SpliceAI prediction score for acceptor gain displayed a correlation with observed PE inclusion levels (Fig. S1), whereas less clear relationships were observed for other score.

### Comparative Analyses of Essential Splicing Signals

To address the effect of SREs on PE activation, we analyzed the auxiliary splicing regulators (ESRseq) score in regions containing activated PEs (Fig. 4). These scores were determined experimentally and reflect the strength of each hexamer, identifying their overall effect on splicing (21).

**Figure 4:**
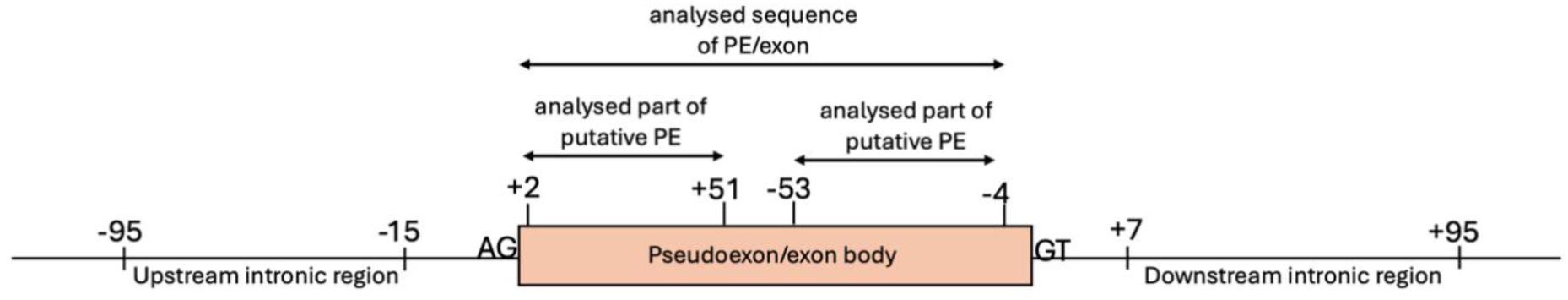
Graphic representation of tested ranges for ESRseq analysis. Following the original suggested testing ranges of Ke *et al.*, 2011 (21), we eliminated the splice sites from our computations. We also removed all the potential duplicities caused by sequence overlaps. For more details, see Methods.

When comparing the ESRseq scores of PEs (n=4) with their flanking intronic regions, PE sequences exhibited higher scores, and therefore greater exonization potential. Importantly, an analogous comparison of *NF1* exons (n=55, with first and last exons excluded) revealed a similar but more pronounced trend, confirming this tendency at least for this genomic context. As expected, PE sequences exhibited lower ESRseq score than true exons, but when compared to intronic sequences flanking the true exons, these scores were significantly discriminating PE sequence from acceptor and donor regions. For both PEs and real exons, the difference was more pronounced for acceptor than for donor regions (Fig. 5A, Tab. S6).

**Figure 5:**
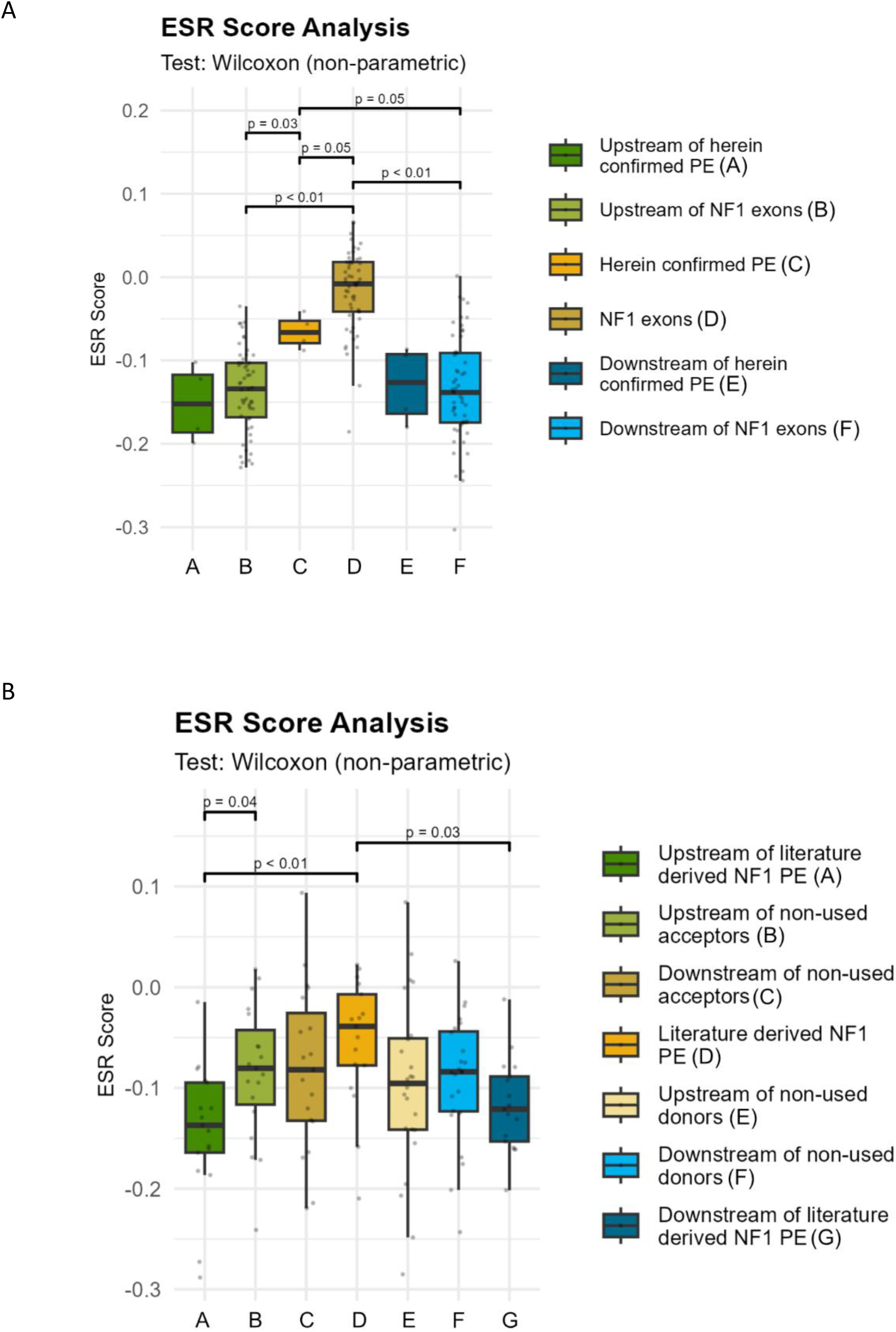
Statistical analysis of ESRseq scores. A) Comparison of PEs validated in this study and exons 2-56 of the *NF1* gene with their flanking intronic sequences. P-value is indicated only in case of statistical significance. The boxplots show medians and interquartile ranges (IQR). Whiskers correspond to the minimum and maximum values within 1.5 times the IQR from the quartiles. B) Comparison of literature derived PEs in the *NF1* gene with negative group of putative PEs.

To further characterize our group of PEs, we compared them with a set of putative PEs. Those were selected from a group of experimentally introduced splice sites not activated in our experiment, as a distance-curated subset of variants outside of activation-prone regions, thus eliminating any potential overlap. Again, we carefully curated the selected regions so each of them was only included once. Because only one splice site position was known, we analysed acceptor (n=19) and donor (n=24) sites separately. When comparing the putative pseudoexonic regions (50 bp downstream/upstream from the splice site) with their flanking introns, we observed no significant difference in exonization potential in terms of ESRseq scores (Fig. S3A). We also observed differences in ESRseq scores between intronic sequences of putative and validated PEs, with the latter exhibiting more negative scores.

Furthermore, we tested a subset of all PEs identified in the *NF1* gene (29), here marked as “literature derived PEs”, following the criteria described above. We observed a significant difference when comparing the ESRseq score of the literature derived PE body with its flanking intronic regions. Comparison with the validated but non-used regions (considered as a negative control group) yielded the same results as observed while using only the smaller set of PEs validated in this study (n=4), but with more pronounced differences (Fig. 5B, Fig. S3B, Tab. S6).

## DISCUSSION

In this study, we employed a minigene assay to perform a high-throughput analysis of more than 500 deep intronic variants within intron 3 of the *NF1* gene that were predicted to alter splicing and promote PE inclusion. Minigene assay was previously utilized to study deep intronic variants (30–33) but, to our knowledge, this is the first time it has been used for their high-throughput analysis. Despite the known limitations (34, 35), minigene-based approach enabled controlled analysis independent of patient-derived RNA, improved sensitivity for low-frequency events, and facilitated manipulation of long intronic regions. The inclusion of two previously reported variants (22, 23) as positive controls confirmed the robustness of this approach.

In addition to these two positive controls, previously reported in patients with NF1, we identified nine variants capable of activating PE inclusion (threshold ≥0.5%). Two of those variants are listed in the ClinVar database lacking published experimental validation (variants 545 and 594; variant 594 described with neurofibromatosis type 1 (NF1) condition), and the remaining seven are described herein for the first time. None of these variants is present in the gnomAD population database (version 4.1).

PE inclusion has potential clinical relevance, especially when introducing PTCs. One variant (545A>G) activated PE inclusion in approximately 21% of transcripts (nearly 50% in fragment analysis) exceeding the inclusion levels observed for the positive control 2025T>G (previously described in patient with NF1 (22) and generated transcripts containing PTCs, likely triggering nonsense-mediated decay and supporting its classification as likely pathogenic. In contrast, variants producing in-frame insertions (e.g., 1156C>G and 594A>G) remain difficult to interpret, as protein modelling did not predict substantial alterations in protein folding (data can be sent upon request). Variants with low inclusion (<5%) present an additional challenge, as the threshold for pathogenic transcript reduction in NF1 remains undefined. Nevertheless, all activating variants provide insight into splicing regulation.

Observed discrepancies between NGS and fragment analysis likely reflect inherent methodological differences. NGS libraries included a proportion of wild-type reads, whereas fragment analysis assessed fully variant-specific constructs, leading to higher inclusion estimates. We therefore consider fragment analysis more representative of quantitative splicing outcomes, particularly for variants with potential clinical relevance.

In agreement with observations that the same PEs can be activated by multiple variants (7), the 11 activating variants repeatedly preferred the same five partner splice sites, suggesting that certain cryptic splice sites and certain intronic regions are intrinsically more susceptible to activation. Clustering of activating variants within four locations (Fig. 3) may indicate the vulnerability of these regions to even more subtle changes, such as auxiliary regulators variants, what we accidently showed in library 5 (Supplemental Text 2). These activation-prone “hot spots” represent high-priority targets for genomic diagnostics, particularly in patients with a clinical diagnosis but no causative variant identified in coding regions or canonical splice sites.

Furthermore, the distribution of aberrant splicing events in wild type settings was non-random (Tab. S4). Instead, these background events co-localized with the same activation-prone hotspots identified in our screen. We hypothesize that such trace-level background splicing, when detected repeatedly, reflects a sensitized genetic background where deep intronic variants are more likely to trigger pathogenic splicing shifts. Identifying these sensitized regions could significantly improve the prioritization and detection of deep intronic variation in clinical diagnostics.

In addition to variants-induced PEs, we identified a low-level wild-type pseudoexon (PEwt; c.288+3769_3844) that was present independently of introduced variants and detectable in untransfected HeLa cells. Unlike prior reports (SpliceVault online tool (36)), suggesting partial intron retention, our data indicate coordinated usage of both splice sites, supporting genuine exonization. We showed that variants strengthening the associated splice sites substantially increase PEwt inclusion (∼55% for variant 3850TG>T), which, together with the frameshift introduction, may suggest potential clinical relevance. This region may therefore represent an additional splicing-sensitive hotspot.

High-throughput approach is invaluable for rapid and efficient analysis of large datasets, but it is also sensitive to subtle sequence changes. Introduction of AsiSI restriction sites occasionally altered splicing, either by disrupting cryptic splice sites (library 18) or by facilitating PE activation in combination with otherwise neutral variants (libraries 5 and 8). These findings highlight the importance of sequence context and demonstrate how even minor modifications can influence the splicing outcome.

In recent years, the SpliceAI predictor (26) has emerged as a powerful tool widely used in both clinical practice and research. It outperforms other *in silico* prediction tools in its specificity and sensitivity and has shown promising performance in predicting PE inclusion (16). Recent discussions have underscored the lack of a universal SpliceAI threshold, with recommended thresholds ranging from 0.5 to as low as 0.02 score for predicting PEs inclusion (16, 28). Using the generally recommended Δscore >0.2, we observed three false negative and six false positive predictions. Requiring a Δscore >0.2 at both partner splice sites (25) reduced the number of false positives to three while increasing false negatives to four. Raising the threshold to the clinically recommended Δscore >0.5 further improved precision at the expense of sensitivity, resulting in six false negatives. Applying the SpliceAI-10k calculator recommendations (28) yielded up to 100% sensitivity, depending on whether the Δscore >0.02 or >0.05 was applied to the mutated or partner splice site. This was offset by lower precision, given the increased number of false positives (Tab. 3). However, all our positive variants were detected, making this approach the most sensitive option in clinical practice.

Notably, our classification was based on an arbitrary 0.5% PE inclusion threshold, increasing the likelihood that most of the false negatives may be clinically irrelevant. However, applying the Δscore >0.5 threshold would miss one of the variants with a rather high inclusion level (1156C>G, 8.9% by NGS). These findings highlight the need for context-aware interpretation.

Beyond splice-site strength, we also assessed splicing regulatory elements using ESRseq scores (21). Despite the limited sample size, ESRseq scores clearly distinguished between exonic regions from flanking intronic sequences in both *NF1* exons and PEs, including those confirmed here and reported previously (29), whereas no such distinction was observed for non-activated putative pseudoexons. These findings suggest that ESRseq scores can estimate the exonization potential of intronic regions under certain conditions, a conclusion supported by similar observations for PE-activating deep intronic variants in genes associated with hereditary breast and ovarian cancer in both germline and somatic settings (37). Together, these results extend the relevance of ESRseq-based prioritization beyond NF1 and support its potential utility in diagnostic settings.

With this study, we aimed to contribute to the challenging and labour-intensive process of annotating and identifying regions susceptible to aberrant splicing. Such datasets can support prioritization of deep intronic variants of uncertain significance identified by genome sequencing and improve diagnostic interpretation. Moreover, clinically relevant PEs represent attractive targets for personalized antisense-based therapy. Their deep intronic location enables the delivery of sequence-specific antisense molecules without interfering with canonical splicing (5, 37), further emphasizing the clinical importance of identifying and characterizing deep intronic variants, their effect on splicing, and their potential treatability.

## ACKNOWLEDGEMENTS

We would like to thank to Filip Pardy for technical support with sequencing. NGS data were obtained in collaboration with CF Genomics CEITEC MU.

## AUTHOR CONTRIBUTIONS

Eliška Svobodová: Investigation, Data curation, Formal analysis, Methodology, Writing – original draft, Writing – review & editing. Lucie Kajan Grodecká: Supervision, Conceptualization, Data curation, Formal analysis, Writing – review & editing. Přemysl Souček: Methodology, Formal analysis, Writing – review & editing. Kamila Réblová: Formal analysis. Zdeněk Kubát: Investigation. Marie Krátká: Formal analysis. Tomáš Freiberger: Supervision, Funding acquisition, Writing – review & editing. All authors have read and agreed to the published version of the manuscript.

## SUPPLEMENTARY DATA

Supplementary Data are available online.

## DECLARATION OF INTERESTS

The authors declare no competing interests.

## FUNDING

This work was supported by the Czech Science Foundation [GF24-11219K]. Conceptual development of research organization was provided by the Ministry of Health of the Czech Republic [FNBr 65269705]. Computational resources were provided by the e-INFRA CZ project [ID:90254], supported by the Ministry of Education, Youth and Sports of the Czech Republic and ELIXIR-CZ project [ID:90255], part of the international ELIXIR infrastructure.

## DATA AVAILABILITY

All sequencing data, code and primer sequences are available at Zenodo https://doi.org/10.5281/zenodo.20744004.

## DECLARATION OF GENERATIVE AI AND AI-ASSISTED TECHNOLOGIES IN THE MANUSCRIPT PREPARATION PROCESS

During the preparation of this work the authors used Google Gemini in order to assist with R script development and language editing. The author(s) reviewed and edited the output as needed and take full responsibility for the content of the published article.

## Notes

https://doi.org/10.5281/zenodo.20744004

### Competing Interest Statement

The authors have declared no competing interest.

