## Supplemental material for "Decoding pseudoexon activation by deep intronic variants using a high-throughput minigene assay in the *NF1* gene"

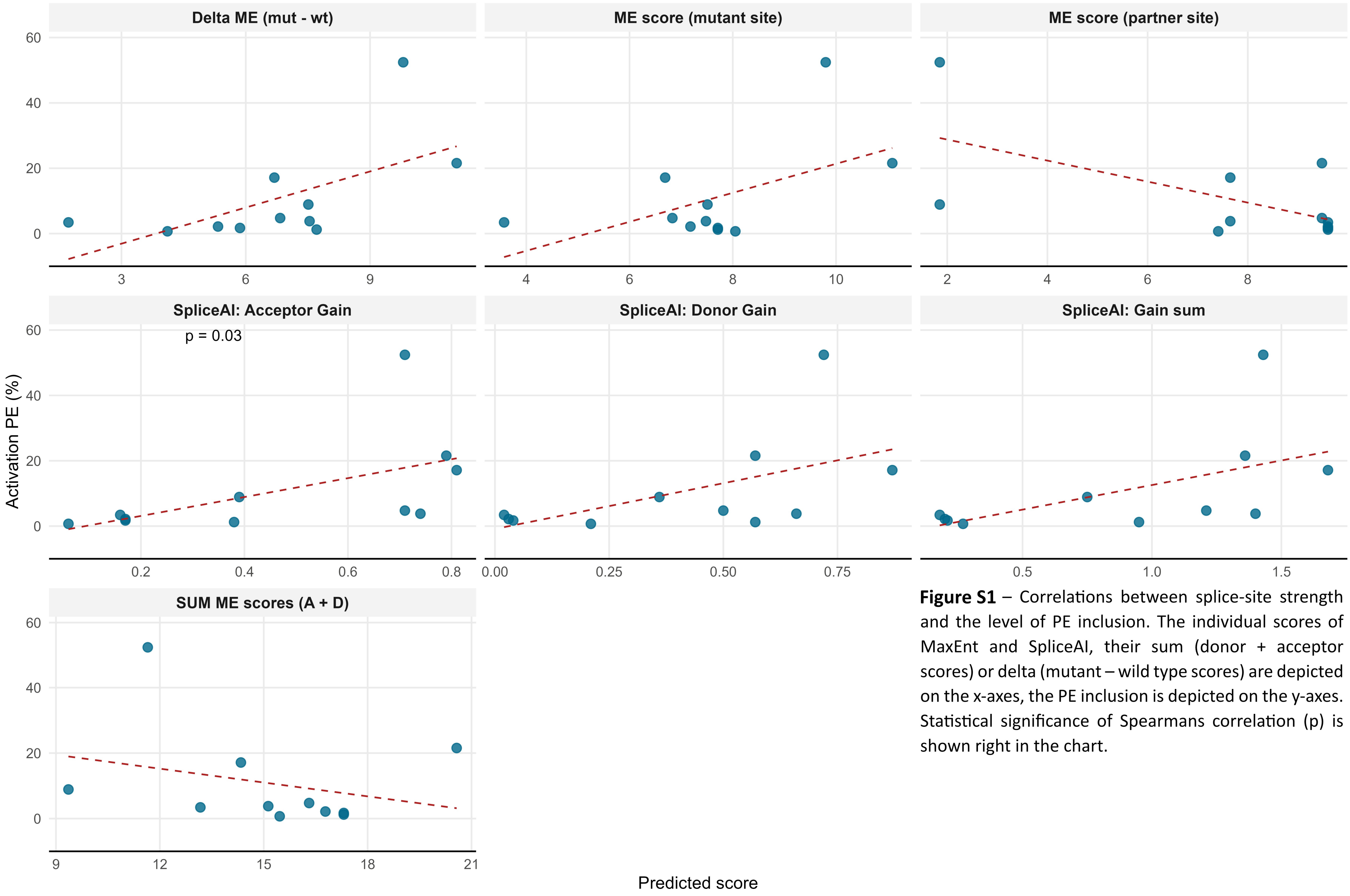

**Figure S1** – Correlations between splice-site strength and the level of PE inclusion. The individual scores of MaxEnt and SpliceAI, their sum (donor + acceptor scores) or delta (mutant – wild type scores) are depicted on the x-axes, the PE inclusion is depicted on the y-axes. Statistical significance of Spearmans correlation (p) is shown right in the chart.

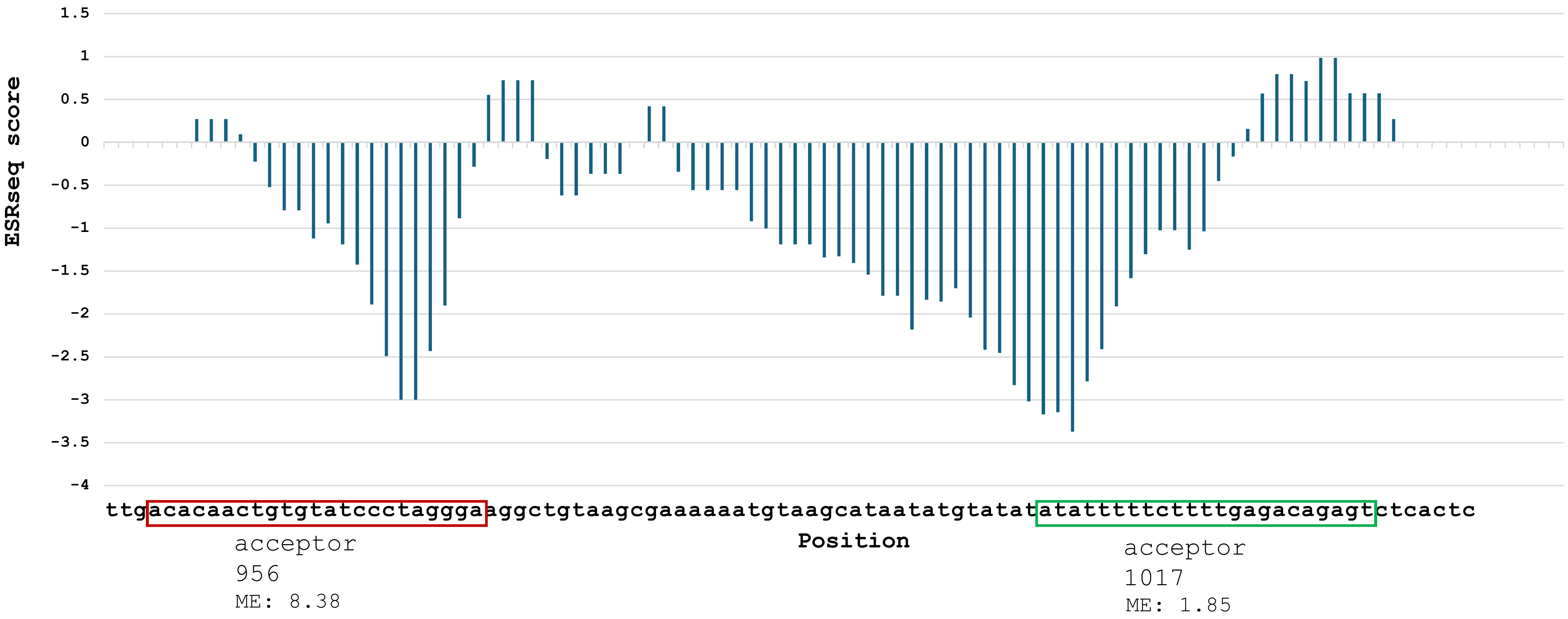

**Figure S2** – graphical representation of ESRseq score for each hexamer in intronic region of used acceptor 1017 and unused acceptor 956.

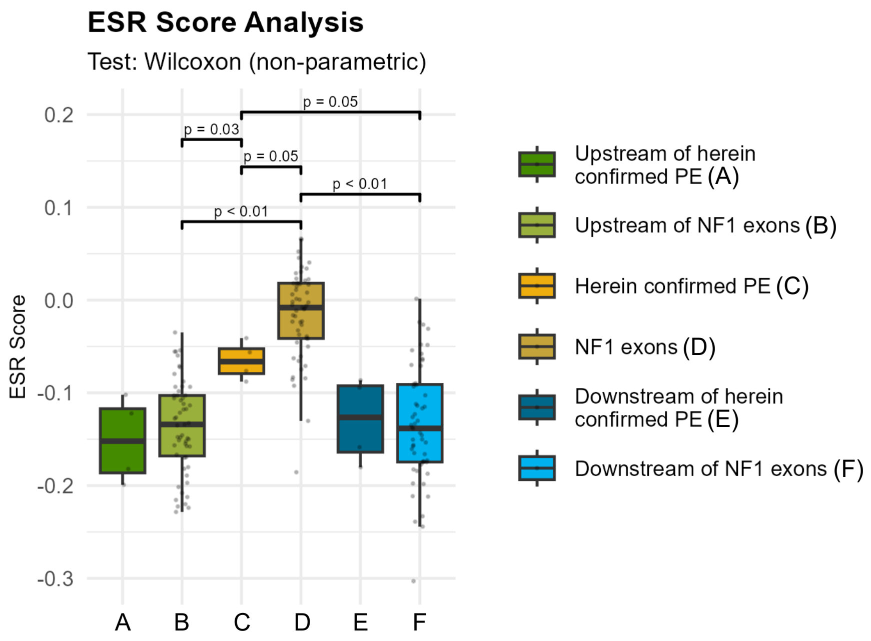

**Figure S3A** – Statistical analysis of ESRseq scores. Comparison of PEs validated in this study and negative group of non-used splice sites. The boxplots show medians and interquartile ranges (IQR). Whiskers correspond to the minimum and maximum values within 1.5 times the IQR from the quartiles.

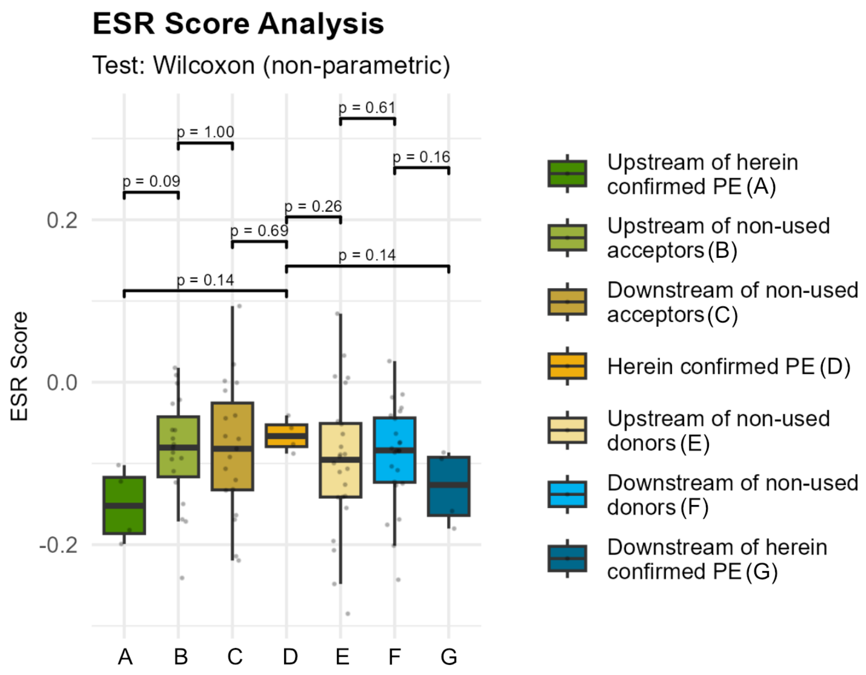

**Figure S3B** – Statistical analysis of ESRseq scores. Comparison of literature derived PEs in the NF1 gene with exons 2-56 of the NF1 gene with their flanking intronic sequences. The boxplots show medians and interquartile ranges (IQR). Whiskers correspond to the minimum and maximum values within 1.5 times the IQR from the quartiles.

**Supplementary text 1**

***Nanopore sequencing and data processing***

Plasmid libraries were linearized using the PstI restriction enzyme and purified by ethanol precipitation. Sequencing libraries were prepared using the Ligation Sequencing Kit V14 (SQK-LSK114, Oxford Nanopore Technologies) according to the manufacturer's protocol, starting with 100 fmol of DNA per library for adapter ligation. For each run, two libraries were pooled and loaded onto a Flongle Flow Cell (FLO-FLG114). Sequencing was performed on a MinION Mk-1B device (Oxford Nanopore Technologies) for 24 hours.

Raw Nanopore sequencing data were basecalled using Dorado v0.9.1 (Oxford Nanopore Technologie) in duplex mode with the SUP (super-accuracy) model (dna_r10.4.1_e8.2_400bps_sup@v5.0.0), using the stereo duplex model (dna_r10.4.1_e8.2_5khz_stereo@v1.3) for duplex pairing. Reads were aligned to a custom plasmid reference sequence during basecalling. The resulting BAM file was sorted and indexed using SAMtools v1.21 [1]. Quality metrics were assessed using NanoPlot with a maximum read length filter of 20,000 bp [2]. Variants were called with custom script, using SAM tools.

After basecalling and quality checking, sequencing libraries were demultiplexed by scoring each read against library-specific restriction site sequences (AsiSI sites) introduced at defined positions within the minigene construct. For each library, a scoring function was applied: reads matching the restriction site sequence at variable positions received a positive score (+1 per matching base), while reads matching the original reference sequence received a negative score (−1 per matching base). Reads with a positive net score were assigned to the corresponding library.

Barcodes were extracted by hard-clipping reads to the 6 bp barcode region (positions 482–488 of the reference) using samtools ampliconclip with --both-ends and --hard-clip flags. Barcode quality scores were calculated as a composite Phred-based probability of all bases being correctly called. The barcode sequence CTAGCT was excluded as a no-template control. Barcodes with fewer than 6 recovered bases were discarded.

Variants were called by iterating over each read aligned to the reference and querying the base at each variant position defined in a VCF file. Reads were classified as carrying the alternative allele (1), reference allele (0), or an unexpected base (9) at each position. Reads with no coverage at a given position (e.g., due to deletions or reference skips) were assigned a missing value (−9). Per-barcode variant tables were generated separately for each library, reporting mutated read depth, total sequencing depth, and probabilistic support scores for mutation presence and absence, derived from per-base quality values.

**Supplementary text 2**

***Splicing influenced by AsiSI sites addition***

Following the splicing changes observed after insertion of the *AsiSI* restriction site in library 18, we investigated the effect of introducing this restriction site into other libraries. Although experimentally tested during project design, combined effect of the AsiSI sequences and introduced mutations cannot be excluded. Closer inspection revealed that the sequence modification affected several cryptic splice sites, predominantly acceptors. We assessed each affected site, in terms of predicted strength (Tab. S7), and although no significant effects were expected, we considered the potential impact on splicing during subsequent data analysis.

In library 8, we observed an occurrence of 140 bp PE in position [1534-1673], with the most prominent activation in variant 1677T>G (29.8%). However, this was not confirmed in fragment analysis (using a plasmid without any *AsiSI* restriction site). Moreover, this PE activation was not detected in any other library. In fact, this AsiSi introduction possibly substituted naturally present binding site for the splicing silencer MBNL1 with binding site for the enhancer SRp20, as suggested by the SpliceAID2 database [1], what may have resulted in changed splicing environment leading to false positive results. Since PE inclusion was not confirmed in fragment analysis, all variants activating this PE were labelled as false positive and excluded from bioinformatic tools assessment.

A more complex situation of the *AsiSI* restriction site was observed in library 5. The substitution left the used splice site sequence unaffected but influenced the balance of SREs to be more splicing-supporting. We detected an increase in activation of the [1018-NA] inclusion across the entire library 5 (Tab. S4). Yet, not all of these events could be classified as false positive and excluded from the analysis. Several of them were confirmed by fragmentation analysis, including one of the positive controls.

**Table S1 -** A list of variants selected based on their ME score for pseudoexon inclusion study. Chromosomal position is using the hg19 human coordinates. DD -*de novo* donor, DA - *de novo* acceptor, CD - cryptic donor, CA - cryptic acceptor. The effect on splicing is assessed according to the rules applied in the article (>3 reads, >0.5% inclusion level). The "characteristics" column details the position of the splice site dinucleotide and the associated mutation. For *de novo* splice sites( e.g. TT>GT:58) the number denotes the second nucleotide of the dinucleotide (spanning intronic positions 57–58). For cryptic sites (e.g., 70-68A>G), the first number indicates the end of the dinucleotide (spanning positions 69–70), while the second value (e.g., -68) specifies the position of the mutation.

**Library 1**

| intronic position | type | chrom. position | characteristics | ref | alt | MaxEnt score | SpliceAI prediction | | | | acquired | effect on splicing |
|---|---|---|---|---|---|---|---|---|---|---|---|---|
| | | | | | | | acceptor gain | acceptor loss | donor gain | donor loss | | |
| 57 | DD | 29486168 | TT>GT:58 | T | G | 5,94 | 0.00 | 0.00 | 0.00 | 0.00 | no | N/A |
| 67 | DA | 29486178 | AA>AG:67 | A | G | 4,68 | 0.00 | 0.00 | 0.00 | 0.00 | yes | none |
| 68 | DA | 29486179 | AA>AG:68 | A | G | 5,79 | 0.00 | 0.00 | 0.00 | 0.01 | yes | none |
| 68 | CD | 29486179 | 70-68A>G | A | G | 8,04 | 0.00 | 0.00 | 0.00 | 0.00 | yes | none |
| 75 | DD | 29486186 | AT>GT:76 | A | G | 6,34 | 0.00 | 0.00 | 0.00 | 0.01 | yes | none |
| 90 | DD | 29486201 | GA>GT:90 | A | T | 7,7 | 0.00 | 0.00 | 0.00 | 0.00 | yes | none |
| 111 | DA | 29486222 | TG>AG:112 | T | A | 3,46 | 0.00 | 0.00 | 0.00 | 0.00 | yes | none |
| 125 | DD | 29486236 | AT>GT:126 | A | G | 7,8 | 0.05 | 0.00 | 0.14 | 0.02 | yes | none |
| 128 | CA | 29486239 | 143-128A>G | A | G | 4,34 | 0.00 | 0.00 | 0.00 | 0.00 | yes | none |
| 128 | CA | 29486239 | 143-128A>T | A | T | 5,75 | 0.00 | 0.00 | 0.00 | 0.00 | yes | none |
| 129 | CA | 29486240 | 143-129G>T | G | T | 4,92 | 0.00 | 0.00 | 0.00 | 0.00 | yes | none |
| 130 | DD | 29486241 | TT>GT:131 | T | G | 6,36 | 0.01 | 0.00 | 0.03 | 0.01 | yes | none |
| 133 | CA | 29486244 | 143-133T>C | T | C | 3,53 | 0.00 | 0.00 | 0.00 | 0.00 | yes | none |
| 156 | CD | 29486267 | 153-156C>G | C | G | 7,54 | 0.00 | 0.00 | 0.00 | 0.00 | yes | none |
| 191 | DA | 29486302 | TG>AG:192 | T | A | 5,58 | 0.00 | 0.00 | 0.00 | 0.00 | yes | none |
| 195 | DA | 29486306 | AA>AG:195 | A | G | 9,06 | 0.00 | 0.00 | 0.00 | 0.00 | yes | none |
| 196 | DA | 29486307 | AA>AG:196 | A | G | 4,9 | 0.00 | 0.00 | 0.00 | 0.00 | yes | none |
| 196 | CD | 29486307 | 193-196A>G | A | G | 7,4 | 0.00 | 0.00 | 0.00 | 0.00 | yes | none |
| 200 | DA | 29486311 | AA>AG:200 | A | G | 4,77 | 0.00 | 0.00 | 0.00 | 0.00 | yes | none |
| 205 | DD | 29486316 | AT>GT:206 | A | G | 7,26 | 0.00 | 0.00 | 0.00 | 0.00 | yes | none |
| 208 | DD | 29486319 | TT>GT:209 | T | G | 8,65 | 0.00 | 0.00 | 0.00 | 0.00 | yes | none |
| 228 | DD | 29486339 | TT>GT:229 | T | G | 8,35 | 0.00 | 0.00 | 0.00 | 0.00 | yes | none |
| 231 | DA | 29486342 | AT>AG:231 | T | G | 4,25 | 0.00 | 0.00 | 0.00 | 0.00 | yes | none |
| 242 | CD | 29486353 | 244-242C>G | C | G | 7,33 | 0.00 | 0.00 | 0.00 | 0.00 | yes | none |
| 246 | CD | 29486357 | 244-246T>A | T | A | 5,9 | 0.00 | 0.00 | 0.00 | 0.00 | yes | none |
| 247 | CD | 29486358 | 244-247C>G | C | G | 7,26 | 0.00 | 0.00 | 0.00 | 0.00 | yes | none |
| 256 | CA | 29486367 | 265-256A>G | A | G | 10,38 | 0.08 | 0.00 | 0.01 | 0.00 | yes | none |
| 257 | CA | 29486368 | 265-257G>C | G | C | 12,31 | 0.29 | 0.00 | 0.08 | 0.00 | yes | none |

**Library 1** **continued**

| intronic position | type | chrom. position | characteristics | ref | alt | MaxEnt score | SpliceAI prediction | | | | acquired | effect on splicing |
|---|---|---|---|---|---|---|---|---|---|---|---|---|
| | | | | | | | acceptor gain | acceptor loss | donor gain | donor loss | | |
| 258 | CA | 29486369 | 265-258G>C | G | C | 5,64 | 0.01 | 0.00 | 0.00 | 0.00 | yes | none |
| 257 | DA | 29486368 | GG>AG:258 | G | A | 4,68 | 0.02 | 0.00 | 0.00 | 0.00 | yes | none |
| 260 | CD | 29486371 | 259-260T>A | T | A | 7,51 | 0.00 | 0.00 | 0.00 | 0.00 | insufficient read depth | none |
| 262 | CD | 29486373 | 259-262T>G | T | G | 7,44 | 0.01 | 0.00 | 0.00 | 0.00 | yes | none |
| 267 | DD | 29486378 | GA>GT:267 | A | T | 6,62 | 0.00 | 0.00 | 0.00 | 0.00 | yes | none |
| 268 | CA | 29486379 | 265-268A>C | A | C | 4,83 | 0.01 | 0.00 | 0.01 | 0.00 | yes | none |
| 278 | DA | 29486389 | AT>AG:278 | T | G | 4,06 | 0.00 | 0.00 | 0.00 | 0.00 | yes | none |

**Library 2**

| intronic position | type | chrom. position | characteristics | ref | alt | MaxEnt score | SpliceAI prediction | | | | acquired | effect on splicing |
|---|---|---|---|---|---|---|---|---|---|---|---|---|
| | | | | | | | acceptor gain | acceptor loss | donor gain | donor loss | | |
| 303 | CA | 29486414 | 313-303G>C | G | C | 5,62 | 0.00 | 0.00 | 0.00 | 0.00 | yes | none |
| 302 | DA | 29486413 | CG>AG:303 | C | A | 5,37 | 0.00 | 0.00 | 0.00 | 0.00 | yes | none |
| 304 | CA | 29486415 | 313-304G>C | G | C | 4,78 | 0.00 | 0.00 | 0.00 | 0.00 | yes | none |
| 304 | CA | 29486415 | 313-304G>T | G | T | 6,64 | 0.00 | 0.00 | 0.00 | 0.00 | yes | none |
| 303 | DA | 29486414 | GG>AG:304 | G | A | 7,76 | 0.01 | 0.00 | 0.00 | 0.00 | yes | none |
| 305 | CA | 29486416 | 313-305A>C | A | C | 4,18 | 0.00 | 0.00 | 0.00 | 0.00 | yes | none |
| 314 | DD | 29486425 | GA>GT:314 | A | T | 7,68 | 0.00 | 0.00 | 0.01 | 0.00 | no | N/A |
| 325 | DD | 29486436 | GA>GT:325 | A | T | 8,73 | 0.29 | 0.00 | 0.48 | 0.00 | no | N/A |
| 353 | CD | 29486464 | 350-353T>G | T | G | 6,31 | 0.46 | 0.00 | 0.64 | 0.00 | yes | none |
| 370 | DA | 29486481 | AT>AG:370 | T | G | 8,26 | 0.00 | 0.00 | 0.00 | 0.00 | yes | none |
| 372 | DA | 29486483 | AT>AG:372 | T | G | 10,89 | 0.00 | 0.00 | 0.00 | 0.00 | yes | none |
| 372 | DA | 29486483 | TG>AG:373 | T | A | 4,63 | 0.00 | 0.00 | 0.00 | 0.00 | yes | none |
| 376 | DA | 29486487 | AT>AG:376 | T | G | 4,47 | 0.01 | 0.00 | 0.00 | 0.00 | yes | none |
| 378 | DA | 29486489 | AT>AG:378 | T | G | 7,81 | 0.00 | 0.00 | 0.00 | 0.00 | no | N/A |
| 394 | DA | 29486505 | AC>AG:394 | C | G | 6,22 | 0.00 | 0.00 | 0.00 | 0.00 | yes | none |
| 398 | DA | 29486509 | TG>AG:399 | T | A | 7,9 | 0.00 | 0.00 | 0.00 | 0.00 | yes | none |
| 402 | DA | 29486513 | AA>AG:402 | A | G | 5,82 | 0.00 | 0.00 | 0.00 | 0.00 | no | N/A |
| 404 | DD | 29486515 | TT>GT:405 | T | G | 5,87 | 0.00 | 0.00 | 0.01 | 0.00 | yes | none |

**Library 2** **continued**

| intronic position | type | chrom. position | characteristics | ref | alt | MaxEnt score | SpliceAI prediction: acceptor gain | acceptor loss | donor gain | donor loss | acquired | effect on splicing |
|---|---|---|---|---|---|---|---|---|---|---|---|---|
| 407 | DA | 29486518 | AT>AG:407 | T | G | 7,17 | 0.01 | 0.00 | 0.01 | 0.00 | no | N/A |
| 407 | CD | 29486518 | 409-407T>G | T | G | 6,62 | 0.01 | 0.00 | 0.01 | 0.00 | | |
| 411 | DA | 29486522 | AA>AG:411 | A | G | 4,48 | 0.00 | 0.00 | 0.00 | 0.00 | no | N/A |
| 412 | CD | 29486523 | 409-412A>G | A | G | 6,69 | 0.02 | 0.00 | 0.07 | 0.00 | yes | none |
| 423 | CA | 29486534 | 440-423A>T | A | T | 3,64 | 0.00 | 0.00 | 0.00 | 0.00 | yes | none |
| 429 | CA | 29486540 | 440-429A>T | A | T | 4,49 | 0.00 | 0.00 | 0.00 | 0.00 | yes | none |
| 431 | CA | 29486542 | 440-431A>C | A | C | 3,4 | 0.00 | 0.00 | 0.00 | 0.00 | yes | none |
| 438 | CA | 29486549 | 440-438A>T | A | T | 5,71 | 0.00 | 0.00 | 0.00 | 0.00 | yes | none |
| 442 | DD | 29486553 | GA>GT:442 | A | T | 6,12 | 0.00 | 0.00 | 0.00 | 0.00 | yes | none |
| 493 | CD | 29486604 | 497-493T>G | T | G | 9,81 | 0.00 | 0.00 | 0.01 | 0.00 | yes | none |
| 493 | CD | 29486604 | 497-493T>C | T | C | 9,8 | 0.00 | 0.00 | 0.00 | 0.00 | yes | none |
| 493 | CD | 29486604 | 497-493T>A | T | A | 9,79 | 0.00 | 0.00 | 0.00 | 0.00 | yes | none |
| 492 | DD | 29486603 | AT>GT:493 | A | G | 9 | 0.00 | 0.00 | 0.00 | 0.00 | yes | none |
| 494 | CD | 29486605 | 497-494A>C | A | C | 9,22 | 0.03 | 0.00 | 0.03 | 0.00 | no | N/A |

**Library 3**

| intronic position | type | chrom. position | characteristics | ref | alt | MaxEnt score | SpliceAI prediction: acceptor gain | acceptor loss | donor gain | donor loss | acquired | effect on splicing |
|---|---|---|---|---|---|---|---|---|---|---|---|---|
| 499 | CD | 29486610 | 497-499T>A | T | A | 9,66 | 0.00 | 0.00 | 0.00 | 0.00 | no | N/A |
| 499 | CD | 29486610 | 497-499T>C | T | C | 8,17 | 0.00 | 0.00 | 0.00 | 0.00 | no | N/A |
| 499 | CD | 29486610 | 497-499T>G | T | G | 7,92 | 0.00 | 0.00 | 0.00 | 0.00 | no | N/A |
| 510 | DD | 29486621 | GA>GT:510 | A | T | 10,03 | 0.00 | 0.00 | 0.01 | 0.00 | yes | none |
| 533 | DA | 29486644 | TG>AG:534 | T | A | 6,69 | 0.00 | 0.00 | 0.00 | 0.00 | yes | none |
| 534 | DA | 29486645 | GG>AG:535 | G | A | 4,33 | 0.00 | 0.00 | 0.00 | 0.00 | yes | none |
| 539 | CD | 29486650 | 543-539C>A | C | A | 8,59 | 0.00 | 0.00 | 0.00 | 0.00 | yes | none |
| 539 | CD | 29486650 | 543-539C>G | C | G | 8,56 | 0.00 | 0.00 | 0.00 | 0.00 | yes | none |
| 539 | CD | 29486650 | 543-539C>T | C | T | 7,96 | 0.00 | 0.00 | 0.00 | 0.00 | yes | none |
| 540 | CD | 29486651 | 543-540C>G | C | G | 8,58 | 0.00 | 0.00 | 0.00 | 0.00 | yes | none |
| 540 | CD | 29486651 | 543-540C>A | C | A | 8,31 | 0.00 | 0.00 | 0.00 | 0.00 | yes | none |
| 540 | CD | 29486651 | 543-540C>T | C | T | 8 | 0.00 | 0.00 | 0.00 | 0.00 | no | N/A |
| 541 | CD | 29486652 | 543-541T>G | T | G | 10,91 | 0.01 | 0.00 | 0.05 | 0.00 | no | N/A |

**Library 3** **continued**

| intronic position | type | chrom. position | characteristics | ref | alt | MaxEnt score | SpliceAI prediction acceptor gain | acceptor loss | donor gain | donor loss | acquired | effect on splicing |
|---|---|---|---|---|---|---|---|---|---|---|---|---|
| 541 | CD | 29486652 | 543-541T>A | T | A | 9,09 | 0.00 | 0.00 | 0.00 | 0.00 | no | N/A |
| 541 | CD | 29486652 | 543-541T>C | T | C | 9,09 | 0.00 | 0.00 | 0.00 | 0.00 | yes | none |
| 541 | DA | 29486652 | TG>AG:542 | T | A | 6,3 | 0.00 | 0.00 | 0.00 | 0.00 | no | N/A |
| 544 | CA | 29486655 | 546-544A>T | A | T | 8,16 | 0.01 | 0.00 | 0.00 | 0.00 | yes | none |
| 544 | CA | 29486655 | 546-544A>C | A | C | 9,37 | 0.02 | 0.00 | 0.00 | 0.00 | yes | none |
| 545 | DA | 29486656 | AA>AG:545 | A | G | 11,09 | 0.79 | 0.00 | 0.57 | 0.00 | yes | PE545 - 21.5% inclusion |
| 549 | CA | 29486660 | 546-549G>A | G | A | 5,78 | 0.00 | 0.00 | 0.00 | 0.00 | yes | none |
| 549 | CA | 29486660 | 546-549G>T | G | T | 6,39 | 0.00 | 0.00 | 0.00 | 0.00 | yes | none |
| 572 | DA | 29486683 | AT>AG:572 | T | G | 4,8 | 0.00 | 0.00 | 0.01 | 0.00 | no | N/A |
| 578 | CA | 29486689 | 595-578A>T | A | T | 3,45 | 0.00 | 0.00 | 0.00 | 0.00 | no | N/A |
| 593 | CA | 29486704 | 595-593A>T | A | T | 6,18 | 0.08 | 0.00 | 0.01 | 0.00 | no | N/A |
| 594 | DA | 29486705 | AA>AG:594 | A | G | 6,83 | 0.71 | 0.00 | 0.50 | 0.00 | yes | PE594 - 4.75 % inclusion |
| 596 | CA | 29486707 | 595-596A>C | A | C | 4,23 | 0.00 | 0.00 | 0.00 | 0.00 | yes | none |
| 596 | CA | 29486707 | 595-596A>G | A | G | 5,28 | 0.01 | 0.00 | 0.00 | 0.00 | yes | none |
| 601 | CD | 29486712 | 598-601T>G | T | G | 5,66 | 0.00 | 0.00 | 0.00 | 0.00 | yes | none |
| 616 | DD | 29486727 | GA>GT:616 | A | T | 9,72 | 0.00 | 0.00 | 0.01 | 0.00 | yes | none |
| 618 | CD | 29486729 | 620-618A>G | A | G | 7,64 | 0.00 | 0.00 | 0.00 | 0.00 | yes | none |
| 623 | CD | 29486734 | 620-623T>G | T | G | 6,93 | 0.00 | 0.00 | 0.00 | 0.00 | yes | none |
| 644 | DA | 29486755 | AA>AG:644 | A | G | 4,44 | 0.00 | 0.00 | 0.00 | 0.00 | yes | none |
| 652 | DA | 29486763 | TG>AG:653 | T | A | 7,96 | 0.01 | 0.00 | 0.00 | 0.00 | no | N/A |
| 653 | DA | 29486764 | GG>AG:654 | G | A | 7,38 | 0.00 | 0.00 | 0.00 | 0.00 | yes | none |
| 657 | DA | 29486768 | AC>AG:657 | C | G | 5,5 | 0.00 | 0.00 | 0.00 | 0.00 | yes | none |
| 659 | DA | 29486770 | AT>AG:659 | T | G | 5,96 | 0.00 | 0.00 | 0.00 | 0.00 | yes | none |
| 659 | CD | 29486770 | 662-659T>C | T | C | 10,65 | 0.00 | 0.00 | 0.01 | 0.00 | yes | none |
| 659 | CD | 29486770 | 662-659T>A | T | A | 10,57 | 0.00 | 0.00 | 0.04 | 0.00 | no | N/A |
| 666 | CD | 29486777 | 662-666A>T | A | T | 11,01 | 0.00 | 0.00 | 0.05 | 0.00 | yes | none |
| 704 | DA | 29486815 | AA>AG:704 | A | G | 5,81 | 0.00 | 0.00 | 0.00 | 0.00 | no | N/A |
| 705 | DA | 29486816 | AT>AG:705 | T | G | 7,82 | 0.00 | 0.00 | 0.00 | 0.00 | yes | none |
| 705 | DA | 29486816 | TG>AG:706 | T | A | 5,56 | 0.00 | 0.00 | 0.00 | 0.00 | yes | none |
| 710 | DA | 29486821 | TG>AG:711 | T | A | 5,16 | 0.00 | 0.00 | 0.00 | 0.00 | yes | none |
| 716 | DA | 29486827 | AA>AG:716 | A | G | 4,07 | 0.00 | 0.00 | 0.00 | 0.00 | yes | none |

**Library 4**

| intronic position | type | chrom. position | characteristics | ref | alt | MaxEnt score | SpliceAI prediction | | | | acquired | effect on splicing |
|---|---|---|---|---|---|---|---|---|---|---|---|---|
| | | | | | | | acceptor gain | acceptor loss | donor gain | donor loss | | |
| 721 | DA | 29486832 | AT>AG:721 | T | G | 5,95 | 0.00 | 0.00 | 0.00 | 0.00 | yes | none |
| 727 | DA | 29486838 | AT>AG:727 | T | G | 4,58 | 0.00 | 0.00 | 0.00 | 0.00 | yes | none |
| 728 | DA | 29486839 | TG>AG:729 | T | A | 3,75 | 0.00 | 0.00 | 0.00 | 0.00 | yes | none |
| 748 | DA | 29486859 | AT>AG:748 | T | G | 6,28 | 0.00 | 0.00 | 0.00 | 0.00 | yes | none |
| 763 | DA | 29486874 | TG>AG:764 | T | A | 11,49 | 0.12 | 0.00 | 0.09 | 0.00 | yes | none |
| 764 | DA | 29486875 | GG>AG:765 | G | A | 10,98 | 0.02 | 0.00 | 0.01 | 0.00 | yes | none |
| 769 | CA | 29486880 | 777-769A>G | A | G | 3,97 | 0.01 | 0.00 | 0.00 | 0.00 | yes | none |
| 769 | CA | 29486880 | 777-769A>C | A | C | 6,1 | 0.01 | 0.00 | 0.00 | 0.00 | yes | none |
| 770 | CA | 29486881 | 777-770G>A | G | A | 4,77 | 0.00 | 0.00 | 0.00 | 0.00 | yes | none |
| 770 | CA | 29486881 | 777-770G>C | G | C | 5,47 | 0.01 | 0.00 | 0.00 | 0.00 | yes | none |
| 813 | DA | 29486924 | AC>AG:813 | C | G | 8,65 | 0.00 | 0.00 | 0.00 | 0.00 | yes | none |
| 815 | DA | 29486926 | AT>AG:815 | T | G | 9,94 | 0.01 | 0.00 | 0.01 | 0.00 | yes | none |
| 814 | DD | 29486925 | AT>GT:815 | A | G | 7,26 | 0.01 | 0.00 | 0.00 | 0.00 | yes | none |
| 817 | DA | 29486928 | AT>AG:817 | T | G | 10,27 | 0.00 | 0.00 | 0.00 | 0.00 | yes | none |
| 820 | DA | 29486931 | TG>AG:821 | T | A | 5,42 | 0.00 | 0.00 | 0.00 | 0.00 | yes | none |
| 823 | CA | 29486934 | 834-823A>G | A | G | 5,93 | 0.01 | 0.00 | 0.00 | 0.00 | yes | none |
| 823 | CA | 29486934 | 834-823A>C | A | C | 7,5 | 0.01 | 0.00 | 0.00 | 0.00 | yes | none |
| 824 | CA | 29486935 | 834-824G>T | G | T | 7 | 0.00 | 0.00 | 0.00 | 0.00 | yes | none |
| 835 | CA | 29486946 | 834-835T>G | T | G | 3,64 | 0.00 | 0.00 | 0.00 | 0.00 | yes | none |
| 882 | DD | 29486993 | TT>GT:883 | T | G | 6,04 | 0.00 | 0.00 | 0.00 | 0.00 | yes | none |
| 906 | CD | 29487017 | 910-906G>C | G | C | 6,91 | 0.00 | 0.00 | 0.02 | 0.00 | yes | none |
| 906 | CD | 29487017 | 910-906G>A | G | A | 5,76 | 0.00 | 0.00 | 0.00 | 0.00 | yes | none |
| 907 | CD | 29487018 | 910-907G>A | G | A | 7,81 | 0.05 | 0.00 | 0.13 | 0.00 | yes | none |
| 907 | CD | 29487018 | 910-907G>C | G | C | 6,22 | 0.01 | 0.00 | 0.01 | 0.00 | yes | none |
| 912 | CD | 29487023 | 910-912G>A | G | A | 8,05 | 0.06 | 0.00 | 0.21 | 0.00 | yes | PE912 - 0.7% inclusion |
| 912 | CD | 29487023 | 910-912G>C | G | C | 6,13 | 0.01 | 0.00 | 0.03 | 0.00 | yes | none |
| 912 | CD | 29487023 | 910-912G>T | G | T | 5,84 | 0.01 | 0.00 | 0.01 | 0.00 | yes | none |
| 914 | CD | 29487025 | 910-914G>T | G | T | 6,59 | 0.02 | 0.00 | 0.11 | 0.00 | yes | none |
| 949 | CA | 29487060 | 956-949A>T | A | T | 9,53 | 0.01 | 0.00 | 0.00 | 0.00 | yes | none |
| 949 | CA | 29487060 | 956-949A>C | A | C | 10,21 | 0.00 | 0.00 | 0.00 | 0.00 | yes | none |

**Library 5**

| intronic position | type | chrom. position | characteristics | ref | alt | MaxEnt score | SpliceAI prediction | | | | acquired | effect on splicing |
|---|---|---|---|---|---|---|---|---|---|---|---|---|
| | | | | | | | acceptor gain | acceptor loss | donor gain | donor loss | | |
| 956 | DA | 29487067 | GG>AG:957 | G | A | 5 | 0.00 | 0.00 | 0.00 | 0.00 | no | N/A |
| 958 | DD | 29487069 | GG>GT:958 | G | T | 9,31 | 0.03 | 0.00 | 0.00 | 0.00 | yes | none |
| 959 | CA | 29487070 | 956-959A>T | A | T | 8,54 | 0.01 | 0.00 | 0.00 | 0.00 | yes | none |
| 959 | CA | 29487070 | 956-959A>C | A | C | 8,78 | 0.00 | 0.00 | 0.00 | 0.00 | yes | none |
| 959 | DD | 29487070 | GA>GT:959 | A | T | 7,26 | 0.01 | 0.00 | 0.00 | 0.00 | yes | none |
| 963 | CD | 29487074 | 966-963C>A | C | A | 6,16 | 0.01 | 0.00 | 0.00 | 0.00 | yes | none |
| 964 | CD | 29487075 | 966-964T>G | T | G | 8,99 | 0.01 | 0.00 | 0.00 | 0.00 | yes | none |
| 970 | CD | 29487081 | 966-970C>T | C | T | 8,56 | 0.01 | 0.00 | 0.00 | 0.00 | no | N/A |
| 976 | CD | 29487087 | 980-976A>G | A | G | 6,16 | 0.00 | 0.00 | 0.00 | 0.00 | no | N/A |
| 978 | CD | 29487089 | 980-978T>G | T | G | 10,22 | 0.01 | 0.00 | 0.06 | 0.00 | yes | none |
| 978 | CD | 29487089 | 980-978T>C | T | C | 7,35 | 0.01 | 0.00 | 0.00 | 0.00 | yes | none |
| 978 | CD | 29487089 | 980-978T>A | T | A | 7,31 | 0.00 | 0.00 | 0.00 | 0.00 | yes | none |
| 984 | CD | 29487095 | 980-984C>T | C | T | 8,62 | 0.03 | 0.00 | 0.04 | 0.00 | yes | none |
| 984 | CD | 29487095 | 980-984C>G | C | G | 6,17 | 0.00 | 0.00 | 0.00 | 0.00 | yes | none |
| 1010 | DA | 29487121 | TG>AG:1011 | T | A | 7,11 | 0.26 | 0.00 | 0.01 | 0.00 | yes | none |
| 1012 | CA | 29487123 | 1017-1012A>G | A | G | 5,96 | 0.00 | 0.00 | 0.00 | 0.00 | yes | none |
| 1012 | CA | 29487123 | 1017-1012A>C | A | C | 7,18 | 0.17 | 0.00 | 0.03 | 0.00 | yes | PE1012C - 2.16% inclusion level |
| 1012 | CA | 29487123 | 1017-1012A>T | A | T | 7,71 | 0.17 | 0.00 | 0.04 | 0.00 | yes | PE1012T - 1.72% inclusion |
| 1019 | CA | 29487130 | 1017-1019G>T | G | T | 3,57 | 0.16 | 0.00 | 0.02 | 0.00 | yes | PE1019 - 3.43% inclusion |
| 1031 | DA | 29487142 | GG>AG:1032 | G | A | 3,71 | 0.00 | 0.00 | 0.00 | 0.00 | yes | none |
| 1068 | DA | 29487179 | AT>AG:1068 | T | G | 4,66 | 0.02 | 0.00 | 0.00 | 0.00 | no | N/A |
| 1069 | DA | 29487180 | TG>AG:1070 | T | A | 4,74 | 0.00 | 0.00 | 0.00 | 0.00 | no | N/A |
| 1073 | DA | 29487184 | AA>AG:1073 | A | G | 6,5 | 0.00 | 0.00 | 0.00 | 0.00 | no | N/A |
| 1085 | DA | 29487196 | CG>AG:1086 | C | A | 7,05 | 0.00 | 0.00 | 0.00 | 0.00 | yes | none |
| 1086 | DA | 29487197 | GG>AG:1087 | G | A | 6,19 | 0.00 | 0.00 | 0.00 | 0.00 | yes | none |
| 1093 | DA | 29487204 | AA>AG:1093 | A | G | 7,41 | 0.00 | 0.00 | 0.00 | 0.00 | yes | none |
| 1095 | DD | 29487206 | CT>GT:1096 | C | G | 6,8 | 0.06 | 0.00 | 0.08 | 0.00 | yes | none |
| 1119 | CD | 29487230 | 1121-1119A>G | A | G | 6,52 | 0.00 | 0.00 | 0.05 | 0.00 | yes, in combination with 1156C>G | N/A |
| 1124 | CD | 29487235 | 1121-1124C>G | C | G | 7,33 | 0.05 | 0.00 | 0.09 | 0.00 | no | N/A |
| 1137 | DD | 29487248 | GC>GT:1137 | C | T | 9,8 | 0.71 | 0.00 | 0.72 | 0.00 | yes | PE1137, positive control, 52.4 % incl |
| 1139 | CD | 29487250 | 1141-1139T>G | T | G | 7,88 | 0.07 | 0.00 | 0.12 | 0.00 | yes | none |
| 1156 | DD | 29487267 | CT>GT:1157 | C | G | 7,51 | 0.39 | 0.00 | 0.36 | 0.00 | yes | PE1156 - 8.9% inclusion |

**Library 5** **continued**

| intronic position | type | chrom. position | characteristics | ref | alt | MaxEnt score | SpliceAI prediction acceptor gain | acceptor loss | donor gain | donor loss | acquired | effect on splicing |
|---|---|---|---|---|---|---|---|---|---|---|---|---|
| 1170 | DA | 29487281 | AT>AG:1170 | T | G | 5,91 | 0.02 | 0.00 | 0.01 | 0.00 | yes | none |
| 1176 | DA | 29487287 | AT>AG:1176 | T | G | 8,27 | 0.02 | 0.00 | 0.00 | 0.00 | yes | none |
| 1178 | DA | 29487289 | AA>AG:1178 | A | G | 10,26 | 0.05 | 0.00 | 0.00 | 0.00 | yes | none |

**Library 6**

| intronic position | type | chrom. position | characteristics | ref | alt | MaxEnt score | SpliceAI prediction acceptor gain | acceptor loss | donor gain | donor loss | acquired | effect on splicing |
|---|---|---|---|---|---|---|---|---|---|---|---|---|
| 1199 | DA | 29487310 | TG>AG:1200 | T | A | 6,28 | 0.00 | 0.00 | 0.00 | 0.00 | yes | none |
| 1203 | CD | 29487314 | 1202-1203C>A | C | A | 8,92 | 0.00 | 0.00 | 0.01 | 0.00 | yes | none |
| 1203 | CD | 29487314 | 1202-1203C>G | C | G | 6,9 | 0.00 | 0.00 | 0.00 | 0.00 | yes | none |
| 1207 | DD | 29487318 | GC>GT:1207 | C | T | 8,02 | 0.00 | 0.00 | 0.00 | 0.00 | yes | none |
| 1229 | CA | 29487340 | 1248-1229A>C | A | C | 3,4 | 0.00 | 0.00 | 0.00 | 0.00 | no | N/A |
| 1230 | CA | 29487341 | 1248-1230A>T | A | T | 5,37 | 0.00 | 0.00 | 0.01 | 0.00 | no | N/A |
| 1230 | DA | 29487341 | AA>AG:1230 | A | G | 8,88 | 0.00 | 0.00 | 0.02 | 0.00 | | |
| 1231 | CA | 29487342 | 1248-1231G>A | G | A | 3,9 | 0.00 | 0.00 | 0.00 | 0.00 | yes | none |
| 1231 | CA | 29487342 | 1248-1231G>C | G | C | 4,45 | 0.00 | 0.00 | 0.00 | 0.00 | yes | none |
| 1235 | CD | 29487346 | 1232-1235T>G | T | G | 6,39 | 0.00 | 0.00 | 0.00 | 0.00 | yes | none |
| 1262 | DA | 29487373 | TG>AG:1263 | T | A | 5,27 | 0.00 | 0.00 | 0.00 | 0.00 | yes | none |
| 1263 | DA | 29487374 | GG>AG:1264 | G | A | 6,03 | 0.00 | 0.00 | 0.01 | 0.00 | yes | none |
| 1285 | CA | 29487396 | 1298-1285G>C | G | C | 5,36 | 0.00 | 0.00 | 0.00 | 0.00 | yes | none |
| 1296 | CA | 29487407 | 1298-1296A>C | A | C | 8,33 | 0.00 | 0.00 | 0.01 | 0.00 | yes | none |
| 1297 | DA | 29487408 | AA>AG:1297 | A | G | 6,92 | 0.00 | 0.00 | 0.00 | 0.00 | yes | none |
| 1300 | CA | 29487411 | 1298-1300A>T | A | T | 5,96 | 0.00 | 0.00 | 0.00 | 0.00 | yes | none |
| 1300 | CA | 29487411 | 1298-1300A>G | A | G | 6,38 | 0.00 | 0.00 | 0.00 | 0.00 | yes | none |
| 1301 | DA | 29487412 | AT>AG:1301 | T | G | 5,84 | 0.00 | 0.00 | 0.03 | 0.00 | no | N/A |
| 1311 | DA | 29487422 | AA>AG:1311 | A | G | 3,7 | 0.00 | 0.00 | 0.02 | 0.00 | yes | none |
| 1321 | DA | 29487432 | AT>AG:1321 | T | G | 4,97 | 0.01 | 0.00 | 0.14 | 0.00 | yes, in combination with 1322T>A | N/A |

**Library 6** **continued**

| intronic position | type | chrom. position | characteristics | ref | alt | MaxEnt score | SpliceAI prediction: acceptor gain | acceptor loss | donor gain | donor loss | acquired | effect on splicing |
|---|---|---|---|---|---|---|---|---|---|---|---|---|
| 1322 | DA | 29487433 | TG>AG:1323 | T | A | 7,71 | 0.38 | 0.00 | 0.57 | 0.00 | yes | PE1322 - 1.23% inclusion |
| 1326 | DA | 29487437 | CG>AG:1327 | C | A | 7,33 | 0.08 | 0.00 | 0.29 | 0.00 | yes, in combination with 1329T>A | N/A |
| 1329 | DA | 29487440 | TG>AG:1330 | T | A | 6,76 | 0.02 | 0.00 | 0.14 | 0.00 | yes | none |
| 1337 | DA | 29487448 | AT>AG:1337 | T | G | 4,93 | 0.00 | 0.00 | 0.01 | 0.00 | yes | none |
| 1349 | DA | 29487460 | GG>AG:1350 | G | A | 5,02 | 0.00 | 0.00 | 0.00 | 0.00 | yes | none |
| 1362 | DD | 29487473 | AT>GT:1363 | A | G | 7,4 | 0.00 | 0.00 | 0.00 | 0.00 | yes | none |
| 1378 | DA | 29487489 | AA>AG:1378 | A | G | 5,07 | 0.00 | 0.00 | 0.00 | 0.00 | yes | none |
| 1379 | DA | 29487490 | AA>AG:1379 | A | G | 4,25 | 0.00 | 0.00 | 0.02 | 0.00 | yes | none |
| 1389 | CD | 29487500 | 1388-1389G>A | G | A | 10,22 | 0.00 | 0.00 | 0.08 | 0.00 | insufficient read depth | none |
| 1392 | CD | 29487503 | 1388-1392C>T | C | T | 10,47 | 0.00 | 0.00 | 0.08 | 0.00 | yes | none |
| 1392 | DD | 29487503 | CT>GT:1393 | C | G | 5,77 | 0.00 | 0.00 | 0.00 | 0.00 | yes | none |

**Library 7**

| intronic position | type | chrom. position | characteristics | ref | alt | MaxEnt score | SpliceAI prediction: acceptor gain | acceptor loss | donor gain | donor loss | acquired | effect on splicing |
|---|---|---|---|---|---|---|---|---|---|---|---|---|
| 1407 | CD | 29487518 | 1410-1407T>A | T | A | 7,64 | 0.00 | 0.00 | 0.00 | 0.00 | yes, in combination with 1533G>T | none |
| 1412 | CD | 29487523 | 1410-1412T>A | T | A | 6,49 | 0.00 | 0.00 | 0.00 | 0.00 | yes | none |
| 1413 | CD | 29487524 | 1410-1413T>G | T | G | 8,35 | 0.00 | 0.00 | 0.00 | 0.00 | no | N/A |
| 1419 | DD | 29487530 | TT>GT:1420 | T | G | 6,39 | 0.00 | 0.00 | 0.00 | 0.00 | yes | none |
| 1425 | DA | 29487536 | TG>AG:1426 | T | A | 4,23 | 0.00 | 0.00 | 0.00 | 0.00 | yes | none |
| 1429 | DD | 29487540 | AT>GT:1430 | A | G | 5,97 | 0.00 | 0.00 | 0.00 | 0.00 | yes | none |
| 1432 | DA | 29487543 | AT>AG:1432 | T | G | 4,75 | 0.00 | 0.00 | 0.00 | 0.00 | yes | none |
| 1438 | DA | 29487549 | TG>AG:1439 | T | A | 3,91 | 0.00 | 0.00 | 0.00 | 0.00 | insufficient read depth | none |
| 1444 | DA | 29487555 | AT>AG:1444 | T | G | 6,7 | 0.01 | 0.00 | 0.00 | 0.00 | yes | none |

**Library 7** **continued**

| intronic position | type | chrom. position | characteristics | ref | alt | MaxEnt score | SpliceAI prediction | | | | acquired | effect on splicing |
|---|---|---|---|---|---|---|---|---|---|---|---|---|
| | | | | | | | acceptor gain | acceptor loss | donor gain | donor loss | | |
| 1450 | DA | 29487561 | AA>AG:1450 | A | G | 6,79 | 0.00 | 0.00 | 0.00 | 0.00 | yes | none |
| 1454 | DA | 29487565 | GG>AG:1455 | G | A | 3,4 | 0.00 | 0.00 | 0.00 | 0.00 | no | N/A |
| 1492 | DD | 29487603 | GG>GT:1492 | G | T | 6,09 | 0.00 | 0.00 | 0.00 | 0.00 | yes | none |
| 1499 | DD | 29487610 | GG>GT:1499 | G | T | 9,45 | 0.00 | 0.00 | 0.02 | 0.00 | yes | none |
| 1503 | DD | 29487614 | AT>GT:1504 | A | G | 7,93 | 0.00 | 0.00 | 0.00 | 0.00 | yes | none |
| 1520 | CA | 29487631 | 1533-1520T>C | T | C | 6,09 | 0.00 | 0.00 | 0.00 | 0.00 | yes | none |
| 1524 | CA | 29487635 | 1533-1524C>T | C | T | 6,51 | 0.00 | 0.00 | 0.00 | 0.00 | yes | none |
| 1526 | DA | 29487637 | TG>AG:1527 | T | A | 6,47 | 0.00 | 0.00 | 0.00 | 0.00 | yes, in combination with 1530T>G | none |
| 1530 | DA | 29487641 | AT>AG:1530 | T | G | 4,31 | 0.00 | 0.00 | 0.00 | 0.00 | yes | none |
| 1532 | CA | 29487643 | 1543-1532A>T | A | T | 5,82 | 0.06 | 0.00 | 0.06 | 0.00 | yes | none |
| 1533 | CA | 29487644 | 1543-1533G>A | G | A | 3,73 | 0.01 | 0.00 | 0.01 | 0.00 | yes | none |
| 1533 | CA | 29487644 | 1543-1533G>C | G | C | 3,99 | 0.01 | 0.00 | 0.01 | 0.00 | yes | none |
| 1533 | CA | 29487644 | 1543-1533G>T | G | T | 4,64 | 0.03 | 0.00 | 0.03 | 0.00 | yes | none |
| 1534 | CA | 29487645 | 1533-1534T>A | T | A | 6,94 | 0.00 | 0.00 | 0.00 | 0.00 | yes | none |
| 1534 | CA | 29487645 | 1533-1534T>G | T | G | 7,94 | 0.00 | 0.00 | 0.00 | 0.00 | yes | none |
| 1558 | DA | 29487669 | TG>AG:1559 | T | A | 5,14 | 0.00 | 0.00 | 0.00 | 0.00 | yes | none |
| 1562 | DA | 29487673 | TG>AG:1563 | T | A | 4,73 | 0.00 | 0.00 | 0.00 | 0.00 | yes | none |
| 1566 | DA | 29487677 | AA>AG:1566 | A | G | 7,18 | 0.00 | 0.00 | 0.00 | 0.00 | yes | none |
| 1578 | CD | 29487689 | 1575-1578A>G | A | G | 5,8 | 0.00 | 0.00 | 0.00 | 0.00 | yes | none |
| 1600 | CD | 29487711 | 1603-1600T>A | T | A | 5,87 | 0.00 | 0.00 | 0.00 | 0.00 | yes | none |
| 1604 | CD | 29487715 | 1603-1604G>A | G | A | 5,88 | 0.00 | 0.00 | 0.00 | 0.00 | insufficient read depth | none |
| 1605 | CD | 29487716 | 1603-1605G>A | G | A | 7,61 | 0.01 | 0.00 | 0.01 | 0.00 | yes | none |
| 1605 | CD | 29487716 | 1603-1605G>C | G | C | 5,75 | 0.01 | 0.00 | 0.02 | 0.00 | yes | none |
| 1606 | DD | 29487717 | GG>GT:1606 | G | T | 6,46 | 0.00 | 0.00 | 0.00 | 0.00 | insufficient read depth | none |
| 1607 | CD | 29487718 | 1603-1607G>T | G | T | 6,18 | 0.00 | 0.00 | 0.00 | 0.00 | insufficient read depth | none |
| 1624 | CA | 29487735 | 1634-1624G>C | G | C | 7,56 | 0.00 | 0.00 | 0.00 | 0.00 | no | N/A |
| 1624 | CA | 29487735 | 1634-1624G>T | G | T | 8,25 | 0.00 | 0.00 | 0.00 | 0.00 | no | N/A |
| 1629 | CA | 29487740 | 1634-1629G>T | G | T | 9,1 | 0.00 | 0.00 | 0.00 | 0.00 | no | N/A |

**Library 8**

| intronic position | type | chrom. position | characteristics | ref | alt | MaxEnt score | SpliceAI prediction acceptor gain | acceptor loss | donor gain | donor loss | acquired | effect on splicing |
|---|---|---|---|---|---|---|---|---|---|---|---|---|
| 1637 | CA | 29487748 | 1634-1637T>C | T | C | 6,97 | 0.00 | 0.00 | 0.00 | 0.00 | yes | none |
| 1640 | DA | 29487751 | AC>AG:1640 | C | G | 3,83 | 0.00 | 0.00 | 0.00 | 0.00 | yes | none |
| 1656 | DA | 29487767 | AA>AG:1656 | A | G | 3,83 | 0.00 | 0.00 | 0.00 | 0.00 | yes | none |
| 1657 | DA | 29487768 | AC>AG:1657 | C | G | 5,27 | 0.00 | 0.00 | 0.00 | 0.00 | yes, in combination with 1656A>G | none |
| 1659 | DA | 29487770 | AT>AG:1659 | T | G | 6,77 | 0.00 | 0.00 | 0.00 | 0.00 | yes | none |
| 1661 | CD | 29487772 | 1663-1661T>G | T | G | 5,73 | 0.00 | 0.00 | 0.00 | 0.00 | yes | none |
| 1661 | DA | 29487772 | TG>AG:1662 | T | A | 3,7 | 0.00 | 0.00 | 0.00 | 0.00 | yes | none |
| 1665 | DA | 29487776 | AA>AG:1665 | A | G | 3,78 | 0.00 | 0.00 | 0.00 | 0.00 | yes | none |
| 1666 | CD | 29487777 | 1663-1666T>G | T | G | 7,4 | 0.00 | 0.00 | 0.00 | 0.00 | yes | none |
| 1670 | CD | 29487781 | 1674-1670G>C | G | C | 8,55 | 0.00 | 0.00 | 0.00 | 0.00 | yes | none |
| 1670 | CD | 29487781 | 1674-1670G>A | G | A | 8,49 | 0.00 | 0.00 | 0.00 | 0.00 | yes | none |
| 1677 | CD | 29487788 | 1674-1677T>G | T | G | 10,06 | 0.00 | 0.00 | 0.02 | 0.00 | yes | none |
| 1677 | CD | 29487788 | 1674-1677T>C | T | C | 8,1 | 0.00 | 0.00 | 0.00 | 0.00 | yes | none |
| 1678 | CD | 29487789 | 1674-1678A>T | A | T | 8,88 | 0.00 | 0.00 | 0.00 | 0.00 | yes | none |
| 1678 | CD | 29487789 | 1674-1678A>G | A | G | 8,73 | 0.00 | 0.00 | 0.00 | 0.00 | yes | none |
| 1682 | DD | 29487793 | GA>GT:1682 | A | T | 5,87 | 0.00 | 0.00 | 0.00 | 0.00 | yes | none |
| 1684 | CD | 29487795 | 1686-1684T>G | T | G | 7,64 | 0.00 | 0.00 | 0.00 | 0.00 | yes | none |
| 1689 | CD | 29487800 | 1686-1689T>G | T | G | 5,87 | 0.00 | 0.00 | 0.00 | 0.00 | yes | none |
| 1692 | DA | 29487803 | GG>AG:1693 | G | A | 5,33 | 0.00 | 0.00 | 0.00 | 0.00 | yes | none |
| 1714 | DA | 29487825 | TG>AG:1715 | T | A | 3,8 | 0.00 | 0.00 | 0.00 | 0.00 | yes | none |
| 1721 | DA | 29487832 | TG>AG:1722 | T | A | 3,5 | 0.00 | 0.00 | 0.00 | 0.00 | yes | none |
| 1725 | DA | 29487836 | TG>AG:1726 | T | A | 4,92 | 0.00 | 0.00 | 0.00 | 0.00 | yes | none |
| 1732 | DA | 29487843 | AA>AG:1732 | A | G | 6,44 | 0.00 | 0.00 | 0.00 | 0.00 | yes | none |
| 1733 | DA | 29487844 | AT>AG:1733 | T | G | 4,88 | 0.00 | 0.00 | 0.00 | 0.00 | yes | none |
| 1735 | DA | 29487846 | AC>AG:1735 | C | G | 6,54 | 0.00 | 0.00 | 0.00 | 0.00 | yes | none |
| 1741 | CA | 29487852 | 1744-1741G>T | G | T | 5,02 | 0.00 | 0.00 | 0.00 | 0.00 | yes | none |
| 1742 | CA | 29487853 | 1744-1742A>T | A | T | 6,17 | 0.00 | 0.00 | 0.00 | 0.00 | no | N/A |
| 1742 | CA | 29487853 | 1744-1742A>C | A | C | 7,82 | 0.00 | 0.00 | 0.00 | 0.00 | yes | none |
| 1745 | DD | 29487856 | AT>GT:1746 | A | G | 7,58 | 0.00 | 0.00 | 0.00 | 0.00 | yes | none |
| 1747 | CA | 29487858 | 1744-1747A>C | A | C | 4,51 | 0.00 | 0.00 | 0.00 | 0.00 | yes | none |
| 1765 | DD | 29487876 | AT>GT:1766 | A | G | 8,78 | 0.00 | 0.00 | 0.00 | 0.00 | yes | none |

**Library 8** **continued**

| intronic position | type | chrom. position | characteristics | ref | alt | MaxEnt score | SpliceAI prediction | | | | acquired | effect on splicing |
|---|---|---|---|---|---|---|---|---|---|---|---|---|
| | | | | | | | acceptor gain | acceptor loss | donor gain | donor loss | | |
| 1785 | DD | 29487896 | TT>GT:1786 | T | G | 7,69 | 0.00 | 0.00 | 0.00 | 0.00 | yes | none |
| 1788 | CD | 29487899 | 1790-1788T>G | T | G | 7,57 | 0.00 | 0.00 | 0.00 | 0.00 | yes | none |
| 1799 | DA | 29487910 | AA>AG:1799 | A | G | 3,65 | 0.00 | 0.00 | 0.00 | 0.00 | yes | none |
| 1802 | DA | 29487913 | AT>AG:1802 | T | G | 3,61 | 0.00 | 0.00 | 0.00 | 0.00 | yes | none |
| 1802 | CD | 29487913 | 1804-1802T>G | T | G | 7,84 | 0.00 | 0.00 | 0.00 | 0.00 | yes | none |
| 1807 | CD | 29487918 | 1804-1807A>G | A | G | 5,87 | 0.00 | 0.00 | 0.00 | 0.00 | yes | none |
| 1817 | DA | 29487928 | CG>AG:1818 | C | A | 6,91 | 0.00 | 0.00 | 0.00 | 0.00 | yes | none |
| 1826 | DA | 29487937 | AC>AG:1826 | C | G | 6,14 | 0.00 | 0.00 | 0.00 | 0.00 | yes | none |
| 1827 | DA | 29487938 | TG>AG:1828 | T | A | 7,53 | 0.00 | 0.00 | 0.00 | 0.00 | yes, in combination with 1826C>G | none |
| 1830 | DA | 29487941 | TG>AG:1831 | T | A | 8,2 | 0.00 | 0.00 | 0.00 | 0.00 | yes | none |
| 1834 | DA | 29487945 | AA>AG:1834 | A | G | 4,86 | 0.00 | 0.00 | 0.00 | 0.00 | yes | none |
| 1839 | CD | 29487950 | 1841-1839T>G | T | G | 6,57 | 0.00 | 0.00 | 0.00 | 0.00 | yes | none |

**Library 9**

| intronic position | type | chrom. position | characteristics | ref | alt | MaxEnt score | SpliceAI prediction | | | | acquired | effect on splicing |
|---|---|---|---|---|---|---|---|---|---|---|---|---|
| | | | | | | | acceptor gain | acceptor loss | donor gain | donor loss | | |
| 1853 | DA | 29487964 | AT>AG:1853 | T | G | 6,39 | 0.00 | 0.00 | 0.00 | 0.00 | yes | none |
| 1856 | DA | 29487967 | AA>AG:1856 | A | G | 7,26 | 0.00 | 0.00 | 0.00 | 0.00 | yes | none |
| 1857 | DA | 29487968 | AA>AG:1857 | A | G | 5,19 | 0.00 | 0.00 | 0.00 | 0.00 | yes | none |
| 1858 | DA | 29487969 | AT>AG:1858 | T | G | 4,69 | 0.00 | 0.00 | 0.00 | 0.00 | yes | none |
| 1862 | DA | 29487973 | AC>AG:1862 | C | G | 6,33 | 0.00 | 0.00 | 0.00 | 0.00 | yes | none |
| 1865 | DA | 29487976 | AT>AG:1865 | T | G | 3,71 | 0.00 | 0.00 | 0.00 | 0.00 | yes | none |
| 1869 | DA | 29487980 | AA>AG:1869 | A | G | 6,07 | 0.00 | 0.00 | 0.00 | 0.00 | yes | none |
| 1872 | DA | 29487983 | AA>AG:1872 | A | G | 5,23 | 0.00 | 0.00 | 0.00 | 0.00 | yes | none |
| 1872 | CD | 29487983 | 1874-1872A>G | A | G | 8,99 | 0.00 | 0.00 | 0.00 | 0.00 | yes | none |
| 1877 | CD | 29487988 | 1874-1877T>G | T | G | 6,6 | 0.00 | 0.00 | 0.00 | 0.00 | yes | none |
| 1892 | DA | 29488003 | TG>AG:1893 | T | A | 5,98 | 0.00 | 0.00 | 0.00 | 0.00 | yes | none |
| 1896 | DA | 29488007 | AC>AG:1896 | C | G | 3,66 | 0.00 | 0.00 | 0.00 | 0.00 | yes | none |
| 1898 | CA | 29488009 | 1916-1898A>T | A | T | 8,49 | 0.00 | 0.00 | 0.00 | 0.00 | yes | none |

**Library 9** **continued**

| intronic position | type | chrom. position | characteristics | ref | alt | MaxEnt score | SpliceAI prediction acceptor gain | acceptor loss | donor gain | donor loss | acquired | effect on splicing |
|---|---|---|---|---|---|---|---|---|---|---|---|---|
| 1898 | DA | 29488009 | AA>AG:1898 | A | G | 6,56 | 0.00 | 0.00 | 0.00 | 0.00 | yes | none |
| 1905 | CA | 29488016 | 1916-1905T>C | T | C | 7,69 | 0.00 | 0.00 | 0.00 | 0.00 | yes | none |
| 1907 | DA | 29488018 | AT>AG:1907 | T | G | 4,62 | 0.00 | 0.00 | 0.00 | 0.00 | yes | none |
| 1917 | CA | 29488028 | 1916-1917T>C | T | C | 7,98 | 0.00 | 0.00 | 0.00 | 0.00 | yes | none |
| 1917 | CA | 29488028 | 1916-1917T>G | T | G | 9,53 | 0.02 | 0.00 | 0.01 | 0.00 | yes | none |
| 1920 | CD | 29488031 | 1917-1920T>G | T | G | 7,68 | 0.00 | 0.00 | 0.00 | 0.00 | yes | none |
| 1945 | DA | 29488056 | AC>AG:1945 | C | G | 4,49 | 0.00 | 0.00 | 0.00 | 0.00 | yes | none |
| 1951 | DA | 29488062 | AA>AG:1951 | A | G | 9,66 | 0.03 | 0.00 | 0.01 | 0.00 | yes | none |
| 1952 | DA | 29488063 | AA>AG:1952 | A | G | 4,85 | 0.00 | 0.00 | 0.00 | 0.00 | yes | none |
| 1953 | DA | 29488064 | AT>AG:1953 | T | G | 5,65 | 0.00 | 0.00 | 0.00 | 0.00 | yes | none |
| 1956 | DA | 29488067 | AA>AG:1956 | A | G | 4,56 | 0.00 | 0.00 | 0.00 | 0.00 | yes | none |
| 1957 | DA | 29488068 | AA>AG:1957 | A | G | 3,92 | 0.02 | 0.00 | 0.01 | 0.00 | ufficient read dep | none |
| 1957 | CD | 29488068 | 1959-1957A>G | A | G | 8,88 | 0.02 | 0.00 | 0.01 | 0.00 | | none |
| 1961 | DA | 29488072 | AA>AG:1961 | A | G | 3,51 | 0.00 | 0.00 | 0.00 | 0.00 | yes | none |
| 1962 | CD | 29488073 | 1959-1962A>G | A | G | 9,72 | 0.03 | 0.00 | 0.01 | 0.00 | yes | none |
| 1969 | DD | 29488080 | GC>GT:1969 | C | T | 5,77 | 0.00 | 0.00 | 0.00 | 0.00 | yes | none |
| 1994 | CD | 29488105 | 1996-1994A>G | A | G | 7,48 | 0.74 | 0.00 | 0.66 | 0.00 | yes | PE1994 - 3.8% inclusion |
| 1997 | CA | 29488108 | 2015-1997A>C | A | C | 4,64 | 0.01 | 0.00 | 0.00 | 0.00 | yes | none |
| 1997 | CA | 29488108 | 2015-1997A>T | A | T | 5,22 | 0.00 | 0.00 | 0.00 | 0.00 | yes | none |
| 2011 | DA | 29488122 | AC>AG:2011 | C | G | 4,79 | 0.00 | 0.00 | 0.00 | 0.00 | yes | none |
| 2013 | DA | 29488124 | AT>AG:2013 | T | G | 3,89 | 0.01 | 0.00 | 0.00 | 0.00 | yes | none |
| 2016 | CA | 29488127 | 2015-2016T>A | T | A | 5,91 | 0.00 | 0.00 | 0.00 | 0.00 | yes | none |
| 2016 | CA | 29488127 | 2015-2016T>G | T | G | 7,25 | 0.00 | 0.00 | 0.00 | 0.00 | yes | none |
| 2025 | DD | 29488136 | TT>GT:2026 | T | G | 6,69 | 0.81 | 0.00 | 0.87 | 0.00 | yes | PE2025, positive control 2, 17.13% ir |
| 2050 | DD | 29488161 | AT>GT:2051 | A | G | 5,8 | 0.01 | 0.00 | 0.01 | 0.00 | yes | none |
| 2053 | DA | 29488164 | AT>AG:2053 | T | G | 3,98 | 0.00 | 0.00 | 0.00 | 0.00 | yes | none |
| 2055 | DD | 29488166 | TT>GT:2056 | T | G | 6,56 | 0.03 | 0.00 | 0.06 | 0.00 | yes | none |
| 2057 | DD | 29488168 | AT>GT:2058 | A | G | 6,59 | 0.04 | 0.00 | 0.01 | 0.00 | yes, in combination with 2055T>G | none |
| 2059 | CA | 29488170 | 2074-2059G>T | G | T | 3,75 | 0.00 | 0.00 | 0.00 | 0.00 | yes | none |
| 2060 | CA | 29488171 | 2074-2060A>T | A | T | 6,48 | 0.01 | 0.00 | 0.00 | 0.00 | yes | none |
| 2065 | CA | 29488176 | 2074-2065A>G | A | G | 3,99 | 0.00 | 0.00 | 0.00 | 0.00 | yes | none |
| 2065 | CA | 29488176 | 2074-2065A>T | A | T | 4,92 | 0.00 | 0.00 | 0.00 | 0.00 | yes | none |

**Library 10**

| intronic position | type | chrom. position | characteristics | ref | alt | MaxEnt score | SpliceAI prediction | | | | acquired | effect on splicing |
|---|---|---|---|---|---|---|---|---|---|---|---|---|
| | | | | | | | acceptor gain | acceptor loss | donor gain | donor loss | | |
| 2084 | DD | 29488195 | GA>GT:2084 | A | T | 7,27 | 0.01 | 0.00 | 0.02 | 0.00 | yes | none |
| 2088 | DD | 29488199 | TT>GT:2089 | T | G | 10,44 | 0.30 | 0.00 | 0.38 | 0.00 | yes | variant didn't meet set criteria, not included in the study, validated by fragment analysis as PE activating, used for SpliceAI evaluation |
| 2105 | DA | 29488216 | TG>AG:2106 | T | A | 4,4 | 0.00 | 0.00 | 0.00 | 0.00 | yes | none |
| 2110 | CD | 29488221 | 2107-2110A>G | A | G | 7,21 | 0.05 | 0.00 | 0.07 | 0.00 | yes | none |
| 2111 | DA | 29488222 | AT>AG:2111 | T | G | 6,56 | 0.00 | 0.00 | 0.00 | 0.00 | yes | none |
| 2113 | DA | 29488224 | AT>AG:2113 | T | G | 8,92 | 0.00 | 0.00 | 0.00 | 0.00 | yes | none |
| 2145 | CA | 29488256 | 2160-2145G>A | G | A | 4,2 | 0.00 | 0.00 | 0.00 | 0.00 | no | N/A |
| 2145 | CA | 29488256 | 2160-2145G>T | G | T | 5,56 | 0.00 | 0.00 | 0.00 | 0.00 | no | N/A |
| 2154 | CA | 29488265 | 2160-2154T>G | T | G | 3,55 | 0.00 | 0.00 | 0.00 | 0.00 | yes | none |
| 2156 | CA | 29488267 | 2160-2156G>T | G | T | 7,85 | 0.00 | 0.00 | 0.00 | 0.00 | yes | none |
| 2163 | DD | 29488275 | TT>GT:2164 | T | G | 8,1 | 0.00 | 0.00 | 0.00 | 0.00 | yes | none |
| 2180 | DA | 29488291 | TG>AG:2181 | T | A | 4,95 | 0.00 | 0.00 | 0.00 | 0.00 | yes | none |
| 2181 | DA | 29488292 | GG>AG:2182 | G | A | 6,31 | 0.00 | 0.00 | 0.00 | 0.00 | yes | none |
| 2184 | CD | 29488295 | 2183-2184T>A | T | A | 8,72 | 0.00 | 0.00 | 0.00 | 0.00 | yes | none |
| 2184 | CD | 29488295 | 2183-2184T>G | T | G | 7,43 | 0.00 | 0.00 | 0.00 | 0.00 | yes | none |
| 2214 | CD | 29488325 | 2216-2214T>G | T | G | 6,38 | 0.00 | 0.00 | 0.00 | 0.00 | yes | none |
| 2219 | CD | 29488330 | 2216-2219A>G | A | G | 7,8 | 0.00 | 0.00 | 0.00 | 0.00 | yes | none |
| 2229 | CD | 29488340 | 2228-2229T>A | T | A | 7,51 | 0.00 | 0.00 | 0.00 | 0.00 | yes | none |
| 2231 | CD | 29488342 | 2228-2231T>G | T | G | 7,44 | 0.00 | 0.00 | 0.00 | 0.00 | yes | none |
| 2240 | CD | 29488351 | 2239-2240C>A | C | A | 6,97 | 0.00 | 0.00 | 0.00 | 0.00 | yes | none |
| 2247 | CD | 29488358 | 2249-2247T>G | T | G | 5,9 | 0.00 | 0.00 | 0.00 | 0.00 | yes | none |
| 2252 | DA | 29488363 | TG>AG:2253 | T | A | 3,76 | 0.00 | 0.00 | 0.00 | 0.00 | yes | none |
| 2257 | DD | 29488368 | CT>GT:2258 | C | G | 5,87 | 0.00 | 0.00 | 0.00 | 0.00 | yes | none |
| 2260 | DA | 29488371 | AT>AG:2260 | T | G | 10,41 | 0.00 | 0.00 | 0.00 | 0.00 | yes | none |
| 2260 | DA | 29488371 | TG>AG:2261 | T | A | 4,25 | 0.00 | 0.00 | 0.00 | 0.00 | yes | none |
| 2278 | DA | 29488389 | AT>AG:2278 | T | G | 4,76 | 0.00 | 0.00 | 0.00 | 0.00 | yes | none |
| 2282 | DD | 29488393 | GA>GT:2282 | A | T | 7 | 0.00 | 0.00 | 0.00 | 0.00 | yes | none |
| 2284 | CA | 29488395 | 2299-2284A>C | A | C | 4,56 | 0.00 | 0.00 | 0.00 | 0.00 | yes | none |
| 2284 | CA | 29488395 | 2299-2284A>T | A | T | 4,79 | 0.00 | 0.00 | 0.00 | 0.00 | yes | none |

**Library 11**

| intronic position | type | chrom. position | characteristics | ref | alt | MaxEnt score | SpliceAI prediction | | | | acquired | effect on splicing |
|---|---|---|---|---|---|---|---|---|---|---|---|---|
| | | | | | | | acceptor gain | acceptor loss | donor gain | donor loss | | |
| 2292 | CA | 29488403 | 2299-2292G>C | G | C | 3,5 | 0.00 | 0.00 | 0.00 | 0.00 | no | N/A |
| 2292 | CA | 29488403 | 2299-2292G>T | G | T | 3,6 | 0.00 | 0.00 | 0.00 | 0.00 | no | N/A |
| 2300 | DD | 29488411 | TT>GT:2301 | T | G | 8,68 | 0.02 | 0.00 | 0.02 | 0.00 | no | N/A |
| 2338 | DA | 29488449 | TG>AG:2339 | T | A | 4,08 | 0.00 | 0.00 | 0.00 | 0.00 | yes | none |
| 2345 | DA | 29488456 | TG>AG:2346 | T | A | 5 | 0.00 | 0.00 | 0.00 | 0.00 | no | N/A |
| 2350 | DA | 29488461 | AT>AG:2350 | T | G | 7,71 | 0.00 | 0.00 | 0.00 | 0.00 | yes | none |
| 2353 | DA | 29488464 | TG>AG:2354 | T | A | 7,59 | 0.00 | 0.00 | 0.00 | 0.00 | yes | none |
| 2378 | CA | 29488489 | 2387-2378C>T | C | T | 3,51 | 0.00 | 0.00 | 0.00 | 0.00 | yes | none |
| 2381 | DD | 29488492 | GC>GT:2381 | C | T | 5,87 | 0.00 | 0.00 | 0.00 | 0.00 | yes | none |
| 2383 | DA | 29488494 | AT>AG:2383 | T | G | 9,21 | 0.00 | 0.00 | 0.00 | 0.00 | yes | none |
| 2383 | CD | 29488494 | 2385-2383T>G | T | G | 6,3 | 0.00 | 0.00 | 0.00 | 0.00 | yes | none |
| 2388 | CA | 29488499 | 2387-2388T>C | T | C | 4,08 | 0.00 | 0.00 | 0.00 | 0.00 | yes | none |
| 2388 | CA | 29488499 | 2387-2388T>A | T | A | 5,1 | 0.00 | 0.00 | 0.00 | 0.00 | yes | none |
| 2388 | CA | 29488499 | 2387-2388T>G | T | G | 6,21 | 0.00 | 0.00 | 0.00 | 0.00 | yes | none |
| 2400 | DD | 29488511 | TT>GT:2401 | T | G | 8,37 | 0.00 | 0.00 | 0.00 | 0.00 | no | N/A |
| 2406 | CD | 29488517 | 2405-2406G>A | G | A | 6,02 | 0.00 | 0.00 | 0.00 | 0.00 | no | N/A |
| 2408 | CD | 29488519 | 2405-2408A>G | A | G | 6,95 | 0.00 | 0.00 | 0.00 | 0.00 | yes | none |
| 2459 | CD | 29488570 | 2457-2459T>A | T | A | 7,35 | 0.00 | 0.00 | 0.00 | 0.00 | yes | none |
| 2461 | DD | 29488572 | TT>GT:2462 | T | G | 5,8 | 0.00 | 0.00 | 0.00 | 0.00 | yes | none |
| 2464 | DA | 29488575 | AA>AG:2464 | A | G | 3,77 | 0.00 | 0.00 | 0.00 | 0.00 | yes | none |
| 2466 | DA | 29488577 | TG>AG:2467 | T | A | 3,53 | 0.00 | 0.00 | 0.00 | 0.00 | yes | none |
| 2470 | DA | 29488581 | AA>AG:2470 | A | G | 7,24 | 0.00 | 0.00 | 0.00 | 0.00 | yes | none |
| 2471 | DA | 29488582 | AA>AG:2471 | A | G | 4,12 | 0.00 | 0.00 | 0.00 | 0.00 | yes | none |
| 2471 | CD | 29488582 | 2473-2471A>G | A | G | 7,84 | 0.00 | 0.00 | 0.00 | 0.00 | yes | none |
| 2476 | CD | 29488587 | 2473-2476A>G | A | G | 6,93 | 0.00 | 0.00 | 0.00 | 0.00 | yes | none |
| 2486 | CD | 29488597 | 2488-2486A>G | A | G | 6,38 | 0.00 | 0.00 | 0.00 | 0.00 | yes | none |

**Library 12**

| intronic position | type | chrom. position | characteristics | ref | alt | MaxEnt score | SpliceAI prediction | | | | acquired | effect on splicing |
|---|---|---|---|---|---|---|---|---|---|---|---|---|
| | | | | | | | acceptor gain | acceptor loss | donor gain | donor loss | | |
| 2514 | DD | 29488625 | TT>GT:2515 | T | G | 7,22 | 0.00 | 0.00 | 0.00 | 0.00 | yes | none |
| 2519 | CA | 29488630 | 2538-2519A>G | A | G | 6,04 | 0.00 | 0.00 | 0.00 | 0.00 | yes | none |
| 2519 | CA | 29488630 | 2538-2519A>C | A | C | 6,29 | 0.00 | 0.00 | 0.00 | 0.00 | yes | none |
| 2524 | DA | 29488635 | TG>AG:2525 | T | A | 3,43 | 0.00 | 0.00 | 0.00 | 0.00 | yes | none |
| 2532 | CA | 29488643 | 2538-2532A>C | A | C | 6,98 | 0.00 | 0.00 | 0.00 | 0.00 | yes | none |
| 2532 | CA | 29488643 | 2538-2532A>T | A | T | 7,54 | 0.00 | 0.00 | 0.00 | 0.00 | yes | none |
| 2551 | DD | 29488662 | TT>GT:2552 | T | G | 6,35 | 0.00 | 0.00 | 0.00 | 0.00 | yes | none |
| 2554 | DA | 29488665 | AT>AG:2554 | T | G | 4,03 | 0.00 | 0.00 | 0.00 | 0.00 | yes | none |
| 2562 | DA | 29488673 | AT>AG:2562 | T | G | 8,69 | 0.00 | 0.00 | 0.00 | 0.00 | yes | none |
| 2562 | DA | 29488673 | TG>AG:2563 | T | A | 7,02 | 0.00 | 0.00 | 0.00 | 0.00 | yes | none |
| 2564 | DA | 29488675 | GG>AG:2564 | G | A | 6,77 | 0.00 | 0.00 | 0.00 | 0.00 | no | N/A |
| 2566 | CD | 29488677 | 2565-2566G>A | G | A | 6,56 | 0.00 | 0.00 | 0.00 | 0.00 | yes | none |
| 2567 | CD | 29488678 | 2565-2567T>A | T | A | 7,23 | 0.00 | 0.00 | 0.00 | 0.00 | yes | none |
| 2570 | CD | 29488681 | 2572-2570T>G | T | G | 7,96 | 0.00 | 0.00 | 0.00 | 0.00 | yes | none |
| 2579 | DA | 29488690 | AA>AG:2579 | A | G | 3,42 | 0.00 | 0.00 | 0.00 | 0.00 | yes | none |
| 2584 | DA | 29488695 | AT>AG:2584 | T | G | 4 | 0.00 | 0.00 | 0.00 | 0.00 | yes | none |
| 2590 | CD | 29488701 | 2587-2590A>G | A | G | 6,97 | 0.00 | 0.00 | 0.00 | 0.00 | yes | none |
| 2601 | DA | 29488712 | TG>AG:2602 | T | A | 3,82 | 0.00 | 0.00 | 0.00 | 0.00 | no | N/A |
| 2608 | DA | 29488719 | AC>AG:2608 | C | G | 3,48 | 0.00 | 0.00 | 0.00 | 0.00 | yes | none |
| 2613 | DA | 29488724 | AA>AG:2613 | A | G | 3,59 | 0.00 | 0.00 | 0.00 | 0.00 | no | N/A |
| 2618 | DA | 29488729 | AA>AG:2618 | A | G | 4,88 | 0.00 | 0.00 | 0.00 | 0.00 | no | N/A |
| 2642 | CA | 29488753 | 2650-2642A>G | A | G | 6,96 | 0.02 | 0.00 | 0.02 | 0.00 | yes | none |
| 2642 | CA | 29488753 | 2650-2642A>T | A | T | 7,84 | 0.07 | 0.00 | 0.08 | 0.00 | yes | none |
| 2643 | CA | 29488754 | 2650-2643G>C | G | C | 8,64 | 0.05 | 0.00 | 0.06 | 0.00 | yes | none |
| 2643 | CA | 29488754 | 2650-2643G>T | G | T | 9,42 | 0.12 | 0.00 | 0.16 | 0.00 | yes | none |

**Library 13**

| intronic position | type | chrom. position | characteristics | ref | alt | MaxEnt score | SpliceAI prediction acceptor gain | acceptor loss | donor gain | donor loss | acquired | effect on splicing |
|---|---|---|---|---|---|---|---|---|---|---|---|---|
| 2720 | DD | 29488831 | GA>GT:2720 | A | T | 8,02 | 0.00 | 0.00 | 0.00 | 0.00 | yes | none |
| 2727 | CD | 29488838 | 2731-2727G>C | G | C | 7,2 | 0.00 | 0.00 | 0.01 | 0.00 | yes | none |
| 2727 | CD | 29488838 | 2731-2727G>A | G | A | 6,23 | 0.00 | 0.00 | 0.01 | 0.00 | yes | none |
| 2732 | CD | 29488843 | 2731-2732C>A | C | A | 10,28 | 0.36 | 0.00 | 0.43 | 0.00 | yes | none |
| 2732 | CD | 29488843 | 2731-2732C>G | C | G | 8,41 | 0.11 | 0.00 | 0.15 | 0.00 | yes | none |
| 2735 | CD | 29488846 | 2731-2735G>T | G | T | 7,7 | 0.12 | 0.00 | 0.22 | 0.00 | yes | none |
| 2735 | CD | 29488846 | 2731-2735G>C | G | C | 5,66 | 0.00 | 0.00 | 0.01 | 0.00 | yes | none |
| 2736 | DD | 29488847 | GA>GT:2736 | A | T | 7,91 | 0.01 | 0.00 | 0.02 | 0.00 | yes | none |
| 2802 | DD | 29488913 | AT>GT:2803 | A | G | 6,34 | 0.00 | 0.00 | 0.00 | 0.00 | yes | none |
| 2810 | CA | 29488921 | 2823-2810G>T | G | T | 4,35 | 0.00 | 0.00 | 0.00 | 0.00 | yes | none |
| 2812 | CA | 29488923 | 2823-2812A>C | A | C | 3,51 | 0.00 | 0.00 | 0.00 | 0.00 | yes | none |
| 2814 | CA | 29488925 | 2823-2814A>C | A | C | 3,64 | 0.00 | 0.00 | 0.00 | 0.00 | yes | none |
| 2815 | CD | 29488926 | 2817-2815T>G | T | G | 6,84 | 0.00 | 0.00 | 0.00 | 0.00 | yes | none |
| 2818 | CD | 29488929 | 2817-2818C>A | C | A | 5,71 | 0.00 | 0.00 | 0.00 | 0.00 | yes | none |
| 2822 | CA | 29488933 | 2829-2822A>T | A | T | 3,68 | 0.00 | 0.00 | 0.00 | 0.00 | yes | none |
| 2822 | CA | 29488933 | 2829-2822A>C | A | C | 4,07 | 0.00 | 0.00 | 0.00 | 0.00 | yes | none |
| 2823 | CA | 29488934 | 2829-2823G>C | G | C | 3,6 | 0.00 | 0.00 | 0.00 | 0.00 | yes | none |
| 2823 | CA | 29488934 | 2829-2823G>T | G | T | 3,75 | 0.00 | 0.00 | 0.00 | 0.00 | insufficient read depth | none |
| 2824 | CA | 29488935 | 2823-2824T>G | T | G | 4,21 | 0.00 | 0.00 | 0.00 | 0.00 | insufficient read depth | none |
| 2830 | DD | 29488941 | CT>GT:2831 | C | G | 8,17 | 0.00 | 0.00 | 0.00 | 0.00 | no | N/A |
| 2838 | DD | 29488949 | GG>GT:2838 | G | T | 5,85 | 0.00 | 0.00 | 0.00 | 0.00 | insufficient read depth | none |
| 2842 | DD | 29488953 | CT>GT:2843 | C | G | 7,13 | 0.00 | 0.00 | 0.00 | 0.00 | insufficient read depth | none |
| 2848 | DD | 29488959 | GC>GT:2848 | C | T | 8,24 | 0.00 | 0.00 | 0.00 | 0.00 | yes | none |
| 2881 | CD | 29488992 | 2880-2881T>A | T | A | 8,24 | 0.00 | 0.00 | 0.00 | 0.00 | yes | none |
| 2884 | CD | 29488995 | 2880-2884A>T | A | T | 6,36 | 0.00 | 0.00 | 0.00 | 0.00 | yes | none |
| 2884 | CD | 29488995 | 2886-2884A>G | A | G | 5,97 | 0.00 | 0.00 | 0.00 | 0.00 | yes | none |
| 2890 | CD | 29489001 | 2886-2890C>T | C | T | 6,05 | 0.00 | 0.00 | 0.00 | 0.00 | yes | none |
| 2913 | CA | 29489024 | 2926-2913A>C | A | C | 4,39 | 0.00 | 0.00 | 0.00 | 0.00 | no | N/A |
| 2913 | CA | 29489024 | 2926-2913A>G | A | G | 4,58 | 0.00 | 0.00 | 0.00 | 0.00 | insufficient read depth | none |

**Library 13** **continued**

| intronic position | type | chrom. position | characteristics | ref | alt | MaxEnt score | SpliceAI prediction: acceptor gain | SpliceAI prediction: acceptor loss | SpliceAI prediction: donor gain | SpliceAI prediction: donor loss | acquired | effect on splicing |
|---|---|---|---|---|---|---|---|---|---|---|---|---|
| 2913 | CA | 29489024 | 2926-2913A>T | A | T | 5,11 | 0.00 | 0.00 | 0.00 | 0.00 | insufficient read depth | none |
| 2914 | CA | 29489025 | 2926-2914G>C | G | C | 3,61 | 0.00 | 0.00 | 0.00 | 0.00 | insufficient read depth | none |

**Library 14**

| intronic position | type | chrom. position | characteristics | ref | alt | MaxEnt score | SpliceAI prediction: acceptor gain | SpliceAI prediction: acceptor loss | SpliceAI prediction: donor gain | SpliceAI prediction: donor loss | acquired | effect on splicing |
|---|---|---|---|---|---|---|---|---|---|---|---|---|
| 2984 | CD | 29489095 | 2986-2984A>G | A | G | 7,53 | 0.00 | 0.00 | 0.00 | 0.00 | no | N/A |
| 2989 | CD | 29489100 | 2986-2989C>G | C | G | 9,49 | 0.00 | 0.00 | 0.01 | 0.00 | yes | none |
| 3007 | DD | 29489118 | CT>GT:3008 | C | G | 7,69 | 0.00 | 0.00 | 0.00 | 0.00 | yes | none |
| 3016 | DA | 29489127 | TG>AG:3017 | T | A | 4,22 | 0.00 | 0.00 | 0.00 | 0.00 | yes | none |
| 3017 | DA | 29489128 | GG>AG:3018 | G | A | 3,53 | 0.00 | 0.00 | 0.00 | 0.00 | yes | none |
| 3019 | DD | 29489130 | GA>GT:3019 | A | T | 6,06 | 0.00 | 0.00 | 0.00 | 0.00 | yes | none |
| 3037 | DD | 29489148 | AT>GT:3038 | A | G | 7 | 0.00 | 0.00 | 0.00 | 0.00 | yes | none |
| 3050 | CD | 29489161 | 3052-3050T>G | T | G | 10,48 | 0.00 | 0.00 | 0.02 | 0.00 | yes | none |
| 3050 | CD | 29489161 | 3052-3050T>C | T | C | 6,06 | 0.00 | 0.00 | 0.00 | 0.00 | yes | none |
| 3054 | CA | 29489165 | 3059-3054A>C | A | C | 4,06 | 0.00 | 0.00 | 0.00 | 0.00 | yes | none |
| 3054 | CA | 29489165 | 3059-3054A>T | A | T | 5,23 | 0.00 | 0.00 | 0.00 | 0.00 | yes | none |
| 3055 | CA | 29489166 | 3059-3055G>T | G | T | 4,24 | 0.00 | 0.00 | 0.00 | 0.00 | yes | none |
| 3056 | CD | 29489167 | 3052-3056G>T | G | T | 7,96 | 0.00 | 0.00 | 0.01 | 0.00 | yes | none |
| 3057 | DD | 29489168 | GC>GT:3057 | C | T | 6,77 | 0.00 | 0.00 | 0.01 | 0.00 | yes | none |
| 3081 | DD | 29489192 | GG>GT:3081 | G | T | 6,41 | 0.00 | 0.00 | 0.00 | 0.00 | yes | none |
| 3084 | DD | 29489195 | AT>GT:3085 | A | G | 6,42 | 0.00 | 0.00 | 0.00 | 0.00 | yes | none |
| 3097 | CD | 29489208 | 3094-3097A>G | A | G | 8,95 | 0.00 | 0.00 | 0.03 | 0.00 | yes | none |

**Library 15**

| intronic position | type | chrom. position | characteristics | ref | alt | MaxEnt score | SpliceAI prediction | | | | acquired | effect on splicing |
|---|---|---|---|---|---|---|---|---|---|---|---|---|
| | | | | | | | acceptor gain | acceptor loss | donor gain | donor loss | | |
| 3138 | DA | 29489249 | AC>AG:3138 | C | G | 4,15 | 0.00 | 0.00 | 0.00 | 0.00 | yes | none |
| 3144 | DD | 29489255 | GG>GT:3144 | G | T | 5,85 | 0.00 | 0.00 | 0.00 | 0.00 | yes | none |
| 3154 | DD | 29489265 | GA>GT:3154 | A | T | 6,32 | 0.00 | 0.00 | 0.00 | 0.00 | no | N/A |
| 3166 | DD | 29489277 | TT>GT:3167 | T | G | 5,78 | 0.00 | 0.00 | 0.00 | 0.00 | yes | none |
| 3168 | CD | 29489279 | 3172-3168G>C | G | C | 7,2 | 0.00 | 0.00 | 0.00 | 0.00 | no | N/A |
| 3168 | CD | 29489279 | 3172-3168G>A | G | A | 6,23 | 0.00 | 0.00 | 0.00 | 0.00 | no | N/A |
| 3173 | CD | 29489284 | 3172-3173C>A | C | A | 10,28 | 0.00 | 0.00 | 0.02 | 0.00 | yes | none |
| 3173 | CD | 29489284 | 3172-3173C>G | C | G | 8,41 | 0.00 | 0.00 | 0.00 | 0.00 | yes | none |
| 3176 | CD | 29489287 | 3172-3176G>T | G | T | 7,7 | 0.00 | 0.00 | 0.00 | 0.00 | yes | none |
| 3176 | CD | 29489287 | 3172-3176G>C | G | C | 5,66 | 0.00 | 0.00 | 0.00 | 0.00 | yes | none |
| 3223 | DA | 29489334 | AA>AG:3223 | A | G | 4,03 | 0.00 | 0.00 | 0.00 | 0.00 | yes | none |
| 3229 | DA | 29489340 | AC>AG:3229 | C | G | 3,83 | 0.00 | 0.00 | 0.00 | 0.00 | yes | none |
| 3244 | DD | 29489355 | GC>GT:3244 | C | T | 6,23 | 0.00 | 0.00 | 0.00 | 0.00 | yes | none |
| 3269 | CA | 29489380 | 3277-3269A>G | A | G | 3,8 | 0.00 | 0.00 | 0.00 | 0.00 | no | N/A |
| 3269 | CA | 29489380 | 3277-3269A>C | A | C | 5,25 | 0.00 | 0.00 | 0.00 | 0.00 | yes, in combination with 3149C>T (variant not intended) | none |
| 3269 | CA | 29489380 | 3277-3269A>T | A | T | 5,69 | 0.00 | 0.00 | 0.00 | 0.00 | yes | none |
| 3270 | CA | 29489381 | 3277-3270G>C | G | C | 4,77 | 0.00 | 0.00 | 0.00 | 0.00 | no | N/A |
| 3271 | DD | 29489382 | CT>GT:3272 | C | G | 7,26 | 0.00 | 0.00 | 0.00 | 0.00 | no | N/A |
| 3279 | DD | 29489390 | GG>GT:3279 | G | T | 7,72 | 0.00 | 0.00 | 0.00 | 0.00 | yes | none |
| 3283 | DD | 29489394 | CT>GT:3284 | C | G | 7,13 | 0.00 | 0.00 | 0.00 | 0.00 | yes | none |
| 3289 | DD | 29489400 | GC>GT:3289 | C | T | 8,24 | 0.00 | 0.00 | 0.00 | 0.00 | yes | none |
| 3293 | DD | 29489404 | GA>GT:3293 | A | T | 6,6 | 0.00 | 0.00 | 0.00 | 0.00 | yes | none |
| 3311 | CA | 29489422 | 3313-3311G>T | G | T | 6,32 | 0.00 | 0.00 | 0.00 | 0.00 | yes | none |
| 3311 | CA | 29489422 | 3313-3311G>C | G | C | 7,38 | 0.00 | 0.00 | 0.00 | 0.00 | yes | none |
| 3310 | DA | 29489421 | GG>AG:3311 | G | A | 3,66 | 0.00 | 0.00 | 0.00 | 0.00 | yes | none |
| 3317 | DD | 29489428 | GG>GT:3317 | G | T | 8,07 | 0.00 | 0.00 | 0.00 | 0.00 | yes | none |
| 3325 | CD | 29489436 | 3327-3325A>G | A | G | 8,02 | 0.00 | 0.00 | 0.00 | 0.00 | yes | none |
| 3331 | CD | 29489442 | 3327-3331C>T | C | T | 7,39 | 0.00 | 0.00 | 0.00 | 0.00 | yes | none |
| 3331 | DD | 29489442 | CT>GT:3332 | C | G | 7,66 | 0.00 | 0.00 | 0.01 | 0.00 | yes | none |

**Library 16**

| intronic position | type | chrom. position | characteristics | ref | alt | MaxEnt score | SpliceAI prediction: acceptor gain | SpliceAI prediction: acceptor loss | SpliceAI prediction: donor gain | SpliceAI prediction: donor loss | acquired | effect on splicing |
|---|---|---|---|---|---|---|---|---|---|---|---|---|
| 3347 | CA | 29489458 | 3352-3347G>T | G | T | 5,13 | 0.00 | 0.00 | 0.00 | 0.00 | yes | none |
| 3347 | CA | 29489458 | 3352-3347G>C | G | C | 5,99 | 0.00 | 0.00 | 0.00 | 0.00 | yes | none |
| 3348 | CA | 29489459 | 3352-3348C>T | C | T | 3,7 | 0.00 | 0.00 | 0.00 | 0.00 | yes | none |
| 3349 | CA | 29489460 | 3352-3349A>G | A | G | 4,12 | 0.00 | 0.00 | 0.00 | 0.00 | yes | none |
| 3350 | DA | 29489461 | AT>AG:3350 | T | G | 6,37 | 0.00 | 0.00 | 0.00 | 0.00 | yes | none |
| 3363 | DD | 29489474 | CT>GT:3364 | C | G | 6,6 | 0.00 | 0.00 | 0.00 | 0.00 | yes | none |
| 3369 | DD | 29489480 | GA>GT:3369 | A | T | 8,92 | 0.00 | 0.00 | 0.03 | 0.00 | yes | none |
| 3386 | DA | 29489497 | CG>AG:3387 | C | A | 4,55 | 0.00 | 0.00 | 0.01 | 0.00 | yes | none |
| 3392 | CD | 29489503 | 3389-3392A>G | A | G | 9,1 | 0.00 | 0.00 | 0.05 | 0.00 | yes | none |
| 3399 | DA | 29489510 | TG>AG:3400 | T | A | 5,85 | 0.00 | 0.00 | 0.00 | 0.00 | yes | none |
| 3400 | DA | 29489511 | GG>AG:3401 | G | A | 3,97 | 0.00 | 0.00 | 0.00 | 0.00 | yes | none |
| 3409 | DA | 29489520 | AT>AG:3409 | T | G | 4,09 | 0.00 | 0.00 | 0.00 | 0.00 | yes | none |
| 3410 | DA | 29489521 | TG>AG:3411 | T | A | 7,49 | 0.00 | 0.00 | 0.00 | 0.00 | yes | none |
| 3489 | DD | 29489600 | GG>GT:3489 | G | T | 8,83 | 0.00 | 0.00 | 0.02 | 0.00 | no | N/A |
| 3507 | CD | 29489618 | 3510-3507T>C | T | C | 5,93 | 0.00 | 0.00 | 0.00 | 0.00 | yes | none |
| 3507 | DA | 29489618 | TG>AG:3508 | T | A | 7,47 | 0.00 | 0.00 | 0.00 | 0.00 | yes | none |
| 3508 | DA | 29489619 | GG>AG:3509 | G | A | 7,85 | 0.00 | 0.00 | 0.00 | 0.00 | yes | none |
| 3511 | CD | 29489622 | 3510-3511G>A | G | A | 7,12 | 0.00 | 0.00 | 0.00 | 0.00 | yes | none |
| 3512 | CD | 29489623 | 3510-3512T>A | T | A | 9,27 | 0.00 | 0.00 | 0.00 | 0.00 | yes | none |
| 3512 | CD | 29489623 | 3510-3512T>C | T | C | 6,18 | 0.00 | 0.00 | 0.00 | 0.00 | yes | none |
| 3514 | DD | 29489625 | TT>GT:3515 | T | G | 5,8 | 0.00 | 0.00 | 0.00 | 0.00 | no | N/A |
| 3516 | CA | 29489627 | 3528-3516A>C | A | C | 4,32 | 0.00 | 0.00 | 0.00 | 0.00 | no | N/A |
| 3517 | DA | 29489628 | AA>AG:3517 | A | G | 5,92 | 0.00 | 0.00 | 0.00 | 0.00 | yes, in combination with 3508G>A | none |
| 3518 | DA | 29489629 | AT>AG:3518 | T | G | 5,18 | 0.00 | 0.00 | 0.00 | 0.00 | no | N/A |
| 3519 | DA | 29489630 | TG>AG:3520 | T | A | 5,15 | 0.00 | 0.00 | 0.00 | 0.00 | yes | none |
| 3522 | CA | 29489633 | 3528-3522A>G | A | G | 3,63 | 0.00 | 0.00 | 0.00 | 0.00 | yes | none |
| 3523 | DA | 29489634 | AT>AG:3523 | T | G | 5,53 | 0.00 | 0.00 | 0.00 | 0.00 | yes | none |
| 3524 | DA | 29489635 | TG>AG:3525 | T | A | 3,53 | 0.00 | 0.00 | 0.00 | 0.00 | yes | none |
| 3526 | CA | 29489637 | 3528-3526A>T | A | T | 5,61 | 0.00 | 0.00 | 0.00 | 0.00 | yes | none |
| 3526 | CA | 29489637 | 3528-3526A>C | A | C | 6,76 | 0.00 | 0.00 | 0.00 | 0.00 | yes | none |
| 3529 | DD | 29489640 | TT>GT:3530 | T | G | 7,87 | 0.00 | 0.00 | 0.00 | 0.00 | yes | none |

**Library 17**

| intronic position | type | chrom. position | characteristics | ref | alt | MaxEnt score | SpliceAI prediction: acceptor gain | acceptor loss | donor gain | donor loss | acquired | effect on splicing |
|---|---|---|---|---|---|---|---|---|---|---|---|---|
| 3565 | DD | 29489676 | GC>GT:3565 | C | T | 7,87 | 0.00 | 0.00 | 0.00 | 0.00 | yes | none |
| 3577 | CD | 29489688 | 3579-3577A>G | A | G | 6,35 | 0.00 | 0.00 | 0.00 | 0.00 | yes | none |
| 3597 | DA | 29489708 | AC>AG:3597 | C | G | 6,09 | 0.01 | 0.00 | 0.00 | 0.00 | yes | none |
| 3601 | CA | 29489712 | 3606-3601G>C | G | C | 3,68 | 0.00 | 0.00 | 0.00 | 0.00 | yes | none |
| 3601 | CA | 29489712 | 3606-3601G>T | G | T | 4,03 | 0.01 | 0.00 | 0.00 | 0.00 | yes | none |
| 3600 | DA | 29489711 | TG>AG:3601 | T | A | 3,62 | 0.00 | 0.00 | 0.00 | 0.00 | yes | none |
| 3604 | CA | 29489715 | 3606-3604A>T | A | T | 4,68 | 0.01 | 0.00 | 0.00 | 0.00 | yes | none |
| 3604 | CA | 29489715 | 3606-3604A>C | A | C | 5,77 | 0.00 | 0.00 | 0.00 | 0.00 | yes | none |
| 3604 | DA | 29489715 | AA>AG:3604 | A | G | 3,95 | 0.00 | 0.00 | 0.00 | 0.00 | yes | none |
| 3605 | DA | 29489716 | AA>AG:3605 | A | G | 5,14 | 0.00 | 0.00 | 0.00 | 0.00 | yes | none |
| 3631 | CD | 29489742 | 3635-3631T>C | T | C | 5,76 | 0.00 | 0.00 | 0.00 | 0.00 | yes | none |
| 3630 | DD | 29489741 | TT>GT:3631 | T | G | 7,94 | 0.00 | 0.00 | 0.00 | 0.00 | yes | none |
| 3633 | CD | 29489744 | 3635-3633A>G | A | G | 6,86 | 0.00 | 0.00 | 0.00 | 0.00 | yes | none |
| 3635 | CD | 29489746 | 3639-3635T>C | T | C | 6,13 | 0.00 | 0.00 | 0.00 | 0.00 | yes | none |
| 3637 | CD | 29489748 | 3635-3637G>A | G | A | 8,82 | 0.00 | 0.00 | 0.00 | 0.00 | yes, in combination with del3629 (variant not intended) | none |
| 3640 | CA | 29489751 | 3657-3640A>C | A | C | 4,2 | 0.00 | 0.00 | 0.00 | 0.00 | yes | none |
| 3641 | CA | 29489752 | 3657-3641G>A | G | A | 3,52 | 0.00 | 0.00 | 0.00 | 0.00 | yes | none |
| 3641 | CD | 29489752 | 3639-3641G>A | G | A | 7,57 | 0.00 | 0.00 | 0.00 | 0.00 | yes | none |
| 3641 | CD | 29489752 | 3639-3641G>C | G | C | 5,9 | 0.00 | 0.00 | 0.00 | 0.00 | yes | none |
| 3642 | CD | 29489753 | 3639-3642T>G | T | G | 6,53 | 0.00 | 0.00 | 0.00 | 0.00 | yes | none |
| 3650 | CD | 29489761 | 3654-3650T>C | T | C | 6,6 | 0.00 | 0.00 | 0.00 | 0.00 | yes | none |
| 3650 | CD | 29489761 | 3654-3650T>A | T | A | 6,57 | 0.00 | 0.00 | 0.00 | 0.00 | no | N/A |
| 3651 | CD | 29489762 | 3654-3651T>C | T | C | 7,03 | 0.01 | 0.00 | 0.02 | 0.00 | yes | none |
| 3651 | CD | 29489762 | 3654-3651T>A | T | A | 6,39 | 0.00 | 0.00 | 0.00 | 0.00 | yes | none |
| 3653 | CA | 29489764 | 3657-3653G>A | G | A | 3,78 | 0.00 | 0.00 | 0.01 | 0.00 | yes | none |
| 3653 | CA | 29489764 | 3657-3653G>T | G | T | 4,67 | 0.01 | 0.00 | 0.02 | 0.00 | yes | none |
| 3655 | CD | 29489766 | 3654-3655C>A | C | A | 10,47 | 0.00 | 0.00 | 0.00 | 0.00 | yes | none |
| 3655 | CD | 29489766 | 3654-3655C>G | C | G | 9,27 | 0.00 | 0.00 | 0.00 | 0.00 | yes | none |
| 3670 | DD | 29489781 | GC>GT:3670 | C | T | 9,72 | 0.00 | 0.00 | 0.00 | 0.00 | yes | none |
| 3692 | DD | 29489803 | GA>GT:3692 | A | T | 7,51 | 0.00 | 0.00 | 0.00 | 0.00 | yes | none |

**Library 17** **continued**

| intronic position | type | chrom. position | characteristics | ref | alt | MaxEnt score | SpliceAI prediction: acceptor gain | acceptor loss | donor gain | donor loss | acquired | effect on splicing |
|---|---|---|---|---|---|---|---|---|---|---|---|---|
| 3742 | DA | 29489853 | AT>AG:3742 | T | G | 5,14 | 0.00 | 0.00 | 0.00 | 0.00 | yes | none |
| 3746 | DD | 29489857 | TT>GT:3747 | T | G | 9,82 | 0.03 | 0.00 | 0.08 | 0.00 | yes | none |
| 3750 | CA | 29489861 | 3768-3750G>A | G | A | 3,62 | 0.06 | 0.00 | 0.02 | 0.00 | no | variant validated by fragment analysis for SpliceAI evaluation, no effect on splicing |
| 3750 | CA | 29489861 | 3768-3750G>C | G | C | 4,03 | 0.24 | 0.00 | 0.27 | 0.00 | no | variant validated by fragment analysis for SpliceAI evaluation as PEwt activating |
| 3753 | CA | 29489864 | 3768-3753G>C | G | C | 4,57 | 0.01 | 0.00 | 0.01 | 0.00 | yes, in combination with 3670C>T | none |
| 3758 | CD | 29489869 | 3760-3758T>G | T | G | 7,06 | 0.00 | 0.00 | 0.00 | 0.00 | yes, in combination with del3751 (variant not intended) | none |
| 3759 | CA | 29489870 | 3768-3759G>T | G | T | 5,13 | 0.34 | 0.00 | 0.32 | 0.00 | no | variant validated by fragment analysis for SpliceAI evaluation as PEwt activating |
| 3763 | CD | 29489874 | 3760-3763T>G | T | G | 7,96 | 0.00 | 0.00 | 0.00 | 0.00 | no | N/A |

**Library 18**

| intronic position | type | chrom. position | characteristics | ref | alt | MaxEnt score | SpliceAI prediction: acceptor gain | acceptor loss | donor gain | donor loss | acquired | effect on splicing |
|---|---|---|---|---|---|---|---|---|---|---|---|---|
| 3797 | DD | 29489908 | GG>GT:3797 | G | T | 11,08 | 0.00 | 0.00 | 0.06 | 0.00 | yes | non-evaluable |
| 3798 | DD | 29489909 | GA>GT:3798 | A | T | 5,76 | 0.00 | 0.00 | 0.00 | 0.00 | yes, in combination with 3818A>G | non-evaluable |
| 3815 | CD | 29489926 | 3812-3815A>G | A | G | 7,43 | 0.00 | 0.00 | 0.00 | 0.00 | yes | non-evaluable |
| 3818 | DA | 29489929 | AA>AG:3818 | A | G | 3,43 | 0.00 | 0.00 | 0.00 | 0.00 | yes | none |
| 3820 | DD | 29489931 | GA>GT:3820 | A | T | 6,39 | 0.00 | 0.00 | 0.00 | 0.00 | yes | non-evaluable |
| 3824 | DD | 29489935 | GC>GT:3824 | C | T | 6,31 | 0.00 | 0.00 | 0.00 | 0.00 | yes | non-evaluable |
| 3832 | DD | 29489943 | GA>GT:3832 | A | T | 9,99 | 0.00 | 0.00 | 0.12 | 0.00 | yes | non-evaluable |
| 3837 | DD | 29489948 | GG>GT:3837 | G | T | 6,66 | 0.00 | 0.00 | 0.01 | 0.00 | yes | non-evaluable |
| 3843 | CD | 29489954 | 3846-3843T>A | T | A | 11,08 | 0.16 | 0.00 | 0.20 | 0.00 | yes | variant validated by fragment analysis for SpliceAI evaluation as PEwt activating |
| 3843 | CD | 29489954 | 3846-3843T>C | T | C | 10,19 | 0.07 | 0.00 | 0.07 | 0.00 | yes | non-evaluable |
| 3843 | CD | 29489954 | 3846-3843T>G | T | G | 10,17 | 0.00 | 0.00 | 0.01 | 0.00 | no | N/A |
| 3850 | CD | 29489961 | 3846-3850G>T | G | T | 10,65 | 0.37 | 0.00 | 0.38 | 0.00 | yes, in combination with 3851A>T | variant validated by fragment analysis for SpliceAI evaluation as PEwt activating, tested only once |
| 3851 | DD | 29489962 | GA>GT:3851 | A | T | 6,1 | 0.00 | 0.00 | 0.01 | 0.00 | yes | non-evaluable |
| 3855 | DD | 29489966 | GC>GT:3855 | C | T | 5,88 | 0.00 | 0.00 | 0.00 | 0.00 | yes | non-evaluable |
| 3880 | CD | 29489991 | 3877-3880T>G | T | G | 7,35 | 0.00 | 0.00 | 0.01 | 0.00 | yes | non-evaluable |
| 3883 | DA | 29489994 | AT>AG:3883 | T | G | 4,51 | 0.00 | 0.00 | 0.02 | 0.00 | yes | none |
| 3884 | DA | 29489995 | GG>AG:3885 | G | A | 3,44 | 0.00 | 0.00 | 0.00 | 0.00 | yes | none |
| 3889 | CD | 29490000 | 3888-3889C>A | C | A | 7,83 | 0.00 | 0.00 | 0.00 | 0.00 | yes | non-evaluable |
| 3911 | CA | 29490022 | 3926-3911A>G | A | G | 5,27 | 0.01 | 0.00 | 0.01 | 0.00 | yes | none |
| 3911 | CA | 29490022 | 3926-3911A>T | A | T | 6,19 | 0.00 | 0.00 | 0.01 | 0.00 | yes | none |
| 3916 | CA | 29490027 | 3926-3916A>C | A | C | 5,64 | 0.00 | 0.00 | 0.00 | 0.00 | yes | none |
| 3916 | DA | 29490027 | AA>AG:3916 | A | G | 4,14 | 0.00 | 0.00 | 0.00 | 0.00 | no | N/A |
| 3920 | DA | 29490031 | AT>AG:3920 | T | G | 4,56 | 0.00 | 0.00 | 0.00 | 0.00 | yes, in combination with 3924T>G | none |
| 3923 | CA | 29490034 | 3926-3923A>T | A | T | 4,84 | 0.00 | 0.00 | 0.00 | 0.00 | no | N/A |
| 3924 | DA | 29490035 | AT>AG:3924 | T | G | 3,53 | 0.00 | 0.00 | 0.00 | 0.00 | yes | none |

**Library 19**

| intronic position | type | chrom. position | characteristics | ref | alt | MaxEnt score | SpliceAI prediction | | | | acquired | effect on splicing |
|---|---|---|---|---|---|---|---|---|---|---|---|---|
| | | | | | | | acceptor gain | acceptor loss | donor gain | donor loss | | |
| 3953 | DD | 29490064 | TT>GT:3954 | T | G | 6,72 | 0.00 | 0.00 | 0.00 | 0.00 | yes | none |
| 3958 | CA | 29490069 | 3972-3958A>G | A | G | 7,26 | 0.01 | 0.00 | 0.00 | 0.00 | yes | none |
| 3958 | CA | 29490069 | 3972-3958A>T | A | T | 9,15 | 0.01 | 0.00 | 0.01 | 0.00 | yes | none |
| 3959 | CA | 29490070 | 3972-3959G>T | G | T | 8,71 | 0.01 | 0.00 | 0.01 | 0.00 | yes | none |
| 3961 | CA | 29490072 | 3972-3961T>C | T | C | 6,22 | 0.00 | 0.00 | 0.00 | 0.00 | yes | none |
| 3974 | CD | 29490085 | 3977-3974T>A | T | A | 6,14 | 0.00 | 0.00 | 0.00 | 0.00 | yes | none |
| 3974 | CD | 29490085 | 3977-3974T>C | T | C | 5,95 | 0.00 | 0.00 | 0.00 | 0.00 | yes | none |
| 3978 | CD | 29490089 | 3977-3978G>A | G | A | 7,64 | 0.00 | 0.00 | 0.00 | 0.00 | yes | none |
| 3979 | CD | 29490090 | 3977-3979T>A | T | A | 8,95 | 0.00 | 0.00 | 0.01 | 0.00 | no | N/A |
| 3979 | CD | 29490090 | 3981-3979T>G | T | G | 7,43 | 0.00 | 0.00 | 0.00 | 0.00 | yes | none |
| 3979 | CD | 29490090 | 3977-3979T>C | T | C | 6,46 | 0.00 | 0.00 | 0.00 | 0.00 | yes | none |
| 3981 | CD | 29490092 | 3985-3981T>C | T | C | 6,57 | 0.00 | 0.00 | 0.00 | 0.00 | yes | none |
| 3981 | CD | 29490092 | 3985-3981T>G | T | G | 6,35 | 0.00 | 0.00 | 0.00 | 0.00 | no | N/A |
| 3981 | CD | 29490092 | 3985-3981T>A | T | A | 6,17 | 0.00 | 0.00 | 0.00 | 0.00 | yes | none |
| 3983 | CD | 29490094 | 3985-3983T>G | T | G | 9,31 | 0.00 | 0.00 | 0.01 | 0.00 | yes | none |
| 3983 | CD | 29490094 | 3981-3983T>A | T | A | 7,65 | 0.00 | 0.00 | 0.00 | 0.00 | yes | none |
| 3983 | CD | 29490094 | 3985-3983T>C | T | C | 6,01 | 0.00 | 0.00 | 0.00 | 0.00 | no | N/A |
| 3989 | CD | 29490100 | 3985-3989G>T | G | T | 6,69 | 0.00 | 0.00 | 0.00 | 0.00 | yes | none |
| 3991 | CD | 29490102 | 3990-3991G>A | G | A | 6,8 | 0.00 | 0.00 | 0.01 | 0.00 | yes | none |
| 3993 | CD | 29490104 | 3990-3993T>G | T | G | 6,3 | 0.00 | 0.00 | 0.00 | 0.00 | yes | none |
| 4003 | DA | 29490114 | TG>AG:4004 | T | A | 5,63 | 0.00 | 0.00 | 0.01 | 0.00 | yes | none |
| 4008 | CD | 29490119 | 4010-4008T>G | T | G | 5,93 | 0.00 | 0.00 | 0.00 | 0.00 | yes | none |
| 4008 | DA | 29490119 | TG>AG:4009 | T | A | 6,81 | 0.00 | 0.00 | 0.01 | 0.00 | yes | none |
| 4012 | CD | 29490123 | 4010-4012T>A | T | A | 7 | 0.00 | 0.00 | 0.01 | 0.00 | yes | none |
| 4014 | CD | 29490125 | 4012-4014T>A | T | A | 5,84 | 0.00 | 0.00 | 0.01 | 0.00 | yes | none |
| 4016 | CD | 29490127 | 4014-4016T>A | T | A | 5,84 | 0.00 | 0.00 | 0.01 | 0.00 | yes | none |
| 4020 | DA | 29490131 | TG>AG:4021 | T | A | 8,03 | 0.00 | 0.00 | 0.00 | 0.00 | yes | none |
| 4023 | DA | 29490134 | AA>AG:4023 | A | G | 4,79 | 0.00 | 0.00 | 0.00 | 0.00 | yes | none |
| 4024 | DA | 29490135 | AA>AG:4024 | A | G | 7,79 | 0.00 | 0.00 | 0.00 | 0.00 | yes | none |
| 4025 | DA | 29490136 | AA>AG:4025 | A | G | 7,27 | 0.00 | 0.00 | 0.00 | 0.00 | yes | none |
| 4026 | DA | 29490137 | AT>AG:4026 | T | G | 5,4 | 0.00 | 0.00 | 0.00 | 0.00 | yes | none |
| 4032 | DA | 29490143 | AT>AG:4032 | T | G | 6,31 | 0.00 | 0.00 | 0.01 | 0.00 | yes | none |

**Table S2** - A list of primers used within the study

| Primer pair | 5' -> 3' sequence | Purpose |
|---|---|---|
| NF1_inF_F2 | CCCAGGATAGGTACCGGGCCCTCTGTTTGCCTTAGACTTTAGT | Insert cloning |
| NF1_inF_R2 | CGCGGTGGCGGCCGCTCTAGATTAAAGCCTGCTCACGGCA | |
| NF1_1137T_F | TGAGCATGGTGGTACATACCTGTAATCCCAGCTAC | mutagenesis |
| NF1_1137T_R | GTAGCTGGGATTACAGGTATGTACCACCATGCTCA | |
| NF1_2025G_F | AAGTATTTTTCTTTCACTTACATATTCCAACTATGTATAACAAACGTATACTTGTGAG | mutagenesis |
| NF1_2025G_R | CTCACAAGTATACGTTTGTTATACATAGTTGGAATATGTAAGTGAAAGAAAAATACTT | |
| pET1A(F) | CAGCACCTTTGTGGTTCTCA(-FAM) | Splicing analysis of pET |
| rpET_PS02 | GCACTGATCCACGATG | Splicing analysis of pET |
| NF1_i3_part1 | TGTGGGAGAGTGCGATTCAG | Verification of the inserted *NF1* sequence |
| NF1_i3_part2 | ACCTCTGTTTTAATGATCTGCCTAA | |
| NF1_i3_part3 | TTACCAGTATATCAGCCACACCC | |
| NF1_i3_part4 | AATGTGGAACAGCATACCTACA | |
| NF1_i3_part5 | AAGTCATGGGAAATAAGAGTGGT | |
| pETseqI1f | TTTCTGTGGGCAGCTAGA | Verification of the pET01 vector sequence |
| pETseqI2f | CTGATGCTCTGCCCTGTG | |
| pETseqI1r | TATGTCTAGCTGCCCACA | |
| NF1_i3_AsiSI_1_F | GGTCCACATCTGTACTTTGGGACATGCGATCGCTCTCAAGATCAAATTCACAAAGCCT | mutagenesis - *AsiSI* restriction site substitution |
| NF1_i3_AsiSI_1_R | AGGCTTTGTGAATTTGATCTTGAGAGCGATCGCATGTCCCAAAGTACAGATGTGGACC | |
| NF1_i3_AsiSI_2_F | TCTGAGAGATCCGTTAAATCAAATAGCGATCGCTTCATAAAATATTTTCCTAAAAACC | mutagenesis - *AsiSI* restriction site substitution |
| NF1_i3_AsiSI_2_R | GGTTTTTAGGAAAATATTTTATGAAGCGATCGCTATTTGATTTAACGGATCTCTCAGA | |
| NF1_i3_AsiSI_3_F | TACTCTCCTCCTCCACATAGCGATCGCATATTTGTGTGTCT | mutagenesis - *AsiSI* restriction site substitution |
| NF1_i3_AsiSI_3_R | AGACACACAAATATGCGATCGCTATGTGGAGGAGGAGAGTA | |
| NF1_i3_AsiSI_4_F | GAGAGAACATAAGAAATGAGTTAATTTCAATACACAGCGATCGCGCAGATCATTAAAACAGAGG | mutagenesis - *AsiSI* restriction site substitution |
| NF1_i3_AsiSI_4_R | CCTCTGTTTTAATGATCTGCGCGATCGCTGTGTATTGAAATTAACTCATTTCTTATGTTCTCTC | |
| NF1_i3_AsiSI_5_F | ACATTTTTTCGCTTACAGCCTTCCCTAGCGATCGCCAGTTGTGTCAATATAATTTTTTTTTCA | mutagenesis - *AsiSI* restriction site substitution |
| NF1_i3_AsiSI_5_R | TTGAAAAAAAAATTATATTGACACAACTGGCGATCGCTAGGGAAGGCTGTAAGCGAAAAAATGT | |
| NF1_i3_AsiSI_6_F | CTAGCCTGACCAACATAGTGAACCGCGATCGCTTATGAAAATACAAAAAAAATA | mutagenesis - *AsiSI* restriction site substitution |
| NF1_i3_AsiSI_6_R | TATTTTTTTTGTATTTTCATAAGCGATCGCGGTTCACTATGTTGGTCAGGCTAG | |
| NF1_i3_AsiSI_7_F | ATATCACAAGTTAACGGAAAATACCATGCGATCGCAACTAGCTCACCTTCGATTTTAAGTAGA | mutagenesis - *AsiSI* restriction site substitution |
| NF1_i3_AsiSI_7_R | TCTACTTAAAATCGAAGGTGAGCTAGTTGCGATCGCATGGTATTTTCCGTTAACTTGTGATAT | |
| NF1_i3_AsiSI_8_F | CTGTGAAACTGAACAAGGCGATCGCTGACCCCACGCCCCACCATTCAG | mutagenesis - *AsiSI* restriction site substitution |
| NF1_i3_AsiSI_8_R | CTGAATGGTGGGGCGTGGGGTCAGCGATCGCCTTGTTCAGTTTCACAG | |

| Primer pair | 5' -> 3' sequence | Purpose |
|---|---|---|
| NF1_i3_AsiSI_9_F | GCATTACTTTTTAAATAGTAAATTTAATGAAAAAGCGATCGCATTTTTCAACAGTAGAGCACGGCAAGATAATATA | mutagenesis - *AsiSI* restriction site substitution |
| NF1_i3_AsiSI_9_R | TATATTATCTTGCCGTGCTCTACTGTTGAAAAATGCGATCGCTTTTTCATTAAATTTACTATTTAAAAAGTAATGC | |
| NF1_i3_AsiSI_10_F | ACTTACCAATATGTTATGAGTGCGATCGCGTAGAATTTGAGGAATCTTAAGTATTTTTGTTCC | mutagenesis - *AsiSI* restriction site substitution |
| NF1_i3_AsiSI_10_R | CAGGAACAAAAATACTTAAGATTCCTCAAATTCTACGCGATCGCACTCATAACATATTGGTAA | |
| NF1_i3_AsiSI_11_F | TCAATTCTGTAACTGCAATCCAAGCGATCGCAGATGCAAATACACTGAAACATAG | mutagenesis - *AsiSI* restriction site substitution |
| NF1_i3_AsiSI_11_R | CTATGTTTCAGTGTATTTGCATCTGCGATCGCTTGGATTGCAGTTACAGAATTGA | |
| NF1_i3_AsiSI_12_F | ACCAAGTTTGATAACTTCGGTGTTAACAGTGCGATCGCTACTTTGTAAATATATACTTTACAATTAAC | mutagenesis - *AsiSI* restriction site substitution |
| NF1_i3_AsiSI_12_R | GTTAATTGTAAAGTATATATTTACAAAGTAGCGATCGCACTGTTAACACCGAAGTTATCAAACTTGGT | |
| NF1_i3_AsiSI_13_F | CTCTCCGCCTGTCGCGATCGCCCAAAGTGCTGG | mutagenesis - *AsiSI* restriction site substitution |
| NF1_i3_AsiSI_13_R | CCAGCACTTTGGGCGATCGCGACAGGCGGAGAG | |
| NF1_i3_AsiSI_14_F | GAGTCTCGCTCTGTGGCGATCGCTGGAGTGCAGTGGCA | mutagenesis - *AsiSI* restriction site substitution |
| NF1_i3_AsiSI_14_R | TGCCACTGCACTCCAGCGATCGCCACAGAGCGAGACTC | |
| NF1_i3_AsiSI_15_F | CCTCCCAAAGTGCTGCGATCGCAGGCGTGAGCCACT | mutagenesis - *AsiSI* restriction site substitution |
| NF1_i3_AsiSI_15_R | AGTGGCTCACGCCTGCGATCGCAGCACTTTGGGAGG | |
| NF1_i3_AsiSI_16_F | ATTCTATGCAAGGGCGCGATCGCAGCTCACTGCAACTTC | mutagenesis - *AsiSI* restriction site substitution |
| NF1_i3_AsiSI_16_R | GAAGTTGCAGTGAGCTGCGATCGCGCCCTTGCATAGAAT | |
| NF1_i3_AsiSI_17_F | CTTCACCCCATGTGCCTTGCGATCGCTTAAAAGAATTTTCTCTAATGTAACTTCA | mutagenesis - *AsiSI* restriction site substitution |
| NF1_i3_AsiSI_17_R | TGAAGTTACATTAGAGAAAATTCTTTTAAGCGATCGCAAGGCACATGGGGTGAAG | |
| NF1_i3_AsiSI_18_F | CCCAGTTTGCTGTGAGTTCTACAATTACGCGATCGCCTTAATTCATATTGATTGATGAGTGCAT | mutagenesis - *AsiSI* restriction site substitution |
| NF1_i3_AsiSI_18_R | ATGCACTCATCAATCAATATGAATTAAGGCGATCGCGTAATTGTAGAACTCACAGCAAACTGGG | |
| NF1_i3_AsiSI_19_F | GATTTAACAACTGTTAACGTTTCATCATATTTGCGATCGCTGTCTATAGATAGTTTAATAAAGAGGATGGTTC | mutagenesis - *AsiSI* restriction site substitution |
| NF1_i3_AsiSI_19_R | GAACCATCCTCTTTATTAAACTATCTATAGACAGCGATCGCAAATATGATGAAACGTTAACAGTTGTTAAATC | |
| 6n_BC_mut_F | GCGAAGTGNNNNNNCAGATCTCGAGGAC | barcoding |
| 6n_BC_mut_R | GTCCTCGAGATCTGNNNNNNCACTTCGC | |
| fpET_PS01 | CAGGCTTTTGTCAAACAG | Verification of the barcode |
| NF1_seq_F | TCGTCGGCAGCGTCAGATGTGTATAAGAGACAGTGGGGAGCGTGGATTCT | NGS sequencing |
| NF1_seq_upr_1R | GTCTCGTGGGCTCGGAGATGTGTATAAGAGACAGCCGTTAAATCAAATAGTGATCTC | NGS sequencing with Illumina adaptors |
| NF1_seq_upr_2R | GTCTCGTGGGCTCGGAGATGTGTATAAGAGACAGACTCTCCTCCTCCACATACCT | NGS sequencing with Illumina adaptors |
| NF1_seq_upr_3R | GTCTCGTGGGCTCGGAGATGTGTATAAGAGACAGGAGTTAATTTCAATACACATG | NGS sequencing with Illumina adaptors |
| NF1_seq_upr_4R | GTCTCGTGGGCTCGGAGATGTGTATAAGAGACAGACAGCCTTCCCTAGGGATACA | NGS sequencing with Illumina adaptors |
| NF1_seq_upr_5R | GTCTCGTGGGCTCGGAGATGTGTATAAGAGACAGGACCAACATAGTGAACCCCC | NGS sequencing with Illumina adaptors |

| Primer pair | 5' -> 3' sequence | Purpose |
| --- | --- | --- |
| NF1_seq_upr_6R | TCTCGTGGGCTCGGAGATGTGTATAAGAGACAGACGGAAAATACCATGCTAAA | NGS sequencing with Illumina adaptors |
| NF1_seq_upr_7R | GTCTCGTGGGCTCGGAGATGTGTATAAGAGACAGGAAACTGAACAAGGCAATATA | NGS sequencing with Illumina adaptors |
| NF1_seq_upr_8R | GTCTCGTGGGCTCGGAGATGTGTATAAGAGACAGGTAAATTTAATGAAAAAGCATA | NGS sequencing with Illumina adaptors |
| NF1_seq_upr_9R | GTCTCGTGGGCTCGGAGATGTGTATAAGAGACAGAGATTCCTCAAATTCTACAAA | NGS sequencing with Illumina adaptors |
| NF1_seq_upr_10R | GTCTCGTGGGCTCGGAGATGTGTATAAGAGACAGTCTGTAACTGCAATCCAAGCAC | NGS sequencing with Illumina adaptors |
| NF1_seq_upr11R | GTCTCGTGGGCTCGGAGATGTGTATAAGAGACAGACTTCGGTGTTAACAGTGAA | NGS sequencing with Illumina adaptors |
| NF1_seq_upr_12R | GTCTCGTGGGCTCGGAGATGTGTATAAGAGACAGTCTCCGCCTGTCTCGATCT | NGS sequencing with Illumina adaptors |
| NF1_seq_upr_13R | GTCTCGTGGGCTCGGAGATGTGTATAAGAGACAGGAGTCTCGCTCTGTGGCCC | NGS sequencing with Illumina adaptors |
| NF1_seq_upr_14R | GTCTCGTGGGCTCGGAGATGTGTATAAGAGACAGGCCTCCCAAAGTGCTGGGATTA | NGS sequencing with Illumina adaptors |
| NF1_seq_upr_15R | GTCTCGTGGGCTCGGAGATGTGTATAAGAGACAGACTATTCTATGCAAGGGCGCA | NGS sequencing with Illumina adaptors |
| NF1_seq_upr_16R | GTCTCGTGGGCTCGGAGATGTGTATAAGAGACAGTTCACCCCATGTGCCTTGGC | NGS sequencing with Illumina adaptors |
| NF1_seq_upr_17R | GTCTCGTGGGCTCGGAGATGTGTATAAGAGACAGTGCTGTGAGTTCTACAATTACAGA | NGS sequencing with Illumina adaptors |
| NF1_seq_upr_18R | GTCTCGTGGGCTCGGAGATGTGTATAAGAGACAGACGTTTCATCATATTTGCTACA | NGS sequencing with Illumina adaptors |
| NF1_seq_upr_19R | GTCTCGTGGGCTCGGAGATGTGTATAAGAGACAGCTCATTGTGTCCTTTGGTTG | NGS sequencing with Illumina adaptors |
| fPET_PS03_ill_f.adapt<br>rPET_PS04_ill_r.adapt | TCGTCGGCAGCGTCAGATGTGTATAAGAGACAGTGGGGAGCGTGGATTCT<br>GTCTCGTGGGCTCGGAGATGTGTATAAGAGACAGCCACCTCCAGTGCCAAG | NGS sequencing of RNA libraries |
| NF1_ex4_seqR_Miseq | TGCATGCTGGTTTCCTTCACGACA | Custom reverse primer pro Miseq seqeuncing |
| NF1_ex3_F | GGATACACTGGAAAAATGTCTTGC | Sanger sequencing for PEwt verification |
| NF1_545A>G_F<br>NF1_545A>G_R | GAAATACACTGTGGTAACAACCTACAGGAAACCAGAGA<br>TCTCTGGTTTCCTGTAGGTTGTTACCACAGTGTATTTC | mutagenesis (variant FA verification) |
| NF1_594A>G_F<br>NF1_594A>G_R | CATGACTTAATTTGAACTTTTTCAGGAGTATTTCTATTGTGTAAAGA<br>TCTTTACACAATAGAAATACTCCTGAAAAAGTTCAAATTAAGTCATG | mutagenesis (variant FA verification) |

| | | |
|---|---|---|
| NF1_912G>A_F | GTGTTGGTGGAAGCATTGGGGTAAGGCATTGAAAAAAAAATT | mutagenesis (variant FA verification) |
| NF1_912G>A_R | AATTTTTTTTTCAATGCCTTACCCCAATGCTTCCACCAACAC | |
| NF1_1012A>C_F | ATGTATATATATTTTTCTTTTGCGACAGAGTCTCACTCTGCGGC | mutagenesis (variant FA verification) |
| NF1_1012A>C_R | GCCGCAGAGTGAGACTCTGTCGCAAAAGAAAAATATATATACAT | |
| NF1_1012A>T_F | ATGTATATATATTTTTCTTTTGTGACAGAGTCTCACTCTGCGGC | mutagenesis (variant FA verification) |
| NF1_1012A>T_R | GCCGCAGAGTGAGACTCTGTCACAAAAGAAAAATATATATACAT | |
| NF1_1019G>T_F | TATTTTTCTTTTGAGACAGATTCTCACTCTGCGGCAC | mutagenesis (variant FA verification) |
| NF1_1019G>T_R | GTGCCGCAGAGTGAGAATCTGTCTCAAAAGAAAAATA | |
| NF1_1156C>G_F | AGGCATGTACCACCATGCTCAGGTATTTTTTTTGTATTTTCATAAG | mutagenesis (variant FA verification) |
| NF1_1156C>G_R | CTTATGAAAATACAAAAAAAATACCTGAGCATGGTGGTACATGCCT | |
| NF1_1322T>A_F | TGCCCTAAGCATAATATTTTAAATTTTCTTATAGACCGTTGACC | mutagenesis (variant FA verification) |
| NF1_1322T>A_R | GGTCAACGGTCTATAAGAAAATTTAAAATATTATGCTTAGGGCA | |
| NF1_1677T>G_F | GTAATCATTCTCCAAAATACATTCATTCTTACCTCTCAATTACAAATGTTAATGGA | mutagenesis (variant FA verification) |
| NF1_1677T>G_R | TCCATTAACATTTGTAATTGAGAGGTAAGAATGAATGTATTTTGGAGAATGATTAC | |
| NF1_1994A>G_F | GTTTAAGAAGAACGGAGCTCACAGGTATACGTTTGTTATACATAGTT | mutagenesis (variant FA verification) |
| NF1_1994A>G_R | AACTATGTATAACAAACGTATACCTGTGAGCTCCGTTCTTCTTAAAC | |
| NF1_2088T>G_F | TGCAATTTTGTAGAATTTGAGGAATCGTAAGTATTTTTGTTCCTGTGAATATG | mutagenesis (variant FA verification) |
| NF1_2088T>G_R | CATATTCACAGGAACAAAAATACTTACGATTCCTCAAATTCTACAAAATTGCA | |
| NF1_2732C>A_F | AGGCGGAGAGCTTGAGGTAAGGAATCCAAGAC | mutagenesis (variant FA verification) |
| NF1_2732C>A_R | GTCTTGGATTCCTTACCTCAAGCTCTCCGCCT | |
| NF1_2735G>T_F | GAGAGCTTGAGGTCAGTAATCCAAGACCAGCCT | mutagenesis (variant FA verification) |
| NF1_2735G>T_R | AGGCTGGTCTTGGATTACTGACCTCAAGCTCTC | |
| NF1_3750G>A_F | CACAATGCACTCATCAATCAATATGAATTAAATCGATTCTGTAATTGT | mutagenesis (variant FA verification) |
| NF1_3750G>A_R | ACAATTACAGAATCGATTTAATTCATATTGATTGATGAGTGCATTGTG | |
| NF1_3750G>C_F | CACAATGCACTCATCAATCAATATGAATTAACTCGATTCTGTAATTGT | mutagenesis (variant FA verification) |
| NF1_3750G>C_R | ACAATTACAGAATCGAGTTAATTCATATTGATTGATGAGTGCATTGTG | |
| NF1_3759G>T_F | GAATTAAGTCGATTCTTTAATTGTAGAACTCACAGCAAACTG | mutagenesis (variant FA verification) |
| NF1_3759G>T_R | CAGTTTGCTGTGAGTTCTACAATTAAAGAATCGACTTAATTC | |
| NF1_3843T>A_F | CAGGAATGGGAGGACAGGTAAGGATGGCAGG | mutagenesis (variant FA verification) |
| NF1_3843T>A_R | CCTGCCATCCTTACCTGTCCTCCCATTCCTG | |
| NF1_3850G>T_F | GGGAGGACTGGTAAGTATGGCAGGGGGATTG | mutagenesis (variant FA verification) |
| NF1_3850G>T_R | CAATCCCCCTGCCATACTTACCAGTCCTCCC | |

**Table S3A -** Comparison Illumina and Nanopore sequencing results, library 8

| comparison without any filtering | # | % |
|---|---|---|
| matching | 376 | 97.16% |
| non-matching | 11 | 2.84% |

| Comparison after NP data filtering | | | |
|---|---|---|---|
| *condition 1:* | mut_depth > 9 | **AND** | |
| *condition 2:* | mut support > 0,75 | | |
| | **#** | **% of all** | **% of filtered** |
| matching | 333 | 86.05% | 99.40% |
| non-matching | 2 | 0.52% | 0.60% |

**Table S3B** - Comparison Illumina and Nanopore sequencing results, library 8, detailed

| Barcode | Illumina | | | | NanoPore | | | | | |
|---|---|---|---|---|---|---|---|---|---|---|
| | DNA variant (min 30 reads) | Variant reads | Total reads | | DNA variant | variant reads | total reads | mutation support | | Illumina vs NP Match |
| ACTGGG | 1637T>C | 253 | 324 | | 1637T>C | 38 | 41 | 1.000 | | 1 |
| CACATT | 1637T>C | 58 | 83 | | 1637T>C | 13 | 13 | 1.000 | | 1 |
| CAGGGG | 1637T>C | 72 | 100 | | 1637T>C | 21 | 21 | 1.000 | | 1 |
| CCTTGG | 1637T>C | 73 | 140 | | 1637T>C | 16 | 24 | 1.000 | | 1 |
| CTAGAA | 1637T>C | 34 | 47 | | 1637T>C | 12 | 16 | 0.978 | | 1 |
| CTATCT | 1637T>C | 234 | 339 | | 1637T>C | 68 | 74 | 1.000 | | 1 |
| GACCAG | 1637T>C | 94 | 131 | | 1637T>C | 23 | 24 | 1.000 | | 1 |
| GAGAAT | 1637T>C | 182 | 239 | | 1637T>C | 56 | 57 | 1.000 | | 1 |
| GGGGAT | 1637T>C | 89 | 122 | | 1637T>C | 16 | 17 | 1.000 | | 1 |
| TAGGGT | 1637T>C | 224 | 295 | | 1637T>C | 38 | 43 | 1.000 | | 1 |
| TCACAC | 1637T>C | 333 | 444 | | 1637T>C | 74 | 75 | 1.000 | | 1 |
| TCCTCC | 1637T>C | 84 | 135 | | 1637T>C | 31 | 35 | 1.000 | | 1 |
| TGATTA | 1637T>C | 421 | 577 | | 1637T>C | 110 | 114 | 1.000 | | 1 |
| TTCGAC | 1637T>C | 123 | 280 | | 1637T>C | 27 | 54 | 1.000 | | 1 |
| ACCATC | 1640C>G | 117 | 150 | | 1640C>G | 25 | 29 | 1.000 | | 1 |
| AGGGGT | 1640C>G | 245 | 322 | | 1640C>G | 56 | 60 | 1.000 | | 1 |
| ATAAGG | 1640C>G | 122 | 169 | | 1640C>G | 42 | 43 | 1.000 | | 1 |
| CAAATT | 1640C>G | 197 | 256 | | 1640C>G | 59 | 71 | 1.000 | | 1 |
| CTATCA | 1640C>G | 123 | 166 | | 1640C>G | 35 | 38 | 1.000 | | 1 |
| CTCAAA | 1640C>G | 201 | 283 | | 1640C>G | 52 | 56 | 1.000 | | 1 |
| GCGGAT | 1640C>G | 36 | 47 | | 1640C>G | 12 | 13 | 1.000 | | 1 |
| GGTATT | 1640C>G | 148 | 203 | | 1640C>G | 35 | 35 | 1.000 | | 1 |
| TCGGAT | 1640C>G | 72 | 100 | | 1640C>G | 14 | 18 | 1.000 | | 1 |
| AATTTT | 1656A>G | 168 | 243 | | 1656A>G | 44 | 47 | 1.000 | | 1 |
| CATCAA | 1656A>G | 100 | 141 | | 1656A>G | 38 | 41 | 1.000 | | 1 |
| CGGGCC | 1656A>G | 98 | 146 | | 1656A>G | 42 | 43 | 1.000 | | 1 |
| CTAGTC | 1656A>G | 26 | 34 | | 1656A>G | 4 | 31 | 1.000 | | 1 |
| GGTAAT | 1656A>G | 115 | 151 | | 1656A>G | 21 | 22 | 1.000 | | 1 |
| TAGCAG | 1656A>G | 143 | 191 | | 1656A>G | 29 | 31 | 1.000 | | 1 |
| ATAGAG | 1659T>G | 48 | 75 | | 1659T>G | 19 | 19 | 1.000 | | 1 |
| CCTGAC | 1659T>G | 27 | 29 | | 1659T>G | 7 | 8 | 0.993 | | 1 |
| CGGAAA | 1659T>G | 314 | 419 | | 1659T>G | 84 | 87 | 1.000 | | 1 |
| CGGTCG | 1659T>G | 131 | 170 | | 1659T>G | 32 | 32 | 1.000 | | 1 |
| CTAATA | 1659T>G | 50 | 60 | | 1659T>G | 20 | 21 | 1.000 | | 1 |
| GCCCAC | 1659T>G | 139 | 190 | | 1659T>G | 23 | 23 | 1.000 | | 1 |
| TGGTGT | 1659T>G | 274 | 403 | | 1659T>G | 76 | 82 | 1.000 | | 1 |
| GCCATA | 1661T>A | 156 | 226 | | 1661T>A | 27 | 27 | 1.000 | | 1 |
| GTCCAG | 1661T>A | 75 | 112 | | 1661T>A | 30 | 31 | 1.000 | | 1 |
| GTGCAT | 1661T>A | 79 | 123 | | 1661T>A | 30 | 30 | 1.000 | | 1 |
| AAGTCG | 1661T>G | 171 | 228 | | 1661T>G | 46 | 47 | 1.000 | | 1 |
| ATATCA | 1661T>G | 135 | 218 | | 1661T>G | 45 | 46 | 1.000 | | 1 |
| ATGCGA | 1661T>G | 263 | 496 | | 1661T>G | 51 | 69 | 1.000 | | 1 |
| CATTAG | 1661T>G | 237 | 316 | | 1661T>G | 64 | 66 | 1.000 | | 1 |
| CTGCGT | 1661T>G | 57 | 77 | | 1661T>G | 14 | 14 | 1.000 | | 1 |
| GCACTT | 1661T>G | 315 | 432 | | 1661T>G | 93 | 93 | 1.000 | | 1 |
| GTGAAG | 1661T>G | 30 | 38 | | 1661T>G | 12 | 13 | 1.000 | | 1 |
| ACACAC | 1665A>G | 26 | 32 | | 1665A>G | 16 | 16 | 1.000 | | 1 |
| ACTCAG | 1665A>G | 81 | 114 | | 1665A>G | 21 | 21 | 1.000 | | 1 |
| AGGTGT | 1665A>G | 16 | 28 | | 1665A>G | 9 | 9 | 1.000 | | 1 |

| Barcode | Illumina | | | | NanoPore | | | | | |
|---|---|---|---|---|---|---|---|---|---|---|
| | DNA variant (min 30 reads) | Variant reads | Total reads | | DNA variant | variant reads | total reads | mutation support | | Illumina vs NP Match |
| ATTACG | 1665A>G | 408 | 544 | | 1665A>G | 104 | 110 | 1.000 | | 1 |
| CGAATT | 1665A>G | 151 | 201 | | 1665A>G | 30 | 32 | 1.000 | | 1 |
| CTAACG | 1665A>G | 150 | 203 | | 1665A>G | 21 | 26 | 1.000 | | 1 |
| CTGAAT | 1665A>G | 164 | 349 | | 1665A>G | 60 | 93 | 1.000 | | 1 |
| GGGGGT | 1665A>G | 300 | 417 | | 1665A>G | 74 | 76 | 1.000 | | 1 |
| GTGAAA | 1665A>G | 10 | 17 | | 1665A>G | 3 | 4 | 1.000 | | 1 |
| TGAACG | 1665A>G | 97 | 129 | | 1665A>G | 34 | 38 | 1.000 | | 1 |
| TGCTTA | 1665A>G | 100 | 141 | | 1665A>G | 1 | 1 | 0.794 | | 1 |
| ATGGGT | 1666T>G | 79 | 110 | | 1666T>G | 27 | 30 | 1.000 | | 1 |
| CATACC | 1666T>G | 212 | 286 | | 1666T>G | 59 | 61 | 1.000 | | 1 |
| CCGCGG | 1666T>G | 186 | 266 | | 1666T>G | 38 | 38 | 1.000 | | 1 |
| CGAGGT | 1666T>G | 91 | 128 | | 1666T>G | 22 | 22 | 1.000 | | 1 |
| CTGACC | 1666T>G | 89 | 117 | | 1666T>G | 25 | 27 | 1.000 | | 1 |
| GACACG | 1666T>G | 12 | 32 | | 1666T>G | 6 | 9 | 0.797 | | 1 |
| GAGTCT | 1666T>G | 131 | 205 | | 1666T>G | 31 | 31 | 1.000 | | 1 |
| GGAAGA | 1666T>G | 295 | 567 | | 1666T>G | 62 | 88 | 1.000 | | 1 |
| GGATTG | 1666T>G | 113 | 149 | | 1666T>G | 27 | 30 | 1.000 | | 1 |
| GTTTTT | 1666T>G | 134 | 179 | | 1666T>G | 44 | 45 | 1.000 | | 1 |
| TACGCG | 1666T>G | 156 | 188 | | 1666T>G | 36 | 38 | 1.000 | | 1 |
| TAGGGG | 1666T>G | 273 | 385 | | 1666T>G | 74 | 79 | 1.000 | | 1 |
| TCGGCC | 1666T>G | 66 | 94 | | 1666T>G | 15 | 15 | 1.000 | | 1 |
| TCTACC | 1666T>G | 298 | 397 | | 1765A>G | 47 | 51 | 1.000 | | 0 |
| TGCCAT | 1666T>G | 136 | 186 | | 1666T>G | 34 | 35 | 1.000 | | 1 |
| TTTTAT | 1666T>G | 179 | 267 | | 1666T>G | 60 | 65 | 1.000 | | 1 |
| ACAGCG | 1670G>A | 200 | 294 | | 1670G>A | 68 | 72 | 1.000 | | 1 |
| CACAGA | 1670G>A | 116 | 155 | | 1670G>A | 29 | 32 | 1.000 | | 1 |
| CGACGT | 1670G>A | 131 | 218 | | 1670G>A | 35 | 38 | 1.000 | | 1 |
| CTAAGC | 1670G>A | 84 | 112 | | 1670G>A | 27 | 30 | 1.000 | | 1 |
| CTTAAG | 1670G>A | 90 | 120 | | 1670G>A | 31 | 31 | 1.000 | | 1 |
| CTTTAT | 1670G>A | 40 | 68 | | 1670G>A | 12 | 17 | 0.999 | | 1 |
| GGAAGC | 1670G>A | 21 | 29 | | 1666T>G | 1 | 2 | 0.063 | | 0 |
| TAATGC | 1670G>A | 164 | 238 | | 1670G>A | 45 | 47 | 1.000 | | 1 |
| TCCTTG | 1670G>A | 245 | 336 | | 1670G>A | 67 | 71 | 1.000 | | 1 |
| TGTGGG | 1670G>A | 60 | 90 | | 1670G>A | 22 | 23 | 1.000 | | 1 |
| TTCTGG | 1670G>A | 218 | 316 | | 1670G>A | 78 | 82 | 1.000 | | 1 |
| AGCAAA | 1670G>C | 20 | 36 | | 1670G>C | 10 | 10 | 1.000 | | 1 |
| AGCTGG | 1670G>C | 284 | 369 | | 1670G>C | 61 | 63 | 1.000 | | 1 |
| ATAACG | 1670G>C | 61 | 70 | | 1670G>C | 14 | 16 | 1.000 | | 1 |
| CAATCA | 1670G>C | 14 | 20 | | 1670G>C | 2 | 2 | 0.322 | | 1 |
| CGGCTG | 1670G>C | 223 | 290 | | 1765A>G | 1 | 1 | 0.422 | | 0 |
| GAACGA | 1670G>C | 85 | 102 | | 1670G>C | 22 | 23 | 1.000 | | 1 |
| TACTTT | 1670G>C | 26 | 40 | | 1670G>C | 4 | 5 | 1.000 | | 1 |
| TCTACA | 1670G>C | 83 | 120 | | 1670G>C | 21 | 23 | 1.000 | | 1 |
| TTGCTT | 1670G>C | 152 | 281 | | 1670G>C | 35 | 46 | 1.000 | | 1 |
| AACAGC | 1677T>C | 111 | 156 | | 1677T>C | 34 | 35 | 1.000 | | 1 |
| AAGAAT | 1677T>C | 76 | 121 | | 1677T>C | 20 | 20 | 1.000 | | 1 |
| CATTAT | 1677T>C | 141 | 186 | | 1677T>C | 36 | 38 | 1.000 | | 1 |
| GCTTAG | 1677T>C | 141 | 168 | | 1677T>C | 24 | 25 | 1.000 | | 1 |
| TGTTTT | 1677T>C | 248 | 335 | | 1677T>C | 70 | 71 | 1.000 | | 1 |
| AAAATC | 1677T>G | 33 | 49 | | 1677T>G | 14 | 16 | 1.000 | | 1 |
| AACACG | 1677T>G | 250 | 333 | | 1677T>G | 74 | 82 | 1.000 | | 1 |

| | Illumina | | | | NanoPore | | | | | |
|---|---|---|---|---|---|---|---|---|---|---|
| Barcode | DNA variant (min 30 reads) | Variant reads | Total reads | | DNA variant | variant reads | total reads | mutation support | | Illumina vs NP Match |
| AGAAGT | 1677T>G | 10 | 14 | | 1677T>G | 6 | 7 | 1.000 | | 1 |
| ATATCT | 1677T>G | 44 | 79 | | 1677T>G | 17 | 17 | 1.000 | | 1 |
| CAGTAT | 1677T>G | 106 | 139 | | 1677T>G | 29 | 31 | 1.000 | | 1 |
| CCAGTA | 1677T>G | 76 | 110 | | 1677T>G | 21 | 21 | 1.000 | | 1 |
| CTCTTT | 1677T>G | 62 | 119 | | 1677T>G | 25 | 26 | 1.000 | | 1 |
| CTGACA | 1677T>G | 62 | 93 | | 1677T>G | 8 | 9 | 1.000 | | 1 |
| GACCGT | 1677T>G | 90 | 130 | | 1677T>G | 20 | 24 | 1.000 | | 1 |
| GCTTCA | 1677T>G | 89 | 120 | | 1677T>G | 23 | 23 | 1.000 | | 1 |
| GGGTTA | 1677T>G | 157 | 234 | | 1677T>G | 47 | 50 | 1.000 | | 1 |
| TATAGC | 1677T>G | 328 | 475 | | 1677T>G | 99 | 101 | 1.000 | | 1 |
| TGTTGA | 1677T>G | 236 | 358 | | 1677T>G | 76 | 77 | 1.000 | | 1 |
| ATCACC | 1678A>G | 84 | 111 | | 1678A>G | 15 | 16 | 1.000 | | 1 |
| CGGGGT | 1678A>G | 283 | 422 | | 1678A>G | 47 | 53 | 1.000 | | 1 |
| GTATTA | 1678A>G | 36 | 48 | | 1678A>G | 2 | 3 | 1.000 | | 1 |
| ATTGAC | 1678A>T | 189 | 263 | | 1678A>T | 60 | 60 | 1.000 | | 1 |
| CACGAA | 1678A>T | 48 | 60 | | 1678A>T | 9 | 10 | 1.000 | | 1 |
| CCAGAA | 1678A>T | 20 | 28 | | 1689T>G | 1 | 3 | 0.126 | | 0 |
| CGGAAG | 1678A>T | 73 | 109 | | 1678A>T | 15 | 16 | 1.000 | | 1 |
| GAGTAA | 1678A>T | 239 | 351 | | 1678A>T | 100 | 108 | 1.000 | | 1 |
| GGCAAT | 1678A>T | 134 | 181 | | 1678A>T | 29 | 29 | 1.000 | | 1 |
| GGCGTG | 1678A>T | 147 | 193 | | 1678A>T | 45 | 46 | 1.000 | | 1 |
| GGGCAG | 1678A>T | 112 | 160 | | 1678A>T | 17 | 17 | 1.000 | | 1 |
| GTGGTA | 1678A>T | 97 | 128 | | 1678A>T | 16 | 22 | 1.000 | | 1 |
| TGACAC | 1678A>T | 160 | 222 | | 1678A>T | 29 | 30 | 1.000 | | 1 |
| TTCAAA | 1678A>T | 326 | 502 | | 1678A>T | 84 | 94 | 1.000 | | 1 |
| TTGGAT | 1678A>T | 120 | 148 | | 1678A>T | 28 | 31 | 1.000 | | 1 |
| TTTGAA | 1678A>T | 26 | 30 | | 1678A>T | 8 | 8 | 1.000 | | 1 |
| AATAGT | 1682A>T | 162 | 213 | | 1682A>T | 36 | 37 | 1.000 | | 1 |
| AATATT | 1682A>T | 18 | 21 | | 1682A>T | 3 | 5 | 1.000 | | 1 |
| AATCTA | 1682A>T | 287 | 383 | | 1682A>T | 45 | 46 | 1.000 | | 1 |
| ACGACG | 1682A>T | 162 | 253 | | 1682A>T | 27 | 31 | 1.000 | | 1 |
| ACGCTA | 1682A>T | 50 | 63 | | 1682A>T | 15 | 16 | 1.000 | | 1 |
| AGAAAT | 1682A>T | 119 | 233 | | 1682A>T | 33 | 48 | 1.000 | | 1 |
| AGGTGA | 1682A>T | 58 | 84 | | 1682A>T | 12 | 13 | 1.000 | | 1 |
| AGTAAG | 1682A>T | 267 | 381 | | 1682A>T | 63 | 65 | 1.000 | | 1 |
| AGTTAA | 1682A>T | 102 | 142 | | 1682A>T | 31 | 36 | 1.000 | | 1 |
| ATGAGG | 1682A>T | 6 | 18 | | 1788T>G | 2 | 3 | 0.654 | | 0 |
| ATTCGG | 1682A>T | 128 | 158 | | 1682A>T | 18 | 19 | 1.000 | | 1 |
| CAAGCT | 1682A>T | 72 | 90 | | 1682A>T | 19 | 21 | 1.000 | | 1 |
| CATCGG | 1682A>T | 148 | 203 | | 1682A>T | 35 | 38 | 1.000 | | 1 |
| CATTGC | 1682A>T | 76 | 104 | | 1682A>T | 22 | 25 | 1.000 | | 1 |
| CATTTT | 1682A>T | 352 | 718 | | 1682A>T | 70 | 107 | 1.000 | | 1 |
| CCAGCC | 1682A>T | 16 | 22 | | 1682A>T | 4 | 6 | 0.716 | | 1 |
| CCTTTC | 1682A>T | 148 | 214 | | 1682A>T | 40 | 44 | 1.000 | | 1 |
| CGACTT | 1682A>T | 51 | 73 | | 1682A>T | 14 | 14 | 1.000 | | 1 |
| CGGCGG | 1682A>T | 490 | 662 | | 1682A>T | 83 | 85 | 1.000 | | 1 |
| CTAGCC | 1682A>T | 54 | 110 | | 1682A>T | 7 | 13 | 1.000 | | 1 |
| CTCTTG | 1682A>T | 186 | 245 | | 1682A>T | 25 | 26 | 1.000 | | 1 |
| CTGTAG | 1682A>T | 315 | 404 | | 1682A>T | 49 | 51 | 1.000 | | 1 |
| CTGTGG | 1682A>T | 36 | 44 | | 1682A>T | 6 | 6 | 1.000 | | 1 |
| GTGTGA | 1682A>T | 116 | 159 | | 1682A>T | 28 | 28 | 1.000 | | 1 |

| Barcode | Illumina | | | | NanoPore | | | | | |
|---|---|---|---|---|---|---|---|---|---|---|
| | DNA variant (min 30 reads) | Variant reads | Total reads | | DNA variant | variant reads | total reads | mutation support | | Illumina vs NP Match |
| TAGGGA | 1682A>T | 255 | 345 | | 1682A>T | 64 | 69 | 1.000 | | 1 |
| TCTGAA | 1682A>T | 30 | 40 | | 1682A>T | 5 | 6 | 1.000 | | 1 |
| TGGAGA | 1682A>T | 101 | 144 | | 1682A>T | 13 | 14 | 1.000 | | 1 |
| TTATAA | 1682A>T | 285 | 401 | | 1682A>T | 69 | 75 | 1.000 | | 1 |
| TTCGTC | 1682A>T | 130 | 176 | | 1682A>T | 28 | 28 | 1.000 | | 1 |
| ACATGT | 1684T>G | 121 | 173 | | 1684T>G | 32 | 33 | 1.000 | | 1 |
| ACCTAC | 1684T>G | 26 | 34 | | 1684T>G | 16 | 17 | 1.000 | | 1 |
| AGTACT | 1684T>G | 160 | 235 | | 1684T>G | 50 | 57 | 1.000 | | 1 |
| CAACGG | 1684T>G | 175 | 254 | | 1684T>G | 44 | 47 | 1.000 | | 1 |
| CGAGTT | 1684T>G | 153 | 225 | | 1684T>G | 41 | 47 | 1.000 | | 1 |
| CGGCAG | 1684T>G | 140 | 187 | | 1682A>T | 1 | 1 | 0.159 | | 0 |
| CGGGTT | 1684T>G | 204 | 304 | | 1684T>G | 63 | 64 | 1.000 | | 1 |
| CTATAA | 1684T>G | 99 | 138 | | 1684T>G | 33 | 36 | 1.000 | | 1 |
| CTGGAG | 1684T>G | 95 | 136 | | 1684T>G | 32 | 33 | 1.000 | | 1 |
| CTTCAC | 1684T>G | 66 | 97 | | 1684T>G | 19 | 19 | 1.000 | | 1 |
| CTTGTC | 1684T>G | 179 | 252 | | 1684T>G | 54 | 56 | 1.000 | | 1 |
| GAAATC | 1684T>G | 253 | 477 | | 1684T>G | 98 | 126 | 1.000 | | 1 |
| GATAAG | 1684T>G | 127 | 157 | | 1684T>G | 46 | 47 | 1.000 | | 1 |
| GCGCTG | 1684T>G | 115 | 155 | | 1659T>G | 1 | 1 | 1.000 | | 0 |
| GCGGTA | 1684T>G | 62 | 83 | | 1684T>G | 16 | 16 | 1.000 | | 1 |
| GGAGAC | 1684T>G | 55 | 71 | | 1684T>G | 21 | 21 | 1.000 | | 1 |
| GGTGCC | 1684T>G | 175 | 246 | | 1684T>G | 59 | 62 | 1.000 | | 1 |
| GGTTTG | 1684T>G | 154 | 257 | | 1684T>G | 51 | 52 | 1.000 | | 1 |
| GTAAGC | 1684T>G | 24 | 44 | | 1684T>G | 7 | 7 | 1.000 | | 1 |
| GTCGGC | 1684T>G | 81 | 131 | | 1684T>G | 21 | 23 | 1.000 | | 1 |
| TACCAA | 1684T>G | 84 | 120 | | 1684T>G | 27 | 28 | 1.000 | | 1 |
| TACCGT | 1684T>G | 159 | 252 | | 1684T>G | 51 | 54 | 1.000 | | 1 |
| TAGCTA | 1684T>G | 175 | 240 | | 1684T>G | 39 | 48 | 1.000 | | 1 |
| TATCAA | 1684T>G | 39 | 51 | | 1684T>G | 7 | 8 | 1.000 | | 1 |
| TATTAG | 1684T>G | 69 | 119 | | 1684T>G | 27 | 28 | 1.000 | | 1 |
| TCATAA | 1684T>G | 255 | 345 | | 1684T>G | 55 | 58 | 1.000 | | 1 |
| TCCGGA | 1684T>G | 94 | 132 | | 1684T>G | 24 | 25 | 1.000 | | 1 |
| TTAGAA | 1684T>G | 45 | 66 | | 1684T>G | 19 | 19 | 1.000 | | 1 |
| ACCAAA | 1689T>G | 216 | 277 | | 1689T>G | 52 | 53 | 1.000 | | 1 |
| ACCGAC | 1689T>G | 159 | 224 | | 1689T>G | 46 | 47 | 1.000 | | 1 |
| AGATTG | 1689T>G | 191 | 265 | | 1689T>G | 34 | 35 | 1.000 | | 1 |
| ATACGC | 1689T>G | 447 | 604 | | 1689T>G | 106 | 106 | 1.000 | | 1 |
| ATCCGG | 1689T>G | 54 | 75 | | 1689T>G | 14 | 15 | 1.000 | | 1 |
| GCCAGA | 1689T>G | 26 | 36 | | 1689T>G | 5 | 5 | 1.000 | | 1 |
| GGGGGA | 1689T>G | 260 | 342 | | 1689T>G | 28 | 30 | 1.000 | | 1 |
| TACTCT | 1689T>G | 169 | 235 | | 1689T>G | 37 | 38 | 1.000 | | 1 |
| TCAAAT | 1689T>G | 154 | 226 | | 1689T>G | 36 | 40 | 1.000 | | 1 |
| TCCAGC | 1689T>G | 48 | 64 | | 1689T>G | 4 | 4 | 1.000 | | 1 |
| TTCGGG | 1689T>G | 119 | 175 | | 1689T>G | 26 | 27 | 1.000 | | 1 |
| TTCTTT | 1689T>G | 71 | 106 | | 1689T>G | 21 | 21 | 1.000 | | 1 |
| TTTGGG | 1689T>G | 179 | 256 | | 1689T>G | 32 | 36 | 1.000 | | 1 |
| TTTTTC | 1689T>G | 48 | 62 | | 1689T>G | 11 | 12 | 1.000 | | 1 |
| AAACCC | 1692G>A | 74 | 103 | | 1692G>A | 30 | 33 | 1.000 | | 1 |
| AAAGTC | 1692G>A | 184 | 233 | | 1692G>A | 65 | 69 | 1.000 | | 1 |
| AAGAGA | 1692G>A | 183 | 247 | | 1692G>A | 48 | 52 | 1.000 | | 1 |
| AATGTT | 1692G>A | 174 | 335 | | 1692G>A | 55 | 73 | 1.000 | | 1 |

| Barcode | Illumina | | | | NanoPore | | | | | Illumina vs NP Match |
|---|---|---|---|---|---|---|---|---|---|---|
| | DNA variant (min 30 reads) | Variant reads | Total reads | | DNA variant | variant reads | total reads | mutation support | | |
| ACCAGG | 1692G>A | 27 | 35 | | 1692G>A | 11 | 14 | 1.000 | | 1 |
| AGGATG | 1692G>A | 45 | 74 | | 1692G>A | 12 | 13 | 1.000 | | 1 |
| CAGGCT | 1692G>A | 143 | 193 | | 1692G>A | 29 | 31 | 1.000 | | 1 |
| CAGTTC | 1692G>A | 190 | 268 | | 1692G>A | 44 | 46 | 1.000 | | 1 |
| CGAACA | 1692G>A | 52 | 79 | | 1692G>A | 17 | 17 | 1.000 | | 1 |
| CGGGTC | 1692G>A | 99 | 134 | | 1692G>A | 24 | 28 | 1.000 | | 1 |
| CTCACT | 1692G>A | 66 | 93 | | 1692G>A | 14 | 17 | 1.000 | | 1 |
| CTGCAA | 1692G>A | 81 | 114 | | 1692G>A | 21 | 25 | 1.000 | | 1 |
| CTTGGG | 1692G>A | 217 | 310 | | 1692G>A | 59 | 63 | 1.000 | | 1 |
| GCGGCT | 1692G>A | 72 | 109 | | 1692G>A | 20 | 21 | 1.000 | | 1 |
| GGGATG | 1692G>A | 145 | 250 | | 1692G>A | 47 | 52 | 1.000 | | 1 |
| GGGGTG | 1692G>A | 94 | 131 | | 1692G>A | 36 | 37 | 1.000 | | 1 |
| GTCCCC | 1692G>A | 12 | 14 | | 1692G>A | 3 | 3 | 0.018 | | 1 |
| TATGGT | 1692G>A | 8 | 20 | | 1661T>G | 1 | 3 | 0.140 | | 0 |
| TATTTG | 1692G>A | 73 | 133 | | 1692G>A | 21 | 24 | 1.000 | | 1 |
| TCAAAA | 1692G>A | 82 | 134 | | 1692G>A | 18 | 20 | 1.000 | | 1 |
| TTACAC | 1692G>A | 20 | 24 | | 1692G>A | 4 | 4 | 1.000 | | 1 |
| TTGTGT | 1692G>A | 215 | 413 | | 1692G>A | 60 | 87 | 1.000 | | 1 |
| ACGAAT | 1714T>A | 97 | 159 | | 1714T>A | 20 | 21 | 1.000 | | 1 |
| ACGGGA | 1714T>A | 134 | 170 | | 1714T>A | 21 | 22 | 1.000 | | 1 |
| AGACGC | 1714T>A | 287 | 393 | | 1714T>A | 88 | 94 | 1.000 | | 1 |
| AGCACA | 1714T>A | 29 | 43 | | 1714T>A | 16 | 18 | 1.000 | | 1 |
| CAGAAG | 1714T>A | 8 | 14 | | 1714T>A | 5 | 5 | 1.000 | | 1 |
| CCGTTA | 1714T>A | 120 | 197 | | 1714T>A | 24 | 27 | 1.000 | | 1 |
| GACGGA | 1714T>A | 22 | 29 | | 1714T>A | 4 | 4 | 0.967 | | 1 |
| GCGTCT | 1714T>A | 151 | 240 | | 1714T>A | 36 | 38 | 1.000 | | 1 |
| GGGGCA | 1714T>A | 123 | 173 | | 1714T>A | 20 | 23 | 1.000 | | 1 |
| GTCAAT | 1714T>A | 45 | 73 | | 1714T>A | 19 | 20 | 1.000 | | 1 |
| TAACAG | 1714T>A | 141 | 204 | | 1714T>A | 40 | 41 | 1.000 | | 1 |
| TTGGTA | 1714T>A | 12 | 24 | | 1714T>A | 5 | 5 | 1.000 | | 1 |
| ATGATT | 1721T>A | 175 | 233 | | 1721T>A | 55 | 59 | 1.000 | | 1 |
| CGGTAT | 1721T>A | 28 | 47 | | 1721T>A | 6 | 8 | 1.000 | | 1 |
| CTAGGG | 1721T>A | 208 | 315 | | 1721T>A | 43 | 46 | 1.000 | | 1 |
| CTTATG | 1721T>A | 181 | 270 | | 1721T>A | 39 | 40 | 1.000 | | 1 |
| GTGTAC | 1721T>A | 128 | 171 | | 1721T>A | 30 | 31 | 1.000 | | 1 |
| TCCAGG | 1721T>A | 90 | 129 | | 1721T>A | 29 | 32 | 1.000 | | 1 |
| TGTGCC | 1721T>A | 171 | 228 | | 1721T>A | 37 | 38 | 1.000 | | 1 |
| TTTCGC | 1721T>A | 33 | 55 | | 1721T>A | 14 | 17 | 1.000 | | 1 |
| AAGTAT | 1725T>A | 113 | 164 | | 1725T>A | 32 | 33 | 1.000 | | 1 |
| ACGGGC | 1725T>A | 115 | 149 | | 1725T>A | 14 | 14 | 1.000 | | 1 |
| AGAAGG | 1725T>A | 73 | 88 | | 1725T>A | 11 | 14 | 1.000 | | 1 |
| AGAGAG | 1725T>A | 60 | 85 | | 1725T>A | 15 | 17 | 1.000 | | 1 |
| ATATGA | 1725T>A | 375 | 536 | | 1725T>A | 83 | 87 | 1.000 | | 1 |
| ATGATG | 1725T>A | 136 | 197 | | 1725T>A | 58 | 62 | 1.000 | | 1 |
| ATTCGA | 1725T>A | 16 | 34 | | 1725T>A | 3 | 4 | 1.000 | | 1 |
| ATTTAT | 1725T>A | 85 | 121 | | 1725T>A | 33 | 35 | 1.000 | | 1 |
| CAACAT | 1725T>A | 12 | 16 | | 1725T>A | 3 | 3 | 0.285 | | 1 |
| CAGTCC | 1725T>A | 64 | 80 | | 1725T>A | 12 | 14 | 1.000 | | 1 |
| CCCAAA | 1725T>A | 123 | 192 | | 1725T>A | 23 | 27 | 1.000 | | 1 |
| CCCTTA | 1725T>A | 179 | 229 | | 1725T>A | 37 | 39 | 1.000 | | 1 |
| CGCCGA | 1725T>A | 118 | 171 | | 1725T>A | 24 | 26 | 1.000 | | 1 |

| Barcode | Illumina | | | | NanoPore | | | | | |
|---|---|---|---|---|---|---|---|---|---|---|
| | DNA variant (min 30 reads) | Variant reads | Total reads | | DNA variant | variant reads | total reads | mutation support | | Illumina vs NP Match |
| CGGCTA | 1725T>A | 185 | 253 | | 1725T>A | 40 | 42 | 1.000 | | 1 |
| CGGGTG | 1725T>A | 52 | 74 | | 1725T>A | 11 | 12 | 0.992 | | 1 |
| CGGTTA | 1725T>A | 169 | 241 | | 1725T>A | 36 | 41 | 1.000 | | 1 |
| CGTGAA | 1725T>A | 121 | 168 | | 1725T>A | 20 | 20 | 1.000 | | 1 |
| CGTTTA | 1725T>A | 137 | 196 | | 1725T>A | 34 | 34 | 1.000 | | 1 |
| CTGACG | 1725T>A | 7 | 23 | | 1725T>A | 8 | 10 | 0.252 | | 1 |
| GAGTGC | 1725T>A | 122 | 167 | | 1725T>A | 37 | 38 | 1.000 | | 1 |
| GATTCT | 1725T>A | 63 | 88 | | 1725T>A | 19 | 20 | 1.000 | | 1 |
| GCGAGA | 1725T>A | 289 | 628 | | 1725T>A | 60 | 106 | 1.000 | | 1 |
| GCGCAT | 1725T>A | 170 | 243 | | 1725T>A | 40 | 41 | 1.000 | | 1 |
| GGATCG | 1725T>A | 48 | 145 | | 1725T>A | 13 | 34 | 0.996 | | 1 |
| GGTGCG | 1725T>A | 212 | 353 | | 1725T>A | 57 | 60 | 1.000 | | 1 |
| GTCAAC | 1725T>A | 233 | 333 | | 1725T>A | 69 | 74 | 1.000 | | 1 |
| TATACA | 1725T>A | 94 | 142 | | 1725T>A | 26 | 28 | 1.000 | | 1 |
| TATAGT | 1725T>A | 170 | 221 | | 1725T>A | 45 | 49 | 1.000 | | 1 |
| TCAACA | 1725T>A | 165 | 239 | | 1725T>A | 34 | 34 | 1.000 | | 1 |
| TCCTAA | 1725T>A | 253 | 356 | | 1725T>A | 44 | 48 | 1.000 | | 1 |
| TTGGTG | 1725T>A | 99 | 127 | | 1725T>A | 25 | 25 | 1.000 | | 1 |
| TTGTTC | 1725T>A | 66 | 114 | | 1725T>A | 24 | 25 | 1.000 | | 1 |
| AAGACG | 1732A>G | 195 | 279 | | 1732A>G | 43 | 47 | 1.000 | | 1 |
| ACTTAA | 1732A>G | 42 | 60 | | 1732A>G | 13 | 13 | 1.000 | | 1 |
| CAGGCC | 1732A>G | 101 | 123 | | 1732A>G | 22 | 28 | 1.000 | | 1 |
| CATGGT | 1732A>G | 62 | 141 | | 1732A>G | 15 | 28 | 1.000 | | 1 |
| TACTAT | 1732A>G | 117 | 172 | | 1732A>G | 27 | 31 | 1.000 | | 1 |
| TCCAGT | 1732A>G | 50 | 78 | | 1732A>G | 7 | 9 | 0.834 | | 1 |
| GGTTCT | 1733T>G | 22 | 34 | | 1733T>G | 4 | 5 | 0.867 | | 1 |
| AAGAAG | 1735C>G | 79 | 114 | | 1735C>G | 30 | 33 | 1.000 | | 1 |
| ACCGGG | 1735C>G | 49 | 79 | | 1735C>G | 16 | 19 | 1.000 | | 1 |
| ACGGTT | 1735C>G | 185 | 276 | | 1735C>G | 32 | 35 | 1.000 | | 1 |
| AGTGCA | 1735C>G | 33 | 52 | | 1735C>G | 8 | 10 | 1.000 | | 1 |
| ATTAGT | 1735C>G | 134 | 179 | | 1735C>G | 42 | 46 | 1.000 | | 1 |
| CGAAGA | 1735C>G | 114 | 170 | | 1735C>G | 31 | 31 | 1.000 | | 1 |
| CGAAGC | 1735C>G | 51 | 72 | | 1735C>G | 18 | 19 | 1.000 | | 1 |
| CTCGCC | 1735C>G | 295 | 421 | | 1735C>G | 69 | 72 | 1.000 | | 1 |
| CTGGAC | 1735C>G | 200 | 290 | | 1735C>G | 34 | 39 | 1.000 | | 1 |
| GAAGGC | 1735C>G | 78 | 116 | | 1735C>G | 20 | 22 | 1.000 | | 1 |
| GGATAG | 1735C>G | 121 | 183 | | 1735C>G | 32 | 33 | 1.000 | | 1 |
| GGTGGG | 1735C>G | 183 | 265 | | 1735C>G | 49 | 50 | 1.000 | | 1 |
| TAATTC | 1735C>G | 241 | 371 | | 1735C>G | 62 | 63 | 1.000 | | 1 |
| TCCACA | 1735C>G | 93 | 124 | | 1735C>G | 26 | 28 | 1.000 | | 1 |
| TCTCTC | 1735C>G | 16 | 28 | | 1735C>G | 6 | 7 | 1.000 | | 1 |
| AAAATA | 1741G>T | 38 | 54 | | 1741G>T | 10 | 11 | 1.000 | | 1 |
| ATATTA | 1741G>T | 125 | 181 | | 1741G>T | 22 | 27 | 1.000 | | 1 |
| ATCTAA | 1741G>T | 130 | 210 | | 1741G>T | 19 | 24 | 1.000 | | 1 |
| CATCGA | 1741G>T | 143 | 199 | | 1741G>T | 33 | 39 | 1.000 | | 1 |
| CCGAAA | 1741G>T | 194 | 281 | | 1741G>T | 21 | 26 | 1.000 | | 1 |
| CTGTAT | 1741G>T | 204 | 309 | | 1741G>T | 47 | 49 | 1.000 | | 1 |
| GGCGCT | 1741G>T | 130 | 210 | | 1741G>T | 36 | 40 | 1.000 | | 1 |
| TAAGCG | 1741G>T | 147 | 201 | | 1741G>T | 53 | 54 | 1.000 | | 1 |
| TTATGC | 1741G>T | 54 | 77 | | 1741G>T | 6 | 8 | 1.000 | | 1 |
| TTGGGG | 1741G>T | 47 | 61 | | 1741G>T | 10 | 11 | 1.000 | | 1 |

| Barcode | Illumina | | | | NanoPore | | | | | |
|---|---|---|---|---|---|---|---|---|---|---|
| | DNA variant (min 30 reads) | Variant reads | Total reads | | DNA variant | variant reads | total reads | mutation support | | Illumina vs NP Match |
| AAGTGA | 1742A>C | 178 | 241 | | 1742A>C | 32 | 34 | 1.000 | | 1 |
| AAGAAA | 1745A>G | 176 | 244 | | 1745A>G | 84 | 87 | 1.000 | | 1 |
| CGACAG | 1745A>G | 77 | 105 | | 1745A>G | 10 | 11 | 1.000 | | 1 |
| TGTCAG | 1745A>G | 51 | 72 | | 1745A>G | 11 | 11 | 1.000 | | 1 |
| AAGACA | 1747A>C | 179 | 256 | | 1747A>C | 50 | 63 | 1.000 | | 1 |
| AATCCG | 1747A>C | 255 | 345 | | 1747A>C | 64 | 83 | 1.000 | | 1 |
| ATGTGT | 1747A>C | 24 | 45 | | 1747A>C | 10 | 14 | 1.000 | | 1 |
| CAGGCG | 1747A>C | 129 | 168 | | 1747A>C | 27 | 34 | 1.000 | | 1 |
| GATGTC | 1747A>C | 186 | 278 | | 1747A>C | 30 | 43 | 1.000 | | 1 |
| GTGGGG | 1747A>C | 254 | 379 | | 1747A>C | 31 | 42 | 1.000 | | 1 |
| TCTAGA | 1747A>C | 94 | 112 | | 1747A>C | 15 | 19 | 1.000 | | 1 |
| AACTGA | 1765A>G | 251 | 369 | | 1765A>G | 56 | 58 | 1.000 | | 1 |
| CATGCC | 1765A>G | 321 | 440 | | 1666T>G | 62 | 63 | 1.000 | | 0 |
| CATTTA | 1765A>G | 244 | 344 | | 1765A>G | 50 | 51 | 1.000 | | 1 |
| CCAATC | 1765A>G | 137 | 173 | | 1765A>G | 20 | 22 | 1.000 | | 1 |
| CGCTTG | 1765A>G | 170 | 246 | | 1765A>G | 30 | 30 | 1.000 | | 1 |
| CTTGAC | 1765A>G | 27 | 44 | | 1765A>G | 8 | 8 | 0.984 | | 1 |
| GCGTGG | 1765A>G | 321 | 440 | | 1765A>G | 56 | 57 | 1.000 | | 1 |
| GTTAAT | 1765A>G | 92 | 161 | | 1765A>G | 33 | 42 | 1.000 | | 1 |
| GTTTTC | 1765A>G | 220 | 293 | | 1765A>G | 63 | 67 | 1.000 | | 1 |
| TACATA | 1765A>G | 38 | 50 | | 1765A>G | 6 | 6 | 0.624 | | 1 |
| TACGTA | 1765A>G | 95 | 122 | | 1765A>G | 22 | 22 | 1.000 | | 1 |
| TATGGG | 1765A>G | 312 | 520 | | 1765A>G | 68 | 87 | 1.000 | | 1 |
| TCAAAG | 1765A>G | 112 | 162 | | 1765A>G | 36 | 40 | 1.000 | | 1 |
| TCCTTA | 1765A>G | 100 | 175 | | 1765A>G | 41 | 43 | 1.000 | | 1 |
| TCGATA | 1765A>G | 341 | 669 | | 1765A>G | 83 | 114 | 1.000 | | 1 |
| TGTAAC | 1765A>G | 22 | 24 | | 1765A>G | 6 | 6 | 0.923 | | 1 |
| TGTCGG | 1765A>G | 22 | 30 | | 1765A>G | 5 | 6 | 0.930 | | 1 |
| ACCCAG | 1785T>G | 236 | 369 | | 1785T>G | 54 | 56 | 1.000 | | 1 |
| ATATAT | 1785T>G | 169 | 238 | | 1785T>G | 51 | 53 | 1.000 | | 1 |
| GCGAAC | 1785T>G | 125 | 192 | | 1785T>G | 29 | 30 | 1.000 | | 1 |
| TAAATG | 1785T>G | 14 | 24 | | 1785T>G | 5 | 7 | 1.000 | | 1 |
| TGATCC | 1785T>G | 100 | 132 | | 1785T>G | 26 | 27 | 1.000 | | 1 |
| TTTGGT | 1785T>G | 138 | 265 | | 1785T>G | 19 | 37 | 1.000 | | 1 |
| ATTTCC | 1788T>G | 37 | 50 | | 1788T>G | 14 | 15 | 1.000 | | 1 |
| CGAGGG | 1788T>G | 153 | 247 | | 1788T>G | 48 | 51 | 1.000 | | 1 |
| TGAAGG | 1788T>G | 211 | 310 | | 1788T>G | 75 | 80 | 1.000 | | 1 |
| AAGGGT | 1799A>G | 74 | 112 | | 1799A>G | 32 | 35 | 1.000 | | 1 |
| ATATGC | 1799A>G | 137 | 236 | | 1799A>G | 35 | 40 | 1.000 | | 1 |
| CTTGAT | 1799A>G | 103 | 137 | | 1799A>G | 26 | 27 | 1.000 | | 1 |
| ACAACT | 1802T>G | 76 | 123 | | 1802T>G | 19 | 19 | 1.000 | | 1 |
| ACGAAA | 1802T>G | 90 | 155 | | 1802T>G | 38 | 39 | 1.000 | | 1 |
| CTGGAT | 1802T>G | 76 | 143 | | 1802T>G | 21 | 22 | 1.000 | | 1 |
| TAATGA | 1802T>G | 28 | 48 | | 1802T>G | 15 | 17 | 1.000 | | 1 |
| TCCCCT | 1802T>G | 125 | 212 | | 1802T>G | 24 | 27 | 1.000 | | 1 |
| ACAGTA | 1807A>G | 70 | 106 | | 1807A>G | 12 | 12 | 0.997 | | 1 |
| ATCCAT | 1807A>G | 294 | 414 | | 1807A>G | 74 | 75 | 1.000 | | 1 |
| CTGATA | 1807A>G | 28 | 57 | | 1807A>G | 22 | 26 | 1.000 | | 1 |
| GAATTC | 1807A>G | 215 | 307 | | 1807A>G | 64 | 67 | 1.000 | | 1 |
| GTATAG | 1807A>G | 191 | 303 | | 1807A>G | 60 | 61 | 1.000 | | 1 |
| TACCTG | 1807A>G | 99 | 174 | | 1689T>G | 1 | 1 | 0.631 | | 0 |

| Barcode | Illumina | | | | NanoPore | | | | | |
|---|---|---|---|---|---|---|---|---|---|---|
| | DNA variant (min 30 reads) | Variant reads | Total reads | | DNA variant | variant reads | total reads | mutation support | | Illumina vs NP Match |
| TGTGCA | 1807A>G | 170 | 243 | | 1807A>G | 22 | 23 | 1.000 | | 1 |
| TTAGAC | 1807A>G | 54 | 78 | | 1807A>G | 8 | 9 | 0.969 | | 1 |
| AAGGAT | 1817C>A | 37 | 51 | | 1817C>A | 22 | 23 | 1.000 | | 1 |
| CAATGA | 1817C>A | 57 | 77 | | 1817C>A | 15 | 16 | 1.000 | | 1 |
| CACTTT | 1817C>A | 208 | 335 | | 1817C>A | 59 | 61 | 1.000 | | 1 |
| CAGACC | 1817C>A | 145 | 210 | | 1817C>A | 30 | 34 | 1.000 | | 1 |
| CCATTC | 1817C>A | 4 | 13 | | 1656A>G | 1 | 4 | 0.643 | | 0 |
| CCTTGA | 1817C>A | 119 | 195 | | 1817C>A | 32 | 32 | 1.000 | | 1 |
| CTGTTT | 1817C>A | 125 | 176 | | 1817C>A | 28 | 30 | 1.000 | | 1 |
| GAAAGG | 1817C>A | 101 | 153 | | 1817C>A | 32 | 35 | 1.000 | | 1 |
| GAAGGT | 1817C>A | 100 | 169 | | 1817C>A | 43 | 44 | 1.000 | | 1 |
| GACCAA | 1817C>A | 28 | 62 | | 1817C>A | 13 | 15 | 1.000 | | 1 |
| GGGAAT | 1817C>A | 139 | 190 | | 1817C>A | 37 | 40 | 1.000 | | 1 |
| TACGGA | 1817C>A | 42 | 56 | | 1817C>A | 18 | 18 | 1.000 | | 1 |
| TCAGAA | 1817C>A | 13 | 30 | | 1817C>A | 4 | 4 | 1.000 | | 1 |
| TGGGTA | 1817C>A | 22 | 32 | | 1817C>A | 11 | 11 | 1.000 | | 1 |
| TTAATT | 1817C>A | 146 | 221 | | 1817C>A | 48 | 50 | 1.000 | | 1 |
| TTATCA | 1817C>A | 66 | 90 | | 1817C>A | 14 | 14 | 0.996 | | 1 |
| TTCCGA | 1817C>A | 287 | 442 | | 1817C>A | 62 | 62 | 1.000 | | 1 |
| TTCCGC | 1817C>A | 49 | 84 | | 1817C>A | 18 | 18 | 1.000 | | 1 |
| TTCGAT | 1817C>A | 141 | 186 | | 1817C>A | 16 | 16 | 0.984 | | 1 |
| TTTTGG | 1817C>A | 230 | 377 | | 1817C>A | 70 | 75 | 1.000 | | 1 |
| TCCTCT | 1826C>G | 47 | 81 | | 1826C>G | 29 | 33 | 1.000 | | 1 |
| CAGTCA | 1830T>A | 141 | 217 | | 1830T>A | 70 | 77 | 1.000 | | 1 |
| CCTGGG | 1830T>A | 43 | 73 | | 1830T>A | 20 | 23 | 1.000 | | 1 |
| TCATCA | 1830T>A | 38 | 54 | | 1830T>A | 15 | 16 | 1.000 | | 1 |
| TGGTAG | 1830T>A | 33 | 44 | | 1830T>A | 15 | 20 | 1.000 | | 1 |
| ACCGGT | 1834A>G | 72 | 107 | | 1834A>G | 38 | 41 | 1.000 | | 1 |
| CGTACT | 1834A>G | 47 | 96 | | 1834A>G | 26 | 29 | 1.000 | | 1 |
| CAACCA | 1839T>G | 69 | 105 | | 1839T>G | 26 | 27 | 1.000 | | 1 |
| CGGATA | 1839T>G | 42 | 74 | | 1839T>G | 19 | 21 | 1.000 | | 1 |
| GACTCG | 1839T>G | 79 | 130 | | 1839T>G | 56 | 56 | 1.000 | | 1 |

**Table S4a** - background splicing noise

| Pseudoexon [span] | # of reads with the PE in wts | Total reads in wts (all libraries) | Activation (%) | Detected in mutant settings? | Notes |
|---|---|---|---|---|---|
| [3769-3845] | 1797 | 234705 | 0.766% | YES | = PEwt |
| [1018-NA] | 78 | 234705 | 0.033% | YES | library 5 excluded |
| [1917-2025] | 3 | 234705 | 1.28E-05 | YES | |
| [1917-1995] | 2 | 234705 | 8.52E-06 | YES | |
| [3638-3746] | 2 | 234705 | 8.52E-06 | NO | |
| [3711-3845] | 2 | 234705 | 8.52E-06 | YES | |
| [547-662] | 1 | 234705 | 4.26E-06 | NO | |
| [596-662] | 1 | 234705 | 4.26E-06 | YES | |
| [957-980] | 1 | 234705 | 4.26E-06 | YES | |
| [1249-1387] | 1 | 234705 | 4.26E-06 | NO | |
| [1533-NA] | 1 | 234705 | 4.26E-06 | YES | |
| [1534-1685] | 1 | 234705 | 4.26E-06 | YES | |
| [1534-NA] | 1 | 234705 | 4.31E-06 | YES | library 8 excluded |
| [1959-1995] | 1 | 234705 | 4.26E-06 | YES | |
| [1959-2025] | 1 | 234705 | 4.26E-06 | YES | |
| [3638-3845] | 1 | 234705 | 4.26E-06 | NO | |
| [3715-3845] | 1 | 234705 | 4.26E-06 | YES | |
| [3796-3845] | 1 | 234705 | 4.26E-06 | YES | |
| [4051-4064] | 1 | 234705 | 4.26E-06 | NO | |
| [NA-1387] | 1 | 234705 | 4.26E-06 | YES | |

**Table S4b** - activation of pseudoexon [1018-NA] among wts, comparison of library 5 with all other libraries.

| Pseudoexon | # of RNA reads with the PE (wts of library 5 only) | Total RNA reads in wts (library 5 only) | Activation (library 5 only) | # of RNA reads with the PE (all other libraries) | Total RNA reads in wts (all other libraries) | Activation (all other libraries) | Activation difference | p (adj, fdr) | log odds ratio | se log odds ratio | odds ratio |
|---|---|---|---|---|---|---|---|---|---|---|---|
| [1018-NA] | 2 | 293 | 0.683% | 78 | 234412 | 0.033% | 0.649% | 2.88E-11 | 2.298 | 0.316 | 9.958 |

**Table S4c -** Comparison of pseudoexon activation in variants of the library 5 with the activation in wild types of library 5 only

| Variant | Pseudoexon | # of RNA reads with the PE | Total RNA reads of the variant | Activation | # of RNA reads with the PE (wts of library 5) | Total RNA reads in wt (wts of library 5) | Activation (wts of library 5) | Activation difference | p (adj, fdr) | log odds ratio | se log odds ratio | odds ratio |
|---|---|---|---|---|---|---|---|---|---|---|---|---|
| 1137C>T | [1018-NA] | 151 | 288 | 52.431% | 2 | 293 | 0.683% | 51.748% | 4.66E-08 | 4.495 | 0.418 | 89.565 |
| 1156C>G | [1018-NA] | 120 | 1348 | 8.902% | 2 | 293 | 0.683% | 8.219% | 8.20E-04 | 2.125 | 0.565 | 8.376 |
| 1207C>T | [1018-NA] | 15 | 8304 | 0.181% | 2 | 293 | 0.683% | -0.502% | 2.29E-02 | -1.655 | 0.638 | 0.191 |
| 1297A>G | [1018-NA] | 9 | 5134 | 0.175% | 2 | 293 | 0.683% | -0.507% | 7.88E-04 | -1.711 | 0.429 | 0.181 |
| 4026T>G | [1018-NA] | 2 | 1154 | 0.173% | 2 | 293 | 0.683% | -0.509% | 1.35E-02 | -1.554 | 0.523 | 0.211 |
| 2802A>G | [1018-NA] | 35 | 20865 | 0.168% | 2 | 293 | 0.683% | -0.515% | 1.80E-03 | -1.692 | 0.490 | 0.184 |
| 1917T>G | [1018-NA] | 3 | 1860 | 0.161% | 2 | 293 | 0.683% | -0.521% | 2.22E-02 | -0.850 | 0.329 | 0.427 |
| 1362A>G | [1018-NA] | 16 | 10153 | 0.158% | 2 | 293 | 0.683% | -0.525% | 5.28E-04 | -1.851 | 0.459 | 0.157 |
| 1600T>A | [1018-NA] | 5 | 3217 | 0.155% | 2 | 293 | 0.683% | -0.527% | 2.16E-03 | -1.717 | 0.473 | 0.180 |
| 644A>G | [1018-NA] | 4 | 2589 | 0.154% | 2 | 293 | 0.683% | -0.528% | 4.95E-02 | -1.774 | 0.765 | 0.170 |
| 1605G>A | [1018-NA] | 6 | 4017 | 0.149% | 2 | 293 | 0.683% | -0.533% | 3.98E-02 | -1.771 | 0.743 | 0.170 |
| 3604A>T | [1018-NA] | 3 | 2021 | 0.148% | 2 | 293 | 0.683% | -0.534% | 4.74E-02 | -1.473 | 0.641 | 0.229 |
| 1262T>A | [1018-NA] | 12 | 8512 | 0.141% | 2 | 293 | 0.683% | -0.542% | 5.29E-04 | -1.922 | 0.474 | 0.146 |
| 1530T>G | [1018-NA] | 4 | 2860 | 0.140% | 2 | 293 | 0.683% | -0.543% | 5.46E-03 | -1.688 | 0.521 | 0.185 |
| 2735G>C | [1018-NA] | 10 | 7232 | 0.138% | 2 | 293 | 0.683% | -0.544% | 2.35E-05 | -1.773 | 0.360 | 0.170 |
| 1235T>G | [1018-NA] | 10 | 7503 | 0.133% | 2 | 293 | 0.683% | -0.549% | 8.00E-04 | -2.006 | 0.504 | 0.134 |
| 2470A>G | [1018-NA] | 1 | 767 | 0.130% | 2 | 293 | 0.683% | -0.552% | 1.90E-02 | -1.603 | 0.569 | 0.201 |
| 949A>T | [1018-NA] | 1 | 785 | 0.127% | 2 | 293 | 0.683% | -0.555% | 1.56E-02 | -1.352 | 0.474 | 0.259 |
| 1349G>A | [1018-NA] | 6 | 4749 | 0.126% | 2 | 293 | 0.683% | -0.556% | 1.55E-02 | -1.927 | 0.688 | 0.146 |
| 2884A>G | [1018-NA] | 1 | 798 | 0.125% | 2 | 293 | 0.683% | -0.557% | 2.18E-02 | -1.814 | 0.650 | 0.163 |
| 2643G>C | [1018-NA] | 17 | 13701 | 0.124% | 2 | 293 | 0.683% | -0.559% | 1.20E-03 | -2.072 | 0.558 | 0.126 |
| 2812A>C | [1018-NA] | 5 | 4050 | 0.123% | 2 | 293 | 0.683% | -0.559% | 3.56E-03 | -1.906 | 0.559 | 0.149 |
| 2814A>C | [1018-NA] | 2 | 1650 | 0.121% | 2 | 293 | 0.683% | -0.561% | 1.64E-02 | -1.945 | 0.674 | 0.143 |
| 1558T>A | [1018-NA] | 9 | 7629 | 0.118% | 2 | 293 | 0.683% | -0.565% | 2.29E-04 | -1.984 | 0.462 | 0.138 |
| 3176G>C | [1018-NA] | 9 | 7718 | 0.117% | 2 | 293 | 0.683% | -0.566% | 6.32E-04 | -1.825 | 0.472 | 0.161 |
| 534G>A | [1018-NA] | 3 | 2608 | 0.115% | 2 | 293 | 0.683% | -0.568% | 3.36E-04 | -1.964 | 0.441 | 0.140 |
| 3630T>G | [1018-NA] | 2 | 1758 | 0.114% | 2 | 293 | 0.683% | -0.569% | 5.76E-03 | -1.688 | 0.519 | 0.185 |
| 3989G>T | [1018-NA] | 4 | 3533 | 0.113% | 2 | 293 | 0.683% | -0.569% | 6.33E-03 | -1.947 | 0.609 | 0.143 |
| 1337T>G | [1018-NA] | 6 | 5302 | 0.113% | 2 | 293 | 0.683% | -0.569% | 2.47E-02 | -2.123 | 0.807 | 0.120 |
| 1862C>G | [1018-NA] | 4 | 3556 | 0.112% | 2 | 293 | 0.683% | -0.570% | 1.81E-02 | -0.966 | 0.366 | 0.381 |
| 2562T>A | [1018-NA] | 3 | 2700 | 0.111% | 2 | 293 | 0.683% | -0.571% | 4.34E-03 | -2.141 | 0.611 | 0.117 |
| 2608C>G | [1018-NA] | 5 | 4514 | 0.111% | 2 | 293 | 0.683% | -0.572% | 6.92E-03 | -2.144 | 0.667 | 0.117 |
| 1865T>G | [1018-NA] | 7 | 6526 | 0.107% | 2 | 293 | 0.683% | -0.575% | 1.89E-04 | -1.049 | 0.256 | 0.350 |
| 1311A>G | [1018-NA] | 8 | 7505 | 0.107% | 2 | 293 | 0.683% | -0.576% | 3.38E-05 | -2.100 | 0.421 | 0.122 |
| 2643G>T | [1018-NA] | 7 | 6594 | 0.106% | 2 | 293 | 0.683% | -0.576% | 1.22E-03 | -2.155 | 0.564 | 0.116 |
| 1392C>G | [1018-NA] | 2 | 1900 | 0.105% | 2 | 293 | 0.683% | -0.577% | 4.22E-03 | -2.121 | 0.600 | 0.120 |
| 3144G>T | [1018-NA] | 8 | 7604 | 0.105% | 2 | 293 | 0.683% | -0.577% | 1.50E-04 | -1.921 | 0.443 | 0.146 |
| 242C>G | [1018-NA] | 3 | 2889 | 0.104% | 2 | 293 | 0.683% | -0.579% | 1.93E-03 | -1.884 | 0.517 | 0.152 |
| 3311G>T | [1018-NA] | 1 | 965 | 0.104% | 2 | 293 | 0.683% | -0.579% | 1.34E-02 | -2.004 | 0.662 | 0.135 |
| 3050T>G | [1018-NA] | 2 | 1953 | 0.102% | 2 | 293 | 0.683% | -0.580% | 6.99E-04 | -1.698 | 0.421 | 0.183 |

| | | | | | | | | | | | | |
|---|---|---|---|---|---|---|---|---|---|---|---|---|
| 2736A>T | [1018-NA] | 10 | 9782 | 0.102% | 2 | 293 | 0.683% | -0.580% | 4.96E-05 | -2.036 | 0.436 | 0.131 |
| 2050A>G | [1018-NA] | 1 | 984 | 0.102% | 2 | 293 | 0.683% | -0.581% | 2.63E-02 | -1.135 | 0.443 | 0.322 |
| 1907T>G | [1018-NA] | 2 | 1982 | 0.101% | 2 | 293 | 0.683% | -0.582% | 1.59E-02 | -0.935 | 0.345 | 0.393 |
| 3640A>C | [1018-NA] | 6 | 5991 | 0.100% | 2 | 293 | 0.683% | -0.582% | 5.71E-05 | -1.767 | 0.385 | 0.171 |
| 2466T>A | [1018-NA] | 6 | 6000 | 0.100% | 2 | 293 | 0.683% | -0.583% | 2.36E-03 | -2.039 | 0.583 | 0.130 |
| 3138C>G | [1018-NA] | 7 | 7036 | 0.099% | 2 | 293 | 0.683% | -0.583% | 8.39E-04 | -1.993 | 0.522 | 0.136 |
| 493T>A | [1018-NA] | 1 | 1024 | 0.098% | 2 | 293 | 0.683% | -0.585% | 1.14E-02 | -2.063 | 0.664 | 0.127 |
| 1892T>A | [1018-NA] | 2 | 2053 | 0.097% | 2 | 293 | 0.683% | -0.585% | 1.55E-03 | -1.080 | 0.303 | 0.340 |
| 302C>A | [1018-NA] | 7 | 7231 | 0.097% | 2 | 293 | 0.683% | -0.586% | 1.09E-04 | -1.923 | 0.435 | 0.146 |
| 1857A>G | [1018-NA] | 1 | 1039 | 0.096% | 2 | 293 | 0.683% | -0.586% | 3.25E-02 | -1.115 | 0.453 | 0.328 |
| 303G>A | [1018-NA] | 3 | 3129 | 0.096% | 2 | 293 | 0.683% | -0.587% | 8.83E-03 | -2.021 | 0.660 | 0.133 |
| 3166T>G | [1018-NA] | 3 | 3134 | 0.096% | 2 | 293 | 0.683% | -0.587% | 2.12E-02 | -2.148 | 0.791 | 0.117 |
| 404T>G | [1018-NA] | 1 | 1049 | 0.095% | 2 | 293 | 0.683% | -0.587% | 4.23E-03 | -1.769 | 0.511 | 0.170 |
| 2735G>T | [1018-NA] | 2 | 2107 | 0.095% | 2 | 293 | 0.683% | -0.588% | 1.54E-03 | -2.002 | 0.522 | 0.135 |
| 3283C>G | [1018-NA] | 2 | 2113 | 0.095% | 2 | 293 | 0.683% | -0.588% | 7.08E-03 | -1.923 | 0.605 | 0.146 |
| 1898A>G | [1018-NA] | 2 | 2125 | 0.094% | 2 | 293 | 0.683% | -0.588% | 8.75E-03 | -0.934 | 0.318 | 0.393 |
| 1444T>G | [1018-NA] | 2 | 2132 | 0.094% | 2 | 293 | 0.683% | -0.589% | 1.83E-03 | -2.202 | 0.567 | 0.111 |
| 2567T>A | [1018-NA] | 5 | 5342 | 0.094% | 2 | 293 | 0.683% | -0.589% | 2.26E-05 | -2.268 | 0.418 | 0.104 |
| 2461T>G | [1018-NA] | 7 | 7518 | 0.093% | 2 | 293 | 0.683% | -0.589% | 5.73E-05 | -1.970 | 0.426 | 0.139 |
| 2053T>G | [1018-NA] | 1 | 1074 | 0.093% | 2 | 293 | 0.683% | -0.589% | 7.33E-03 | -1.036 | 0.338 | 0.355 |
| 3883T>G | [1018-NA] | 4 | 4301 | 0.093% | 2 | 293 | 0.683% | -0.590% | 1.05E-04 | -2.066 | 0.442 | 0.127 |
| 2642A>T | [1018-NA] | 3 | 3238 | 0.093% | 2 | 293 | 0.683% | -0.590% | 6.26E-04 | -2.323 | 0.532 | 0.098 |
| 3331C>G | [1018-NA] | 3 | 3303 | 0.091% | 2 | 293 | 0.683% | -0.592% | 9.92E-04 | -2.075 | 0.528 | 0.126 |
| 1566A>G | [1018-NA] | 12 | 13218 | 0.091% | 2 | 293 | 0.683% | -0.592% | 1.17E-03 | -2.005 | 0.555 | 0.135 |
| 1322T>A | [1018-NA] | 3 | 3335 | 0.090% | 2 | 293 | 0.683% | -0.593% | 7.65E-04 | -2.520 | 0.553 | 0.080 |
| 1898A>T | [1018-NA] | 1 | 1113 | 0.090% | 2 | 293 | 0.683% | -0.593% | 3.49E-03 | -1.030 | 0.308 | 0.357 |
| 3981T>A | [1018-NA] | 3 | 3403 | 0.088% | 2 | 293 | 0.683% | -0.594% | 8.03E-03 | -1.977 | 0.645 | 0.139 |
| 2065A>G | [1018-NA] | 1 | 1138 | 0.088% | 2 | 293 | 0.683% | -0.595% | 3.39E-03 | -1.137 | 0.337 | 0.321 |
| 228T>G | [1018-NA] | 3 | 3473 | 0.086% | 2 | 293 | 0.683% | -0.596% | 1.68E-04 | -1.914 | 0.431 | 0.147 |
| 1520T>C | [1018-NA] | 1 | 1162 | 0.086% | 2 | 293 | 0.683% | -0.597% | 7.18E-03 | -1.872 | 0.580 | 0.154 |
| 2388T>C | [1018-NA] | 2 | 2325 | 0.086% | 2 | 293 | 0.683% | -0.597% | 5.47E-04 | -2.019 | 0.480 | 0.133 |
| 493T>G | [1018-NA] | 1 | 1202 | 0.083% | 2 | 293 | 0.683% | -0.599% | 7.17E-03 | -1.906 | 0.590 | 0.149 |
| 1905T>C | [1018-NA] | 2 | 2466 | 0.081% | 2 | 293 | 0.683% | -0.601% | 2.89E-04 | -1.017 | 0.251 | 0.362 |
| 1856A>G | [1018-NA] | 2 | 2518 | 0.079% | 2 | 293 | 0.683% | -0.603% | 3.01E-03 | -1.113 | 0.336 | 0.328 |
| 1962A>G | [1018-NA] | 1 | 1270 | 0.079% | 2 | 293 | 0.683% | -0.604% | 7.11E-04 | -1.103 | 0.283 | 0.332 |
| 1329T>A | [1018-NA] | 6 | 7629 | 0.079% | 2 | 293 | 0.683% | -0.604% | 1.01E-03 | -2.280 | 0.599 | 0.102 |
| 1499G>T | [1018-NA] | 2 | 2544 | 0.079% | 2 | 293 | 0.683% | -0.604% | 2.20E-04 | -2.191 | 0.474 | 0.112 |
| 423A>T | [1018-NA] | 4 | 5149 | 0.078% | 2 | 293 | 0.683% | -0.605% | 3.78E-03 | -2.045 | 0.618 | 0.129 |
| 1807A>G | [1018-NA] | 1 | 1292 | 0.077% | 2 | 293 | 0.683% | -0.605% | 2.89E-03 | -1.543 | 0.442 | 0.214 |
| 3223A>G | [1018-NA] | 8 | 10383 | 0.077% | 2 | 293 | 0.683% | -0.606% | 9.54E-04 | -2.090 | 0.564 | 0.124 |
| 2554T>G | [1018-NA] | 12 | 15587 | 0.077% | 2 | 293 | 0.683% | -0.606% | 7.89E-05 | -2.382 | 0.526 | 0.092 |
| 3978G>A | [1018-NA] | 3 | 3934 | 0.076% | 2 | 293 | 0.683% | -0.606% | 4.63E-04 | -2.231 | 0.530 | 0.107 |
| 2727G>C | [1018-NA] | 4 | 5272 | 0.076% | 2 | 293 | 0.683% | -0.607% | 8.07E-06 | -2.226 | 0.405 | 0.108 |
| 196A>G | [1018-NA] | 2 | 2674 | 0.075% | 2 | 293 | 0.683% | -0.608% | 1.68E-02 | -2.083 | 0.747 | 0.125 |
| 3173C>G | [1018-NA] | 8 | 10737 | 0.075% | 2 | 293 | 0.683% | -0.608% | 3.01E-05 | -2.223 | 0.464 | 0.108 |
| 1747A>C | [1018-NA] | 1 | 1350 | 0.074% | 2 | 293 | 0.683% | -0.609% | 3.33E-02 | -1.679 | 0.677 | 0.186 |

| | | | | | | | | | | | | |
|---|---|---|---|---|---|---|---|---|---|---|---|---|
| 1920T>G | [1018-NA] | 2 | 2701 | 0.074% | 2 | 293 | 0.683% | -0.609% | 2.26E-02 | -1.223 | 0.476 | 0.294 |
| 1300A>T | [1018-NA] | 4 | 5407 | 0.074% | 2 | 293 | 0.683% | -0.609% | 3.82E-04 | -2.389 | 0.556 | 0.092 |
| 1524C>T | [1018-NA] | 3 | 4063 | 0.074% | 2 | 293 | 0.683% | -0.609% | 1.02E-02 | -2.025 | 0.688 | 0.132 |
| 3348C>T | [1018-NA] | 2 | 2725 | 0.073% | 2 | 293 | 0.683% | -0.609% | 9.52E-06 | -1.884 | 0.347 | 0.152 |
| 1741G>T | [1018-NA] | 1 | 1367 | 0.073% | 2 | 293 | 0.683% | -0.609% | 1.18E-03 | -1.438 | 0.380 | 0.237 |
| 3818A>G | [1018-NA] | 4 | 5497 | 0.073% | 2 | 293 | 0.683% | -0.610% | 2.04E-02 | -2.268 | 0.849 | 0.104 |
| 2590A>G | [1018-NA] | 32 | 43980 | 0.073% | 2 | 293 | 0.683% | -0.610% | 2.02E-04 | -2.441 | 0.595 | 0.087 |
| 128A>G | [1018-NA] | 12 | 16572 | 0.072% | 2 | 293 | 0.683% | -0.610% | 2.02E-06 | -2.200 | 0.410 | 0.111 |
| 2388T>G | [1018-NA] | 6 | 8319 | 0.072% | 2 | 293 | 0.683% | -0.610% | 1.99E-04 | -2.169 | 0.510 | 0.114 |
| 2566G>A | [1018-NA] | 2 | 2785 | 0.072% | 2 | 293 | 0.683% | -0.611% | 2.40E-03 | -2.701 | 0.682 | 0.067 |
| 1429A>G | [1018-NA] | 3 | 4206 | 0.071% | 2 | 293 | 0.683% | -0.611% | 5.12E-03 | -2.206 | 0.678 | 0.110 |
| 1425T>A | [1018-NA] | 4 | 5612 | 0.071% | 2 | 293 | 0.683% | -0.611% | 2.47E-04 | -2.207 | 0.514 | 0.110 |
| 3601G>T | [1018-NA] | 4 | 5632 | 0.071% | 2 | 293 | 0.683% | -0.612% | 3.85E-06 | -1.788 | 0.337 | 0.167 |
| 2464A>G | [1018-NA] | 17 | 24022 | 0.071% | 2 | 293 | 0.683% | -0.612% | 2.36E-05 | -2.236 | 0.477 | 0.107 |
| 258G>C | [1018-NA] | 5 | 7083 | 0.071% | 2 | 293 | 0.683% | -0.612% | 4.08E-05 | -2.175 | 0.455 | 0.114 |
| 1896C>G | [1018-NA] | 2 | 2848 | 0.070% | 2 | 293 | 0.683% | -0.612% | 3.40E-04 | -1.010 | 0.254 | 0.364 |
| 3653G>T | [1018-NA] | 4 | 5704 | 0.070% | 2 | 293 | 0.683% | -0.612% | 4.70E-05 | -1.990 | 0.424 | 0.137 |
| 3350T>G | [1018-NA] | 4 | 5722 | 0.070% | 2 | 293 | 0.683% | -0.613% | 7.58E-05 | -1.894 | 0.420 | 0.150 |
| 2486A>G | [1018-NA] | 58 | 82987 | 0.070% | 2 | 293 | 0.683% | -0.613% | 1.32E-06 | -2.194 | 0.420 | 0.111 |
| 1951A>G | [1018-NA] | 1 | 1469 | 0.068% | 2 | 293 | 0.683% | -0.615% | 2.51E-02 | -1.026 | 0.404 | 0.359 |
| 257G>A | [1018-NA] | 3 | 4441 | 0.068% | 2 | 293 | 0.683% | -0.615% | 1.30E-03 | -2.260 | 0.600 | 0.104 |
| 3317G>T | [1018-NA] | 17 | 25228 | 0.067% | 2 | 293 | 0.683% | -0.615% | 7.08E-05 | -2.253 | 0.514 | 0.105 |
| 125A>G | [1018-NA] | 3 | 4453 | 0.067% | 2 | 293 | 0.683% | -0.615% | 4.46E-05 | -2.044 | 0.425 | 0.130 |
| 3597C>G | [1018-NA] | 2 | 2990 | 0.067% | 2 | 293 | 0.683% | -0.616% | 4.65E-06 | -1.765 | 0.322 | 0.171 |
| 710T>A | [1018-NA] | 1 | 1505 | 0.066% | 2 | 293 | 0.683% | -0.616% | 2.42E-03 | -2.130 | 0.576 | 0.119 |
| 75A>G | [1018-NA] | 4 | 6051 | 0.066% | 2 | 293 | 0.683% | -0.616% | 2.23E-06 | -2.038 | 0.367 | 0.130 |
| 133T>C | [1018-NA] | 5 | 7578 | 0.066% | 2 | 293 | 0.683% | -0.617% | 2.22E-05 | -2.165 | 0.441 | 0.115 |
| 1665A>G | [1018-NA] | 1 | 1549 | 0.065% | 2 | 293 | 0.683% | -0.618% | 2.86E-04 | -1.490 | 0.350 | 0.225 |
| 2818C>A | [1018-NA] | 2 | 3103 | 0.064% | 2 | 293 | 0.683% | -0.618% | 5.27E-04 | -2.161 | 0.522 | 0.115 |
| 2584T>G | [1018-NA] | 5 | 7815 | 0.064% | 2 | 293 | 0.683% | -0.619% | 1.28E-05 | -2.564 | 0.469 | 0.077 |
| 3244C>T | [1018-NA] | 9 | 14079 | 0.064% | 2 | 293 | 0.683% | -0.619% | 2.70E-02 | -1.827 | 0.740 | 0.161 |
| 2260T>A | [1018-NA] | 15 | 23900 | 0.063% | 2 | 293 | 0.683% | -0.620% | 6.33E-04 | -2.031 | 0.543 | 0.131 |
| 533T>A | [1018-NA] | 6 | 9572 | 0.063% | 2 | 293 | 0.683% | -0.620% | 2.50E-06 | -2.362 | 0.423 | 0.094 |
| 4023A>G | [1018-NA] | 1 | 1597 | 0.063% | 2 | 293 | 0.683% | -0.620% | 9.63E-04 | -2.336 | 0.560 | 0.097 |
| 3983T>G | [1018-NA] | 4 | 6507 | 0.061% | 2 | 293 | 0.683% | -0.621% | 1.45E-05 | -2.303 | 0.445 | 0.100 |
| 1678A>T | [1018-NA] | 1 | 1637 | 0.061% | 2 | 293 | 0.683% | -0.622% | 8.57E-05 | -1.523 | 0.327 | 0.218 |
| 492A>G | [1018-NA] | 1 | 1640 | 0.061% | 2 | 293 | 0.683% | -0.622% | 2.66E-03 | -2.129 | 0.587 | 0.119 |
| 2562T>G | [1018-NA] | 4 | 6570 | 0.061% | 2 | 293 | 0.683% | -0.622% | 8.46E-06 | -2.446 | 0.446 | 0.087 |
| 540C>G | [1018-NA] | 1 | 1646 | 0.061% | 2 | 293 | 0.683% | -0.622% | 1.68E-03 | -2.538 | 0.632 | 0.079 |
| 3961T>C | [1018-NA] | 6 | 10047 | 0.060% | 2 | 293 | 0.683% | -0.623% | 2.40E-08 | -2.185 | 0.331 | 0.112 |
| 3289C>T | [1018-NA] | 4 | 6758 | 0.059% | 2 | 293 | 0.683% | -0.623% | 3.11E-06 | -2.317 | 0.413 | 0.099 |
| 2476A>G | [1018-NA] | 14 | 24501 | 0.057% | 2 | 293 | 0.683% | -0.625% | 3.27E-08 | -2.331 | 0.374 | 0.097 |
| 205A>G | [1018-NA] | 3 | 5261 | 0.057% | 2 | 293 | 0.683% | -0.626% | 2.38E-06 | -2.253 | 0.392 | 0.105 |
| 2184T>G | [1018-NA] | 2 | 3529 | 0.057% | 2 | 293 | 0.683% | -0.626% | 3.06E-05 | -1.885 | 0.386 | 0.152 |
| 716A>G | [1018-NA] | 1 | 1806 | 0.055% | 2 | 293 | 0.683% | -0.627% | 4.93E-04 | -2.072 | 0.488 | 0.126 |
| 3815A>G | [1018-NA] | 7 | 12742 | 0.055% | 2 | 293 | 0.683% | -0.628% | 1.20E-04 | -2.505 | 0.569 | 0.082 |

| | | | | | | | | | | | | |
|---|---|---|---|---|---|---|---|---|---|---|---|---|
| 3349A>G | [1018-NA] | 3 | 5461 | 0.055% | 2 | 293 | 0.683% | -0.628% | 2.85E-06 | -2.011 | 0.368 | 0.134 |
| 3843T>A | [1018-NA] | 3 | 5527 | 0.054% | 2 | 293 | 0.683% | -0.628% | 3.58E-06 | -2.428 | 0.421 | 0.088 |
| 129G>T | [1018-NA] | 1 | 1859 | 0.054% | 2 | 293 | 0.683% | -0.629% | 1.24E-03 | -2.660 | 0.640 | 0.070 |
| 128A>T | [1018-NA] | 3 | 5658 | 0.053% | 2 | 293 | 0.683% | -0.630% | 4.65E-05 | -2.451 | 0.499 | 0.086 |
| 1562T>A | [1018-NA] | 4 | 7563 | 0.053% | 2 | 293 | 0.683% | -0.630% | 2.29E-08 | -2.347 | 0.337 | 0.096 |
| 2338T>A | [1018-NA] | 1 | 1891 | 0.053% | 2 | 293 | 0.683% | -0.630% | 7.08E-04 | -2.505 | 0.581 | 0.082 |
| 1532A>T | [1018-NA] | 5 | 9495 | 0.053% | 2 | 293 | 0.683% | -0.630% | 1.68E-06 | -2.259 | 0.408 | 0.104 |
| 824G>T | [1018-NA] | 2 | 3809 | 0.053% | 2 | 293 | 0.683% | -0.630% | 5.12E-06 | -1.916 | 0.355 | 0.147 |
| 442A>T | [1018-NA] | 24 | 46811 | 0.051% | 2 | 293 | 0.683% | -0.631% | 4.47E-10 | -2.462 | 0.353 | 0.085 |
| 3512T>A | [1018-NA] | 3 | 5856 | 0.051% | 2 | 293 | 0.683% | -0.631% | 1.96E-07 | -2.303 | 0.357 | 0.100 |
| 1735C>G | [1018-NA] | 1 | 1982 | 0.050% | 2 | 293 | 0.683% | -0.632% | 1.79E-04 | -1.547 | 0.355 | 0.213 |
| 3508G>A | [1018-NA] | 10 | 19835 | 0.050% | 2 | 293 | 0.683% | -0.632% | 2.05E-07 | -2.124 | 0.368 | 0.120 |
| 3097A>G | [1018-NA] | 4 | 7964 | 0.050% | 2 | 293 | 0.683% | -0.632% | 8.28E-07 | -2.169 | 0.379 | 0.114 |
| 1853T>G | [1018-NA] | 2 | 3983 | 0.050% | 2 | 293 | 0.683% | -0.632% | 1.99E-05 | -1.299 | 0.272 | 0.273 |
| 3057C>T | [1018-NA] | 3 | 5979 | 0.050% | 2 | 293 | 0.683% | -0.632% | 5.41E-07 | -2.244 | 0.371 | 0.106 |
| 1994A>G | [1018-NA] | 1 | 2000 | 0.050% | 2 | 293 | 0.683% | -0.633% | 9.31E-04 | -1.055 | 0.284 | 0.348 |
| 3347G>T | [1018-NA] | 2 | 4010 | 0.050% | 2 | 293 | 0.683% | -0.633% | 4.60E-03 | -2.124 | 0.655 | 0.120 |
| 3310G>A | [1018-NA] | 1 | 2018 | 0.050% | 2 | 293 | 0.683% | -0.633% | 1.44E-03 | -2.296 | 0.594 | 0.101 |
| 3820A>T | [1018-NA] | 5 | 10100 | 0.050% | 2 | 293 | 0.683% | -0.633% | 8.35E-06 | -2.636 | 0.493 | 0.072 |
| 429A>T | [1018-NA] | 4 | 8094 | 0.049% | 2 | 293 | 0.683% | -0.633% | 2.03E-02 | -2.185 | 0.835 | 0.112 |
| 3279G>T | [1018-NA] | 1 | 2032 | 0.049% | 2 | 293 | 0.683% | -0.633% | 2.52E-03 | -2.303 | 0.633 | 0.100 |
| 3176G>T | [1018-NA] | 6 | 12207 | 0.049% | 2 | 293 | 0.683% | -0.633% | 1.22E-06 | -2.304 | 0.416 | 0.100 |
| 3837G>T | [1018-NA] | 4 | 8160 | 0.049% | 2 | 293 | 0.683% | -0.634% | 6.40E-03 | -2.505 | 0.806 | 0.082 |
| 1869A>G | [1018-NA] | 2 | 4083 | 0.049% | 2 | 293 | 0.683% | -0.634% | 1.18E-04 | -1.058 | 0.249 | 0.347 |
| 1419T>G | [1018-NA] | 2 | 4131 | 0.048% | 2 | 293 | 0.683% | -0.634% | 2.45E-05 | -2.517 | 0.474 | 0.081 |
| 278T>G | [1018-NA] | 12 | 24797 | 0.048% | 2 | 293 | 0.683% | -0.634% | 1.52E-07 | -2.252 | 0.387 | 0.105 |
| 2278T>G | [1018-NA] | 1 | 2068 | 0.048% | 2 | 293 | 0.683% | -0.634% | 7.15E-05 | -1.779 | 0.376 | 0.169 |
| 3019A>T | [1018-NA] | 4 | 8338 | 0.048% | 2 | 293 | 0.683% | -0.635% | 1.22E-06 | -2.299 | 0.405 | 0.100 |
| 3653G>A | [1018-NA] | 2 | 4181 | 0.048% | 2 | 293 | 0.683% | -0.635% | 4.78E-07 | -2.085 | 0.339 | 0.124 |
| 1578A>G | [1018-NA] | 8 | 16849 | 0.047% | 2 | 293 | 0.683% | -0.635% | 1.17E-07 | -2.339 | 0.388 | 0.096 |
| 3410T>A | [1018-NA] | 10 | 21132 | 0.047% | 2 | 293 | 0.683% | -0.635% | 1.59E-05 | -2.065 | 0.437 | 0.127 |
| 4014T>A | [1018-NA] | 2 | 4279 | 0.047% | 2 | 293 | 0.683% | -0.636% | 1.59E-04 | -2.296 | 0.514 | 0.101 |
| 721T>G | [1018-NA] | 2 | 4346 | 0.046% | 2 | 293 | 0.683% | -0.637% | 1.39E-02 | -1.876 | 0.676 | 0.153 |
| 3526A>C | [1018-NA] | 1 | 2190 | 0.046% | 2 | 293 | 0.683% | -0.637% | 2.45E-05 | -1.986 | 0.383 | 0.137 |
| 2180T>A | [1018-NA] | 1 | 2202 | 0.045% | 2 | 293 | 0.683% | -0.637% | 2.02E-02 | -2.657 | 0.952 | 0.070 |
| 1533G>A | [1018-NA] | 1 | 2283 | 0.044% | 2 | 293 | 0.683% | -0.639% | 1.44E-04 | -2.204 | 0.473 | 0.110 |
| 247C>G | [1018-NA] | 1 | 2295 | 0.044% | 2 | 293 | 0.683% | -0.639% | 7.13E-05 | -2.311 | 0.464 | 0.099 |
| 257G>C | [1018-NA] | 2 | 4608 | 0.043% | 2 | 293 | 0.683% | -0.639% | 2.97E-05 | -2.121 | 0.435 | 0.120 |
| 2570T>G | [1018-NA] | 2 | 4644 | 0.043% | 2 | 293 | 0.683% | -0.640% | 1.91E-04 | -2.710 | 0.588 | 0.067 |
| 398T>A | [1018-NA] | 4 | 9326 | 0.043% | 2 | 293 | 0.683% | -0.640% | 2.66E-10 | -2.250 | 0.292 | 0.105 |
| 1503A>G | [1018-NA] | 3 | 7053 | 0.043% | 2 | 293 | 0.683% | -0.640% | 3.33E-07 | -2.347 | 0.383 | 0.096 |
| 2524T>A | [1018-NA] | 15 | 35274 | 0.043% | 2 | 293 | 0.683% | -0.640% | 2.18E-03 | -2.785 | 0.820 | 0.062 |
| 3522A>G | [1018-NA] | 3 | 7153 | 0.042% | 2 | 293 | 0.683% | -0.641% | 1.10E-06 | -2.156 | 0.381 | 0.116 |
| 268A>C | [1018-NA] | 3 | 7195 | 0.042% | 2 | 293 | 0.683% | -0.641% | 7.88E-09 | -2.429 | 0.333 | 0.088 |
| 1533G>C | [1018-NA] | 1 | 2422 | 0.041% | 2 | 293 | 0.683% | -0.641% | 2.17E-04 | -2.439 | 0.529 | 0.087 |
| 2551T>G | [1018-NA] | 2 | 4854 | 0.041% | 2 | 293 | 0.683% | -0.641% | 5.03E-04 | -2.608 | 0.627 | 0.074 |

| | | | | | | | | | | | | |
|---|---|---|---|---|---|---|---|---|---|---|---|---|
| 2105T>A | [1018-NA] | 3 | 7457 | 0.040% | 2 | 293 | 0.683% | -0.642% | 1.16E-09 | -2.043 | 0.281 | 0.130 |
| 90A>T | [1018-NA] | 3 | 7480 | 0.040% | 2 | 293 | 0.683% | -0.642% | 6.25E-08 | -2.303 | 0.354 | 0.100 |
| 2065A>T | [1018-NA] | 1 | 2526 | 0.040% | 2 | 293 | 0.683% | -0.643% | 3.80E-06 | -1.264 | 0.239 | 0.282 |
| 596A>C | [1018-NA] | 1 | 2551 | 0.039% | 2 | 293 | 0.683% | -0.643% | 2.60E-04 | -2.804 | 0.590 | 0.061 |
| 111T>A | [1018-NA] | 1 | 2572 | 0.039% | 2 | 293 | 0.683% | -0.644% | 1.54E-04 | -2.231 | 0.487 | 0.107 |
| 2532A>C | [1018-NA] | 8 | 20858 | 0.038% | 2 | 293 | 0.683% | -0.644% | 1.05E-09 | -2.807 | 0.378 | 0.060 |
| 1765A>G | [1018-NA] | 1 | 2609 | 0.038% | 2 | 293 | 0.683% | -0.644% | 4.48E-05 | -1.730 | 0.363 | 0.177 |
| 3293A>T | [1018-NA] | 3 | 7911 | 0.038% | 2 | 293 | 0.683% | -0.645% | 2.99E-08 | -2.349 | 0.351 | 0.096 |
| 763T>A | [1018-NA] | 1 | 2668 | 0.037% | 2 | 293 | 0.683% | -0.645% | 3.63E-06 | -2.033 | 0.357 | 0.131 |
| 3229C>G | [1018-NA] | 4 | 10732 | 0.037% | 2 | 293 | 0.683% | -0.645% | 1.87E-07 | -2.366 | 0.391 | 0.094 |
| 3386C>A | [1018-NA] | 3 | 8113 | 0.037% | 2 | 293 | 0.683% | -0.646% | 1.47E-07 | -2.119 | 0.348 | 0.120 |
| 4012T>A | [1018-NA] | 2 | 5439 | 0.037% | 2 | 293 | 0.683% | -0.646% | 4.82E-08 | -2.481 | 0.353 | 0.084 |
| 3512T>C | [1018-NA] | 3 | 8171 | 0.037% | 2 | 293 | 0.683% | -0.646% | 1.37E-04 | -2.260 | 0.525 | 0.104 |
| 1817C>A | [1018-NA] | 1 | 2724 | 0.037% | 2 | 293 | 0.683% | -0.646% | 9.60E-07 | -1.627 | 0.280 | 0.197 |
| 3577A>G | [1018-NA] | 7 | 19845 | 0.035% | 2 | 293 | 0.683% | -0.647% | 6.67E-04 | -2.052 | 0.551 | 0.128 |
| 3529T>G | [1018-NA] | 15 | 43213 | 0.035% | 2 | 293 | 0.683% | -0.648% | 5.91E-10 | -2.148 | 0.316 | 0.117 |
| 2642A>G | [1018-NA] | 3 | 8740 | 0.034% | 2 | 293 | 0.683% | -0.648% | 2.43E-07 | -2.791 | 0.429 | 0.061 |
| 3565C>T | [1018-NA] | 3 | 8759 | 0.034% | 2 | 293 | 0.683% | -0.648% | 2.65E-04 | -2.238 | 0.547 | 0.107 |
| 3911A>G | [1018-NA] | 7 | 20670 | 0.034% | 2 | 293 | 0.683% | -0.649% | 1.52E-05 | -2.682 | 0.552 | 0.068 |
| 3392A>G | [1018-NA] | 2 | 5915 | 0.034% | 2 | 293 | 0.683% | -0.649% | 1.76E-10 | -2.102 | 0.264 | 0.122 |
| 1692G>A | [1018-NA] | 1 | 2979 | 0.034% | 2 | 293 | 0.683% | -0.649% | 6.09E-07 | -1.657 | 0.281 | 0.191 |
| 2532A>T | [1018-NA] | 6 | 17953 | 0.033% | 2 | 293 | 0.683% | -0.649% | 3.27E-09 | -2.938 | 0.398 | 0.053 |
| 3924T>G | [1018-NA] | 7 | 21610 | 0.032% | 2 | 293 | 0.683% | -0.650% | 1.88E-11 | -2.613 | 0.329 | 0.073 |
| 3084A>G | [1018-NA] | 2 | 6343 | 0.032% | 2 | 293 | 0.683% | -0.651% | 5.86E-09 | -2.242 | 0.314 | 0.106 |
| 4008T>A | [1018-NA] | 1 | 3196 | 0.031% | 2 | 293 | 0.683% | -0.651% | 2.47E-05 | -2.509 | 0.475 | 0.081 |
| 3974T>C | [1018-NA] | 3 | 9624 | 0.031% | 2 | 293 | 0.683% | -0.651% | 2.74E-06 | -2.634 | 0.478 | 0.072 |
| 3974T>A | [1018-NA] | 2 | 6461 | 0.031% | 2 | 293 | 0.683% | -0.652% | 3.45E-08 | -2.416 | 0.355 | 0.089 |
| 2282A>T | [1018-NA] | 2 | 6577 | 0.030% | 2 | 293 | 0.683% | -0.652% | 4.57E-09 | -2.254 | 0.314 | 0.105 |
| 764G>A | [1018-NA] | 1 | 3314 | 0.030% | 2 | 293 | 0.683% | -0.652% | 1.02E-05 | -1.853 | 0.359 | 0.157 |
| 3884G>A | [1018-NA] | 1 | 3314 | 0.030% | 2 | 293 | 0.683% | -0.652% | 1.18E-03 | -2.577 | 0.667 | 0.076 |
| 3311G>C | [1018-NA] | 1 | 3349 | 0.030% | 2 | 293 | 0.683% | -0.653% | 1.75E-05 | -2.930 | 0.507 | 0.053 |
| 2989C>G | [1018-NA] | 1 | 3389 | 0.030% | 2 | 293 | 0.683% | -0.653% | 1.52E-05 | -2.423 | 0.453 | 0.089 |
| 3173C>A | [1018-NA] | 1 | 3409 | 0.029% | 2 | 293 | 0.683% | -0.653% | 1.39E-05 | -2.707 | 0.482 | 0.067 |
| 3037A>G | [1018-NA] | 10 | 34387 | 0.029% | 2 | 293 | 0.683% | -0.654% | 1.91E-12 | -2.362 | 0.297 | 0.094 |
| 4032T>G | [1018-NA] | 8 | 27713 | 0.029% | 2 | 293 | 0.683% | -0.654% | 6.75E-05 | -2.518 | 0.576 | 0.081 |
| 2163T>G | [1018-NA] | 1 | 3471 | 0.029% | 2 | 293 | 0.683% | -0.654% | 2.20E-05 | -2.229 | 0.438 | 0.108 |
| 2383T>G | [1018-NA] | 1 | 3493 | 0.029% | 2 | 293 | 0.683% | -0.654% | 1.93E-06 | -2.630 | 0.422 | 0.072 |
| 3692A>T | [1018-NA] | 2 | 6991 | 0.029% | 2 | 293 | 0.683% | -0.654% | 3.22E-08 | -2.095 | 0.323 | 0.123 |
| 2350T>G | [1018-NA] | 1 | 3507 | 0.029% | 2 | 293 | 0.683% | -0.654% | 6.08E-05 | -2.541 | 0.517 | 0.079 |
| 3991G>A | [1018-NA] | 2 | 7107 | 0.028% | 2 | 293 | 0.683% | -0.654% | 2.13E-03 | -2.540 | 0.728 | 0.079 |
| 2154T>G | [1018-NA] | 1 | 3574 | 0.028% | 2 | 293 | 0.683% | -0.655% | 5.63E-07 | -2.080 | 0.342 | 0.125 |
| 3824C>T | [1018-NA] | 2 | 7158 | 0.028% | 2 | 293 | 0.683% | -0.655% | 8.31E-06 | -2.869 | 0.523 | 0.057 |
| 3605A>G | [1018-NA] | 3 | 10853 | 0.028% | 2 | 293 | 0.683% | -0.655% | 1.67E-09 | -2.167 | 0.310 | 0.114 |
| 3007C>G | [1018-NA] | 4 | 14559 | 0.027% | 2 | 293 | 0.683% | -0.655% | 2.09E-12 | -2.373 | 0.282 | 0.093 |
| 4020T>A | [1018-NA] | 1 | 3681 | 0.027% | 2 | 293 | 0.683% | -0.655% | 6.26E-05 | -2.619 | 0.532 | 0.073 |
| 3331C>T | [1018-NA] | 1 | 3760 | 0.027% | 2 | 293 | 0.683% | -0.656% | 1.56E-08 | -2.449 | 0.322 | 0.086 |

| | | | | | | | | | | | | |
|---|---|---|---|---|---|---|---|---|---|---|---|---|
| 156C>G | [1018-NA] | 1 | 3821 | 0.026% | 2 | 293 | 0.683% | -0.656% | 1.55E-06 | -2.490 | 0.411 | 0.083 |
| 304G>C | [1018-NA] | 2 | 7739 | 0.026% | 2 | 293 | 0.683% | -0.657% | 1.61E-06 | -2.440 | 0.436 | 0.087 |
| 1684T>G | [1018-NA] | 1 | 3906 | 0.026% | 2 | 293 | 0.683% | -0.657% | 1.66E-05 | -1.572 | 0.323 | 0.208 |
| 1450A>G | [1018-NA] | 1 | 4160 | 0.024% | 2 | 293 | 0.683% | -0.659% | 3.00E-07 | -2.550 | 0.387 | 0.078 |
| 2408A>G | [1018-NA] | 2 | 8475 | 0.024% | 2 | 293 | 0.683% | -0.659% | 4.65E-08 | -2.531 | 0.385 | 0.080 |
| 3993T>G | [1018-NA] | 1 | 4280 | 0.023% | 2 | 293 | 0.683% | -0.659% | 4.95E-06 | -2.529 | 0.450 | 0.080 |
| 1725T>A | [1018-NA] | 1 | 4376 | 0.023% | 2 | 293 | 0.683% | -0.660% | 6.77E-09 | -1.675 | 0.245 | 0.187 |
| 3843T>C | [1018-NA] | 2 | 9236 | 0.022% | 2 | 293 | 0.683% | -0.661% | 1.69E-07 | -2.816 | 0.440 | 0.060 |
| 3670C>T | [1018-NA] | 2 | 9338 | 0.021% | 2 | 293 | 0.683% | -0.661% | 7.28E-03 | -2.243 | 0.750 | 0.106 |
| 2284A>T | [1018-NA] | 2 | 9434 | 0.021% | 2 | 293 | 0.683% | -0.661% | 8.24E-13 | -2.138 | 0.246 | 0.118 |
| 304G>T | [1018-NA] | 1 | 4898 | 0.020% | 2 | 293 | 0.683% | -0.662% | 3.41E-07 | -2.531 | 0.399 | 0.080 |
| 3511G>A | [1018-NA] | 3 | 16888 | 0.018% | 2 | 293 | 0.683% | -0.665% | 2.68E-15 | -2.298 | 0.242 | 0.100 |
| 3507T>C | [1018-NA] | 2 | 11604 | 0.017% | 2 | 293 | 0.683% | -0.665% | 7.34E-17 | -2.359 | 0.219 | 0.094 |
| 3953T>G | [1018-NA] | 2 | 11751 | 0.017% | 2 | 293 | 0.683% | -0.666% | 1.08E-10 | -2.882 | 0.351 | 0.056 |
| 3959G>T | [1018-NA] | 1 | 5998 | 0.017% | 2 | 293 | 0.683% | -0.666% | 8.31E-08 | -2.597 | 0.390 | 0.074 |
| 3797G>T | [1018-NA] | 1 | 6036 | 0.017% | 2 | 293 | 0.683% | -0.666% | 1.50E-06 | -3.391 | 0.512 | 0.034 |
| 3400G>A | [1018-NA] | 2 | 12139 | 0.016% | 2 | 293 | 0.683% | -0.666% | 2.43E-13 | -2.235 | 0.255 | 0.107 |
| 3642T>G | [1018-NA] | 1 | 6592 | 0.015% | 2 | 293 | 0.683% | -0.667% | 2.28E-06 | -2.268 | 0.416 | 0.103 |
| 3409T>G | [1018-NA] | 3 | 20235 | 0.015% | 2 | 293 | 0.683% | -0.668% | 1.58E-13 | -2.370 | 0.275 | 0.094 |
| 2257C>G | [1018-NA] | 1 | 7119 | 0.014% | 2 | 293 | 0.683% | -0.669% | 2.17E-13 | -2.222 | 0.237 | 0.108 |
| 3507T>A | [1018-NA] | 2 | 14416 | 0.014% | 2 | 293 | 0.683% | -0.669% | 4.96E-19 | -2.419 | 0.211 | 0.089 |
| 305A>C | [1018-NA] | 1 | 7366 | 0.014% | 2 | 293 | 0.683% | -0.669% | 1.54E-13 | -2.479 | 0.250 | 0.084 |
| 2219A>G | [1018-NA] | 1 | 7838 | 0.013% | 2 | 293 | 0.683% | -0.670% | 5.70E-10 | -2.277 | 0.306 | 0.103 |
| 748T>G | [1018-NA] | 1 | 8402 | 0.012% | 2 | 293 | 0.683% | -0.671% | 1.48E-12 | -2.059 | 0.242 | 0.128 |
| 303G>C | [1018-NA] | 1 | 8567 | 0.012% | 2 | 293 | 0.683% | -0.671% | 3.71E-13 | -2.727 | 0.277 | 0.065 |
| 3363C>G | [1018-NA] | 1 | 8594 | 0.012% | 2 | 293 | 0.683% | -0.671% | 1.24E-08 | -2.330 | 0.349 | 0.097 |
| 394C>G | [1018-NA] | 1 | 8841 | 0.011% | 2 | 293 | 0.683% | -0.671% | 2.20E-12 | -2.788 | 0.297 | 0.062 |
| 2252T>A | [1018-NA] | 1 | 9290 | 0.011% | 2 | 293 | 0.683% | -0.672% | 1.09E-14 | -2.333 | 0.237 | 0.097 |
| 4016T>A | [1018-NA] | 1 | 10320 | 0.010% | 2 | 293 | 0.683% | -0.673% | 1.32E-11 | -2.790 | 0.322 | 0.061 |
| 3526A>T | [1018-NA] | 1 | 12550 | 0.008% | 2 | 293 | 0.683% | -0.675% | 1.25E-15 | -2.430 | 0.243 | 0.088 |
| 3399T>A | [1018-NA] | 0 | 8742 | 0.000% | 2 | 293 | 0.683% | -0.683% | 3.98E-17 | -2.527 | 0.219 | 0.080 |
| 2260T>G | [1018-NA] | 0 | 11260 | 0.000% | 2 | 293 | 0.683% | -0.683% | 3.73E-15 | -2.265 | 0.235 | 0.104 |
| 2231T>G | [1018-NA] | 0 | 7536 | 0.000% | 2 | 293 | 0.683% | -0.683% | 1.36E-14 | -2.439 | 0.240 | 0.087 |
| 728T>A | [1018-NA] | 0 | 5963 | 0.000% | 2 | 293 | 0.683% | -0.683% | 1.70E-14 | -1.975 | 0.203 | 0.139 |
| 3600T>A | [1018-NA] | 0 | 5922 | 0.000% | 2 | 293 | 0.683% | -0.683% | 1.83E-13 | -2.358 | 0.241 | 0.095 |
| 835T>G | [1018-NA] | 0 | 5293 | 0.000% | 2 | 293 | 0.683% | -0.683% | 5.07E-13 | -2.124 | 0.229 | 0.120 |
| 882T>G | [1018-NA] | 0 | 5229 | 0.000% | 2 | 293 | 0.683% | -0.683% | 4.93E-12 | -2.062 | 0.238 | 0.127 |
| 2240C>A | [1018-NA] | 0 | 4911 | 0.000% | 2 | 293 | 0.683% | -0.683% | 5.87E-12 | -2.231 | 0.250 | 0.107 |
| 770G>C | [1018-NA] | 0 | 4060 | 0.000% | 2 | 293 | 0.683% | -0.683% | 1.46E-11 | -2.072 | 0.238 | 0.126 |
| 820T>A | [1018-NA] | 0 | 4185 | 0.000% | 2 | 293 | 0.683% | -0.683% | 1.70E-11 | -2.256 | 0.253 | 0.105 |
| 2084A>T | [1018-NA] | 0 | 4388 | 0.000% | 2 | 293 | 0.683% | -0.683% | 6.67E-11 | -2.304 | 0.271 | 0.100 |
| 2181G>A | [1018-NA] | 0 | 5184 | 0.000% | 2 | 293 | 0.683% | -0.683% | 1.67E-10 | -2.527 | 0.303 | 0.080 |
| 3524T>A | [1018-NA] | 0 | 4045 | 0.000% | 2 | 293 | 0.683% | -0.683% | 1.71E-10 | -2.260 | 0.272 | 0.104 |
| 1682A>T | [1018-NA] | 0 | 4041 | 0.000% | 2 | 293 | 0.683% | -0.683% | 1.97E-10 | -1.817 | 0.234 | 0.163 |
| 814A>G | [1018-NA] | 0 | 3583 | 0.000% | 2 | 293 | 0.683% | -0.683% | 3.75E-10 | -2.013 | 0.254 | 0.134 |
| 208T>G | [1018-NA] | 0 | 5196 | 0.000% | 2 | 293 | 0.683% | -0.683% | 4.47E-10 | -3.060 | 0.345 | 0.047 |

| | | | | | | | | | | | | |
|---|---|---|---|---|---|---|---|---|---|---|---|---|
| 1605G>C | [1018-NA] | 0 | 4061 | 0.000% | 2 | 293 | 0.683% | -0.683% | 1.64E-09 | -2.718 | 0.329 | 0.066 |
| 3979T>C | [1018-NA] | 0 | 4378 | 0.000% | 2 | 293 | 0.683% | -0.683% | 2.28E-09 | -2.652 | 0.334 | 0.070 |
| 1961A>G | [1018-NA] | 0 | 3159 | 0.000% | 2 | 293 | 0.683% | -0.683% | 3.24E-09 | -1.361 | 0.196 | 0.256 |
| 3659C>G | [1018-NA] | 0 | 3387 | 0.000% | 2 | 293 | 0.683% | -0.683% | 3.94E-09 | -2.205 | 0.292 | 0.110 |
| 3601G>C | [1018-NA] | 0 | 3389 | 0.000% | 2 | 293 | 0.683% | -0.683% | 5.95E-09 | -2.397 | 0.314 | 0.091 |
| 231T>G | [1018-NA] | 0 | 3702 | 0.000% | 2 | 293 | 0.683% | -0.683% | 6.41E-09 | -2.721 | 0.344 | 0.066 |
| 2247T>G | [1018-NA] | 0 | 5577 | 0.000% | 2 | 293 | 0.683% | -0.683% | 9.42E-09 | -2.228 | 0.324 | 0.108 |
| 1666T>G | [1018-NA] | 0 | 2575 | 0.000% | 2 | 293 | 0.683% | -0.683% | 1.16E-08 | -1.889 | 0.263 | 0.151 |
| 623T>G | [1018-NA] | 0 | 3559 | 0.000% | 2 | 293 | 0.683% | -0.683% | 1.52E-08 | -2.445 | 0.333 | 0.087 |
| 769A>G | [1018-NA] | 0 | 2527 | 0.000% | 2 | 293 | 0.683% | -0.683% | 2.85E-08 | -2.003 | 0.285 | 0.135 |
| 3832A>T | [1018-NA] | 0 | 4348 | 0.000% | 2 | 293 | 0.683% | -0.683% | 3.66E-08 | -3.105 | 0.410 | 0.045 |
| 549G>A | [1018-NA] | 0 | 2953 | 0.000% | 2 | 293 | 0.683% | -0.683% | 3.89E-08 | -2.400 | 0.334 | 0.091 |
| 813C>G | [1018-NA] | 0 | 2456 | 0.000% | 2 | 293 | 0.683% | -0.683% | 4.49E-08 | -2.129 | 0.302 | 0.119 |
| 3633A>G | [1018-NA] | 0 | 3476 | 0.000% | 2 | 293 | 0.683% | -0.683% | 4.73E-08 | -2.625 | 0.366 | 0.072 |
| 2284A>C | [1018-NA] | 0 | 2535 | 0.000% | 2 | 293 | 0.683% | -0.683% | 5.19E-08 | -2.081 | 0.301 | 0.125 |
| 823A>C | [1018-NA] | 0 | 2472 | 0.000% | 2 | 293 | 0.683% | -0.683% | 5.56E-08 | -2.056 | 0.298 | 0.128 |
| 4008T>G | [1018-NA] | 0 | 4867 | 0.000% | 2 | 293 | 0.683% | -0.683% | 6.50E-08 | -3.177 | 0.434 | 0.042 |
| 1637T>C | [1018-NA] | 0 | 2581 | 0.000% | 2 | 293 | 0.683% | -0.683% | 6.60E-08 | -2.025 | 0.299 | 0.132 |
| 914G>T | [1018-NA] | 0 | 2451 | 0.000% | 2 | 293 | 0.683% | -0.683% | 7.91E-08 | -1.861 | 0.280 | 0.156 |
| 3325A>G | [1018-NA] | 0 | 3227 | 0.000% | 2 | 293 | 0.683% | -0.683% | 7.98E-08 | -2.807 | 0.385 | 0.060 |
| 2184T>A | [1018-NA] | 0 | 2394 | 0.000% | 2 | 293 | 0.683% | -0.683% | 1.00E-07 | -1.926 | 0.290 | 0.146 |
| 2214T>G | [1018-NA] | 0 | 3039 | 0.000% | 2 | 293 | 0.683% | -0.683% | 1.95E-07 | -2.459 | 0.368 | 0.085 |
| 3855C>T | [1018-NA] | 0 | 3230 | 0.000% | 2 | 293 | 0.683% | -0.683% | 3.47E-07 | -2.690 | 0.406 | 0.068 |
| 657C>G | [1018-NA] | 0 | 4425 | 0.000% | 2 | 293 | 0.683% | -0.683% | 4.28E-07 | -3.410 | 0.490 | 0.033 |
| 4003T>A | [1018-NA] | 0 | 3699 | 0.000% | 2 | 293 | 0.683% | -0.683% | 5.95E-07 | -2.986 | 0.453 | 0.051 |
| 2563G>A | [1018-NA] | 0 | 5593 | 0.000% | 2 | 293 | 0.683% | -0.683% | 6.92E-07 | -4.049 | 0.563 | 0.017 |
| 1689T>G | [1018-NA] | 0 | 2120 | 0.000% | 2 | 293 | 0.683% | -0.683% | 8.78E-07 | -1.852 | 0.308 | 0.157 |
| 510A>T | [1018-NA] | 0 | 3326 | 0.000% | 2 | 293 | 0.683% | -0.683% | 1.15E-06 | -3.307 | 0.491 | 0.037 |
| 2353T>A | [1018-NA] | 0 | 2647 | 0.000% | 2 | 293 | 0.683% | -0.683% | 1.21E-06 | -2.896 | 0.443 | 0.055 |
| 2459T>A | [1018-NA] | 0 | 3089 | 0.000% | 2 | 293 | 0.683% | -0.683% | 1.43E-06 | -2.996 | 0.466 | 0.050 |
| 2519A>C | [1018-NA] | 0 | 6129 | 0.000% | 2 | 293 | 0.683% | -0.683% | 1.65E-06 | -3.735 | 0.582 | 0.024 |
| 68A>G | [1018-NA] | 0 | 3022 | 0.000% | 2 | 293 | 0.683% | -0.683% | 1.81E-06 | -3.211 | 0.490 | 0.040 |
| 2229T>A | [1018-NA] | 0 | 2348 | 0.000% | 2 | 293 | 0.683% | -0.683% | 1.85E-06 | -2.371 | 0.392 | 0.093 |
| 823A>G | [1018-NA] | 0 | 2036 | 0.000% | 2 | 293 | 0.683% | -0.683% | 2.03E-06 | -2.029 | 0.345 | 0.132 |
| 594A>G | [1018-NA] | 0 | 2906 | 0.000% | 2 | 293 | 0.683% | -0.683% | 2.80E-06 | -2.989 | 0.481 | 0.050 |
| 438A>T | [1018-NA] | 0 | 2591 | 0.000% | 2 | 293 | 0.683% | -0.683% | 4.34E-06 | -3.057 | 0.493 | 0.047 |
| 1670G>A | [1018-NA] | 0 | 1639 | 0.000% | 2 | 293 | 0.683% | -0.683% | 4.52E-06 | -1.874 | 0.331 | 0.153 |
| 2011C>G | [1018-NA] | 0 | 1496 | 0.000% | 2 | 293 | 0.683% | -0.683% | 4.93E-06 | -1.327 | 0.246 | 0.265 |
| 618A>G | [1018-NA] | 0 | 2639 | 0.000% | 2 | 293 | 0.683% | -0.683% | 7.09E-06 | -2.739 | 0.475 | 0.065 |
| 3054A>T | [1018-NA] | 0 | 1806 | 0.000% | 2 | 293 | 0.683% | -0.683% | 7.68E-06 | -2.360 | 0.412 | 0.094 |
| 1969C>T | [1018-NA] | 0 | 1548 | 0.000% | 2 | 293 | 0.683% | -0.683% | 9.49E-06 | -1.341 | 0.258 | 0.262 |
| 596A>G | [1018-NA] | 0 | 2351 | 0.000% | 2 | 293 | 0.683% | -0.683% | 1.39E-05 | -2.960 | 0.516 | 0.052 |
| 2013T>G | [1018-NA] | 0 | 1501 | 0.000% | 2 | 293 | 0.683% | -0.683% | 1.58E-05 | -1.532 | 0.298 | 0.216 |
| 2514T>G | [1018-NA] | 0 | 2883 | 0.000% | 2 | 293 | 0.683% | -0.683% | 1.66E-05 | -2.982 | 0.537 | 0.051 |
| 493T>C | [1018-NA] | 0 | 1939 | 0.000% | 2 | 293 | 0.683% | -0.683% | 2.22E-05 | -2.431 | 0.455 | 0.088 |
| 601T>G | [1018-NA] | 0 | 2316 | 0.000% | 2 | 293 | 0.683% | -0.683% | 2.56E-05 | -2.945 | 0.537 | 0.053 |

| | | | | | | | | | | | | |
|---|---|---|---|---|---|---|---|---|---|---|---|---|
| 1714T>A | [1018-NA] | 0 | 1607 | 0.000% | 2 | 293 | 0.683% | -0.683% | 4.00E-05 | -1.855 | 0.374 | 0.156 |
| 2113T>G | [1018-NA] | 0 | 1614 | 0.000% | 2 | 293 | 0.683% | -0.683% | 4.10E-05 | -2.297 | 0.446 | 0.101 |
| 2156G>T | [1018-NA] | 0 | 1508 | 0.000% | 2 | 293 | 0.683% | -0.683% | 4.20E-05 | -2.229 | 0.434 | 0.108 |
| 2471A>G | [1018-NA] | 0 | 2066 | 0.000% | 2 | 293 | 0.683% | -0.683% | 4.25E-05 | -3.054 | 0.565 | 0.047 |
| 545A>G | [1018-NA] | 0 | 1396 | 0.000% | 2 | 293 | 0.683% | -0.683% | 4.45E-05 | -2.103 | 0.413 | 0.122 |
| 549G>T | [1018-NA] | 0 | 2091 | 0.000% | 2 | 293 | 0.683% | -0.683% | 4.50E-05 | -3.066 | 0.570 | 0.047 |
| 1640C>G | [1018-NA] | 0 | 1279 | 0.000% | 2 | 293 | 0.683% | -0.683% | 5.52E-05 | -1.765 | 0.360 | 0.171 |
| 539C>G | [1018-NA] | 0 | 1775 | 0.000% | 2 | 293 | 0.683% | -0.683% | 5.73E-05 | -2.679 | 0.519 | 0.069 |
| 727T>G | [1018-NA] | 0 | 1263 | 0.000% | 2 | 293 | 0.683% | -0.683% | 6.19E-05 | -1.870 | 0.381 | 0.154 |
| 3523T>G | [1018-NA] | 0 | 1394 | 0.000% | 2 | 293 | 0.683% | -0.683% | 9.79E-05 | -2.256 | 0.464 | 0.105 |
| 1661T>G | [1018-NA] | 0 | 1259 | 0.000% | 2 | 293 | 0.683% | -0.683% | 1.35E-04 | -2.000 | 0.427 | 0.135 |
| 1534T>G | [1018-NA] | 0 | 1579 | 0.000% | 2 | 293 | 0.683% | -0.683% | 1.59E-04 | -2.562 | 0.539 | 0.077 |
| 3880T>G | [1018-NA] | 0 | 1627 | 0.000% | 2 | 293 | 0.683% | -0.683% | 1.78E-04 | -2.815 | 0.586 | 0.060 |
| 3651T>A | [1018-NA] | 0 | 1210 | 0.000% | 2 | 293 | 0.683% | -0.683% | 1.79E-04 | -2.009 | 0.437 | 0.134 |
| 3519T>A | [1018-NA] | 0 | 1714 | 0.000% | 2 | 293 | 0.683% | -0.683% | 1.89E-04 | -2.644 | 0.564 | 0.071 |
| 1872A>G | [1018-NA] | 0 | 1077 | 0.000% | 2 | 293 | 0.683% | -0.683% | 2.30E-04 | -1.522 | 0.348 | 0.218 |
| 267A>T | [1018-NA] | 0 | 1795 | 0.000% | 2 | 293 | 0.683% | -0.683% | 2.50E-04 | -3.201 | 0.671 | 0.041 |
| 130T>G | [1018-NA] | 0 | 1179 | 0.000% | 2 | 293 | 0.683% | -0.683% | 2.65E-04 | -2.270 | 0.499 | 0.103 |
| 906G>A | [1018-NA] | 0 | 1232 | 0.000% | 2 | 293 | 0.683% | -0.683% | 3.07E-04 | -1.978 | 0.452 | 0.138 |
| 2579A>G | [1018-NA] | 0 | 1673 | 0.000% | 2 | 293 | 0.683% | -0.683% | 3.12E-04 | -2.843 | 0.623 | 0.058 |
| 1721T>A | [1018-NA] | 0 | 1078 | 0.000% | 2 | 293 | 0.683% | -0.683% | 4.05E-04 | -1.712 | 0.404 | 0.181 |
| 817T>G | [1018-NA] | 0 | 1014 | 0.000% | 2 | 293 | 0.683% | -0.683% | 4.17E-04 | -1.784 | 0.418 | 0.168 |
| 256A>G | [1018-NA] | 0 | 1171 | 0.000% | 2 | 293 | 0.683% | -0.683% | 4.78E-04 | -2.332 | 0.537 | 0.097 |
| 1379A>G | [1018-NA] | 0 | 1408 | 0.000% | 2 | 293 | 0.683% | -0.683% | 5.75E-04 | -2.958 | 0.671 | 0.052 |
| 3347G>C | [1018-NA] | 0 | 1481 | 0.000% | 2 | 293 | 0.683% | -0.683% | 8.05E-04 | -2.721 | 0.652 | 0.066 |
| 1945C>G | [1018-NA] | 0 | 986 | 0.000% | 2 | 293 | 0.683% | -0.683% | 8.41E-04 | -1.089 | 0.283 | 0.337 |
| 2016T>G | [1018-NA] | 0 | 882 | 0.000% | 2 | 293 | 0.683% | -0.683% | 9.07E-04 | -1.289 | 0.332 | 0.276 |
| 3655C>G | [1018-NA] | 0 | 991 | 0.000% | 2 | 293 | 0.683% | -0.683% | 9.67E-04 | -2.320 | 0.567 | 0.098 |
| 1659T>G | [1018-NA] | 0 | 951 | 0.000% | 2 | 293 | 0.683% | -0.683% | 1.11E-03 | -1.769 | 0.454 | 0.171 |
| 246T>A | [1018-NA] | 0 | 1052 | 0.000% | 2 | 293 | 0.683% | -0.683% | 1.56E-03 | -2.667 | 0.674 | 0.069 |
| 1432T>G | [1018-NA] | 0 | 1043 | 0.000% | 2 | 293 | 0.683% | -0.683% | 1.78E-03 | -2.658 | 0.682 | 0.070 |
| 544A>T | [1018-NA] | 0 | 1408 | 0.000% | 2 | 293 | 0.683% | -0.683% | 1.86E-03 | -3.363 | 0.847 | 0.035 |
| 912G>C | [1018-NA] | 0 | 1130 | 0.000% | 2 | 293 | 0.683% | -0.683% | 1.86E-03 | -2.451 | 0.643 | 0.086 |
| 1670G>C | [1018-NA] | 0 | 785 | 0.000% | 2 | 293 | 0.683% | -0.683% | 2.11E-03 | -1.577 | 0.433 | 0.207 |
| 659T>G | [1018-NA] | 0 | 1277 | 0.000% | 2 | 293 | 0.683% | -0.683% | 2.32E-03 | -3.266 | 0.846 | 0.038 |
| 3958A>G | [1018-NA] | 0 | 939 | 0.000% | 2 | 293 | 0.683% | -0.683% | 2.42E-03 | -2.353 | 0.633 | 0.095 |
| 1785T>G | [1018-NA] | 0 | 758 | 0.000% | 2 | 293 | 0.683% | -0.683% | 2.44E-03 | -1.765 | 0.485 | 0.171 |
| 3017G>A | [1018-NA] | 0 | 1170 | 0.000% | 2 | 293 | 0.683% | -0.683% | 3.41E-03 | -3.178 | 0.863 | 0.042 |
| 3055G>T | [1018-NA] | 0 | 817 | 0.000% | 2 | 293 | 0.683% | -0.683% | 3.48E-03 | -2.414 | 0.670 | 0.089 |
| 1677T>C | [1018-NA] | 0 | 677 | 0.000% | 2 | 293 | 0.683% | -0.683% | 3.72E-03 | -1.717 | 0.494 | 0.180 |
| 540C>A | [1018-NA] | 0 | 994 | 0.000% | 2 | 293 | 0.683% | -0.683% | 4.29E-03 | -3.015 | 0.843 | 0.049 |
| 666A>T | [1018-NA] | 0 | 962 | 0.000% | 2 | 293 | 0.683% | -0.683% | 4.64E-03 | -2.983 | 0.843 | 0.051 |
| 370T>G | [1018-NA] | 0 | 669 | 0.000% | 2 | 293 | 0.683% | -0.683% | 4.67E-03 | -1.927 | 0.564 | 0.146 |
| 769A>C | [1018-NA] | 0 | 726 | 0.000% | 2 | 293 | 0.683% | -0.683% | 4.82E-03 | -2.009 | 0.590 | 0.134 |
| 2016T>A | [1018-NA] | 0 | 716 | 0.000% | 2 | 293 | 0.683% | -0.683% | 4.90E-03 | -1.150 | 0.352 | 0.317 |
| 3979T>G | [1018-NA] | 0 | 1058 | 0.000% | 2 | 293 | 0.683% | -0.683% | 5.01E-03 | -3.078 | 0.879 | 0.046 |

| | | | | | | | | | | | | |
|---|---|---|---|---|---|---|---|---|---|---|---|---|
| 539C>A | [1018-NA] | 0 | 840 | 0.000% | 2 | 293 | 0.683% | -0.683% | 6.65E-03 | -2.847 | 0.845 | 0.058 |
| 3851A>T | [1018-NA] | 0 | 826 | 0.000% | 2 | 293 | 0.683% | -0.683% | 6.72E-03 | -2.830 | 0.842 | 0.059 |
| 2026T>G | [1018-NA] | 0 | 1635 | 0.000% | 2 | 293 | 0.683% | -0.683% | 7.56E-03 | -4.206 | 1.244 | 0.015 |
| 412A>G | [1018-NA] | 0 | 804 | 0.000% | 2 | 293 | 0.683% | -0.683% | 7.57E-03 | -2.803 | 0.849 | 0.061 |
| 3650T>C | [1018-NA] | 0 | 647 | 0.000% | 2 | 293 | 0.683% | -0.683% | 8.18E-03 | -2.181 | 0.679 | 0.113 |
| 1802T>G | [1018-NA] | 0 | 579 | 0.000% | 2 | 293 | 0.683% | -0.683% | 8.52E-03 | -1.561 | 0.500 | 0.210 |
| 705T>A | [1018-NA] | 0 | 747 | 0.000% | 2 | 293 | 0.683% | -0.683% | 8.63E-03 | -2.730 | 0.842 | 0.065 |
| 1656A>G | [1018-NA] | 0 | 585 | 0.000% | 2 | 293 | 0.683% | -0.683% | 8.81E-03 | -1.446 | 0.468 | 0.236 |
| 1830T>A | [1018-NA] | 0 | 582 | 0.000% | 2 | 293 | 0.683% | -0.683% | 9.30E-03 | -1.788 | 0.574 | 0.167 |
| 1732A>G | [1018-NA] | 0 | 579 | 0.000% | 2 | 293 | 0.683% | -0.683% | 9.67E-03 | -1.379 | 0.453 | 0.252 |
| 544A>C | [1018-NA] | 0 | 712 | 0.000% | 2 | 293 | 0.683% | -0.683% | 1.03E-02 | -2.682 | 0.850 | 0.068 |
| 200A>G | [1018-NA] | 0 | 765 | 0.000% | 2 | 293 | 0.683% | -0.683% | 1.17E-02 | -2.754 | 0.890 | 0.064 |
| 906G>C | [1018-NA] | 0 | 560 | 0.000% | 2 | 293 | 0.683% | -0.683% | 1.19E-02 | -1.596 | 0.535 | 0.203 |
| 1203C>G | [1018-NA] | 0 | 1159 | 0.000% | 2 | 293 | 0.683% | -0.683% | 1.30E-02 | -3.862 | 1.244 | 0.021 |
| 3641G>C | [1018-NA] | 0 | 526 | 0.000% | 2 | 293 | 0.683% | -0.683% | 1.43E-02 | -1.974 | 0.668 | 0.139 |
| 353T>G | [1018-NA] | 0 | 620 | 0.000% | 2 | 293 | 0.683% | -0.683% | 1.46E-02 | -2.544 | 0.852 | 0.079 |
| 3655C>A | [1018-NA] | 0 | 637 | 0.000% | 2 | 293 | 0.683% | -0.683% | 1.46E-02 | -2.571 | 0.861 | 0.076 |
| 3742T>G | [1018-NA] | 0 | 535 | 0.000% | 2 | 293 | 0.683% | -0.683% | 1.59E-02 | -1.991 | 0.687 | 0.136 |
| 1412T>A | [1018-NA] | 0 | 1002 | 0.000% | 2 | 293 | 0.683% | -0.683% | 1.62E-02 | -3.716 | 1.244 | 0.024 |
| 2822A>T | [1018-NA] | 0 | 588 | 0.000% | 2 | 293 | 0.683% | -0.683% | 1.83E-02 | -2.491 | 0.866 | 0.083 |
| 912G>A | [1018-NA] | 0 | 568 | 0.000% | 2 | 293 | 0.683% | -0.683% | 1.86E-02 | -2.456 | 0.857 | 0.086 |
| 3016T>A | [1018-NA] | 0 | 460 | 0.000% | 2 | 293 | 0.683% | -0.683% | 2.12E-02 | -1.554 | 0.567 | 0.211 |
| 912G>T | [1018-NA] | 0 | 481 | 0.000% | 2 | 293 | 0.683% | -0.683% | 2.29E-02 | -1.685 | 0.621 | 0.185 |
| 1788T>G | [1018-NA] | 0 | 485 | 0.000% | 2 | 293 | 0.683% | -0.683% | 2.50E-02 | -1.893 | 0.705 | 0.151 |
| 815T>G | [1018-NA] | 0 | 457 | 0.000% | 2 | 293 | 0.683% | -0.683% | 2.51E-02 | -1.834 | 0.684 | 0.160 |
| 3631T>C | [1018-NA] | 0 | 459 | 0.000% | 2 | 293 | 0.683% | -0.683% | 2.97E-02 | -2.243 | 0.853 | 0.106 |
| 3369A>T | [1018-NA] | 0 | 432 | 0.000% | 2 | 293 | 0.683% | -0.683% | 2.98E-02 | -1.411 | 0.549 | 0.244 |
| 3604A>C | [1018-NA] | 0 | 450 | 0.000% | 2 | 293 | 0.683% | -0.683% | 3.10E-02 | -2.224 | 0.851 | 0.108 |
| 1839T>G | [1018-NA] | 0 | 399 | 0.000% | 2 | 293 | 0.683% | -0.683% | 3.40E-02 | -1.699 | 0.670 | 0.183 |
| 907G>A | [1018-NA] | 0 | 423 | 0.000% | 2 | 293 | 0.683% | -0.683% | 3.44E-02 | -2.162 | 0.845 | 0.115 |
| 3916A>C | [1018-NA] | 0 | 583 | 0.000% | 2 | 293 | 0.683% | -0.683% | 3.74E-02 | -3.175 | 1.244 | 0.042 |
| 1378A>G | [1018-NA] | 0 | 581 | 0.000% | 2 | 293 | 0.683% | -0.683% | 3.77E-02 | -3.171 | 1.244 | 0.042 |
| 262T>G | [1018-NA] | 0 | 547 | 0.000% | 2 | 293 | 0.683% | -0.683% | 4.11E-02 | -3.111 | 1.244 | 0.045 |
| 3651T>C | [1018-NA] | 0 | 384 | 0.000% | 2 | 293 | 0.683% | -0.683% | 4.34E-02 | -2.065 | 0.848 | 0.127 |
| 3269A>T | [1018-NA] | 0 | 372 | 0.000% | 2 | 293 | 0.683% | -0.683% | 4.57E-02 | -2.033 | 0.844 | 0.131 |
| 3641G>A | [1018-NA] | 0 | 374 | 0.000% | 2 | 293 | 0.683% | -0.683% | 4.61E-02 | -2.039 | 0.847 | 0.130 |
| 2381C>T | [1018-NA] | 0 | 490 | 0.000% | 2 | 293 | 0.683% | -0.683% | 4.85E-02 | -3.001 | 1.244 | 0.050 |

**Table S4d** - activation of pseudoexons among wts, comparison of library 8 with all other libraries.

| Pseudoexon | # of RNA reads with the PE (wts of library 8 only) | Total RNA reads in wts (library 8 only) | Activation (library 8 only) | # of RNA reads with the PE (all other libraries) | Total RNA reads in wt (all other libraries) | Activation (all other libraries) | Activation difference | p (adj, fdr) | log odds ratio | se log odds ratio | odds ratio |
|---|---|---|---|---|---|---|---|---|---|---|---|
| [1534-NA] | 6 | 2716 | 0.2209% | 1 | 231989 | 0.0004% | 0.2205% | 7.51E-23 | 1.620 | 0.156 | 5.053 |
| [957-949] | 1 | 2716 | 0.0368% | 0 | 231989 | 0.0000% | 0.0368% | 7.86E-08 | 1.101 | 0.188 | 3.008 |
| [1533-NA] | 1 | 2716 | 0.0368% | 0 | 231989 | 0.0000% | 0.0368% | 7.86E-08 | 1.101 | 0.187 | 3.008 |
| [1534-1685] | 1 | 2716 | 0.0368% | 0 | 231989 | 0.0000% | 0.0368% | 7.86E-08 | 1.101 | 0.188 | 3.008 |
| [3769-3845] | 31 | 2716 | 1.1414% | 1766 | 231989 | 0.7612% | 0.3801% | 2.84E-01 | 0.482 | 0.450 | 1.619 |

**Table S4e** - Comparison of pseudoexon activation in variants of the library 8 with the activation in wild types of library 8 only

| Variant | Pseudoexon | # of RNA reads with the PE | Total RNA reads of the variant | Activation | # of RNA reads with the PE (wts of library 8) | Total RNA reads in wt (wts of library 8) | Activation (wts of library 8) | Activation difference | p (adj, fdr) | log odds ratio | se log odds ratio | odds ratio |
|---|---|---|---|---|---|---|---|---|---|---|---|---|
| 1677T>G | [1534-NA] | 117 | 392 | 29.847% | 6 | 2716 | 0.221% | 29.626% | 5.85E-19 | 4.553 | 0.354 | 94.963 |
| 1684T>G;1689T>G | [1534-NA] | 10 | 59 | 16.949% | 6 | 2716 | 0.221% | 16.728% | 1.26E-09 | 3.832 | 0.486 | 46.138 |
| 1677T>G | [1534-1673] | 23 | 392 | 5.867% | 0 | 2716 | 0.000% | 5.867% | 2.35E-20 | 3.379 | 0.247 | 29.335 |
| 1684T>G;1689T>G | [1534-1685] | 2 | 42 | 4.762% | 1 | 2716 | 0.037% | 4.725% | 8.53E-13 | 3.016 | 0.304 | 20.401 |
| 3176G>T;3317G>T | [1534-NA] | 1 | 37 | 2.703% | 6 | 2716 | 0.221% | 2.482% | 1.59E-01 | 1.906 | 1.077 | 6.729 |
| 1677T>G | [1533-NA] | 7 | 290 | 2.414% | 1 | 2716 | 0.037% | 2.377% | 2.00E-16 | 2.410 | 0.207 | 11.138 |
| 1666T>G | [1534-NA] | 31 | 2575 | 1.204% | 6 | 2716 | 0.221% | 0.983% | 6.57E-03 | 1.132 | 0.357 | 3.102 |
| 1799A>G | [1534-NA] | 3 | 328 | 0.915% | 6 | 2716 | 0.221% | 0.694% | 2.20E-01 | 0.972 | 0.622 | 2.643 |
| 1661T>A | [1534-NA] | 2 | 289 | 0.692% | 6 | 2716 | 0.221% | 0.471% | 3.94E-01 | 0.800 | 0.704 | 2.224 |
| 1670G>A | [1534-NA] | 10 | 1639 | 0.610% | 6 | 2716 | 0.221% | 0.389% | 3.10E-01 | 0.572 | 0.432 | 1.773 |
| 1742A>C | [1534-NA] | 1 | 183 | 0.546% | 6 | 2716 | 0.221% | 0.326% | 7.53E-01 | 0.454 | 0.983 | 1.575 |
| 1677T>G | [1533-1662] | 1 | 185 | 0.541% | 0 | 2716 | 0.000% | 0.541% | 1.29E-06 | 1.491 | 0.254 | 4.440 |
| 1637T>C | [1534-NA] | 13 | 2581 | 0.504% | 6 | 2716 | 0.221% | 0.283% | 4.67E-01 | 0.390 | 0.397 | 1.477 |
| 1640C>G;1684T>G | [1534-NA] | 2 | 435 | 0.460% | 6 | 2716 | 0.221% | 0.239% | 7.97E-01 | 0.281 | 0.734 | 1.324 |
| 1677T>C | [1534-NA] | 3 | 677 | 0.443% | 6 | 2716 | 0.221% | 0.222% | 6.31E-01 | 0.386 | 0.567 | 1.471 |
| 1684T>G;1725T>A | [1534-1673] | 1 | 227 | 0.441% | 0 | 2716 | 0.000% | 0.441% | 1.72E-03 | 1.116 | 0.301 | 3.051 |
| 1659T>G | [1534-NA] | 4 | 951 | 0.421% | 6 | 2716 | 0.221% | 0.200% | 6.46E-01 | 0.334 | 0.511 | 1.396 |
| 1725T>A | [1534-NA] | 18 | 4376 | 0.411% | 6 | 2716 | 0.221% | 0.190% | 5.38E-01 | 0.315 | 0.373 | 1.370 |
| 1678A>G | [1534-NA] | 1 | 275 | 0.364% | 6 | 2716 | 0.221% | 0.143% | 8.12E-01 | 0.312 | 0.883 | 1.367 |
| 1830T>A | [1534-NA] | 2 | 582 | 0.344% | 6 | 2716 | 0.221% | 0.123% | 8.39E-01 | 0.195 | 0.643 | 1.216 |
| 1765A>G;1817C>A | [1534-NA] | 1 | 308 | 0.325% | 6 | 2716 | 0.221% | 0.104% | 9.67E-01 | -0.068 | 0.980 | 0.934 |
| 1692G>A | [1534-NA] | 9 | 2979 | 0.302% | 6 | 2716 | 0.221% | 0.081% | 8.49E-01 | 0.107 | 0.374 | 1.113 |
| 1741G>T | [1534-NA] | 4 | 1367 | 0.293% | 6 | 2716 | 0.221% | 0.072% | 8.59E-01 | 0.124 | 0.461 | 1.132 |
| 1732A>G;1733T>G | [1534-NA] | 3 | 1037 | 0.289% | 6 | 2716 | 0.221% | 0.068% | 8.76E-01 | 0.140 | 0.585 | 1.151 |
| 1670G>C | [1534-NA] | 2 | 785 | 0.255% | 6 | 2716 | 0.221% | 0.034% | 8.91E-01 | 0.118 | 0.558 | 1.125 |
| 1839T>G | [1534-1673] | 1 | 399 | 0.251% | 0 | 2716 | 0.000% | 0.251% | 2.06E-02 | 0.722 | 0.261 | 2.058 |
| 1839T>G | [1534-NA] | 1 | 399 | 0.251% | 6 | 2716 | 0.221% | 0.030% | 9.70E-01 | 0.051 | 0.807 | 1.052 |
| 1678A>T | [1534-NA] | 4 | 1637 | 0.244% | 6 | 2716 | 0.221% | 0.023% | 9.99E-01 | -0.001 | 0.429 | 0.999 |
| 1661T>G | [1534-NA] | 3 | 1259 | 0.238% | 6 | 2716 | 0.221% | 0.017% | 8.88E-01 | -0.111 | 0.511 | 0.895 |
| 1788T>G | [1534-NA] | 1 | 485 | 0.206% | 6 | 2716 | 0.221% | -0.015% | 9.09E-01 | -0.145 | 0.813 | 0.865 |
| 1689T>G | [1534-NA] | 4 | 2120 | 0.189% | 6 | 2716 | 0.221% | -0.032% | 7.91E-01 | -0.170 | 0.432 | 0.844 |
| 1684T>G | [1534-NA] | 7 | 3906 | 0.179% | 6 | 2716 | 0.221% | -0.042% | 8.57E-01 | -0.101 | 0.368 | 0.904 |
| 1732A>G | [1534-NA] | 1 | 579 | 0.173% | 6 | 2716 | 0.221% | -0.048% | 9.49E-01 | 0.064 | 0.639 | 1.066 |
| 1656A>G | [1534-NA] | 1 | 585 | 0.171% | 6 | 2716 | 0.221% | -0.050% | 9.88E-01 | 0.018 | 0.664 | 1.018 |
| 209T>G | [1534-NA] | 3 | 1913 | 0.157% | 6 | 2716 | 0.221% | -0.064% | 4.02E-01 | -0.772 | 0.689 | 0.462 |
| 1656A>G;1657C>G | [1534-NA] | 1 | 644 | 0.155% | 6 | 2716 | 0.221% | -0.066% | 7.97E-01 | -0.282 | 0.741 | 0.754 |
| 2532A>C;2551T>G | [1534-NA] | 1 | 655 | 0.153% | 6 | 2716 | 0.221% | -0.068% | 4.94E-01 | -0.991 | 1.063 | 0.371 |
| 1994A>G | [1534-NA] | 1 | 697 | 0.143% | 6 | 2716 | 0.221% | -0.077% | 8.22E-01 | 0.183 | 0.544 | 1.201 |
| 1785T>G | [1534-1673] | 1 | 758 | 0.132% | 0 | 2716 | 0.000% | 0.132% | 2.81E-01 | 0.393 | 0.281 | 1.481 |
| 1637T>C | [1544-1673] | 1 | 787 | 0.127% | 0 | 2716 | 0.000% | 0.127% | 2.68E-01 | 0.275 | 0.192 | 1.317 |
| 1682A>T | [1534-NA] | 5 | 4041 | 0.124% | 6 | 2716 | 0.221% | -0.097% | 5.79E-01 | -0.335 | 0.436 | 0.715 |
| 1725T>A | [1534-1673] | 5 | 4376 | 0.114% | 0 | 2716 | 0.000% | 0.114% | 2.71E-01 | 0.330 | 0.233 | 1.391 |
| 1741G>T | [1533-NA] | 1 | 906 | 0.110% | 1 | 2716 | 0.037% | 0.074% | 6.36E-01 | 0.209 | 0.312 | 1.233 |
| 1817C>A | [1534-NA] | 3 | 2724 | 0.110% | 6 | 2716 | 0.221% | -0.111% | 6.61E-01 | -0.362 | 0.578 | 0.697 |

| | | | | | | | | | | | | |
|---|---|---|---|---|---|---|---|---|---|---|---|---|
| 1735C>G | [1534-NA] | 2 | 1982 | 0.101% | 6 | 2716 | 0.221% | -0.120% | 5.83E-01 | -0.371 | 0.486 | 0.690 |
| 1732A>G;1733T>G | [1534-1673] | 1 | 1037 | 0.096% | 0 | 2716 | 0.000% | 0.096% | 1.83E-01 | 0.347 | 0.207 | 1.415 |
| 1670G>A | [1533-NA] | 1 | 1047 | 0.096% | 1 | 2716 | 0.037% | 0.059% | 7.80E-01 | 0.098 | 0.236 | 1.103 |
| 1678A>T | [1534-1685] | 1 | 1068 | 0.094% | 1 | 2716 | 0.037% | 0.057% | 7.67E-01 | 0.109 | 0.250 | 1.116 |
| 1721T>A | [1534-NA] | 1 | 1078 | 0.093% | 6 | 2716 | 0.221% | -0.128% | 6.93E-01 | -0.363 | 0.640 | 0.696 |
| 1640C>G | [1534-1673] | 1 | 1279 | 0.078% | 0 | 2716 | 0.000% | 0.078% | 3.41E-01 | 0.222 | 0.178 | 1.248 |
| 1640C>G | [1534-NA] | 1 | 1279 | 0.078% | 6 | 2716 | 0.221% | -0.143% | 5.57E-01 | -0.449 | 0.556 | 0.638 |
| 1807A>G | [1534-NA] | 1 | 1292 | 0.077% | 6 | 2716 | 0.221% | -0.144% | 5.00E-01 | -0.544 | 0.592 | 0.581 |
| 1765A>G | [1534-NA] | 2 | 2609 | 0.077% | 6 | 2716 | 0.221% | -0.144% | 3.28E-01 | -0.568 | 0.444 | 0.567 |
| 1725T>A | [1534-3845] | 1 | 1337 | 0.075% | 0 | 2716 | 0.000% | 0.075% | 2.35E-01 | 0.255 | 0.168 | 1.291 |
| 1605G>C | [1534-NA] | 1 | 1377 | 0.073% | 6 | 2716 | 0.221% | -0.148% | 2.73E-01 | -1.088 | 0.766 | 0.337 |
| 1725T>A | [1533-1673] | 1 | 1443 | 0.069% | 0 | 2716 | 0.000% | 0.069% | 3.90E-01 | 0.179 | 0.157 | 1.196 |
| 2570T>G | [1534-NA] | 1 | 1601 | 0.062% | 6 | 2716 | 0.221% | -0.158% | 1.68E-01 | -1.511 | 0.873 | 0.221 |
| 1714T>A | [1534-NA] | 1 | 1607 | 0.062% | 6 | 2716 | 0.221% | -0.159% | 4.01E-01 | -0.575 | 0.514 | 0.563 |
| 1670G>A | [1534-1673] | 1 | 1639 | 0.061% | 0 | 2716 | 0.000% | 0.061% | 7.85E-01 | 0.077 | 0.189 | 1.080 |
| 1297A>G | [1534-NA] | 1 | 1649 | 0.061% | 6 | 2716 | 0.221% | -0.160% | 1.66E-01 | -1.423 | 0.817 | 0.241 |
| #HODNOTA! | [1534-NA] | 2 | 3507 | 0.057% | 6 | 2716 | 0.221% | -0.164% | 8.56E-02 | -1.218 | 0.579 | 0.296 |
| 1735C>G | [1534-1673] | 1 | 1982 | 0.050% | 0 | 2716 | 0.000% | 0.050% | 6.89E-01 | 0.124 | 0.215 | 1.132 |
| 1689T>G | [1534-1673] | 1 | 2120 | 0.047% | 0 | 2716 | 0.000% | 0.047% | 9.03E-01 | 0.037 | 0.196 | 1.038 |
| #HODNOTA! | [1534-NA] | 1 | 2271 | 0.044% | 6 | 2716 | 0.221% | -0.177% | 1.02E-01 | -1.266 | 0.630 | 0.282 |
| 1666T>G | [1534-1662] | 1 | 2575 | 0.039% | 0 | 2716 | 0.000% | 0.039% | 9.40E-01 | -0.029 | 0.237 | 0.972 |
| 1666T>G | [1534-1673] | 1 | 2575 | 0.039% | 0 | 2716 | 0.000% | 0.039% | 9.21E-01 | -0.029 | 0.182 | 0.972 |
| 1765A>G | [1534-1662] | 1 | 2609 | 0.038% | 0 | 2716 | 0.000% | 0.038% | 8.44E-01 | -0.059 | 0.200 | 0.943 |
| 1765A>G | [1534-1673] | 1 | 2609 | 0.038% | 0 | 2716 | 0.000% | 0.038% | 8.35E-01 | -0.059 | 0.189 | 0.943 |
| 3293A>T | [1534-NA] | 1 | 2622 | 0.038% | 6 | 2716 | 0.221% | -0.183% | 6.98E-02 | -1.343 | 0.608 | 0.261 |
| 1817C>A | [1534-1662] | 1 | 2724 | 0.037% | 0 | 2716 | 0.000% | 0.037% | 9.12E-01 | 0.044 | 0.254 | 1.045 |
| 1817C>A | [1534-1673] | 1 | 2724 | 0.037% | 0 | 2716 | 0.000% | 0.037% | 8.42E-01 | 0.044 | 0.148 | 1.045 |
| 1692G>A | [1534-1673] | 1 | 2979 | 0.034% | 0 | 2716 | 0.000% | 0.034% | 9.57E-01 | 0.014 | 0.156 | 1.014 |
| 3507T>A | [1534-NA] | 1 | 3609 | 0.028% | 6 | 2716 | 0.221% | -0.193% | 4.72E-02 | -1.072 | 0.451 | 0.342 |
| 1684T>G | [1534-1673] | 1 | 3906 | 0.026% | 0 | 2716 | 0.000% | 0.026% | 7.38E-01 | 0.099 | 0.204 | 1.104 |
| 1682A>T | [1534-1673] | 1 | 4041 | 0.025% | 0 | 2716 | 0.000% | 0.025% | 9.27E-01 | -0.029 | 0.199 | 0.971 |
| 3317G>T | [1534-NA] | 1 | 7837 | 0.013% | 6 | 2716 | 0.221% | -0.208% | 2.22E-04 | -1.745 | 0.409 | 0.175 |
| 3037A>G | [1534-NA] | 1 | 13185 | 0.008% | 6 | 2716 | 0.221% | -0.213% | 7.52E-07 | -1.701 | 0.301 | 0.182 |
| 1637T>C | [1533-1662] | 0 | 860 | 0.000% | 0 | 2716 | 0.000% | 0.000% | 1.69E-01 | -0.353 | 0.205 | 0.702 |
| 1640C>G | [1533-1662] | 0 | 452 | 0.000% | 0 | 2716 | 0.000% | 0.000% | 6.64E-01 | -0.152 | 0.246 | 0.859 |
| 1640C>G;1684T>G | [1533-1662] | 0 | 171 | 0.000% | 0 | 2716 | 0.000% | 0.000% | 3.02E-01 | -0.684 | 0.507 | 0.505 |
| 1656A>G | [1533-1662] | 0 | 201 | 0.000% | 0 | 2716 | 0.000% | 0.000% | 5.37E-01 | 0.252 | 0.297 | 1.286 |
| 1656A>G;1657C>G | [1533-1662] | 0 | 250 | 0.000% | 0 | 2716 | 0.000% | 0.000% | 4.31E-01 | -0.371 | 0.350 | 0.690 |
| 1659T>G | [1533-1662] | 0 | 316 | 0.000% | 0 | 2716 | 0.000% | 0.000% | 9.19E-01 | -0.046 | 0.287 | 0.955 |
| 1659T>G;1785T>G | [1533-1662] | 0 | 19 | 0.000% | 0 | 2716 | 0.000% | 0.000% | 3.77E-01 | 0.816 | 0.699 | 2.262 |
| 1661T>A | [1533-1662] | 0 | 93 | 0.000% | 0 | 2716 | 0.000% | 0.000% | 5.56E-01 | 0.329 | 0.404 | 1.389 |
| 1661T>A;1725T>A | [1533-1662] | 0 | 109 | 0.000% | 0 | 2716 | 0.000% | 0.000% | 3.10E-01 | -0.926 | 0.697 | 0.396 |
| 1661T>G | [1533-1662] | 0 | 439 | 0.000% | 0 | 2716 | 0.000% | 0.000% | 3.02E-01 | -0.374 | 0.278 | 0.688 |
| 1665A>G | [1533-1662] | 0 | 552 | 0.000% | 0 | 2716 | 0.000% | 0.000% | 6.47E-01 | -0.152 | 0.233 | 0.859 |
| 1666T>G | [1533-1662] | 0 | 871 | 0.000% | 0 | 2716 | 0.000% | 0.000% | 3.43E-01 | -0.233 | 0.187 | 0.792 |
| 1670G>A | [1533-1662] | 0 | 532 | 0.000% | 0 | 2716 | 0.000% | 0.000% | 7.28E-01 | -0.115 | 0.227 | 0.892 |
| 1670G>C | [1533-1662] | 0 | 294 | 0.000% | 0 | 2716 | 0.000% | 0.000% | 9.53E-01 | 0.026 | 0.280 | 1.026 |
| 1677T>C | [1533-1662] | 0 | 232 | 0.000% | 0 | 2716 | 0.000% | 0.000% | 8.80E-01 | -0.073 | 0.314 | 0.929 |
| 1678A>G | [1533-1662] | 0 | 98 | 0.000% | 0 | 2716 | 0.000% | 0.000% | 8.65E-01 | -0.128 | 0.496 | 0.880 |

| | | | | | | | | | | | | |
|---|---|---|---|---|---|---|---|---|---|---|---|---|
| 1678A>T | [1533-1662] | 0 | 538 | 0.000% | 0 | 2716 | 0.000% | 0.000% | 6.97E-01 | -0.126 | 0.226 | 0.882 |
| 1682A>T | [1533-1662] | 0 | 1401 | 0.000% | 0 | 2716 | 0.000% | 0.000% | 4.10E-01 | -0.185 | 0.168 | 0.831 |
| 1684T>G | [1533-1662] | 0 | 1371 | 0.000% | 0 | 2716 | 0.000% | 0.000% | 8.01E-01 | -0.058 | 0.156 | 0.943 |
| 1684T>G;1689T>G | [1533-1662] | 0 | 22 | 0.000% | 0 | 2716 | 0.000% | 0.000% | 4.80E-01 | 0.671 | 0.698 | 1.955 |
| 1684T>G;1725T>A | [1533-1662] | 0 | 78 | 0.000% | 0 | 2716 | 0.000% | 0.000% | 8.97E-01 | 0.100 | 0.495 | 1.105 |
| 1689T>G | [1533-1662] | 0 | 700 | 0.000% | 0 | 2716 | 0.000% | 0.000% | 6.18E-01 | -0.148 | 0.210 | 0.862 |
| 1692G>A | [1533-1662] | 0 | 1054 | 0.000% | 0 | 2716 | 0.000% | 0.000% | 4.00E-01 | -0.200 | 0.179 | 0.818 |
| 1692G>A;1695G>A | [1533-1662] | 0 | 89 | 0.000% | 0 | 2716 | 0.000% | 0.000% | 4.42E-01 | -0.724 | 0.697 | 0.485 |
| 1692G>A;1807A>G | [1533-1662] | 0 | 75 | 0.000% | 0 | 2716 | 0.000% | 0.000% | 5.68E-01 | -0.553 | 0.697 | 0.575 |
| 1714T>A | [1533-1662] | 0 | 588 | 0.000% | 0 | 2716 | 0.000% | 0.000% | 7.21E-01 | -0.128 | 0.246 | 0.880 |
| 1721T>A | [1533-1662] | 0 | 381 | 0.000% | 0 | 2716 | 0.000% | 0.000% | 8.09E-01 | -0.099 | 0.276 | 0.905 |
| 1721T>A;1817C>A | [1533-1662] | 0 | 90 | 0.000% | 0 | 2716 | 0.000% | 0.000% | 4.34E-01 | -0.735 | 0.697 | 0.480 |
| 1725T>A | [1533-1662] | 0 | 1443 | 0.000% | 0 | 2716 | 0.000% | 0.000% | 5.88E-01 | -0.109 | 0.145 | 0.896 |
| 1732A>G | [1533-1662] | 0 | 189 | 0.000% | 0 | 2716 | 0.000% | 0.000% | 4.20E-01 | 0.313 | 0.290 | 1.367 |
| 1732A>G;1733T>G | [1533-1662] | 0 | 378 | 0.000% | 0 | 2716 | 0.000% | 0.000% | 8.13E-01 | -0.091 | 0.261 | 0.913 |
| 1735C>G | [1533-1662] | 0 | 675 | 0.000% | 0 | 2716 | 0.000% | 0.000% | 7.12E-01 | -0.112 | 0.209 | 0.894 |
| 1741G>T | [1533-1662] | 0 | 489 | 0.000% | 0 | 2716 | 0.000% | 0.000% | 7.07E-01 | -0.126 | 0.232 | 0.882 |
| 1742A>C | [1533-1662] | 0 | 61 | 0.000% | 0 | 2716 | 0.000% | 0.000% | 7.33E-01 | -0.346 | 0.697 | 0.707 |
| 1745A>G | [1533-1662] | 0 | 121 | 0.000% | 0 | 2716 | 0.000% | 0.000% | 9.19E-01 | 0.066 | 0.413 | 1.069 |
| 1747A>C | [1533-1662] | 0 | 462 | 0.000% | 0 | 2716 | 0.000% | 0.000% | 2.40E-01 | -0.425 | 0.282 | 0.654 |
| 1747A>C;1765A>G | [1533-1662] | 0 | 29 | 0.000% | 0 | 2716 | 0.000% | 0.000% | 6.93E-01 | 0.395 | 0.698 | 1.485 |
| 1765A>G | [1533-1662] | 0 | 917 | 0.000% | 0 | 2716 | 0.000% | 0.000% | 1.75E-01 | -0.349 | 0.205 | 0.706 |
| 1765A>G;1817C>A | [1533-1662] | 0 | 113 | 0.000% | 0 | 2716 | 0.000% | 0.000% | 2.89E-01 | -0.962 | 0.697 | 0.382 |
| 1785T>G | [1533-1662] | 0 | 234 | 0.000% | 0 | 2716 | 0.000% | 0.000% | 8.23E-01 | 0.100 | 0.301 | 1.105 |
| 1788T>G | [1533-1662] | 0 | 147 | 0.000% | 0 | 2716 | 0.000% | 0.000% | 8.37E-01 | -0.128 | 0.416 | 0.880 |
| 1799A>G | [1533-1662] | 0 | 127 | 0.000% | 0 | 2716 | 0.000% | 0.000% | 9.79E-01 | 0.018 | 0.405 | 1.018 |
| 1802T>G | [1533-1662] | 0 | 185 | 0.000% | 0 | 2716 | 0.000% | 0.000% | 7.43E-01 | 0.152 | 0.318 | 1.165 |
| 1807A>G | [1533-1662] | 0 | 415 | 0.000% | 0 | 2716 | 0.000% | 0.000% | 6.20E-01 | -0.185 | 0.264 | 0.831 |
| 1817C>A | [1533-1662] | 0 | 965 | 0.000% | 0 | 2716 | 0.000% | 0.000% | 6.61E-01 | -0.112 | 0.180 | 0.894 |
| 1826C>G | [1533-1662] | 0 | 28 | 0.000% | 0 | 2716 | 0.000% | 0.000% | 6.64E-01 | 0.430 | 0.698 | 1.538 |
| 1826C>G;1827T>A | [1533-1662] | 0 | 68 | 0.000% | 0 | 2716 | 0.000% | 0.000% | 7.45E-01 | 0.237 | 0.501 | 1.267 |
| 1830T>A | [1533-1662] | 0 | 196 | 0.000% | 0 | 2716 | 0.000% | 0.000% | 8.10E-01 | -0.128 | 0.356 | 0.880 |
| 1834A>G | [1533-1662] | 0 | 89 | 0.000% | 0 | 2716 | 0.000% | 0.000% | 9.70E-01 | -0.032 | 0.493 | 0.969 |
| 1839T>G | [1533-1662] | 0 | 130 | 0.000% | 0 | 2716 | 0.000% | 0.000% | 9.95E-01 | -0.005 | 0.408 | 0.995 |
| 1637T>C | [1533-1673] | 0 | 860 | 0.000% | 0 | 2716 | 0.000% | 0.000% | 1.69E-01 | -0.353 | 0.205 | 0.702 |
| 1640C>G | [1533-1673] | 0 | 452 | 0.000% | 0 | 2716 | 0.000% | 0.000% | 6.64E-01 | -0.152 | 0.246 | 0.859 |
| 1640C>G;1684T>G | [1533-1673] | 0 | 171 | 0.000% | 0 | 2716 | 0.000% | 0.000% | 3.02E-01 | -0.684 | 0.507 | 0.505 |
| 1656A>G | [1533-1673] | 0 | 201 | 0.000% | 0 | 2716 | 0.000% | 0.000% | 5.37E-01 | 0.252 | 0.297 | 1.286 |
| 1656A>G;1657C>G | [1533-1673] | 0 | 250 | 0.000% | 0 | 2716 | 0.000% | 0.000% | 4.31E-01 | -0.371 | 0.350 | 0.690 |
| 1659T>G | [1533-1673] | 0 | 316 | 0.000% | 0 | 2716 | 0.000% | 0.000% | 9.19E-01 | -0.046 | 0.287 | 0.955 |
| 1659T>G;1785T>G | [1533-1673] | 0 | 19 | 0.000% | 0 | 2716 | 0.000% | 0.000% | 3.77E-01 | 0.816 | 0.699 | 2.262 |
| 1661T>A | [1533-1673] | 0 | 93 | 0.000% | 0 | 2716 | 0.000% | 0.000% | 5.56E-01 | 0.329 | 0.404 | 1.389 |
| 1661T>A;1725T>A | [1533-1673] | 0 | 109 | 0.000% | 0 | 2716 | 0.000% | 0.000% | 3.10E-01 | -0.926 | 0.697 | 0.396 |
| 1661T>G | [1533-1673] | 0 | 439 | 0.000% | 0 | 2716 | 0.000% | 0.000% | 3.02E-01 | -0.374 | 0.278 | 0.688 |
| 1665A>G | [1533-1673] | 0 | 552 | 0.000% | 0 | 2716 | 0.000% | 0.000% | 6.47E-01 | -0.152 | 0.233 | 0.859 |
| 1666T>G | [1533-1673] | 0 | 871 | 0.000% | 0 | 2716 | 0.000% | 0.000% | 3.43E-01 | -0.233 | 0.187 | 0.792 |
| 1670G>A | [1533-1673] | 0 | 532 | 0.000% | 0 | 2716 | 0.000% | 0.000% | 7.28E-01 | -0.115 | 0.227 | 0.892 |
| 1670G>C | [1533-1673] | 0 | 294 | 0.000% | 0 | 2716 | 0.000% | 0.000% | 9.53E-01 | 0.026 | 0.280 | 1.026 |
| 1677T>C | [1533-1673] | 0 | 232 | 0.000% | 0 | 2716 | 0.000% | 0.000% | 8.80E-01 | -0.073 | 0.314 | 0.929 |

| | | | | | | | | | | | | |
|---|---|---|---|---|---|---|---|---|---|---|---|---|
| 1677T>G | [1533-1673] | 0 | 185 | 0.000% | 0 | 2716 | 0.000% | 0.000% | 1.03E-02 | 0.738 | 0.242 | 2.092 |
| 1678A>G | [1533-1673] | 0 | 98 | 0.000% | 0 | 2716 | 0.000% | 0.000% | 8.65E-01 | -0.128 | 0.496 | 0.880 |
| 1678A>T | [1533-1673] | 0 | 538 | 0.000% | 0 | 2716 | 0.000% | 0.000% | 6.97E-01 | -0.126 | 0.226 | 0.882 |
| 1682A>T | [1533-1673] | 0 | 1401 | 0.000% | 0 | 2716 | 0.000% | 0.000% | 4.10E-01 | -0.185 | 0.168 | 0.831 |
| 1684T>G | [1533-1673] | 0 | 1371 | 0.000% | 0 | 2716 | 0.000% | 0.000% | 8.01E-01 | -0.058 | 0.156 | 0.943 |
| 1684T>G;1689T>G | [1533-1673] | 0 | 22 | 0.000% | 0 | 2716 | 0.000% | 0.000% | 4.80E-01 | 0.671 | 0.698 | 1.955 |
| 1684T>G;1725T>A | [1533-1673] | 0 | 78 | 0.000% | 0 | 2716 | 0.000% | 0.000% | 8.97E-01 | 0.100 | 0.495 | 1.105 |
| 1689T>G | [1533-1673] | 0 | 700 | 0.000% | 0 | 2716 | 0.000% | 0.000% | 6.18E-01 | -0.148 | 0.210 | 0.862 |
| 1692G>A | [1533-1673] | 0 | 1054 | 0.000% | 0 | 2716 | 0.000% | 0.000% | 4.00E-01 | -0.200 | 0.179 | 0.818 |
| 1692G>A;1695G>A | [1533-1673] | 0 | 89 | 0.000% | 0 | 2716 | 0.000% | 0.000% | 4.42E-01 | -0.724 | 0.697 | 0.485 |
| 1692G>A;1807A>G | [1533-1673] | 0 | 75 | 0.000% | 0 | 2716 | 0.000% | 0.000% | 5.68E-01 | -0.553 | 0.697 | 0.575 |
| 1714T>A | [1533-1673] | 0 | 588 | 0.000% | 0 | 2716 | 0.000% | 0.000% | 7.21E-01 | -0.128 | 0.246 | 0.880 |
| 1721T>A | [1533-1673] | 0 | 381 | 0.000% | 0 | 2716 | 0.000% | 0.000% | 8.09E-01 | -0.099 | 0.276 | 0.905 |
| 1721T>A;1817C>A | [1533-1673] | 0 | 90 | 0.000% | 0 | 2716 | 0.000% | 0.000% | 4.34E-01 | -0.735 | 0.697 | 0.480 |
| 1732A>G | [1533-1673] | 0 | 189 | 0.000% | 0 | 2716 | 0.000% | 0.000% | 4.20E-01 | 0.313 | 0.290 | 1.367 |
| 1732A>G;1733T>G | [1533-1673] | 0 | 378 | 0.000% | 0 | 2716 | 0.000% | 0.000% | 8.13E-01 | -0.091 | 0.261 | 0.913 |
| 1735C>G | [1533-1673] | 0 | 675 | 0.000% | 0 | 2716 | 0.000% | 0.000% | 7.12E-01 | -0.112 | 0.209 | 0.894 |
| 1741G>T | [1533-1673] | 0 | 489 | 0.000% | 0 | 2716 | 0.000% | 0.000% | 7.07E-01 | -0.126 | 0.232 | 0.882 |
| 1742A>C | [1533-1673] | 0 | 61 | 0.000% | 0 | 2716 | 0.000% | 0.000% | 7.33E-01 | -0.346 | 0.697 | 0.707 |
| 1745A>G | [1533-1673] | 0 | 121 | 0.000% | 0 | 2716 | 0.000% | 0.000% | 9.19E-01 | 0.066 | 0.413 | 1.069 |
| 1747A>C | [1533-1673] | 0 | 462 | 0.000% | 0 | 2716 | 0.000% | 0.000% | 2.40E-01 | -0.425 | 0.282 | 0.654 |
| 1747A>C;1765A>G | [1533-1673] | 0 | 29 | 0.000% | 0 | 2716 | 0.000% | 0.000% | 6.93E-01 | 0.395 | 0.698 | 1.485 |
| 1765A>G | [1533-1673] | 0 | 917 | 0.000% | 0 | 2716 | 0.000% | 0.000% | 1.75E-01 | -0.349 | 0.205 | 0.706 |
| 1765A>G;1817C>A | [1533-1673] | 0 | 113 | 0.000% | 0 | 2716 | 0.000% | 0.000% | 2.89E-01 | -0.962 | 0.697 | 0.382 |
| 1785T>G | [1533-1673] | 0 | 234 | 0.000% | 0 | 2716 | 0.000% | 0.000% | 8.23E-01 | 0.100 | 0.301 | 1.105 |
| 1788T>G | [1533-1673] | 0 | 147 | 0.000% | 0 | 2716 | 0.000% | 0.000% | 8.37E-01 | -0.128 | 0.416 | 0.880 |
| 1799A>G | [1533-1673] | 0 | 127 | 0.000% | 0 | 2716 | 0.000% | 0.000% | 9.79E-01 | 0.018 | 0.405 | 1.018 |
| 1802T>G | [1533-1673] | 0 | 185 | 0.000% | 0 | 2716 | 0.000% | 0.000% | 7.43E-01 | 0.152 | 0.318 | 1.165 |
| 1807A>G | [1533-1673] | 0 | 415 | 0.000% | 0 | 2716 | 0.000% | 0.000% | 6.20E-01 | -0.185 | 0.264 | 0.831 |
| 1817C>A | [1533-1673] | 0 | 965 | 0.000% | 0 | 2716 | 0.000% | 0.000% | 6.61E-01 | -0.112 | 0.180 | 0.894 |
| 1826C>G | [1533-1673] | 0 | 28 | 0.000% | 0 | 2716 | 0.000% | 0.000% | 6.64E-01 | 0.430 | 0.698 | 1.538 |
| 1826C>G;1827T>A | [1533-1673] | 0 | 68 | 0.000% | 0 | 2716 | 0.000% | 0.000% | 7.45E-01 | 0.237 | 0.501 | 1.267 |
| 1830T>A | [1533-1673] | 0 | 196 | 0.000% | 0 | 2716 | 0.000% | 0.000% | 8.10E-01 | -0.128 | 0.356 | 0.880 |
| 1834A>G | [1533-1673] | 0 | 89 | 0.000% | 0 | 2716 | 0.000% | 0.000% | 9.70E-01 | -0.032 | 0.493 | 0.969 |
| 1839T>G | [1533-1673] | 0 | 130 | 0.000% | 0 | 2716 | 0.000% | 0.000% | 9.95E-01 | -0.005 | 0.408 | 0.995 |
| 1637T>C | [1533-NA] | 0 | 1647 | 0.000% | 1 | 2716 | 0.037% | -0.037% | 3.71E-02 | -0.458 | 0.184 | 0.633 |
| 1640C>G | [1533-NA] | 0 | 850 | 0.000% | 1 | 2716 | 0.037% | -0.037% | 4.10E-01 | -0.239 | 0.216 | 0.788 |
| 1640C>G;1684T>G | [1533-NA] | 0 | 292 | 0.000% | 1 | 2716 | 0.037% | -0.037% | 2.36E-01 | -0.673 | 0.443 | 0.510 |
| 1656A>G | [1533-NA] | 0 | 401 | 0.000% | 1 | 2716 | 0.037% | -0.037% | 9.66E-01 | 0.020 | 0.270 | 1.020 |
| 1656A>G;1657C>G | [1533-NA] | 0 | 445 | 0.000% | 1 | 2716 | 0.037% | -0.037% | 3.27E-01 | -0.402 | 0.312 | 0.669 |
| 1659T>G | [1533-NA] | 0 | 606 | 0.000% | 1 | 2716 | 0.037% | -0.037% | 5.29E-01 | -0.226 | 0.262 | 0.798 |
| 1659T>G;1785T>G | [1533-NA] | 0 | 48 | 0.000% | 1 | 2716 | 0.037% | -0.037% | 6.16E-01 | 0.436 | 0.616 | 1.547 |
| 1661T>A | [1533-NA] | 0 | 185 | 0.000% | 1 | 2716 | 0.037% | -0.037% | 7.19E-01 | 0.187 | 0.357 | 1.205 |
| 1661T>A;1725T>A | [1533-NA] | 0 | 220 | 0.000% | 1 | 2716 | 0.037% | -0.037% | 1.61E-01 | -1.083 | 0.615 | 0.339 |
| 1661T>G | [1533-NA] | 0 | 792 | 0.000% | 1 | 2716 | 0.037% | -0.037% | 1.79E-01 | -0.419 | 0.248 | 0.658 |
| 1665A>G | [1533-NA] | 0 | 1042 | 0.000% | 1 | 2716 | 0.037% | -0.037% | 3.88E-01 | -0.242 | 0.211 | 0.785 |
| 1666T>G | [1533-NA] | 0 | 1657 | 0.000% | 1 | 2716 | 0.037% | -0.037% | 1.01E-01 | -0.331 | 0.164 | 0.719 |
| 1670G>C | [1533-NA] | 0 | 539 | 0.000% | 1 | 2716 | 0.037% | -0.037% | 7.69E-01 | -0.109 | 0.251 | 0.897 |
| 1677T>C | [1533-NA] | 0 | 445 | 0.000% | 1 | 2716 | 0.037% | -0.037% | 6.46E-01 | -0.179 | 0.275 | 0.836 |

| | | | | | | | | | | | | |
|---|---|---|---|---|---|---|---|---|---|---|---|---|
| 1678A>G | [1533-NA] | 0 | 185 | 0.000% | 1 | 2716 | 0.037% | -0.037% | 9.95E-01 | 0.005 | 0.403 | 1.005 |
| 1678A>T | [1533-NA] | 0 | 1068 | 0.000% | 1 | 2716 | 0.037% | -0.037% | 3.06E-01 | -0.266 | 0.199 | 0.766 |
| 1682A>T | [1533-NA] | 0 | 2670 | 0.000% | 1 | 2716 | 0.037% | -0.037% | 1.31E-01 | -0.284 | 0.152 | 0.752 |
| 1684T>G | [1533-NA] | 0 | 2541 | 0.000% | 1 | 2716 | 0.037% | -0.037% | 4.96E-01 | -0.130 | 0.141 | 0.878 |
| 1684T>G;1689T>G | [1533-NA] | 0 | 42 | 0.000% | 1 | 2716 | 0.037% | -0.037% | 4.97E-01 | 0.569 | 0.616 | 1.767 |
| 1684T>G;1725T>A | [1533-NA] | 0 | 144 | 0.000% | 1 | 2716 | 0.037% | -0.037% | 9.65E-01 | 0.032 | 0.435 | 1.033 |
| 1689T>G | [1533-NA] | 0 | 1352 | 0.000% | 1 | 2716 | 0.037% | -0.037% | 2.14E-01 | -0.297 | 0.189 | 0.743 |
| 1692G>A | [1533-NA] | 0 | 1916 | 0.000% | 1 | 2716 | 0.037% | -0.037% | 2.18E-01 | -0.253 | 0.162 | 0.777 |
| 1692G>A;1695G>A | [1533-NA] | 0 | 165 | 0.000% | 1 | 2716 | 0.037% | -0.037% | 3.24E-01 | -0.796 | 0.615 | 0.451 |
| 1692G>A;1807A>G | [1533-NA] | 0 | 147 | 0.000% | 1 | 2716 | 0.037% | -0.037% | 4.09E-01 | -0.680 | 0.615 | 0.506 |
| 1714T>A | [1533-NA] | 0 | 1071 | 0.000% | 1 | 2716 | 0.037% | -0.037% | 2.77E-01 | -0.315 | 0.224 | 0.730 |
| 1721T>A | [1533-NA] | 0 | 716 | 0.000% | 1 | 2716 | 0.037% | -0.037% | 5.84E-01 | -0.185 | 0.242 | 0.831 |
| 1721T>A;1817C>A | [1533-NA] | 0 | 144 | 0.000% | 1 | 2716 | 0.037% | -0.037% | 4.25E-01 | -0.660 | 0.616 | 0.517 |
| 1725T>A | [1533-NA] | 0 | 2780 | 0.000% | 1 | 2716 | 0.037% | -0.037% | 1.77E-01 | -0.220 | 0.130 | 0.803 |
| 1732A>G | [1533-NA] | 0 | 365 | 0.000% | 1 | 2716 | 0.037% | -0.037% | 5.76E-01 | 0.200 | 0.258 | 1.222 |
| 1732A>G;1733T>G | [1533-NA] | 0 | 689 | 0.000% | 1 | 2716 | 0.037% | -0.037% | 6.61E-01 | -0.147 | 0.234 | 0.864 |
| 1733T>G | [1533-NA] | 0 | 13 | 0.000% | 1 | 2716 | 0.037% | -0.037% | 3.65E-01 | 1.046 | 0.873 | 2.846 |
| 1735C>G | [1533-NA] | 0 | 1293 | 0.000% | 1 | 2716 | 0.037% | -0.037% | 3.65E-01 | -0.216 | 0.181 | 0.806 |
| 1742A>C | [1533-NA] | 0 | 120 | 0.000% | 1 | 2716 | 0.037% | -0.037% | 5.76E-01 | -0.478 | 0.615 | 0.620 |
| 1745A>G | [1533-NA] | 0 | 250 | 0.000% | 1 | 2716 | 0.037% | -0.037% | 8.37E-01 | -0.114 | 0.371 | 0.893 |
| 1747A>C | [1533-NA] | 0 | 886 | 0.000% | 1 | 2716 | 0.037% | -0.037% | 8.36E-02 | -0.531 | 0.250 | 0.588 |
| 1747A>C;1765A>G | [1533-NA] | 0 | 78 | 0.000% | 1 | 2716 | 0.037% | -0.037% | 9.64E-01 | -0.048 | 0.616 | 0.953 |
| 1765A>G | [1533-NA] | 0 | 1727 | 0.000% | 1 | 2716 | 0.037% | -0.037% | 6.89E-02 | -0.404 | 0.183 | 0.668 |
| 1765A>G;1817C>A | [1533-NA] | 0 | 216 | 0.000% | 1 | 2716 | 0.037% | -0.037% | 1.68E-01 | -1.065 | 0.615 | 0.345 |
| 1785T>G | [1533-NA] | 0 | 482 | 0.000% | 1 | 2716 | 0.037% | -0.037% | 6.72E-01 | -0.164 | 0.272 | 0.849 |
| 1788T>G | [1533-NA] | 0 | 303 | 0.000% | 1 | 2716 | 0.037% | -0.037% | 5.55E-01 | -0.306 | 0.375 | 0.737 |
| 1799A>G | [1533-NA] | 0 | 227 | 0.000% | 1 | 2716 | 0.037% | -0.037% | 9.76E-01 | -0.017 | 0.355 | 0.983 |
| 1802T>G | [1533-NA] | 0 | 390 | 0.000% | 1 | 2716 | 0.037% | -0.037% | 9.13E-01 | -0.048 | 0.280 | 0.953 |
| 1807A>G | [1533-NA] | 0 | 841 | 0.000% | 1 | 2716 | 0.037% | -0.037% | 2.49E-01 | -0.345 | 0.234 | 0.708 |
| 1817C>A | [1533-NA] | 0 | 1755 | 0.000% | 1 | 2716 | 0.037% | -0.037% | 3.03E-01 | -0.216 | 0.161 | 0.805 |
| 1826C>G | [1533-NA] | 0 | 58 | 0.000% | 1 | 2716 | 0.037% | -0.037% | 7.87E-01 | 0.248 | 0.615 | 1.281 |
| 1826C>G;1827T>A | [1533-NA] | 0 | 117 | 0.000% | 1 | 2716 | 0.037% | -0.037% | 7.12E-01 | 0.239 | 0.447 | 1.270 |
| 1830T>A | [1533-NA] | 0 | 367 | 0.000% | 1 | 2716 | 0.037% | -0.037% | 6.39E-01 | -0.210 | 0.315 | 0.811 |
| 1834A>G | [1533-NA] | 0 | 158 | 0.000% | 1 | 2716 | 0.037% | -0.037% | 9.30E-01 | -0.060 | 0.434 | 0.941 |
| 1839T>G | [1533-NA] | 0 | 275 | 0.000% | 1 | 2716 | 0.037% | -0.037% | 6.84E-01 | -0.209 | 0.358 | 0.812 |
| 1637T>C | [1534-1662] | 0 | 2581 | 0.000% | 0 | 2716 | 0.000% | 0.000% | 3.84E-02 | -0.354 | 0.143 | 0.702 |
| 1640C>G | [1534-1662] | 0 | 1279 | 0.000% | 0 | 2716 | 0.000% | 0.000% | 6.64E-01 | -0.094 | 0.152 | 0.910 |
| 1640C>G;1684T>G | [1534-1662] | 0 | 435 | 0.000% | 0 | 2716 | 0.000% | 0.000% | 1.61E-01 | -0.519 | 0.295 | 0.595 |
| 1656A>G | [1534-1662] | 0 | 585 | 0.000% | 0 | 2716 | 0.000% | 0.000% | 3.49E-01 | 0.225 | 0.183 | 1.252 |
| 1656A>G;1657C>G | [1534-1662] | 0 | 644 | 0.000% | 0 | 2716 | 0.000% | 0.000% | 4.41E-01 | -0.219 | 0.210 | 0.804 |
| 1659T>G | [1534-1662] | 0 | 951 | 0.000% | 0 | 2716 | 0.000% | 0.000% | 7.29E-01 | -0.098 | 0.194 | 0.907 |
| 1659T>G;1785T>G | [1534-1662] | 0 | 72 | 0.000% | 0 | 2716 | 0.000% | 0.000% | 2.64E-01 | 0.584 | 0.404 | 1.793 |
| 1661T>A | [1534-1662] | 0 | 289 | 0.000% | 0 | 2716 | 0.000% | 0.000% | 3.43E-01 | 0.294 | 0.236 | 1.341 |
| 1661T>A;1725T>A | [1534-1662] | 0 | 345 | 0.000% | 0 | 2716 | 0.000% | 0.000% | 4.41E-02 | -0.980 | 0.402 | 0.375 |
| 1661T>G | [1534-1662] | 0 | 1259 | 0.000% | 0 | 2716 | 0.000% | 0.000% | 1.39E-01 | -0.329 | 0.179 | 0.720 |
| 1665A>G | [1534-1662] | 0 | 1549 | 0.000% | 0 | 2716 | 0.000% | 0.000% | 7.19E-01 | -0.085 | 0.163 | 0.919 |
| 1670G>A | [1534-1662] | 0 | 1639 | 0.000% | 0 | 2716 | 0.000% | 0.000% | 2.66E-01 | -0.204 | 0.142 | 0.816 |
| 1670G>C | [1534-1662] | 0 | 785 | 0.000% | 0 | 2716 | 0.000% | 0.000% | 7.15E-01 | 0.094 | 0.177 | 1.098 |
| 1677T>C | [1534-1662] | 0 | 677 | 0.000% | 0 | 2716 | 0.000% | 0.000% | 8.71E-01 | -0.046 | 0.185 | 0.955 |

| | | | | | | | | | | | | |
|---|---|---|---|---|---|---|---|---|---|---|---|---|
| 1677T>G | [1534-1662] | 0 | 392 | 0.000% | 0 | 2716 | 0.000% | 0.000% | 6.92E-06 | 0.880 | 0.166 | 2.412 |
| 1678A>G | [1534-1662] | 0 | 275 | 0.000% | 0 | 2716 | 0.000% | 0.000% | 8.25E-01 | 0.093 | 0.283 | 1.097 |
| 1678A>T | [1534-1662] | 0 | 1637 | 0.000% | 0 | 2716 | 0.000% | 0.000% | 6.03E-01 | -0.110 | 0.151 | 0.896 |
| 1682A>T | [1534-1662] | 0 | 4041 | 0.000% | 0 | 2716 | 0.000% | 0.000% | 4.00E-01 | -0.146 | 0.130 | 0.864 |
| 1684T>G | [1534-1662] | 0 | 3906 | 0.000% | 0 | 2716 | 0.000% | 0.000% | 9.74E-01 | -0.007 | 0.124 | 0.993 |
| 1684T>G;1689T>G | [1534-1662] | 0 | 59 | 0.000% | 0 | 2716 | 0.000% | 0.000% | 1.18E-01 | 0.782 | 0.403 | 2.186 |
| 1684T>G;1725T>A | [1534-1662] | 0 | 227 | 0.000% | 0 | 2716 | 0.000% | 0.000% | 7.57E-01 | 0.130 | 0.286 | 1.139 |
| 1689T>G | [1534-1662] | 0 | 2120 | 0.000% | 0 | 2716 | 0.000% | 0.000% | 3.56E-01 | -0.181 | 0.149 | 0.834 |
| 1692G>A | [1534-1662] | 0 | 2979 | 0.000% | 0 | 2716 | 0.000% | 0.000% | 4.20E-01 | -0.141 | 0.131 | 0.869 |
| 1692G>A;1695G>A | [1534-1662] | 0 | 238 | 0.000% | 0 | 2716 | 0.000% | 0.000% | 2.39E-01 | -0.609 | 0.403 | 0.544 |
| 1692G>A;1807A>G | [1534-1662] | 0 | 198 | 0.000% | 0 | 2716 | 0.000% | 0.000% | 4.34E-01 | -0.425 | 0.403 | 0.654 |
| 1714T>A | [1534-1662] | 0 | 1607 | 0.000% | 0 | 2716 | 0.000% | 0.000% | 4.46E-01 | -0.184 | 0.179 | 0.832 |
| 1721T>A | [1534-1662] | 0 | 1078 | 0.000% | 0 | 2716 | 0.000% | 0.000% | 8.80E-01 | -0.041 | 0.176 | 0.960 |
| 1721T>A;1817C>A | [1534-1662] | 0 | 216 | 0.000% | 0 | 2716 | 0.000% | 0.000% | 3.35E-01 | -0.512 | 0.403 | 0.599 |
| 1725T>A | [1534-1662] | 0 | 4376 | 0.000% | 0 | 2716 | 0.000% | 0.000% | 4.51E-01 | -0.109 | 0.107 | 0.897 |
| 1732A>G | [1534-1662] | 0 | 579 | 0.000% | 0 | 2716 | 0.000% | 0.000% | 1.95E-01 | 0.292 | 0.179 | 1.339 |
| 1732A>G;1733T>G | [1534-1662] | 0 | 1037 | 0.000% | 0 | 2716 | 0.000% | 0.000% | 9.95E-01 | -0.002 | 0.171 | 0.998 |
| 1733T>G | [1534-1662] | 0 | 29 | 0.000% | 0 | 2716 | 0.000% | 0.000% | 7.33E-02 | 1.085 | 0.494 | 2.960 |
| 1735C>G | [1534-1662] | 0 | 1982 | 0.000% | 0 | 2716 | 0.000% | 0.000% | 6.41E-01 | -0.090 | 0.137 | 0.914 |
| 1741G>T | [1534-1662] | 0 | 1367 | 0.000% | 0 | 2716 | 0.000% | 0.000% | 7.97E-01 | -0.055 | 0.145 | 0.946 |
| 1742A>C | [1534-1662] | 0 | 183 | 0.000% | 0 | 2716 | 0.000% | 0.000% | 5.31E-01 | -0.346 | 0.402 | 0.707 |
| 1745A>G | [1534-1662] | 0 | 412 | 0.000% | 0 | 2716 | 0.000% | 0.000% | 8.80E-01 | -0.060 | 0.257 | 0.942 |
| 1747A>C | [1534-1662] | 0 | 1350 | 0.000% | 0 | 2716 | 0.000% | 0.000% | 7.23E-02 | -0.399 | 0.182 | 0.671 |
| 1747A>C;1765A>G | [1534-1662] | 0 | 126 | 0.000% | 0 | 2716 | 0.000% | 0.000% | 9.70E-01 | 0.026 | 0.404 | 1.026 |
| 1765A>G;1817C>A | [1534-1662] | 0 | 308 | 0.000% | 0 | 2716 | 0.000% | 0.000% | 7.96E-02 | -0.866 | 0.402 | 0.420 |
| 1785T>G | [1534-1662] | 0 | 758 | 0.000% | 0 | 2716 | 0.000% | 0.000% | 7.31E-01 | -0.094 | 0.189 | 0.910 |
| 1788T>G | [1534-1662] | 0 | 485 | 0.000% | 0 | 2716 | 0.000% | 0.000% | 5.43E-01 | -0.223 | 0.266 | 0.800 |
| 1799A>G | [1534-1662] | 0 | 328 | 0.000% | 0 | 2716 | 0.000% | 0.000% | 6.14E-01 | 0.168 | 0.236 | 1.182 |
| 1802T>G | [1534-1662] | 0 | 579 | 0.000% | 0 | 2716 | 0.000% | 0.000% | 6.85E-01 | 0.110 | 0.190 | 1.117 |
| 1807A>G | [1534-1662] | 0 | 1292 | 0.000% | 0 | 2716 | 0.000% | 0.000% | 3.14E-01 | -0.222 | 0.169 | 0.801 |
| 1826C>G | [1534-1662] | 0 | 86 | 0.000% | 0 | 2716 | 0.000% | 0.000% | 4.56E-01 | 0.407 | 0.403 | 1.502 |
| 1826C>G;1827T>A | [1534-1662] | 0 | 166 | 0.000% | 0 | 2716 | 0.000% | 0.000% | 2.56E-01 | 0.442 | 0.302 | 1.556 |
| 1830T>A | [1534-1662] | 0 | 582 | 0.000% | 0 | 2716 | 0.000% | 0.000% | 7.03E-01 | -0.118 | 0.214 | 0.889 |
| 1834A>G | [1534-1662] | 0 | 233 | 0.000% | 0 | 2716 | 0.000% | 0.000% | 8.06E-01 | 0.104 | 0.285 | 1.110 |
| 1839T>G | [1534-1662] | 0 | 399 | 0.000% | 0 | 2716 | 0.000% | 0.000% | 9.40E-01 | -0.028 | 0.237 | 0.972 |
| 1637T>C | [1534-1673] | 0 | 2581 | 0.000% | 0 | 2716 | 0.000% | 0.000% | 3.84E-02 | -0.354 | 0.143 | 0.702 |
| 1640C>G;1684T>G | [1534-1673] | 0 | 435 | 0.000% | 0 | 2716 | 0.000% | 0.000% | 1.61E-01 | -0.519 | 0.295 | 0.595 |
| 1656A>G | [1534-1673] | 0 | 585 | 0.000% | 0 | 2716 | 0.000% | 0.000% | 3.49E-01 | 0.225 | 0.183 | 1.252 |
| 1656A>G;1657C>G | [1534-1673] | 0 | 644 | 0.000% | 0 | 2716 | 0.000% | 0.000% | 4.41E-01 | -0.219 | 0.210 | 0.804 |
| 1659T>G | [1534-1673] | 0 | 951 | 0.000% | 0 | 2716 | 0.000% | 0.000% | 7.29E-01 | -0.098 | 0.194 | 0.907 |
| 1659T>G;1785T>G | [1534-1673] | 0 | 72 | 0.000% | 0 | 2716 | 0.000% | 0.000% | 2.64E-01 | 0.584 | 0.404 | 1.793 |
| 1661T>A | [1534-1673] | 0 | 289 | 0.000% | 0 | 2716 | 0.000% | 0.000% | 3.43E-01 | 0.294 | 0.236 | 1.341 |
| 1661T>A;1725T>A | [1534-1673] | 0 | 345 | 0.000% | 0 | 2716 | 0.000% | 0.000% | 4.41E-02 | -0.980 | 0.402 | 0.375 |
| 1661T>G | [1534-1673] | 0 | 1259 | 0.000% | 0 | 2716 | 0.000% | 0.000% | 1.39E-01 | -0.329 | 0.179 | 0.720 |
| 1665A>G | [1534-1673] | 0 | 1549 | 0.000% | 0 | 2716 | 0.000% | 0.000% | 7.19E-01 | -0.085 | 0.163 | 0.919 |
| 1670G>C | [1534-1673] | 0 | 785 | 0.000% | 0 | 2716 | 0.000% | 0.000% | 7.15E-01 | 0.094 | 0.177 | 1.098 |
| 1677T>C | [1534-1673] | 0 | 677 | 0.000% | 0 | 2716 | 0.000% | 0.000% | 8.71E-01 | -0.046 | 0.185 | 0.955 |
| 1678A>G | [1534-1673] | 0 | 275 | 0.000% | 0 | 2716 | 0.000% | 0.000% | 8.25E-01 | 0.093 | 0.283 | 1.097 |
| 1678A>T | [1534-1673] | 0 | 1637 | 0.000% | 0 | 2716 | 0.000% | 0.000% | 6.03E-01 | -0.110 | 0.151 | 0.896 |

| | | | | | | | | | | | | |
|---|---|---|---|---|---|---|---|---|---|---|---|---|
| 1684T>G;1689T>G | [1534-1673] | 0 | 59 | 0.000% | 0 | 2716 | 0.000% | 0.000% | 1.18E-01 | 0.782 | 0.403 | 2.186 |
| 1692G>A;1695G>A | [1534-1673] | 0 | 238 | 0.000% | 0 | 2716 | 0.000% | 0.000% | 2.39E-01 | -0.609 | 0.403 | 0.544 |
| 1692G>A;1807A>G | [1534-1673] | 0 | 198 | 0.000% | 0 | 2716 | 0.000% | 0.000% | 4.34E-01 | -0.425 | 0.403 | 0.654 |
| 1714T>A | [1534-1673] | 0 | 1607 | 0.000% | 0 | 2716 | 0.000% | 0.000% | 4.46E-01 | -0.184 | 0.179 | 0.832 |
| 1721T>A | [1534-1673] | 0 | 1078 | 0.000% | 0 | 2716 | 0.000% | 0.000% | 8.80E-01 | -0.041 | 0.176 | 0.960 |
| 1721T>A;1817C>A | [1534-1673] | 0 | 216 | 0.000% | 0 | 2716 | 0.000% | 0.000% | 3.35E-01 | -0.512 | 0.403 | 0.599 |
| 1732A>G | [1534-1673] | 0 | 579 | 0.000% | 0 | 2716 | 0.000% | 0.000% | 1.95E-01 | 0.292 | 0.179 | 1.339 |
| 1733T>G | [1534-1673] | 0 | 29 | 0.000% | 0 | 2716 | 0.000% | 0.000% | 7.33E-02 | 1.085 | 0.494 | 2.960 |
| 1741G>T | [1534-1673] | 0 | 1367 | 0.000% | 0 | 2716 | 0.000% | 0.000% | 7.97E-01 | -0.055 | 0.145 | 0.946 |
| 1742A>C | [1534-1673] | 0 | 183 | 0.000% | 0 | 2716 | 0.000% | 0.000% | 5.31E-01 | -0.346 | 0.402 | 0.707 |
| 1745A>G | [1534-1673] | 0 | 412 | 0.000% | 0 | 2716 | 0.000% | 0.000% | 8.80E-01 | -0.060 | 0.257 | 0.942 |
| 1747A>C | [1534-1673] | 0 | 1350 | 0.000% | 0 | 2716 | 0.000% | 0.000% | 7.23E-02 | -0.399 | 0.182 | 0.671 |
| 1747A>C;1765A>G | [1534-1673] | 0 | 126 | 0.000% | 0 | 2716 | 0.000% | 0.000% | 9.70E-01 | 0.026 | 0.404 | 1.026 |
| 1765A>G;1817C>A | [1534-1673] | 0 | 308 | 0.000% | 0 | 2716 | 0.000% | 0.000% | 7.96E-02 | -0.866 | 0.402 | 0.420 |
| 1788T>G | [1534-1673] | 0 | 485 | 0.000% | 0 | 2716 | 0.000% | 0.000% | 5.43E-01 | -0.223 | 0.266 | 0.800 |
| 1799A>G | [1534-1673] | 0 | 328 | 0.000% | 0 | 2716 | 0.000% | 0.000% | 6.14E-01 | 0.168 | 0.236 | 1.182 |
| 1802T>G | [1534-1673] | 0 | 579 | 0.000% | 0 | 2716 | 0.000% | 0.000% | 6.85E-01 | 0.110 | 0.190 | 1.117 |
| 1807A>G | [1534-1673] | 0 | 1292 | 0.000% | 0 | 2716 | 0.000% | 0.000% | 3.14E-01 | -0.222 | 0.169 | 0.801 |
| 1826C>G | [1534-1673] | 0 | 86 | 0.000% | 0 | 2716 | 0.000% | 0.000% | 4.56E-01 | 0.407 | 0.403 | 1.502 |
| 1826C>G;1827T>A | [1534-1673] | 0 | 166 | 0.000% | 0 | 2716 | 0.000% | 0.000% | 2.56E-01 | 0.442 | 0.302 | 1.556 |
| 1830T>A | [1534-1673] | 0 | 582 | 0.000% | 0 | 2716 | 0.000% | 0.000% | 7.03E-01 | -0.118 | 0.214 | 0.889 |
| 1834A>G | [1534-1673] | 0 | 233 | 0.000% | 0 | 2716 | 0.000% | 0.000% | 8.06E-01 | 0.104 | 0.285 | 1.110 |
| 1637T>C | [1534-1685] | 0 | 1647 | 0.000% | 1 | 2716 | 0.037% | -0.037% | 9.76E-02 | -0.458 | 0.225 | 0.633 |
| 1640C>G | [1534-1685] | 0 | 850 | 0.000% | 1 | 2716 | 0.037% | -0.037% | 5.20E-01 | -0.239 | 0.272 | 0.788 |
| 1640C>G;1684T>G | [1534-1685] | 0 | 292 | 0.000% | 1 | 2716 | 0.037% | -0.037% | 3.69E-01 | -0.673 | 0.567 | 0.510 |
| 1656A>G | [1534-1685] | 0 | 401 | 0.000% | 1 | 2716 | 0.037% | -0.037% | 9.71E-01 | 0.020 | 0.343 | 1.020 |
| 1656A>G;1657C>G | [1534-1685] | 0 | 445 | 0.000% | 1 | 2716 | 0.037% | -0.037% | 4.57E-01 | -0.402 | 0.399 | 0.669 |
| 1659T>G | [1534-1685] | 0 | 606 | 0.000% | 1 | 2716 | 0.037% | -0.037% | 6.25E-01 | -0.226 | 0.326 | 0.798 |
| 1659T>G;1785T>G | [1534-1685] | 0 | 48 | 0.000% | 1 | 2716 | 0.037% | -0.037% | 7.03E-01 | 0.436 | 0.794 | 1.547 |
| 1661T>A | [1534-1685] | 0 | 185 | 0.000% | 1 | 2716 | 0.037% | -0.037% | 7.85E-01 | 0.187 | 0.458 | 1.205 |
| 1661T>A;1725T>A | [1534-1685] | 0 | 220 | 0.000% | 1 | 2716 | 0.037% | -0.037% | 2.95E-01 | -1.083 | 0.793 | 0.339 |
| 1661T>G | [1534-1685] | 0 | 792 | 0.000% | 1 | 2716 | 0.037% | -0.037% | 3.01E-01 | -0.419 | 0.311 | 0.658 |
| 1665A>G | [1534-1685] | 0 | 1042 | 0.000% | 1 | 2716 | 0.037% | -0.037% | 4.90E-01 | -0.242 | 0.257 | 0.785 |
| 1666T>G | [1534-1685] | 0 | 1657 | 0.000% | 1 | 2716 | 0.037% | -0.037% | 2.00E-01 | -0.331 | 0.205 | 0.719 |
| 1670G>A | [1534-1685] | 0 | 1047 | 0.000% | 1 | 2716 | 0.037% | -0.037% | 3.77E-01 | -0.293 | 0.251 | 0.746 |
| 1670G>C | [1534-1685] | 0 | 539 | 0.000% | 1 | 2716 | 0.037% | -0.037% | 8.18E-01 | -0.109 | 0.317 | 0.897 |
| 1677T>C | [1534-1685] | 0 | 445 | 0.000% | 1 | 2716 | 0.037% | -0.037% | 7.27E-01 | -0.179 | 0.353 | 0.836 |
| 1677T>G | [1534-1685] | 0 | 290 | 0.000% | 1 | 2716 | 0.037% | -0.037% | 7.01E-02 | 0.651 | 0.295 | 1.918 |
| 1678A>G | [1534-1685] | 0 | 185 | 0.000% | 1 | 2716 | 0.037% | -0.037% | 9.96E-01 | 0.005 | 0.512 | 1.005 |
| 1682A>T | [1534-1685] | 0 | 2670 | 0.000% | 1 | 2716 | 0.037% | -0.037% | 2.06E-01 | -0.284 | 0.178 | 0.752 |
| 1684T>G | [1534-1685] | 0 | 2541 | 0.000% | 1 | 2716 | 0.037% | -0.037% | 5.72E-01 | -0.130 | 0.166 | 0.878 |
| 1684T>G;1725T>A | [1534-1685] | 0 | 144 | 0.000% | 1 | 2716 | 0.037% | -0.037% | 9.71E-01 | 0.032 | 0.560 | 1.033 |
| 1689T>G | [1534-1685] | 0 | 1352 | 0.000% | 1 | 2716 | 0.037% | -0.037% | 3.26E-01 | -0.297 | 0.231 | 0.743 |
| 1692G>A | [1534-1685] | 0 | 1916 | 0.000% | 1 | 2716 | 0.037% | -0.037% | 3.21E-01 | -0.253 | 0.195 | 0.777 |
| 1692G>A;1695G>A | [1534-1685] | 0 | 165 | 0.000% | 1 | 2716 | 0.037% | -0.037% | 4.59E-01 | -0.796 | 0.793 | 0.451 |
| 1692G>A;1807A>G | [1534-1685] | 0 | 147 | 0.000% | 1 | 2716 | 0.037% | -0.037% | 5.32E-01 | -0.680 | 0.793 | 0.506 |
| 1714T>A | [1534-1685] | 0 | 1071 | 0.000% | 1 | 2716 | 0.037% | -0.037% | 3.78E-01 | -0.315 | 0.271 | 0.730 |
| 1721T>A | [1534-1685] | 0 | 716 | 0.000% | 1 | 2716 | 0.037% | -0.037% | 6.64E-01 | -0.185 | 0.299 | 0.831 |
| 1721T>A;1817C>A | [1534-1685] | 0 | 144 | 0.000% | 1 | 2716 | 0.037% | -0.037% | 5.46E-01 | -0.660 | 0.793 | 0.517 |

| | | | | | | | | | | | | |
|---|---|---|---|---|---|---|---|---|---|---|---|---|
| 1725T>A | [1534-1685] | 0 | 2780 | 0.000% | 1 | 2716 | 0.037% | -0.037% | 2.80E-01 | -0.220 | 0.157 | 0.803 |
| 1732A>G | [1534-1685] | 0 | 365 | 0.000% | 1 | 2716 | 0.037% | -0.037% | 6.67E-01 | 0.200 | 0.328 | 1.222 |
| 1732A>G;1733T>G | [1534-1685] | 0 | 689 | 0.000% | 1 | 2716 | 0.037% | -0.037% | 7.29E-01 | -0.147 | 0.292 | 0.864 |
| 1733T>G | [1534-1685] | 0 | 13 | 0.000% | 1 | 2716 | 0.037% | -0.037% | 4.95E-01 | 1.046 | 1.126 | 2.846 |
| 1735C>G | [1534-1685] | 0 | 1293 | 0.000% | 1 | 2716 | 0.037% | -0.037% | 4.75E-01 | -0.216 | 0.223 | 0.806 |
| 1741G>T | [1534-1685] | 0 | 906 | 0.000% | 1 | 2716 | 0.037% | -0.037% | 5.78E-01 | -0.197 | 0.255 | 0.821 |
| 1742A>C | [1534-1685] | 0 | 120 | 0.000% | 1 | 2716 | 0.037% | -0.037% | 6.73E-01 | -0.478 | 0.793 | 0.620 |
| 1745A>G | [1534-1685] | 0 | 250 | 0.000% | 1 | 2716 | 0.037% | -0.037% | 8.75E-01 | -0.114 | 0.469 | 0.893 |
| 1747A>C | [1534-1685] | 0 | 886 | 0.000% | 1 | 2716 | 0.037% | -0.037% | 1.75E-01 | -0.531 | 0.312 | 0.588 |
| 1747A>C;1765A>G | [1534-1685] | 0 | 78 | 0.000% | 1 | 2716 | 0.037% | -0.037% | 9.71E-01 | -0.048 | 0.794 | 0.953 |
| 1765A>G | [1534-1685] | 0 | 1727 | 0.000% | 1 | 2716 | 0.037% | -0.037% | 1.42E-01 | -0.404 | 0.221 | 0.668 |
| 1765A>G;1817C>A | [1534-1685] | 0 | 216 | 0.000% | 1 | 2716 | 0.037% | -0.037% | 3.04E-01 | -1.065 | 0.793 | 0.345 |
| 1785T>G | [1534-1685] | 0 | 482 | 0.000% | 1 | 2716 | 0.037% | -0.037% | 7.45E-01 | -0.164 | 0.344 | 0.849 |
| 1788T>G | [1534-1685] | 0 | 303 | 0.000% | 1 | 2716 | 0.037% | -0.037% | 6.49E-01 | -0.306 | 0.472 | 0.737 |
| 1799A>G | [1534-1685] | 0 | 227 | 0.000% | 1 | 2716 | 0.037% | -0.037% | 9.82E-01 | -0.017 | 0.457 | 0.983 |
| 1802T>G | [1534-1685] | 0 | 390 | 0.000% | 1 | 2716 | 0.037% | -0.037% | 9.33E-01 | -0.048 | 0.357 | 0.953 |
| 1807A>G | [1534-1685] | 0 | 841 | 0.000% | 1 | 2716 | 0.037% | -0.037% | 3.70E-01 | -0.345 | 0.291 | 0.708 |
| 1817C>A | [1534-1685] | 0 | 1755 | 0.000% | 1 | 2716 | 0.037% | -0.037% | 4.09E-01 | -0.216 | 0.196 | 0.805 |
| 1826C>G | [1534-1685] | 0 | 58 | 0.000% | 1 | 2716 | 0.037% | -0.037% | 8.35E-01 | 0.248 | 0.794 | 1.281 |
| 1826C>G;1827T>A | [1534-1685] | 0 | 117 | 0.000% | 1 | 2716 | 0.037% | -0.037% | 7.77E-01 | 0.239 | 0.570 | 1.270 |
| 1830T>A | [1534-1685] | 0 | 367 | 0.000% | 1 | 2716 | 0.037% | -0.037% | 7.19E-01 | -0.210 | 0.401 | 0.811 |
| 1834A>G | [1534-1685] | 0 | 158 | 0.000% | 1 | 2716 | 0.037% | -0.037% | 9.46E-01 | -0.060 | 0.560 | 0.941 |
| 1839T>G | [1534-1685] | 0 | 275 | 0.000% | 1 | 2716 | 0.037% | -0.037% | 7.58E-01 | -0.209 | 0.459 | 0.812 |
| 1637T>C | [1534-3845] | 0 | 787 | 0.000% | 0 | 2716 | 0.000% | 0.000% | 3.35E-01 | -0.265 | 0.209 | 0.767 |
| 1640C>G | [1534-3845] | 0 | 398 | 0.000% | 0 | 2716 | 0.000% | 0.000% | 9.48E-01 | -0.025 | 0.244 | 0.975 |
| 1640C>G;1684T>G | [1534-3845] | 0 | 121 | 0.000% | 0 | 2716 | 0.000% | 0.000% | 6.30E-01 | -0.338 | 0.496 | 0.713 |
| 1656A>G | [1534-3845] | 0 | 200 | 0.000% | 0 | 2716 | 0.000% | 0.000% | 8.79E-01 | 0.075 | 0.317 | 1.078 |
| 1656A>G;1657C>G | [1534-3845] | 0 | 195 | 0.000% | 0 | 2716 | 0.000% | 0.000% | 8.18E-01 | -0.123 | 0.358 | 0.885 |
| 1659T>G | [1534-3845] | 0 | 290 | 0.000% | 0 | 2716 | 0.000% | 0.000% | 7.98E-01 | -0.114 | 0.302 | 0.892 |
| 1659T>G;1785T>G | [1534-3845] | 0 | 29 | 0.000% | 0 | 2716 | 0.000% | 0.000% | 6.93E-01 | 0.395 | 0.698 | 1.485 |
| 1661T>A | [1534-3845] | 0 | 92 | 0.000% | 0 | 2716 | 0.000% | 0.000% | 5.45E-01 | 0.340 | 0.408 | 1.404 |
| 1661T>A;1725T>A | [1534-3845] | 0 | 111 | 0.000% | 0 | 2716 | 0.000% | 0.000% | 3.00E-01 | -0.944 | 0.697 | 0.389 |
| 1661T>G | [1534-3845] | 0 | 353 | 0.000% | 0 | 2716 | 0.000% | 0.000% | 6.98E-01 | -0.156 | 0.281 | 0.855 |
| 1665A>G | [1534-3845] | 0 | 490 | 0.000% | 0 | 2716 | 0.000% | 0.000% | 9.31E-01 | -0.033 | 0.239 | 0.968 |
| 1666T>G | [1534-3845] | 0 | 786 | 0.000% | 0 | 2716 | 0.000% | 0.000% | 6.22E-01 | -0.130 | 0.187 | 0.878 |
| 1670G>A | [1534-3845] | 0 | 515 | 0.000% | 0 | 2716 | 0.000% | 0.000% | 5.75E-01 | -0.177 | 0.227 | 0.837 |
| 1670G>C | [1534-3845] | 0 | 245 | 0.000% | 0 | 2716 | 0.000% | 0.000% | 9.06E-01 | 0.054 | 0.291 | 1.056 |
| 1677T>C | [1534-3845] | 0 | 213 | 0.000% | 0 | 2716 | 0.000% | 0.000% | 9.82E-01 | 0.012 | 0.314 | 1.012 |
| 1677T>G | [1534-3845] | 0 | 105 | 0.000% | 0 | 2716 | 0.000% | 0.000% | 1.06E-02 | 0.898 | 0.295 | 2.455 |
| 1678A>G | [1534-3845] | 0 | 87 | 0.000% | 0 | 2716 | 0.000% | 0.000% | 4.88E-01 | 0.395 | 0.419 | 1.485 |
| 1678A>T | [1534-3845] | 0 | 530 | 0.000% | 0 | 2716 | 0.000% | 0.000% | 7.35E-01 | -0.111 | 0.226 | 0.895 |
| 1682A>T | [1534-3845] | 0 | 1269 | 0.000% | 0 | 2716 | 0.000% | 0.000% | 7.28E-01 | -0.086 | 0.171 | 0.917 |
| 1684T>G | [1534-3845] | 0 | 1170 | 0.000% | 0 | 2716 | 0.000% | 0.000% | 6.57E-01 | 0.100 | 0.158 | 1.105 |
| 1684T>G;1689T>G | [1534-3845] | 0 | 20 | 0.000% | 0 | 2716 | 0.000% | 0.000% | 4.12E-01 | 0.765 | 0.699 | 2.150 |
| 1684T>G;1725T>A | [1534-3845] | 0 | 66 | 0.000% | 0 | 2716 | 0.000% | 0.000% | 7.09E-01 | 0.267 | 0.493 | 1.305 |
| 1689T>G | [1534-3845] | 0 | 652 | 0.000% | 0 | 2716 | 0.000% | 0.000% | 6.19E-01 | -0.151 | 0.215 | 0.860 |
| 1692G>A | [1534-3845] | 0 | 862 | 0.000% | 0 | 2716 | 0.000% | 0.000% | 1.00E+00 | 0.000 | 0.184 | 1.000 |
| 1692G>A;1695G>A | [1534-3845] | 0 | 76 | 0.000% | 0 | 2716 | 0.000% | 0.000% | 5.57E-01 | -0.566 | 0.697 | 0.568 |
| 1692G>A;1807A>G | [1534-3845] | 0 | 72 | 0.000% | 0 | 2716 | 0.000% | 0.000% | 6.01E-01 | -0.512 | 0.697 | 0.599 |

| | | | | | | | | | | | | |
|---|---|---|---|---|---|---|---|---|---|---|---|---|
| 1714T>A | [1534-3845] | 0 | 483 | 0.000% | 0 | 2716 | 0.000% | 0.000% | 5.28E-01 | -0.219 | 0.253 | 0.804 |
| 1721T>A | [1534-3845] | 0 | 335 | 0.000% | 0 | 2716 | 0.000% | 0.000% | 9.45E-01 | 0.029 | 0.268 | 1.029 |
| 1721T>A;1817C>A | [1534-3845] | 0 | 54 | 0.000% | 0 | 2716 | 0.000% | 0.000% | 8.29E-01 | -0.225 | 0.697 | 0.799 |
| 1732A>G | [1534-3845] | 0 | 176 | 0.000% | 0 | 2716 | 0.000% | 0.000% | 3.22E-01 | 0.384 | 0.296 | 1.468 |
| 1732A>G;1733T>G | [1534-3845] | 0 | 311 | 0.000% | 0 | 2716 | 0.000% | 0.000% | 7.95E-01 | 0.103 | 0.266 | 1.109 |
| 1733T>G | [1534-3845] | 0 | 13 | 0.000% | 0 | 2716 | 0.000% | 0.000% | 1.75E-01 | 1.193 | 0.699 | 3.299 |
| 1735C>G | [1534-3845] | 0 | 618 | 0.000% | 0 | 2716 | 0.000% | 0.000% | 9.40E-01 | -0.024 | 0.201 | 0.977 |
| 1741G>T | [1534-3845] | 0 | 417 | 0.000% | 0 | 2716 | 0.000% | 0.000% | 9.27E-01 | 0.033 | 0.229 | 1.034 |
| 1742A>C | [1534-3845] | 0 | 59 | 0.000% | 0 | 2716 | 0.000% | 0.000% | 7.61E-01 | -0.313 | 0.697 | 0.731 |
| 1745A>G | [1534-3845] | 0 | 129 | 0.000% | 0 | 2716 | 0.000% | 0.000% | 9.98E-01 | 0.003 | 0.423 | 1.003 |
| 1747A>C | [1534-3845] | 0 | 424 | 0.000% | 0 | 2716 | 0.000% | 0.000% | 3.62E-01 | -0.339 | 0.282 | 0.712 |
| 1747A>C;1765A>G | [1534-3845] | 0 | 49 | 0.000% | 0 | 2716 | 0.000% | 0.000% | 9.07E-01 | -0.128 | 0.697 | 0.880 |
| 1765A>G | [1534-3845] | 0 | 810 | 0.000% | 0 | 2716 | 0.000% | 0.000% | 5.68E-01 | -0.160 | 0.203 | 0.852 |
| 1765A>G;1817C>A | [1534-3845] | 0 | 103 | 0.000% | 0 | 2716 | 0.000% | 0.000% | 3.43E-01 | -0.870 | 0.697 | 0.419 |
| 1785T>G | [1534-3845] | 0 | 248 | 0.000% | 0 | 2716 | 0.000% | 0.000% | 7.62E-01 | -0.140 | 0.314 | 0.869 |
| 1788T>G | [1534-3845] | 0 | 156 | 0.000% | 0 | 2716 | 0.000% | 0.000% | 7.67E-01 | -0.187 | 0.427 | 0.829 |
| 1799A>G | [1534-3845] | 0 | 100 | 0.000% | 0 | 2716 | 0.000% | 0.000% | 6.56E-01 | 0.257 | 0.403 | 1.292 |
| 1802T>G | [1534-3845] | 0 | 205 | 0.000% | 0 | 2716 | 0.000% | 0.000% | 9.21E-01 | 0.050 | 0.318 | 1.051 |
| 1807A>G | [1534-3845] | 0 | 426 | 0.000% | 0 | 2716 | 0.000% | 0.000% | 5.63E-01 | -0.211 | 0.263 | 0.810 |
| 1817C>A | [1534-3845] | 0 | 790 | 0.000% | 0 | 2716 | 0.000% | 0.000% | 9.51E-01 | -0.018 | 0.184 | 0.982 |
| 1826C>G | [1534-3845] | 0 | 30 | 0.000% | 0 | 2716 | 0.000% | 0.000% | 7.21E-01 | 0.362 | 0.698 | 1.436 |
| 1826C>G;1827T>A | [1534-3845] | 0 | 49 | 0.000% | 0 | 2716 | 0.000% | 0.000% | 4.06E-01 | 0.563 | 0.507 | 1.757 |
| 1830T>A | [1534-3845] | 0 | 171 | 0.000% | 0 | 2716 | 0.000% | 0.000% | 9.90E-01 | 0.008 | 0.358 | 1.008 |
| 1834A>G | [1534-3845] | 0 | 69 | 0.000% | 0 | 2716 | 0.000% | 0.000% | 7.61E-01 | 0.222 | 0.493 | 1.249 |
| 1839T>G | [1534-3845] | 0 | 145 | 0.000% | 0 | 2716 | 0.000% | 0.000% | 8.53E-01 | -0.114 | 0.405 | 0.892 |
| 412A>G | [1534-NA] | 0 | 282 | 0.000% | 6 | 2716 | 0.221% | -0.221% | 5.76E-01 | -1.855 | 2.388 | 0.157 |
| 423A>T | [1534-NA] | 0 | 1858 | 0.000% | 6 | 2716 | 0.221% | -0.221% | 9.27E-02 | -1.794 | 0.869 | 0.166 |
| 429A>T | [1534-NA] | 0 | 2984 | 0.000% | 6 | 2716 | 0.221% | -0.221% | 2.56E-02 | -1.729 | 0.651 | 0.177 |
| 431A>C | [1534-NA] | 0 | 81 | 0.000% | 6 | 2716 | 0.221% | -0.221% | 7.97E-01 | -1.301 | 3.391 | 0.272 |
| 438A>T | [1534-NA] | 0 | 976 | 0.000% | 6 | 2716 | 0.221% | -0.221% | 2.58E-01 | -2.180 | 1.493 | 0.113 |
| 442A>T | [1534-NA] | 0 | 17220 | 0.000% | 6 | 2716 | 0.221% | -0.221% | 2.83E-08 | -2.229 | 0.347 | 0.108 |
| 492A>G | [1534-NA] | 0 | 618 | 0.000% | 6 | 2716 | 0.221% | -0.221% | 3.88E-01 | -1.723 | 1.499 | 0.179 |
| 493T>A | [1534-NA] | 0 | 398 | 0.000% | 6 | 2716 | 0.221% | -0.221% | 4.99E-01 | -2.199 | 2.388 | 0.111 |
| 493T>C | [1534-NA] | 0 | 751 | 0.000% | 6 | 2716 | 0.221% | -0.221% | 3.39E-01 | -1.582 | 1.257 | 0.206 |
| 493T>G | [1534-NA] | 0 | 440 | 0.000% | 6 | 2716 | 0.221% | -0.221% | 4.80E-01 | -1.607 | 1.676 | 0.201 |
| 1199T>A | [1534-NA] | 0 | 765 | 0.000% | 6 | 2716 | 0.221% | -0.221% | 3.30E-01 | -2.159 | 1.687 | 0.115 |
| 1203C>A | [1534-NA] | 0 | 92 | 0.000% | 6 | 2716 | 0.221% | -0.221% | 7.76E-01 | -1.428 | 3.391 | 0.240 |
| 1203C>G | [1534-NA] | 0 | 359 | 0.000% | 6 | 2716 | 0.221% | -0.221% | 5.51E-01 | -2.789 | 3.390 | 0.061 |
| 1207C>T | [1534-NA] | 0 | 2793 | 0.000% | 6 | 2716 | 0.221% | -0.221% | 3.41E-02 | -2.133 | 0.839 | 0.119 |
| 1207C>T;1231G>A | [1534-NA] | 0 | 1444 | 0.000% | 6 | 2716 | 0.221% | -0.221% | 3.48E-01 | -4.180 | 3.389 | 0.015 |
| 1231G>A | [1534-NA] | 0 | 833 | 0.000% | 6 | 2716 | 0.221% | -0.221% | 3.06E-01 | -2.244 | 1.676 | 0.106 |
| 1231G>C | [1534-NA] | 0 | 11 | 0.000% | 6 | 2716 | 0.221% | -0.221% | 8.97E-01 | 0.688 | 3.404 | 1.990 |
| 1235T>G | [1534-NA] | 0 | 2439 | 0.000% | 6 | 2716 | 0.221% | -0.221% | 6.99E-02 | -2.625 | 1.186 | 0.072 |
| 1235T>G;1337T>G | [1534-NA] | 0 | 18 | 0.000% | 6 | 2716 | 0.221% | -0.221% | 9.71E-01 | 0.199 | 3.398 | 1.220 |
| 1262T>A | [1534-NA] | 0 | 3130 | 0.000% | 6 | 2716 | 0.221% | -0.221% | 3.36E-02 | -2.556 | 1.002 | 0.078 |
| 1262T>A;1321T>G;1: | [1534-NA] | 0 | 583 | 0.000% | 6 | 2716 | 0.221% | -0.221% | 4.77E-01 | -3.274 | 3.389 | 0.038 |
| 1263G>A | [1534-NA] | 0 | 758 | 0.000% | 6 | 2716 | 0.221% | -0.221% | 3.69E-01 | -2.843 | 2.387 | 0.058 |
| 1285G>C | [1534-NA] | 0 | 129 | 0.000% | 6 | 2716 | 0.221% | -0.221% | 7.20E-01 | -1.766 | 3.390 | 0.171 |
| 1296A>C | [1534-NA] | 0 | 1602 | 0.000% | 6 | 2716 | 0.221% | -0.221% | 1.70E-01 | -2.898 | 1.682 | 0.055 |

| | | | | | | | | | | | | |
|---|---|---|---|---|---|---|---|---|---|---|---|---|
| 1297A>G;1403A>C | [1534-NA] | 0 | 36 | 0.000% | 6 | 2716 | 0.221% | -0.221% | 9.27E-01 | -0.491 | 3.394 | 0.612 |
| 1300A>G | [1534-NA] | 0 | 782 | 0.000% | 6 | 2716 | 0.221% | -0.221% | 3.38E-01 | -2.469 | 1.955 | 0.085 |
| 1300A>T | [1534-NA] | 0 | 1792 | 0.000% | 6 | 2716 | 0.221% | -0.221% | 1.18E-01 | -2.451 | 1.261 | 0.086 |
| 1311A>G | [1534-NA] | 0 | 2601 | 0.000% | 6 | 2716 | 0.221% | -0.221% | 4.78E-02 | -2.371 | 0.992 | 0.093 |
| 1311A>G;1317C>T | [1534-NA] | 0 | 416 | 0.000% | 6 | 2716 | 0.221% | -0.221% | 4.90E-01 | -2.243 | 2.388 | 0.106 |
| 1321T>G;1322T>A | [1534-NA] | 0 | 469 | 0.000% | 6 | 2716 | 0.221% | -0.221% | 4.65E-01 | -2.363 | 2.388 | 0.094 |
| 1322T>A | [1534-NA] | 0 | 1213 | 0.000% | 6 | 2716 | 0.221% | -0.221% | 3.72E-01 | -4.006 | 3.389 | 0.018 |
| 1329T>A | [1534-NA] | 0 | 2627 | 0.000% | 6 | 2716 | 0.221% | -0.221% | 4.43E-02 | -2.294 | 0.946 | 0.101 |
| 1337T>G | [1534-NA] | 0 | 1864 | 0.000% | 6 | 2716 | 0.221% | -0.221% | 1.19E-01 | -2.644 | 1.365 | 0.071 |
| 1349G>A | [1534-NA] | 0 | 1742 | 0.000% | 6 | 2716 | 0.221% | -0.221% | 1.16E-01 | -2.289 | 1.174 | 0.101 |
| 1362A>G | [1534-NA] | 0 | 3489 | 0.000% | 6 | 2716 | 0.221% | -0.221% | 2.24E-02 | -2.578 | 0.946 | 0.076 |
| 1378A>G | [1534-NA] | 0 | 162 | 0.000% | 6 | 2716 | 0.221% | -0.221% | 6.81E-01 | -1.993 | 3.390 | 0.136 |
| 1379A>G | [1534-NA] | 0 | 415 | 0.000% | 6 | 2716 | 0.221% | -0.221% | 4.88E-01 | -1.836 | 1.943 | 0.160 |
| 1392C>G | [1534-NA] | 0 | 772 | 0.000% | 6 | 2716 | 0.221% | -0.221% | 3.42E-01 | -2.456 | 1.964 | 0.086 |
| 1392C>T | [1534-NA] | 0 | 1116 | 0.000% | 6 | 2716 | 0.221% | -0.221% | 2.18E-01 | -2.131 | 1.358 | 0.119 |
| 1407T>C;1533G>T | [1534-NA] | 0 | 31 | 0.000% | 6 | 2716 | 0.221% | -0.221% | 9.49E-01 | -0.342 | 3.394 | 0.710 |
| 1412T>A | [1534-NA] | 0 | 351 | 0.000% | 6 | 2716 | 0.221% | -0.221% | 5.55E-01 | -2.766 | 3.390 | 0.063 |
| 1419T>G | [1534-NA] | 0 | 1411 | 0.000% | 6 | 2716 | 0.221% | -0.221% | 1.60E-01 | -2.212 | 1.254 | 0.110 |
| 1425T>A | [1534-NA] | 0 | 1883 | 0.000% | 6 | 2716 | 0.221% | -0.221% | 9.17E-02 | -1.962 | 0.945 | 0.141 |
| 1429A>G | [1534-NA] | 0 | 1341 | 0.000% | 6 | 2716 | 0.221% | -0.221% | 1.67E-01 | -1.910 | 1.101 | 0.148 |
| 1432T>G | [1534-NA] | 0 | 367 | 0.000% | 6 | 2716 | 0.221% | -0.221% | 5.20E-01 | -1.713 | 1.944 | 0.180 |
| 1444T>G | [1534-NA] | 0 | 762 | 0.000% | 6 | 2716 | 0.221% | -0.221% | 3.39E-01 | -2.443 | 1.942 | 0.087 |
| 1450A>G | [1534-NA] | 0 | 1373 | 0.000% | 6 | 2716 | 0.221% | -0.221% | 1.62E-01 | -1.828 | 1.042 | 0.161 |
| 1492G>T | [1534-NA] | 0 | 726 | 0.000% | 6 | 2716 | 0.221% | -0.221% | 3.50E-01 | -2.395 | 1.946 | 0.091 |
| 1499G>T | [1534-NA] | 0 | 840 | 0.000% | 6 | 2716 | 0.221% | -0.221% | 3.02E-01 | -2.030 | 1.506 | 0.131 |
| 1503A>G | [1534-NA] | 0 | 2352 | 0.000% | 6 | 2716 | 0.221% | -0.221% | 5.51E-02 | -1.779 | 0.766 | 0.169 |
| 1520T>C | [1534-NA] | 0 | 382 | 0.000% | 6 | 2716 | 0.221% | -0.221% | 5.24E-01 | -1.465 | 1.677 | 0.231 |
| 1524C>T | [1534-NA] | 0 | 1331 | 0.000% | 6 | 2716 | 0.221% | -0.221% | 1.74E-01 | -1.615 | 0.946 | 0.199 |
| 1526T>A;1530T>G | [1534-NA] | 0 | 319 | 0.000% | 6 | 2716 | 0.221% | -0.221% | 5.58E-01 | -1.573 | 1.943 | 0.207 |
| 1530T>G | [1534-NA] | 0 | 947 | 0.000% | 6 | 2716 | 0.221% | -0.221% | 2.69E-01 | -1.680 | 1.172 | 0.186 |
| 1532A>T | [1534-NA] | 0 | 3179 | 0.000% | 6 | 2716 | 0.221% | -0.221% | 1.99E-02 | -1.792 | 0.648 | 0.167 |
| 1532A>T;1533G>T | [1534-NA] | 0 | 18 | 0.000% | 6 | 2716 | 0.221% | -0.221% | 9.71E-01 | 0.199 | 3.398 | 1.220 |
| 1533G>A | [1534-NA] | 0 | 760 | 0.000% | 6 | 2716 | 0.221% | -0.221% | 3.35E-01 | -1.593 | 1.256 | 0.203 |
| 1533G>C | [1534-NA] | 0 | 839 | 0.000% | 6 | 2716 | 0.221% | -0.221% | 2.98E-01 | -2.029 | 1.494 | 0.132 |
| 1533G>T | [1534-NA] | 0 | 758 | 0.000% | 6 | 2716 | 0.221% | -0.221% | 3.31E-01 | -2.150 | 1.682 | 0.116 |
| 1534T>A | [1534-NA] | 0 | 511 | 0.000% | 6 | 2716 | 0.221% | -0.221% | 4.38E-01 | -1.756 | 1.678 | 0.173 |
| 1534T>G | [1534-NA] | 0 | 546 | 0.000% | 6 | 2716 | 0.221% | -0.221% | 4.28E-01 | -1.599 | 1.500 | 0.202 |
| 1558T>A | [1534-NA] | 0 | 2529 | 0.000% | 6 | 2716 | 0.221% | -0.221% | 4.60E-02 | -2.176 | 0.905 | 0.113 |
| 1562T>A | [1534-NA] | 0 | 2530 | 0.000% | 6 | 2716 | 0.221% | -0.221% | 4.45E-02 | -1.909 | 0.789 | 0.148 |
| 1566A>G | [1534-NA] | 0 | 4376 | 0.000% | 6 | 2716 | 0.221% | -0.221% | 5.07E-03 | -1.793 | 0.546 | 0.166 |
| 1578A>G | [1534-NA] | 0 | 5576 | 0.000% | 6 | 2716 | 0.221% | -0.221% | 1.25E-03 | -1.843 | 0.492 | 0.158 |
| 1578A>G;1590C>T | [1534-NA] | 0 | 55 | 0.000% | 6 | 2716 | 0.221% | -0.221% | 8.59E-01 | -0.914 | 3.392 | 0.401 |
| 1595C>T;1605G>A | [1534-NA] | 0 | 376 | 0.000% | 6 | 2716 | 0.221% | -0.221% | 5.43E-01 | -2.835 | 3.389 | 0.059 |
| 1600T>A | [1534-NA] | 0 | 1135 | 0.000% | 6 | 2716 | 0.221% | -0.221% | 2.11E-01 | -1.994 | 1.255 | 0.136 |
| 1605G>A | [1534-NA] | 0 | 1279 | 0.000% | 6 | 2716 | 0.221% | -0.221% | 1.77E-01 | -1.980 | 1.169 | 0.138 |
| 1623T>C | [1534-NA] | 0 | 37 | 0.000% | 6 | 2716 | 0.221% | -0.221% | 9.23E-01 | -0.519 | 3.394 | 0.595 |
| 1659T>G;1785T>G | [1534-NA] | 0 | 72 | 0.000% | 6 | 2716 | 0.221% | -0.221% | 9.79E-01 | -0.087 | 1.945 | 0.916 |
| 1661T>A;1725T>A | [1534-NA] | 0 | 345 | 0.000% | 6 | 2716 | 0.221% | -0.221% | 5.37E-01 | -1.651 | 1.942 | 0.192 |
| 1665A>G | [1534-NA] | 0 | 1549 | 0.000% | 6 | 2716 | 0.221% | -0.221% | 2.86E-01 | -0.756 | 0.547 | 0.469 |

| | | | | | | | | | | | | |
|---|---|---|---|---|---|---|---|---|---|---|---|---|
| 1684T>G;1725T>A | [1534-NA] | 0 | 227 | 0.000% | 6 | 2716 | 0.221% | -0.221% | 7.89E-01 | -0.541 | 1.360 | 0.582 |
| 1692G>A;1695G>A | [1534-NA] | 0 | 238 | 0.000% | 6 | 2716 | 0.221% | -0.221% | 6.43E-01 | -1.280 | 1.943 | 0.278 |
| 1692G>A;1807A>G | [1534-NA] | 0 | 198 | 0.000% | 6 | 2716 | 0.221% | -0.221% | 6.93E-01 | -1.096 | 1.943 | 0.334 |
| 1721T>A;1817C>A | [1534-NA] | 0 | 216 | 0.000% | 6 | 2716 | 0.221% | -0.221% | 6.68E-01 | -1.183 | 1.943 | 0.306 |
| 1733T>G | [1534-NA] | 0 | 29 | 0.000% | 6 | 2716 | 0.221% | -0.221% | 9.12E-01 | 0.414 | 2.395 | 1.513 |
| 1745A>G | [1534-NA] | 0 | 412 | 0.000% | 6 | 2716 | 0.221% | -0.221% | 6.40E-01 | -0.731 | 1.103 | 0.481 |
| 1747A>C | [1534-NA] | 0 | 1350 | 0.000% | 6 | 2716 | 0.221% | -0.221% | 2.31E-01 | -1.070 | 0.700 | 0.343 |
| 1747A>C;1765A>G | [1534-NA] | 0 | 126 | 0.000% | 6 | 2716 | 0.221% | -0.221% | 8.23E-01 | -0.645 | 1.944 | 0.525 |
| 1785T>G | [1534-NA] | 0 | 758 | 0.000% | 6 | 2716 | 0.221% | -0.221% | 4.86E-01 | -0.765 | 0.809 | 0.465 |
| 1802T>G | [1534-NA] | 0 | 579 | 0.000% | 6 | 2716 | 0.221% | -0.221% | 6.36E-01 | -0.561 | 0.837 | 0.571 |
| 1826C>G | [1534-NA] | 0 | 86 | 0.000% | 6 | 2716 | 0.221% | -0.221% | 9.31E-01 | -0.264 | 1.945 | 0.768 |
| 1826C>G;1827T>A | [1534-NA] | 0 | 166 | 0.000% | 6 | 2716 | 0.221% | -0.221% | 9.15E-01 | -0.229 | 1.363 | 0.795 |
| 1834A>G | [1534-NA] | 0 | 233 | 0.000% | 6 | 2716 | 0.221% | -0.221% | 7.78E-01 | -0.567 | 1.359 | 0.567 |
| 1853T>G | [1534-NA] | 0 | 1335 | 0.000% | 6 | 2716 | 0.221% | -0.221% | 4.73E-01 | -0.494 | 0.508 | 0.610 |
| 1853T>G;1892T>A | [1534-NA] | 0 | 65 | 0.000% | 6 | 2716 | 0.221% | -0.221% | 8.31E-01 | -1.081 | 3.392 | 0.339 |
| 1853T>G;1898A>G | [1534-NA] | 0 | 26 | 0.000% | 6 | 2716 | 0.221% | -0.221% | 9.76E-01 | -0.167 | 3.395 | 0.846 |
| 1853T>G;1905T>C | [1534-NA] | 0 | 27 | 0.000% | 6 | 2716 | 0.221% | -0.221% | 9.71E-01 | -0.205 | 3.395 | 0.815 |
| 1853T>G;2065A>G | [1534-NA] | 0 | 131 | 0.000% | 6 | 2716 | 0.221% | -0.221% | 7.18E-01 | -1.781 | 3.390 | 0.168 |
| 1856A>G | [1534-NA] | 0 | 838 | 0.000% | 6 | 2716 | 0.221% | -0.221% | 7.49E-01 | -0.272 | 0.583 | 0.762 |
| 1856A>G;1917T>G | [1534-NA] | 0 | 17 | 0.000% | 6 | 2716 | 0.221% | -0.221% | 9.64E-01 | 0.256 | 3.399 | 1.291 |
| 1857A>G | [1534-NA] | 0 | 358 | 0.000% | 6 | 2716 | 0.221% | -0.221% | 7.50E-01 | -0.486 | 1.042 | 0.615 |
| 1857A>G;1858T>G | [1534-NA] | 0 | 605 | 0.000% | 6 | 2716 | 0.221% | -0.221% | 7.96E-01 | -0.269 | 0.697 | 0.764 |
| 1857A>G;1858T>G;1898A>T | [1534-NA] | 0 | 68 | 0.000% | 6 | 2716 | 0.221% | -0.221% | 8.23E-01 | -1.126 | 3.392 | 0.324 |
| 1857A>G;1858T>G;1951A>G | [1534-NA] | 0 | 27 | 0.000% | 6 | 2716 | 0.221% | -0.221% | 9.71E-01 | -0.205 | 3.395 | 0.815 |
| 1857A>G;1858T>G;2053T>G | [1534-NA] | 0 | 50 | 0.000% | 6 | 2716 | 0.221% | -0.221% | 8.75E-01 | -0.819 | 3.392 | 0.441 |
| 1857A>G;1858T>G;2065A>T | [1534-NA] | 0 | 50 | 0.000% | 6 | 2716 | 0.221% | -0.221% | 8.75E-01 | -0.819 | 3.392 | 0.441 |
| 1857A>G;1858T>G;del2053 | [1534-NA] | 0 | 33 | 0.000% | 6 | 2716 | 0.221% | -0.221% | 9.40E-01 | -0.405 | 3.394 | 0.667 |
| 1857A>G;1951A>G | [1534-NA] | 0 | 18 | 0.000% | 6 | 2716 | 0.221% | -0.221% | 9.71E-01 | 0.199 | 3.398 | 1.220 |
| 1858T>G | [1534-NA] | 0 | 25 | 0.000% | 6 | 2716 | 0.221% | -0.221% | 9.82E-01 | -0.128 | 3.396 | 0.880 |
| 1862C>G | [1534-NA] | 0 | 1282 | 0.000% | 6 | 2716 | 0.221% | -0.221% | 6.93E-01 | -0.258 | 0.455 | 0.772 |
| 1862C>G;1898A>G | [1534-NA] | 0 | 33 | 0.000% | 6 | 2716 | 0.221% | -0.221% | 9.40E-01 | -0.405 | 3.394 | 0.667 |
| 1862C>G;1961A>G | [1534-NA] | 0 | 28 | 0.000% | 6 | 2716 | 0.221% | -0.221% | 9.66E-01 | -0.241 | 3.395 | 0.786 |
| 1862C>G;2065A>G | [1534-NA] | 0 | 115 | 0.000% | 6 | 2716 | 0.221% | -0.221% | 7.87E-01 | -0.959 | 2.391 | 0.383 |
| 1865T>G | [1534-NA] | 0 | 2286 | 0.000% | 6 | 2716 | 0.221% | -0.221% | 4.50E-01 | -0.352 | 0.346 | 0.703 |
| 1865T>G;1877T>G | [1534-NA] | 0 | 64 | 0.000% | 6 | 2716 | 0.221% | -0.221% | 8.34E-01 | -1.066 | 3.392 | 0.345 |
| 1865T>G;1892T>A | [1534-NA] | 0 | 16 | 0.000% | 6 | 2716 | 0.221% | -0.221% | 9.53E-01 | 0.316 | 3.399 | 1.372 |
| 1865T>G;1898A>T | [1534-NA] | 0 | 57 | 0.000% | 6 | 2716 | 0.221% | -0.221% | 8.53E-01 | -0.950 | 3.392 | 0.387 |
| 1865T>G;1917T>C | [1534-NA] | 0 | 63 | 0.000% | 6 | 2716 | 0.221% | -0.221% | 9.25E-01 | -0.358 | 2.394 | 0.699 |
| 1865T>G;1962A>G | [1534-NA] | 0 | 57 | 0.000% | 6 | 2716 | 0.221% | -0.221% | 8.53E-01 | -0.950 | 3.392 | 0.387 |
| 1865T>G;1994A>G | [1534-NA] | 0 | 32 | 0.000% | 6 | 2716 | 0.221% | -0.221% | 9.44E-01 | -0.374 | 3.394 | 0.688 |
| 1865T>G;2013T>G | [1534-NA] | 0 | 94 | 0.000% | 6 | 2716 | 0.221% | -0.221% | 7.73E-01 | -1.450 | 3.391 | 0.235 |
| 1865T>G;2050A>G | [1534-NA] | 0 | 19 | 0.000% | 6 | 2716 | 0.221% | -0.221% | 9.80E-01 | 0.145 | 3.398 | 1.156 |
| 1869A>G | [1534-NA] | 0 | 1356 | 0.000% | 6 | 2716 | 0.221% | -0.221% | 7.61E-01 | -0.189 | 0.423 | 0.827 |
| 1869A>G;1898A>G | [1534-NA] | 0 | 38 | 0.000% | 6 | 2716 | 0.221% | -0.221% | 9.71E-01 | 0.145 | 2.393 | 1.156 |

| | | | | | | | | | | | | | |
|---|---|---|---|---|---|---|---|---|---|---|---|---|---|
| 1869A>G;1951A>G | [1534-NA] | | 0 | 81 | 0.000% | 6 | 2716 | 0.221% | -0.221% | 7.97E-01 | -1.301 | 3.391 | 0.272 |
| 1872A>G | [1534-NA] | | 0 | 385 | 0.000% | 6 | 2716 | 0.221% | -0.221% | 7.12E-01 | -0.558 | 1.042 | 0.572 |
| 1872A>G;1917T>G | [1534-NA] | | 0 | 16 | 0.000% | 6 | 2716 | 0.221% | -0.221% | 9.53E-01 | 0.316 | 3.399 | 1.372 |
| 1877T>G | [1534-NA] | | 0 | 81 | 0.000% | 6 | 2716 | 0.221% | -0.221% | 9.77E-01 | 0.082 | 1.680 | 1.085 |
| 1892T>A | [1534-NA] | | 0 | 739 | 0.000% | 6 | 2716 | 0.221% | -0.221% | 6.92E-01 | -0.378 | 0.664 | 0.685 |
| 1892T>A;1898A>G | [1534-NA] | | 0 | 21 | 0.000% | 6 | 2716 | 0.221% | -0.221% | 9.95E-01 | 0.046 | 3.397 | 1.047 |
| 1892T>A;1907T>G | [1534-NA] | | 0 | 22 | 0.000% | 6 | 2716 | 0.221% | -0.221% | 1.00E+00 | -0.001 | 3.397 | 0.999 |
| 1892T>A;del1918 | [1534-NA] | | 0 | 69 | 0.000% | 6 | 2716 | 0.221% | -0.221% | 9.05E-01 | -0.449 | 2.392 | 0.638 |
| 1896C>G | [1534-NA] | | 0 | 994 | 0.000% | 6 | 2716 | 0.221% | -0.221% | 8.15E-01 | -0.173 | 0.500 | 0.841 |
| 1896C>G;1917T>G | [1534-NA] | | 0 | 81 | 0.000% | 6 | 2716 | 0.221% | -0.221% | 7.97E-01 | -1.301 | 3.391 | 0.272 |
| 1896C>G;1920T>G | [1534-NA] | | 0 | 49 | 0.000% | 6 | 2716 | 0.221% | -0.221% | 8.79E-01 | -0.799 | 3.392 | 0.450 |
| 1898A>G | [1534-NA] | | 0 | 734 | 0.000% | 6 | 2716 | 0.221% | -0.221% | 7.87E-01 | -0.249 | 0.620 | 0.779 |
| 1898A>G;1962A>G | [1534-NA] | | 0 | 30 | 0.000% | 6 | 2716 | 0.221% | -0.221% | 9.54E-01 | -0.310 | 3.395 | 0.734 |
| 1898A>T | [1534-NA] | | 0 | 406 | 0.000% | 6 | 2716 | 0.221% | -0.221% | 8.31E-01 | -0.276 | 0.870 | 0.759 |
| 1905T>C | [1534-NA] | | 0 | 861 | 0.000% | 6 | 2716 | 0.221% | -0.221% | 7.79E-01 | -0.233 | 0.561 | 0.792 |
| 1905T>C;2065A>T | [1534-NA] | | 0 | 22 | 0.000% | 6 | 2716 | 0.221% | -0.221% | 1.00E+00 | -0.001 | 3.397 | 0.999 |
| 1907T>G | [1534-NA] | | 0 | 654 | 0.000% | 6 | 2716 | 0.221% | -0.221% | 8.57E-01 | -0.173 | 0.632 | 0.841 |
| 1907T>G;del1911..19 | [1534-NA] | | 0 | 12 | 0.000% | 6 | 2716 | 0.221% | -0.221% | 9.10E-01 | 0.602 | 3.403 | 1.825 |
| 1907T>G;del1920 | [1534-NA] | | 0 | 52 | 0.000% | 6 | 2716 | 0.221% | -0.221% | 8.68E-01 | -0.858 | 3.392 | 0.424 |
| 1917T>C | [1534-NA] | | 0 | 250 | 0.000% | 6 | 2716 | 0.221% | -0.221% | 7.49E-01 | -0.637 | 1.361 | 0.529 |
| 1917T>G | [1534-NA] | | 0 | 647 | 0.000% | 6 | 2716 | 0.221% | -0.221% | 7.73E-01 | -0.290 | 0.679 | 0.748 |
| 1917T>G;1969C>T | [1534-NA] | | 0 | 55 | 0.000% | 6 | 2716 | 0.221% | -0.221% | 8.59E-01 | -0.914 | 3.392 | 0.401 |
| 1917T>G;del1930 | [1534-NA] | | 0 | 37 | 0.000% | 6 | 2716 | 0.221% | -0.221% | 9.23E-01 | -0.519 | 3.394 | 0.595 |
| 1920T>G | [1534-NA] | | 0 | 940 | 0.000% | 6 | 2716 | 0.221% | -0.221% | 5.89E-01 | -0.458 | 0.608 | 0.633 |
| 1920T>G;2053T>G | [1534-NA] | | 0 | 71 | 0.000% | 6 | 2716 | 0.221% | -0.221% | 8.16E-01 | -1.169 | 3.391 | 0.311 |
| 1945C>G | [1534-NA] | | 0 | 336 | 0.000% | 6 | 2716 | 0.221% | -0.221% | 9.49E-01 | -0.087 | 0.870 | 0.916 |
| 1951A>G | [1534-NA] | | 0 | 521 | 0.000% | 6 | 2716 | 0.221% | -0.221% | 8.42E-01 | -0.220 | 0.737 | 0.802 |
| 1952A>G | [1534-NA] | | 0 | 119 | 0.000% | 6 | 2716 | 0.221% | -0.221% | 8.40E-01 | -0.588 | 1.945 | 0.555 |
| 1952A>G;1953T>G | [1534-NA] | | 0 | 137 | 0.000% | 6 | 2716 | 0.221% | -0.221% | 8.00E-01 | -0.729 | 1.944 | 0.483 |
| 1953T>G | [1534-NA] | | 0 | 82 | 0.000% | 6 | 2716 | 0.221% | -0.221% | 9.44E-01 | -0.217 | 1.946 | 0.805 |
| 1956A>G | [1534-NA] | | 0 | 10 | 0.000% | 6 | 2716 | 0.221% | -0.221% | 8.81E-01 | 0.782 | 3.406 | 2.187 |
| 1961A>G | [1534-NA] | | 0 | 1090 | 0.000% | 6 | 2716 | 0.221% | -0.221% | 6.19E-01 | -0.376 | 0.535 | 0.687 |
| 1961A>G;2060A>T | [1534-NA] | | 0 | 29 | 0.000% | 6 | 2716 | 0.221% | -0.221% | 9.61E-01 | -0.276 | 3.395 | 0.759 |
| 1962A>G | [1534-NA] | | 0 | 412 | 0.000% | 6 | 2716 | 0.221% | -0.221% | 8.23E-01 | -0.291 | 0.869 | 0.748 |
| 1969C>T | [1534-NA] | | 0 | 492 | 0.000% | 6 | 2716 | 0.221% | -0.221% | 7.80E-01 | -0.334 | 0.809 | 0.716 |
| 1997A>C | [1534-NA] | | 0 | 110 | 0.000% | 6 | 2716 | 0.221% | -0.221% | 9.33E-01 | -0.223 | 1.680 | 0.800 |
| 1997A>T | [1534-NA] | | 0 | 169 | 0.000% | 6 | 2716 | 0.221% | -0.221% | 9.65E-01 | -0.093 | 1.256 | 0.911 |
| 2011C>G | [1534-NA] | | 0 | 501 | 0.000% | 6 | 2716 | 0.221% | -0.221% | 7.67E-01 | -0.352 | 0.807 | 0.703 |
| 2013T>G | [1534-NA] | | 0 | 473 | 0.000% | 6 | 2716 | 0.221% | -0.221% | 6.99E-01 | -0.502 | 0.906 | 0.605 |
| 2016T>A | [1534-NA] | | 0 | 234 | 0.000% | 6 | 2716 | 0.221% | -0.221% | 9.24E-01 | -0.167 | 1.101 | 0.846 |
| 2016T>G | [1534-NA] | | 0 | 318 | 0.000% | 6 | 2716 | 0.221% | -0.221% | 8.12E-01 | -0.368 | 1.040 | 0.692 |
| 2026T>G | [1534-NA] | | 0 | 590 | 0.000% | 6 | 2716 | 0.221% | -0.221% | 4.76E-01 | -3.285 | 3.389 | 0.037 |
| 2050A>G | [1534-NA] | | 0 | 319 | 0.000% | 6 | 2716 | 0.221% | -0.221% | 7.69E-01 | -0.476 | 1.099 | 0.621 |
| 2053T>G | [1534-NA] | | 0 | 359 | 0.000% | 6 | 2716 | 0.221% | -0.221% | 8.27E-01 | -0.307 | 0.945 | 0.736 |
| 2055T>G | [1534-NA] | | 0 | 75 | 0.000% | 6 | 2716 | 0.221% | -0.221% | 9.70E-01 | -0.128 | 1.946 | 0.880 |
| 2055T>G;2057A>G | [1534-NA] | | 0 | 103 | 0.000% | 6 | 2716 | 0.221% | -0.221% | 8.82E-01 | -0.444 | 1.945 | 0.641 |
| 2059G>T | [1534-NA] | | 0 | 84 | 0.000% | 6 | 2716 | 0.221% | -0.221% | 9.38E-01 | -0.241 | 1.947 | 0.786 |
| 2060A>T | [1534-NA] | | 0 | 92 | 0.000% | 6 | 2716 | 0.221% | -0.221% | 9.88E-01 | -0.045 | 1.679 | 0.956 |
| 2065A>G | [1534-NA] | | 0 | 413 | 0.000% | 6 | 2716 | 0.221% | -0.221% | 7.45E-01 | -0.447 | 0.943 | 0.640 |

| | | | | | | | | | | | | |
|---|---|---|---|---|---|---|---|---|---|---|---|---|
| 2065A>T | [1534-NA] | 0 | 825 | 0.000% | 6 | 2716 | 0.221% | -0.221% | 6.53E-01 | -0.405 | 0.632 | 0.667 |
| 2514T>G | [1534-NA] | 0 | 1003 | 0.000% | 6 | 2716 | 0.221% | -0.221% | 2.50E-01 | -2.025 | 1.368 | 0.132 |
| 2519A>C | [1534-NA] | 0 | 2157 | 0.000% | 6 | 2716 | 0.221% | -0.221% | 1.00E-01 | -2.790 | 1.375 | 0.061 |
| 2519A>G | [1534-NA] | 0 | 414 | 0.000% | 6 | 2716 | 0.221% | -0.221% | 4.91E-01 | -2.238 | 2.389 | 0.107 |
| 2524T>A | [1534-NA] | 0 | 11998 | 0.000% | 6 | 2716 | 0.221% | -0.221% | 8.43E-06 | -2.634 | 0.507 | 0.072 |
| 2524T>A;2532A>C | [1534-NA] | 0 | 374 | 0.000% | 6 | 2716 | 0.221% | -0.221% | 5.44E-01 | -2.830 | 3.389 | 0.059 |
| 2532A>C | [1534-NA] | 0 | 7077 | 0.000% | 6 | 2716 | 0.221% | -0.221% | 6.10E-04 | -2.551 | 0.636 | 0.078 |
| 2532A>C;2590A>G | [1534-NA] | 0 | 164 | 0.000% | 6 | 2716 | 0.221% | -0.221% | 6.79E-01 | -2.006 | 3.390 | 0.135 |
| 2532A>C;2643G>C | [1534-NA] | 0 | 499 | 0.000% | 6 | 2716 | 0.221% | -0.221% | 4.54E-01 | -2.425 | 2.389 | 0.088 |
| 2532A>T | [1534-NA] | 0 | 6201 | 0.000% | 6 | 2716 | 0.221% | -0.221% | 1.87E-03 | -2.693 | 0.739 | 0.068 |
| 2532A>T;2554T>G | [1534-NA] | 0 | 716 | 0.000% | 6 | 2716 | 0.221% | -0.221% | 3.53E-01 | -2.381 | 1.945 | 0.092 |
| 2551T>G | [1534-NA] | 0 | 1626 | 0.000% | 6 | 2716 | 0.221% | -0.221% | 1.28E-01 | -2.220 | 1.173 | 0.109 |
| 2554T>G | [1534-NA] | 0 | 5451 | 0.000% | 6 | 2716 | 0.221% | -0.221% | 3.16E-03 | -2.565 | 0.740 | 0.077 |
| 2562T>A | [1534-NA] | 0 | 963 | 0.000% | 6 | 2716 | 0.221% | -0.221% | 2.89E-01 | -2.677 | 1.942 | 0.069 |
| 2562T>G | [1534-NA] | 0 | 2190 | 0.000% | 6 | 2716 | 0.221% | -0.221% | 7.19E-02 | -2.295 | 1.043 | 0.101 |
| 2563G>A | [1534-NA] | 0 | 1936 | 0.000% | 6 | 2716 | 0.221% | -0.221% | 1.40E-01 | -3.087 | 1.675 | 0.046 |
| 2566G>A | [1534-NA] | 0 | 968 | 0.000% | 6 | 2716 | 0.221% | -0.221% | 4.04E-01 | -3.780 | 3.389 | 0.023 |
| 2566G>A;2567T>A | [1534-NA] | 0 | 635 | 0.000% | 6 | 2716 | 0.221% | -0.221% | 3.76E-01 | -1.973 | 1.684 | 0.139 |
| 2567T>A | [1534-NA] | 0 | 1833 | 0.000% | 6 | 2716 | 0.221% | -0.221% | 1.21E-01 | -2.627 | 1.365 | 0.072 |
| 2567T>A;2590A>G | [1534-NA] | 0 | 295 | 0.000% | 6 | 2716 | 0.221% | -0.221% | 5.83E-01 | -2.592 | 3.390 | 0.075 |
| 2579A>G | [1534-NA] | 0 | 573 | 0.000% | 6 | 2716 | 0.221% | -0.221% | 4.06E-01 | -1.870 | 1.681 | 0.154 |
| 2584T>G | [1534-NA] | 0 | 2690 | 0.000% | 6 | 2716 | 0.221% | -0.221% | 5.50E-02 | -2.723 | 1.169 | 0.066 |
| 2584T>G;2592C>T | [1534-NA] | 0 | 251 | 0.000% | 6 | 2716 | 0.221% | -0.221% | 6.12E-01 | -2.431 | 3.390 | 0.088 |
| 2590A>G | [1534-NA] | 0 | 15026 | 0.000% | 6 | 2716 | 0.221% | -0.221% | 4.77E-07 | -2.611 | 0.444 | 0.073 |
| 2608C>G | [1534-NA] | 0 | 1503 | 0.000% | 6 | 2716 | 0.221% | -0.221% | 1.64E-01 | -2.611 | 1.493 | 0.073 |
| 2642A>G | [1534-NA] | 0 | 2984 | 0.000% | 6 | 2716 | 0.221% | -0.221% | 3.26E-02 | -2.422 | 0.944 | 0.089 |
| 2642A>T | [1534-NA] | 0 | 1106 | 0.000% | 6 | 2716 | 0.221% | -0.221% | 2.64E-01 | -2.815 | 1.944 | 0.060 |
| 2643G>C | [1534-NA] | 0 | 4685 | 0.000% | 6 | 2716 | 0.221% | -0.221% | 9.42E-03 | -2.719 | 0.884 | 0.066 |
| 2643G>T | [1534-NA] | 0 | 2205 | 0.000% | 6 | 2716 | 0.221% | -0.221% | 7.79E-02 | -2.525 | 1.168 | 0.080 |
| 2989C>G | [1534-NA] | 0 | 1246 | 0.000% | 6 | 2716 | 0.221% | -0.221% | 1.84E-01 | -1.836 | 1.098 | 0.159 |
| 3007C>G | [1534-NA] | 0 | 5418 | 0.000% | 6 | 2716 | 0.221% | -0.221% | 1.49E-03 | -1.766 | 0.479 | 0.171 |
| 3016T>A | [1534-NA] | 0 | 170 | 0.000% | 6 | 2716 | 0.221% | -0.221% | 7.93E-01 | -0.657 | 1.677 | 0.518 |
| 3017G>A | [1534-NA] | 0 | 448 | 0.000% | 6 | 2716 | 0.221% | -0.221% | 4.75E-01 | -2.317 | 2.389 | 0.099 |
| 3019A>T | [1534-NA] | 0 | 3072 | 0.000% | 6 | 2716 | 0.221% | -0.221% | 2.31E-02 | -1.891 | 0.700 | 0.151 |
| 3050T>C | [1534-NA] | 0 | 59 | 0.000% | 6 | 2716 | 0.221% | -0.221% | 8.47E-01 | -0.984 | 3.392 | 0.374 |
| 3050T>G | [1534-NA] | 0 | 766 | 0.000% | 6 | 2716 | 0.221% | -0.221% | 3.40E-01 | -1.468 | 1.170 | 0.230 |
| 3054A>C | [1534-NA] | 0 | 54 | 0.000% | 6 | 2716 | 0.221% | -0.221% | 8.62E-01 | -0.896 | 3.392 | 0.408 |
| 3054A>T | [1534-NA] | 0 | 686 | 0.000% | 6 | 2716 | 0.221% | -0.221% | 3.69E-01 | -1.491 | 1.254 | 0.225 |
| 3055G>T | [1534-NA] | 0 | 322 | 0.000% | 6 | 2716 | 0.221% | -0.221% | 5.56E-01 | -1.582 | 1.943 | 0.206 |
| 3056G>T | [1534-NA] | 0 | 111 | 0.000% | 6 | 2716 | 0.221% | -0.221% | 7.96E-01 | -0.923 | 2.389 | 0.397 |
| 3057C>T | [1534-NA] | 0 | 2256 | 0.000% | 6 | 2716 | 0.221% | -0.221% | 5.87E-02 | -1.855 | 0.808 | 0.157 |
| 3081G>T | [1534-NA] | 0 | 238 | 0.000% | 6 | 2716 | 0.221% | -0.221% | 6.79E-01 | -0.993 | 1.677 | 0.371 |
| 3084A>G | [1534-NA] | 0 | 2385 | 0.000% | 6 | 2716 | 0.221% | -0.221% | 5.31E-02 | -1.638 | 0.701 | 0.194 |
| 3097A>G | [1534-NA] | 0 | 3049 | 0.000% | 6 | 2716 | 0.221% | -0.221% | 2.46E-02 | -1.750 | 0.654 | 0.174 |
| 3138C>G | [1534-NA] | 0 | 2263 | 0.000% | 6 | 2716 | 0.221% | -0.221% | 6.00E-02 | -1.858 | 0.813 | 0.156 |
| 3138C>G;3176G>T | [1534-NA] | 0 | 294 | 0.000% | 6 | 2716 | 0.221% | -0.221% | 5.84E-01 | -2.589 | 3.390 | 0.075 |
| 3144G>T | [1534-NA] | 0 | 2393 | 0.000% | 6 | 2716 | 0.221% | -0.221% | 4.95E-02 | -1.742 | 0.735 | 0.175 |
| 3149C>T;3269A>C | [1534-NA] | 0 | 273 | 0.000% | 6 | 2716 | 0.221% | -0.221% | 5.84E-01 | -1.822 | 2.389 | 0.162 |
| 3166T>G | [1534-NA] | 0 | 1015 | 0.000% | 6 | 2716 | 0.221% | -0.221% | 2.48E-01 | -2.219 | 1.494 | 0.109 |

| | | | | | | | | | | | | |
|---|---|---|---|---|---|---|---|---|---|---|---|---|
| 3166T>G;3289C>T | [1534-NA] | 0 | 33 | 0.000% | 6 | 2716 | 0.221% | -0.221% | 9.40E-01 | -0.405 | 3.394 | 0.667 |
| 3173C>A | [1534-NA] | 0 | 1039 | 0.000% | 6 | 2716 | 0.221% | -0.221% | 2.38E-01 | -2.060 | 1.359 | 0.127 |
| 3173C>G | [1534-NA] | 0 | 3322 | 0.000% | 6 | 2716 | 0.221% | -0.221% | 1.71E-02 | -1.970 | 0.696 | 0.140 |
| 3176G>C | [1534-NA] | 0 | 2429 | 0.000% | 6 | 2716 | 0.221% | -0.221% | 4.93E-02 | -1.657 | 0.699 | 0.191 |
| 3176G>C;3293A>T | [1534-NA] | 0 | 78 | 0.000% | 6 | 2716 | 0.221% | -0.221% | 8.02E-01 | -1.263 | 3.391 | 0.283 |
| 3176G>T | [1534-NA] | 0 | 3766 | 0.000% | 6 | 2716 | 0.221% | -0.221% | 9.66E-03 | -1.739 | 0.570 | 0.176 |
| 3176G>T;3244C>T | [1534-NA] | 0 | 193 | 0.000% | 6 | 2716 | 0.221% | -0.221% | 6.54E-01 | -2.168 | 3.390 | 0.114 |
| 3176G>T;3331C>T | [1534-NA] | 0 | 101 | 0.000% | 6 | 2716 | 0.221% | -0.221% | 7.61E-01 | -1.521 | 3.391 | 0.218 |
| 3223A>G | [1534-NA] | 0 | 3312 | 0.000% | 6 | 2716 | 0.221% | -0.221% | 1.71E-02 | -1.753 | 0.620 | 0.173 |
| 3223A>G;3317G>T | [1534-NA] | 0 | 140 | 0.000% | 6 | 2716 | 0.221% | -0.221% | 7.40E-01 | -1.155 | 2.389 | 0.315 |
| 3229C>G | [1534-NA] | 0 | 3420 | 0.000% | 6 | 2716 | 0.221% | -0.221% | 1.48E-02 | -1.711 | 0.593 | 0.181 |
| 3229C>G;3310G>A | [1534-NA] | 0 | 283 | 0.000% | 6 | 2716 | 0.221% | -0.221% | 5.91E-01 | -2.551 | 3.390 | 0.078 |
| 3244C>T | [1534-NA] | 0 | 4303 | 0.000% | 6 | 2716 | 0.221% | -0.221% | 7.32E-02 | -1.147 | 0.528 | 0.318 |
| 3244C>T;3252C>T | [1534-NA] | 0 | 66 | 0.000% | 6 | 2716 | 0.221% | -0.221% | 8.28E-01 | -1.096 | 3.392 | 0.334 |
| 3244C>T;3317G>T | [1534-NA] | 0 | 5 | 0.000% | 6 | 2716 | 0.221% | -0.221% | 7.72E-01 | 1.466 | 3.422 | 4.330 |
| 3269A>T | [1534-NA] | 0 | 126 | 0.000% | 6 | 2716 | 0.221% | -0.221% | 7.66E-01 | -1.050 | 2.389 | 0.350 |
| 3279G>T | [1534-NA] | 0 | 608 | 0.000% | 6 | 2716 | 0.221% | -0.221% | 3.91E-01 | -1.707 | 1.493 | 0.181 |
| 3283C>G | [1534-NA] | 0 | 662 | 0.000% | 6 | 2716 | 0.221% | -0.221% | 4.04E-01 | -1.610 | 1.444 | 0.200 |
| 3283C>G;3317G>T | [1534-NA] | 0 | 104 | 0.000% | 6 | 2716 | 0.221% | -0.221% | 7.56E-01 | -1.551 | 3.391 | 0.212 |
| 3289C>T | [1534-NA] | 0 | 2055 | 0.000% | 6 | 2716 | 0.221% | -0.221% | 7.41E-02 | -1.895 | 0.869 | 0.150 |
| 3310G>A | [1534-NA] | 0 | 641 | 0.000% | 6 | 2716 | 0.221% | -0.221% | 3.74E-01 | -1.760 | 1.496 | 0.172 |
| 3311G>C | [1534-NA] | 0 | 1064 | 0.000% | 6 | 2716 | 0.221% | -0.221% | 2.49E-01 | -2.489 | 1.678 | 0.083 |
| 3311G>T | [1534-NA] | 0 | 280 | 0.000% | 6 | 2716 | 0.221% | -0.221% | 5.78E-01 | -1.848 | 2.388 | 0.158 |
| 3317G>T;3326G>A | [1534-NA] | 0 | 92 | 0.000% | 6 | 2716 | 0.221% | -0.221% | 7.76E-01 | -1.428 | 3.391 | 0.240 |
| 3325A>G | [1534-NA] | 0 | 991 | 0.000% | 6 | 2716 | 0.221% | -0.221% | 2.52E-01 | -1.725 | 1.170 | 0.178 |
| 3331C>G | [1534-NA] | 0 | 1047 | 0.000% | 6 | 2716 | 0.221% | -0.221% | 2.35E-01 | -1.914 | 1.257 | 0.148 |
| 3331C>T | [1534-NA] | 0 | 1190 | 0.000% | 6 | 2716 | 0.221% | -0.221% | 2.00E-01 | -1.685 | 1.040 | 0.185 |
| 3338T>C | [1534-NA] | 0 | 288 | 0.000% | 6 | 2716 | 0.221% | -0.221% | 5.88E-01 | -2.568 | 3.390 | 0.077 |
| 3343C>G;3350T>G | [1534-NA] | 0 | 126 | 0.000% | 6 | 2716 | 0.221% | -0.221% | 7.23E-01 | -1.742 | 3.390 | 0.175 |
| 3347G>C | [1534-NA] | 0 | 360 | 0.000% | 6 | 2716 | 0.221% | -0.221% | 5.44E-01 | -1.406 | 1.685 | 0.245 |
| 3347G>T | [1534-NA] | 0 | 925 | 0.000% | 6 | 2716 | 0.221% | -0.221% | 3.24E-01 | -1.171 | 0.906 | 0.310 |
| 3348C>T | [1534-NA] | 0 | 687 | 0.000% | 6 | 2716 | 0.221% | -0.221% | 4.14E-01 | -1.136 | 1.042 | 0.321 |
| 3349A>G | [1534-NA] | 0 | 1298 | 0.000% | 6 | 2716 | 0.221% | -0.221% | 2.42E-01 | -1.080 | 0.719 | 0.340 |
| 3350T>G | [1534-NA] | 0 | 1383 | 0.000% | 6 | 2716 | 0.221% | -0.221% | 2.27E-01 | -1.048 | 0.681 | 0.351 |
| 3350T>G;3392A>G | [1534-NA] | 0 | 174 | 0.000% | 6 | 2716 | 0.221% | -0.221% | 6.68E-01 | -2.065 | 3.390 | 0.127 |
| 3350T>G;3511G>A | [1534-NA] | 0 | 39 | 0.000% | 6 | 2716 | 0.221% | -0.221% | 9.15E-01 | -0.571 | 3.393 | 0.565 |
| 3350T>G;3529T>G | [1534-NA] | 0 | 39 | 0.000% | 6 | 2716 | 0.221% | -0.221% | 9.15E-01 | -0.571 | 3.393 | 0.565 |
| 3363C>G | [1534-NA] | 0 | 2084 | 0.000% | 6 | 2716 | 0.221% | -0.221% | 1.08E-01 | -1.115 | 0.563 | 0.328 |
| 3363C>G;3399T>A | [1534-NA] | 0 | 63 | 0.000% | 6 | 2716 | 0.221% | -0.221% | 8.37E-01 | -1.050 | 3.392 | 0.350 |
| 3363C>G;3410T>A | [1534-NA] | 0 | 26 | 0.000% | 6 | 2716 | 0.221% | -0.221% | 9.76E-01 | -0.167 | 3.395 | 0.846 |
| 3363C>G;3512T>C | [1534-NA] | 0 | 22 | 0.000% | 6 | 2716 | 0.221% | -0.221% | 1.00E+00 | -0.001 | 3.397 | 0.999 |
| 3369A>T | [1534-NA] | 0 | 87 | 0.000% | 6 | 2716 | 0.221% | -0.221% | 9.98E-01 | 0.011 | 1.679 | 1.011 |
| 3369A>T;3400G>A | [1534-NA] | 0 | 21 | 0.000% | 6 | 2716 | 0.221% | -0.221% | 9.95E-01 | 0.046 | 3.397 | 1.047 |
| 3369A>T;3400G>A;3 | [1534-NA] | 0 | 36 | 0.000% | 6 | 2716 | 0.221% | -0.221% | 9.27E-01 | -0.491 | 3.394 | 0.612 |
| 3369A>T;3526A>T | [1534-NA] | 0 | 42 | 0.000% | 6 | 2716 | 0.221% | -0.221% | 9.03E-01 | -0.645 | 3.393 | 0.525 |
| 3386C>A | [1534-NA] | 0 | 2081 | 0.000% | 6 | 2716 | 0.221% | -0.221% | 1.03E-01 | -1.146 | 0.573 | 0.318 |
| 3386C>A;3507T>C | [1534-NA] | 0 | 63 | 0.000% | 6 | 2716 | 0.221% | -0.221% | 8.37E-01 | -1.050 | 3.392 | 0.350 |
| 3392A>G | [1534-NA] | 0 | 1390 | 0.000% | 6 | 2716 | 0.221% | -0.221% | 2.36E-01 | -1.008 | 0.664 | 0.365 |
| 3392A>G;3529T>G | [1534-NA] | 0 | 261 | 0.000% | 6 | 2716 | 0.221% | -0.221% | 6.50E-01 | -1.085 | 1.679 | 0.338 |

| | | | | | | | | | | | | |
|---|---|---|---|---|---|---|---|---|---|---|---|---|
| 3399T>A | [1534-NA] | 0 | 2179 | 0.000% | 6 | 2716 | 0.221% | -0.221% | 8.73E-02 | -1.226 | 0.585 | 0.294 |
| 3400G>A | [1534-NA] | 0 | 2874 | 0.000% | 6 | 2716 | 0.221% | -0.221% | 5.27E-02 | -1.064 | 0.456 | 0.345 |
| 3400G>A;3409T>G | [1534-NA] | 0 | 65 | 0.000% | 6 | 2716 | 0.221% | -0.221% | 8.31E-01 | -1.081 | 3.392 | 0.339 |
| 3400G>A;3450T>C | [1534-NA] | 0 | 198 | 0.000% | 6 | 2716 | 0.221% | -0.221% | 6.60E-01 | -1.501 | 2.388 | 0.223 |
| 3400G>A;3508G>A | [1534-NA] | 0 | 123 | 0.000% | 6 | 2716 | 0.221% | -0.221% | 7.28E-01 | -1.718 | 3.390 | 0.179 |
| 3400G>A;3529T>G | [1534-NA] | 0 | 60 | 0.000% | 6 | 2716 | 0.221% | -0.221% | 8.44E-01 | -1.001 | 3.392 | 0.367 |
| 3409T>G | [1534-NA] | 0 | 4846 | 0.000% | 6 | 2716 | 0.221% | -0.221% | 5.57E-03 | -1.188 | 0.368 | 0.305 |
| 3409T>G;3421G>T | [1534-NA] | 0 | 50 | 0.000% | 6 | 2716 | 0.221% | -0.221% | 8.75E-01 | -0.819 | 3.392 | 0.441 |
| 3409T>G;3523T>G;3! | [1534-NA] | 0 | 29 | 0.000% | 6 | 2716 | 0.221% | -0.221% | 9.61E-01 | -0.276 | 3.395 | 0.759 |
| 3410T>A | [1534-NA] | 0 | 5135 | 0.000% | 6 | 2716 | 0.221% | -0.221% | 4.96E-03 | -1.120 | 0.343 | 0.326 |
| 3410T>A;3507T>A | [1534-NA] | 0 | 98 | 0.000% | 6 | 2716 | 0.221% | -0.221% | 7.66E-01 | -1.491 | 3.391 | 0.225 |
| 3410T>A;3507T>C | [1534-NA] | 0 | 32 | 0.000% | 6 | 2716 | 0.221% | -0.221% | 9.44E-01 | -0.374 | 3.394 | 0.688 |
| 3410T>A;3508G>A | [1534-NA] | 0 | 76 | 0.000% | 6 | 2716 | 0.221% | -0.221% | 8.06E-01 | -1.237 | 3.391 | 0.290 |
| 3410T>A;3529T>G | [1534-NA] | 0 | 206 | 0.000% | 6 | 2716 | 0.221% | -0.221% | 6.43E-01 | -2.234 | 3.390 | 0.107 |
| 3410T>A;3422del;35 | [1534-NA] | 0 | 18 | 0.000% | 6 | 2716 | 0.221% | -0.221% | 9.71E-01 | 0.199 | 3.398 | 1.220 |
| 3419A>G;3508G>A | [1534-NA] | 0 | 58 | 0.000% | 6 | 2716 | 0.221% | -0.221% | 8.50E-01 | -0.967 | 3.392 | 0.380 |
| 3507T>C | [1534-NA] | 0 | 2763 | 0.000% | 6 | 2716 | 0.221% | -0.221% | 4.67E-02 | -1.193 | 0.500 | 0.303 |
| 3508G>A | [1534-NA] | 0 | 4690 | 0.000% | 6 | 2716 | 0.221% | -0.221% | 5.81E-03 | -1.217 | 0.379 | 0.296 |
| 3508G>A;3517A>G | [1534-NA] | 0 | 67 | 0.000% | 6 | 2716 | 0.221% | -0.221% | 8.25E-01 | -1.111 | 3.392 | 0.329 |
| 3508G>A;3529T>G | [1534-NA] | 0 | 109 | 0.000% | 6 | 2716 | 0.221% | -0.221% | 7.47E-01 | -1.597 | 3.391 | 0.202 |
| 3511G>A | [1534-NA] | 0 | 3996 | 0.000% | 6 | 2716 | 0.221% | -0.221% | 1.54E-02 | -1.122 | 0.394 | 0.326 |
| 3511G>A;3529T>G | [1534-NA] | 0 | 166 | 0.000% | 6 | 2716 | 0.221% | -0.221% | 6.77E-01 | -2.018 | 3.390 | 0.133 |
| 3511G>A;del3529 | [1534-NA] | 0 | 86 | 0.000% | 6 | 2716 | 0.221% | -0.221% | 8.54E-01 | -0.669 | 2.390 | 0.512 |
| 3512T>A | [1534-NA] | 0 | 1475 | 0.000% | 6 | 2716 | 0.221% | -0.221% | 1.51E-01 | -1.563 | 0.872 | 0.209 |
| 3512T>C | [1534-NA] | 0 | 1989 | 0.000% | 6 | 2716 | 0.221% | -0.221% | 9.93E-02 | -1.283 | 0.634 | 0.277 |
| 3512T>C;3526A>C | [1534-NA] | 0 | 34 | 0.000% | 6 | 2716 | 0.221% | -0.221% | 9.36E-01 | -0.434 | 3.394 | 0.648 |
| 3519T>A | [1534-NA] | 0 | 420 | 0.000% | 6 | 2716 | 0.221% | -0.221% | 5.13E-01 | -1.337 | 1.499 | 0.263 |
| 3519T>A;3522A>G | [1534-NA] | 0 | 1034 | 0.000% | 6 | 2716 | 0.221% | -0.221% | 2.85E-01 | -1.209 | 0.870 | 0.299 |
| 3519T>A;3523T>G;3! | [1534-NA] | 0 | 17 | 0.000% | 6 | 2716 | 0.221% | -0.221% | 9.64E-01 | 0.256 | 3.399 | 1.291 |
| 3519T>A;del3534 | [1534-NA] | 0 | 35 | 0.000% | 6 | 2716 | 0.221% | -0.221% | 9.53E-01 | 0.227 | 2.394 | 1.255 |
| 3522A>G | [1534-NA] | 0 | 1722 | 0.000% | 6 | 2716 | 0.221% | -0.221% | 1.45E-01 | -1.180 | 0.650 | 0.307 |
| 3523T>G | [1534-NA] | 0 | 362 | 0.000% | 6 | 2716 | 0.221% | -0.221% | 5.98E-01 | -1.007 | 1.361 | 0.365 |
| 3523T>G;3524T>A | [1534-NA] | 0 | 516 | 0.000% | 6 | 2716 | 0.221% | -0.221% | 4.46E-01 | -1.543 | 1.496 | 0.214 |
| 3524T>A | [1534-NA] | 0 | 958 | 0.000% | 6 | 2716 | 0.221% | -0.221% | 3.64E-01 | -0.939 | 0.783 | 0.391 |
| 3526A>C | [1534-NA] | 0 | 490 | 0.000% | 6 | 2716 | 0.221% | -0.221% | 5.52E-01 | -0.904 | 1.102 | 0.405 |
| 3526A>T | [1534-NA] | 0 | 3084 | 0.000% | 6 | 2716 | 0.221% | -0.221% | 3.39E-02 | -1.203 | 0.476 | 0.300 |
| 3529T>G | [1534-NA] | 0 | 10498 | 0.000% | 6 | 2716 | 0.221% | -0.221% | 1.32E-05 | -1.129 | 0.230 | 0.323 |
| 3558G>C;3559C>G | [1534-NA] | 0 | 25 | 0.000% | 6 | 2716 | 0.221% | -0.221% | 9.82E-01 | -0.128 | 3.396 | 0.880 |
| 67A>G | [1534-NA] | 0 | 20 | 0.000% | 6 | 2716 | 0.221% | -0.221% | 9.87E-01 | 0.094 | 3.397 | 1.099 |
| 68A>G | [1534-NA] | 0 | 1099 | 0.000% | 6 | 2716 | 0.221% | -0.221% | 2.29E-01 | -2.298 | 1.493 | 0.100 |
| 90A>T | [1534-NA] | 0 | 2673 | 0.000% | 6 | 2716 | 0.221% | -0.221% | 3.77E-02 | -1.801 | 0.723 | 0.165 |
| 111T>A | [1534-NA] | 0 | 937 | 0.000% | 6 | 2716 | 0.221% | -0.221% | 2.72E-01 | -1.669 | 1.171 | 0.188 |
| 125A>G | [1534-NA] | 0 | 1600 | 0.000% | 6 | 2716 | 0.221% | -0.221% | 1.29E-01 | -1.645 | 0.872 | 0.193 |
| 128A>G | [1534-NA] | 0 | 6034 | 0.000% | 6 | 2716 | 0.221% | -0.221% | 8.12E-04 | -2.056 | 0.527 | 0.128 |
| 128A>T | [1534-NA] | 0 | 2082 | 0.000% | 6 | 2716 | 0.221% | -0.221% | 7.81E-02 | -2.244 | 1.040 | 0.106 |
| 129G>T | [1534-NA] | 0 | 657 | 0.000% | 6 | 2716 | 0.221% | -0.221% | 3.98E-01 | -2.700 | 2.389 | 0.067 |
| 130T>G | [1534-NA] | 0 | 419 | 0.000% | 6 | 2716 | 0.221% | -0.221% | 5.12E-01 | -1.335 | 1.494 | 0.263 |
| 133T>C | [1534-NA] | 0 | 2773 | 0.000% | 6 | 2716 | 0.221% | -0.221% | 3.34E-02 | -1.889 | 0.741 | 0.151 |
| 156C>G | [1534-NA] | 0 | 1398 | 0.000% | 6 | 2716 | 0.221% | -0.221% | 1.58E-01 | -1.846 | 1.043 | 0.158 |

| | | | | | | | | | | | | |
|---|---|---|---|---|---|---|---|---|---|---|---|---|
| 191T>A | [1534-NA] | 0 | 83 | 0.000% | 6 | 2716 | 0.221% | -0.221% | 7.93E-01 | -1.325 | 3.391 | 0.266 |
| 196A>G | [1534-NA] | 0 | 944 | 0.000% | 6 | 2716 | 0.221% | -0.221% | 2.67E-01 | -1.810 | 1.260 | 0.164 |
| 200A>G | [1534-NA] | 0 | 258 | 0.000% | 6 | 2716 | 0.221% | -0.221% | 5.99E-01 | -1.766 | 2.393 | 0.171 |
| 205A>G | [1534-NA] | 0 | 1899 | 0.000% | 6 | 2716 | 0.221% | -0.221% | 9.15E-02 | -1.890 | 0.910 | 0.151 |
| 228T>G | [1534-NA] | 0 | 1283 | 0.000% | 6 | 2716 | 0.221% | -0.221% | 1.81E-01 | -1.665 | 0.991 | 0.189 |
| 231T>G | [1534-NA] | 0 | 1336 | 0.000% | 6 | 2716 | 0.221% | -0.221% | 1.68E-01 | -1.801 | 1.040 | 0.165 |
| 242C>G | [1534-NA] | 0 | 1016 | 0.000% | 6 | 2716 | 0.221% | -0.221% | 2.44E-01 | -1.750 | 1.170 | 0.174 |
| 246T>A | [1534-NA] | 0 | 352 | 0.000% | 6 | 2716 | 0.221% | -0.221% | 5.31E-01 | -1.671 | 1.944 | 0.188 |
| 247C>G | [1534-NA] | 0 | 835 | 0.000% | 6 | 2716 | 0.221% | -0.221% | 3.00E-01 | -1.842 | 1.359 | 0.159 |
| 256A>G | [1534-NA] | 0 | 408 | 0.000% | 6 | 2716 | 0.221% | -0.221% | 5.24E-01 | -1.308 | 1.497 | 0.270 |
| 257G>A | [1534-NA] | 0 | 1633 | 0.000% | 6 | 2716 | 0.221% | -0.221% | 1.23E-01 | -2.107 | 1.102 | 0.122 |
| 257G>C | [1534-NA] | 0 | 1659 | 0.000% | 6 | 2716 | 0.221% | -0.221% | 1.23E-01 | -1.548 | 0.809 | 0.213 |
| 258G>C | [1534-NA] | 0 | 2588 | 0.000% | 6 | 2716 | 0.221% | -0.221% | 4.24E-02 | -1.931 | 0.791 | 0.145 |
| 262T>G | [1534-NA] | 0 | 219 | 0.000% | 6 | 2716 | 0.221% | -0.221% | 6.33E-01 | -2.295 | 3.390 | 0.101 |
| 267A>T | [1534-NA] | 0 | 693 | 0.000% | 6 | 2716 | 0.221% | -0.221% | 3.60E-01 | -2.348 | 1.943 | 0.096 |
| 268A>C | [1534-NA] | 0 | 2609 | 0.000% | 6 | 2716 | 0.221% | -0.221% | 4.07E-02 | -2.000 | 0.813 | 0.135 |
| 278T>G | [1534-NA] | 0 | 9006 | 0.000% | 6 | 2716 | 0.221% | -0.221% | 1.77E-05 | -1.807 | 0.366 | 0.164 |
| 302C>A | [1534-NA] | 0 | 2711 | 0.000% | 6 | 2716 | 0.221% | -0.221% | 3.41E-02 | -1.815 | 0.716 | 0.163 |
| 303G>A | [1534-NA] | 0 | 1176 | 0.000% | 6 | 2716 | 0.221% | -0.221% | 2.00E-01 | -2.030 | 1.253 | 0.131 |
| 303G>C | [1534-NA] | 0 | 3223 | 0.000% | 6 | 2716 | 0.221% | -0.221% | 2.00E-02 | -1.988 | 0.718 | 0.137 |
| 304G>C | [1534-NA] | 0 | 2875 | 0.000% | 6 | 2716 | 0.221% | -0.221% | 2.93E-02 | -1.825 | 0.701 | 0.161 |
| 304G>T | [1534-NA] | 0 | 1868 | 0.000% | 6 | 2716 | 0.221% | -0.221% | 9.49E-02 | -1.874 | 0.913 | 0.154 |
| 305A>C | [1534-NA] | 0 | 2732 | 0.000% | 6 | 2716 | 0.221% | -0.221% | 3.41E-02 | -1.728 | 0.681 | 0.178 |
| 353T>G | [1534-NA] | 0 | 234 | 0.000% | 6 | 2716 | 0.221% | -0.221% | 6.22E-01 | -1.668 | 2.388 | 0.189 |
| 370T>G | [1534-NA] | 0 | 248 | 0.000% | 6 | 2716 | 0.221% | -0.221% | 6.64E-01 | -1.034 | 1.676 | 0.356 |
| 372T>A | [1534-NA] | 0 | 341 | 0.000% | 6 | 2716 | 0.221% | -0.221% | 5.33E-01 | -2.044 | 2.388 | 0.129 |
| 372T>G | [1534-NA] | 0 | 104 | 0.000% | 6 | 2716 | 0.221% | -0.221% | 7.56E-01 | -1.551 | 3.391 | 0.212 |
| 376T>G | [1534-NA] | 0 | 100 | 0.000% | 6 | 2716 | 0.221% | -0.221% | 7.62E-01 | -1.511 | 3.391 | 0.221 |
| 394C>G | [1534-NA] | 0 | 3312 | 0.000% | 6 | 2716 | 0.221% | -0.221% | 1.84E-02 | -2.067 | 0.738 | 0.127 |
| 404T>G | [1534-NA] | 0 | 366 | 0.000% | 6 | 2716 | 0.221% | -0.221% | 5.38E-01 | -1.423 | 1.680 | 0.241 |
| 1640C>G | [1544-1673] | 0 | 398 | 0.000% | 0 | 2716 | 0.000% | 0.000% | 9.48E-01 | -0.025 | 0.244 | 0.975 |
| 1640C>G;1684T>G | [1544-1673] | 0 | 121 | 0.000% | 0 | 2716 | 0.000% | 0.000% | 6.30E-01 | -0.338 | 0.496 | 0.713 |
| 1656A>G | [1544-1673] | 0 | 200 | 0.000% | 0 | 2716 | 0.000% | 0.000% | 8.79E-01 | 0.075 | 0.317 | 1.078 |
| 1656A>G;1657C>G | [1544-1673] | 0 | 195 | 0.000% | 0 | 2716 | 0.000% | 0.000% | 8.18E-01 | -0.123 | 0.358 | 0.885 |
| 1659T>G | [1544-1673] | 0 | 290 | 0.000% | 0 | 2716 | 0.000% | 0.000% | 7.98E-01 | -0.114 | 0.302 | 0.892 |
| 1659T>G;1785T>G | [1544-1673] | 0 | 29 | 0.000% | 0 | 2716 | 0.000% | 0.000% | 6.93E-01 | 0.395 | 0.698 | 1.485 |
| 1661T>A | [1544-1673] | 0 | 92 | 0.000% | 0 | 2716 | 0.000% | 0.000% | 5.45E-01 | 0.340 | 0.408 | 1.404 |
| 1661T>A;1725T>A | [1544-1673] | 0 | 111 | 0.000% | 0 | 2716 | 0.000% | 0.000% | 3.00E-01 | -0.944 | 0.697 | 0.389 |
| 1661T>G | [1544-1673] | 0 | 353 | 0.000% | 0 | 2716 | 0.000% | 0.000% | 6.98E-01 | -0.156 | 0.281 | 0.855 |
| 1665A>G | [1544-1673] | 0 | 490 | 0.000% | 0 | 2716 | 0.000% | 0.000% | 9.31E-01 | -0.033 | 0.239 | 0.968 |
| 1666T>G | [1544-1673] | 0 | 786 | 0.000% | 0 | 2716 | 0.000% | 0.000% | 6.22E-01 | -0.130 | 0.187 | 0.878 |
| 1670G>A | [1544-1673] | 0 | 515 | 0.000% | 0 | 2716 | 0.000% | 0.000% | 5.75E-01 | -0.177 | 0.227 | 0.837 |
| 1670G>C | [1544-1673] | 0 | 245 | 0.000% | 0 | 2716 | 0.000% | 0.000% | 9.06E-01 | 0.054 | 0.291 | 1.056 |
| 1677T>C | [1544-1673] | 0 | 213 | 0.000% | 0 | 2716 | 0.000% | 0.000% | 9.82E-01 | 0.012 | 0.314 | 1.012 |
| 1677T>G | [1544-1673] | 0 | 105 | 0.000% | 0 | 2716 | 0.000% | 0.000% | 1.06E-02 | 0.898 | 0.295 | 2.455 |
| 1678A>G | [1544-1673] | 0 | 87 | 0.000% | 0 | 2716 | 0.000% | 0.000% | 4.88E-01 | 0.395 | 0.419 | 1.485 |
| 1678A>T | [1544-1673] | 0 | 530 | 0.000% | 0 | 2716 | 0.000% | 0.000% | 7.35E-01 | -0.111 | 0.226 | 0.895 |
| 1682A>T | [1544-1673] | 0 | 1269 | 0.000% | 0 | 2716 | 0.000% | 0.000% | 7.28E-01 | -0.086 | 0.171 | 0.917 |
| 1684T>G | [1544-1673] | 0 | 1170 | 0.000% | 0 | 2716 | 0.000% | 0.000% | 6.57E-01 | 0.100 | 0.158 | 1.105 |

| | | | | | | | | | | | | |
|---|---|---|---|---|---|---|---|---|---|---|---|---|
| 1684T>G;1689T>G | [1544-1673] | 0 | 20 | 0.000% | 0 | 2716 | 0.000% | 0.000% | 4.12E-01 | 0.765 | 0.699 | 2.150 |
| 1684T>G;1725T>A | [1544-1673] | 0 | 66 | 0.000% | 0 | 2716 | 0.000% | 0.000% | 7.09E-01 | 0.267 | 0.493 | 1.305 |
| 1689T>G | [1544-1673] | 0 | 652 | 0.000% | 0 | 2716 | 0.000% | 0.000% | 6.19E-01 | -0.151 | 0.215 | 0.860 |
| 1692G>A | [1544-1673] | 0 | 862 | 0.000% | 0 | 2716 | 0.000% | 0.000% | 1.00E+00 | 0.000 | 0.184 | 1.000 |
| 1692G>A;1695G>A | [1544-1673] | 0 | 76 | 0.000% | 0 | 2716 | 0.000% | 0.000% | 5.57E-01 | -0.566 | 0.697 | 0.568 |
| 1692G>A;1807A>G | [1544-1673] | 0 | 72 | 0.000% | 0 | 2716 | 0.000% | 0.000% | 6.01E-01 | -0.512 | 0.697 | 0.599 |
| 1714T>A | [1544-1673] | 0 | 483 | 0.000% | 0 | 2716 | 0.000% | 0.000% | 5.28E-01 | -0.219 | 0.253 | 0.804 |
| 1721T>A | [1544-1673] | 0 | 335 | 0.000% | 0 | 2716 | 0.000% | 0.000% | 9.45E-01 | 0.029 | 0.268 | 1.029 |
| 1721T>A;1817C>A | [1544-1673] | 0 | 54 | 0.000% | 0 | 2716 | 0.000% | 0.000% | 8.29E-01 | -0.225 | 0.697 | 0.799 |
| 1725T>A | [1544-1673] | 0 | 1337 | 0.000% | 0 | 2716 | 0.000% | 0.000% | 8.87E-01 | -0.033 | 0.150 | 0.967 |
| 1732A>G | [1544-1673] | 0 | 176 | 0.000% | 0 | 2716 | 0.000% | 0.000% | 3.22E-01 | 0.384 | 0.296 | 1.468 |
| 1732A>G;1733T>G | [1544-1673] | 0 | 311 | 0.000% | 0 | 2716 | 0.000% | 0.000% | 7.95E-01 | 0.103 | 0.266 | 1.109 |
| 1733T>G | [1544-1673] | 0 | 13 | 0.000% | 0 | 2716 | 0.000% | 0.000% | 1.75E-01 | 1.193 | 0.699 | 3.299 |
| 1735C>G | [1544-1673] | 0 | 618 | 0.000% | 0 | 2716 | 0.000% | 0.000% | 9.40E-01 | -0.024 | 0.201 | 0.977 |
| 1741G>T | [1544-1673] | 0 | 417 | 0.000% | 0 | 2716 | 0.000% | 0.000% | 9.27E-01 | 0.033 | 0.229 | 1.034 |
| 1742A>C | [1544-1673] | 0 | 59 | 0.000% | 0 | 2716 | 0.000% | 0.000% | 7.61E-01 | -0.313 | 0.697 | 0.731 |
| 1745A>G | [1544-1673] | 0 | 129 | 0.000% | 0 | 2716 | 0.000% | 0.000% | 9.98E-01 | 0.003 | 0.423 | 1.003 |
| 1747A>C | [1544-1673] | 0 | 424 | 0.000% | 0 | 2716 | 0.000% | 0.000% | 3.62E-01 | -0.339 | 0.282 | 0.712 |
| 1747A>C;1765A>G | [1544-1673] | 0 | 49 | 0.000% | 0 | 2716 | 0.000% | 0.000% | 9.07E-01 | -0.128 | 0.697 | 0.880 |
| 1765A>G | [1544-1673] | 0 | 810 | 0.000% | 0 | 2716 | 0.000% | 0.000% | 5.68E-01 | -0.160 | 0.203 | 0.852 |
| 1765A>G;1817C>A | [1544-1673] | 0 | 103 | 0.000% | 0 | 2716 | 0.000% | 0.000% | 3.43E-01 | -0.870 | 0.697 | 0.419 |
| 1785T>G | [1544-1673] | 0 | 248 | 0.000% | 0 | 2716 | 0.000% | 0.000% | 7.62E-01 | -0.140 | 0.314 | 0.869 |
| 1788T>G | [1544-1673] | 0 | 156 | 0.000% | 0 | 2716 | 0.000% | 0.000% | 7.67E-01 | -0.187 | 0.427 | 0.829 |
| 1799A>G | [1544-1673] | 0 | 100 | 0.000% | 0 | 2716 | 0.000% | 0.000% | 6.56E-01 | 0.257 | 0.403 | 1.292 |
| 1802T>G | [1544-1673] | 0 | 205 | 0.000% | 0 | 2716 | 0.000% | 0.000% | 9.21E-01 | 0.050 | 0.318 | 1.051 |
| 1807A>G | [1544-1673] | 0 | 426 | 0.000% | 0 | 2716 | 0.000% | 0.000% | 5.63E-01 | -0.211 | 0.263 | 0.810 |
| 1817C>A | [1544-1673] | 0 | 790 | 0.000% | 0 | 2716 | 0.000% | 0.000% | 9.51E-01 | -0.018 | 0.184 | 0.982 |
| 1826C>G | [1544-1673] | 0 | 30 | 0.000% | 0 | 2716 | 0.000% | 0.000% | 7.21E-01 | 0.362 | 0.698 | 1.436 |
| 1826C>G;1827T>A | [1544-1673] | 0 | 49 | 0.000% | 0 | 2716 | 0.000% | 0.000% | 4.06E-01 | 0.563 | 0.507 | 1.757 |
| 1830T>A | [1544-1673] | 0 | 171 | 0.000% | 0 | 2716 | 0.000% | 0.000% | 9.90E-01 | 0.008 | 0.358 | 1.008 |
| 1834A>G | [1544-1673] | 0 | 69 | 0.000% | 0 | 2716 | 0.000% | 0.000% | 7.61E-01 | 0.222 | 0.493 | 1.249 |
| 1839T>G | [1544-1673] | 0 | 145 | 0.000% | 0 | 2716 | 0.000% | 0.000% | 8.53E-01 | -0.114 | 0.405 | 0.892 |

**Table S5** – predicted ME and SpliceAI scores of activating mutations and the PE inclusion levels

| **mutation** | **type** | **PE activation (%)** | **MaxEnt score** | **MaxEnt partner** | **sum ME** | **delta ME** (mutation-induced change) | **Splice AI A gain** | **Splice AI D gain** | **Splice AI gain sum** |
|---|---|---|---|---|---|---|---|---|---|
| 1012A>T | CA | 1.72 | 7.71 | 9.6 | 17.31 | 5.86 | 0.17 | 0.04 | 0.21 |
| 1012A>C | CA | 2.16 | 7.18 | 9.6 | 16.78 | 5.33 | 0.17 | 0.03 | 0.2 |
| 1019G>T | CA | 3.43 | 3.57 | 9.6 | 13.17 | 1.72 | 0.16 | 0.02 | 0.18 |
| ***average*** | | ***2.44*** | ***6.15*** | ***9.60*** | ***15.75*** | ***4.30*** | ***0.17*** | ***0.03*** | ***0.20*** |
| 912G>A | CD | 0.7 | 8.05 | 7.41 | 15.46 | 4.11 | 0.06 | 0.21 | 0.27 |
| 1994A>G | CD | 3.8 | 7.48 | 7.65 | 15.13 | 7.54 | 0.74 | 0.66 | 1.4 |
| ***average*** | | ***2.25*** | ***7.77*** | ***7.53*** | ***15.30*** | ***5.83*** | ***0.40*** | ***0.44*** | ***0.84*** |
| 1322T>A | DA | 1.23 | 7.71 | 9.6 | 17.31 | 7.71 | 0.38 | 0.57 | 0.95 |
| 594A>G | DA | 4.75 | 6.83 | 9.48 | 16.31 | 6.83 | 0.71 | 0.5 | 1.21 |
| 545A>G | DA | 21.56 | 11.09 | 9.48 | 20.57 | 11.09 | 0.79 | 0.57 | 1.36 |
| ***average*** | | ***9.18*** | ***8.54*** | ***9.52*** | ***18.06*** | ***8.54*** | ***0.63*** | ***0.55*** | ***1.17*** |
| 1156C>G | DD | 8.9 | 7.51 | 1.85 | 9.36 | 7.51 | 0.39 | 0.36 | 0.75 |
| 2025T>G | DD | 17.13 | 6.69 | 7.65 | 14.34 | 6.69 | 0.81 | 0.87 | 1.68 |
| 1137C>T | DD | 52.43 | 9.8 | 1.85 | 11.65 | 9.8 | 0.71 | 0.72 | 1.43 |
| ***average*** | | ***26.15*** | ***8.00*** | ***3.78*** | ***11.78*** | ***8.00*** | ***0.64*** | ***0.65*** | ***1.29*** |

**Table S6** - Comparison of ESRseq scores between exons, pseudoexons and their flanking intronic sequences

| Compared cathegories | Type of test | method | n group 1 | n group 2 | p (adj) | significance | method | p (adj) | signifi-ance |
|---|---|---|---|---|---|---|---|---|---|
| Herein confirmed PEs vs its Upstream | Paired | Wilcoxon signed rank test with continuity correction | 4 | 4 | 0.137 | ns | Paired t-test | 0.057 | ns |
| Herein confirmed vs its Downstream | Paired | | 4 | 4 | 0.137 | ns | | 0.161 | ns |
| **Literature-derived PEs vs its Upstream** | **Paired** | | **17** | **17** | **0.009** | * | | **0.004** | * |
| **Literature-derived PEs vs its Downstream** | **Paired** | | **17** | **17** | **0.033** | * | | **0.011** | * |
| Non-used Acceptors vs its Upstream | Paired | | 19 | 19 | 1 | ns | | 0.894 | ns |
| Non-used Donors vs its Downstream | Paired | | 24 | 24 | 0.613 | ns | | 0.748 | ns |
| **NF1 exons vs its Upstream** | **Paired** | | **55** | **55** | **7.47E-09** | * | | **1.92E-16** | * |
| **NF1 exons vs its Downstream** | **Paired** | | **55** | **55** | **7.47E-09** | * | | **1.80E-14** | * |
| Herein confirmed PEs vs Non-used Acceptors | Unpaired | Wilcoxon rank sum test with continuity correction | 4 | 19 | 0.686 | ns | Welch Two Sample t-test | 0.511 | ns |
| Herein confirmed PEs vs Non-used Donors | Unpaired | | 4 | 24 | 0.259 | ns | | 0.175 | ns |
| Literature-derived PEs vs Non-used Acceptors | Unpaired | | 17 | 19 | 0.211 | ns | | 0.314 | ns |
| Literature-confirmed PSEX vs Neg PSEX (Don) | Unpaired | | 17 | 24 | 0.105 | ns | | 0.116 | ns |
| Upstream: herein confirmed PEs vs Non-used Acceptors | Unpaired | | 4 | 19 | 0.085 | ns | | 0.090 | ns |
| Downstream: herein confirmed PEs vs Non-used Donors | Unpaired | | 4 | 24 | 0.159 | ns | | 0.239 | ns |
| **Upstream: literature-derived PEs vs Non-used Acceptors** | **Unpaired** | | **17** | **19** | **0.036** | * | | **0.031** | * |
| Downstream: literature-derived PEs vs Non-used Donors | Unpaired | | 17 | 24 | 0.130 | ns | | 0.161 | ns |
| **Herein confirmed PEs vs NF1 Exons** | **Unpaired** | | **4** | **55** | **0.049** | * | | **0.018** | * |
| **Herein confirmed PEs vs Upstream of NF1 Exons** | **Unpaired** | | **4** | **55** | **0.027** | * | | **0.004** | * |
| **Herein confirmed PEs vs Downstream of NF1 Exons** | **Unpaired** | | **4** | **55** | **0.049** | * | | **0.003** | * |
| **Literature-derived PEs vs NF1 Exons** | **Unpaired** | | **17** | **55** | **0.049** | * | | 0.057 | ns |
| **Literature-derived PEs vs Upstream of NF1 Exons** | **Unpaired** | | **17** | **55** | **1.13E-04** | * | | **4.98E-04** | * |
| **Literature-derived PEs vs Downstream of NF1 Exons** | **Unpaired** | | **17** | **55** | **4.52E-04** | * | | **5.44E-04** | * |

**Table S7a** - Effect of AsiSI restriction sites introduction on ME scores of cryptic donors

| AsiSI # | intronic position (G from GT dinucleotide) | wt sequence | mut sequence | ME score WT | ME score mut |
|---|---|---|---|---|---|
| 1 | 61 | tatgtccca | catgtccca | -18.82 | -13.11 |
| 3 | 496 | taggtatgt | / | 7.76 | |
| 4 | 722 | catgtgtat | gctgtgtat | -5.45 | -9.26 |
| 5 | 945 | actgtgtat | / | -8.09 | |
| 5 | 947 | tgtgtatcc | / | -5.13 | |
| 6 | 1188 | ggggttcac | gcggttcac | -9.75 | -6.14 |
| 7 | 1395 | ctagttgtt | ctagttgcg | -14.86 | -22.78 |
| 7 | 1398 | gttgtttta | / | -27.53 | |
| 8 | 1614 | ggggtcata | ggggtcagc | -5.23 | 1.33 |
| 9 | 1838 | aatgtgtat | / | -5.65 | |
| 9 | 1840 | tgtgtatgc | / | 0.72 | |
| 10 | 2061 | tgagtgcaa | tgagtgcga | -15.76 | -3.84 |
| 10 | 2071 | tttgtagaa | cgcgtagaa | -14.18 | -12.73 |
| 11 | 2285 | agagtgctt | / | -6.5 | |
| 12 | 2487 | aaagtagag | aaagtagcg | -3.25 | -2.57 |
| 15 | 3126 | cctgtaatc | / | -6.49 | |
| 18 | 3759 | tctgtaatt | cgcgtaatt | -1.71 | 2.46 |
| 19 | 3933 | gatgtagca | / | -3.36 | |

The slash symbol "/" stands for disruption of GT dinucleotide

**Table S7b** - Effect of AsiSI restriction sites introduction on ME scores of cryptic acceptors

| AsiSI # | intronic position (G from AG) | wt sequence | mut sequence | ME score wt | ME score mut |
|---|---|---|---|---|---|
| 1 | 69 | aatgatcttatgtcccaaagtac | agcgatcgcatgtcccaaagtac | 2.17 | -0.25 |
| 1 | 74 | tcttatgtcccaaagtacagatg | tcgcatgtcccaaagtacagatg | 1 | -0.79 |
| 2 | 282 | taggaaaatattttatgaagaga | taggaaaatattttatgaagcga | -3.7 | -2.45 |
| 2 | 284 | ggaaaatattttatgaagagatc | / | -12.73 | / |
| 3 | 495 | gacacacaaatatgcgataggta | / | 2.56 | / |
| 3 | 505 | tatgcgataggtatgtggaggag | tatgcgatcgctatgtggaggag | -11.24 | -3.32 |
| 3 | 508 | gcgataggtatgtggaggaggag | gcgatcgctatgtggaggaggag | -15.32 | -12.82 |
| 3 | 511 | ataggtatgtggaggaggagagt | atcgctatgtggaggaggagagt | -29.28 | -27.01 |
| 3 | 513 | aggtatgtggaggaggagagtac | cgctatgtggaggaggagagtac | -24.26 | -22.58 |
| 5 | 956 | acacaactgtgtatccctaggga | acacaactggcgatcgctaggga | 8.38 | 1.79 |
| 5 | 961 | actgtgtatccctagggaaggct | actggcgatcgctagggaaggct | -1.84 | -5.04 |
| 5 | 969 | tccctagggaaggctgtaagcga | tcgctagggaaggctgtaagcga | -12.63 | -14.21 |
| 6 | 1179 | ttttttgtattttcataagcca | ttttttgtattttcataagcga | 2.5 | 3.76 |
| 6 | 1205 | gggttcactatgttggtcaggct | cggttcactatgttggtcaggct | -0.02 | 0.86 |
| 8 | 1634 | tcatatattgccttgttcagttt | tcagcgatcgccttgttcagttt | 6.83 | 4.27 |
| 8 | 1642 | tgccttgttcagtttcacagctt | cgccttgttcagtttcacagctt | 2.18 | 1.87 |
| 10 | 2074 | ttatgagtgcaattttgtagaat | ttatgagtgcgatcgcgtagaat | 2.89 | 2.19 |
| 10 | 2082 | gcaattttgtagaatttgaggaa | gcgatcgcgtagaatttgaggaa | -5.68 | -7.54 |
| 11 | 2299 | tgagagtgcttggattgcagtta | tgcgatcgcttggattgcagtta | 1.31 | 4.19 |
| 11 | 2305 | tgcttggattgcagttacagaat | cgcttggattgcagttacagaat | -2.07 | -2.08 |
| 12 | 2490 | agtatatatttacaaagtagaga | agtatatatttacaaagtagcga | -2.06 | -2.11 |
| 12 | 2492 | tatatatttacaaagtagagatt | / | -9.3 | / |
| 12 | 2513 | tttcactgttaacaccgaagtta | tcgcactgttaacaccgaagtta | 1.61 | -0.2 |
| 13 | 2705 | gaatcccagcactttgggagatc | / | -13.86 | / |
| 13 | 2711 | cagcactttgggagatcgagaca | / | -19.21 | / |
| 13 | 2715 | actttgggagatcgagacaggcg | actttgggcgatcgcgacaggcg | -8.17 | 3.56 |
| 13 | 2721 | ggagatcgagacaggcggagagc | ggcgatcgcgacaggcggagagc | -31.65 | -23.58 |
| 13 | 2723 | agatcgagacaggcggagagctt | cgatcgcgacaggcggagagctt | -20.97 | -16.61 |
| 13 | 2729 | agacaggcggagagcttgaggtc | cgacaggcggagagcttgaggtc | -26.22 | -25.14 |
| 14 | 2914 | attgtgccactgcactccagcct | attgtgccactgcactccagcga | 2.39 | 3.34 |
| 14 | 2926 | cactccagcctgggccacagagc | cactccagcgatcgccacagagc | 0.35 | 1.76 |
| 14 | 2928 | ctccagcctgggccacagagcga | ctccagcgatcgccacagagcga | -21.03 | -18.66 |
| 14 | 2932 | agcctgggccacagagcgagact | agcgatcgccacagagcgagact | -20.58 | -20.64 |
| 15 | 3135 | gctcacgcctgtaatcccagcac | gctcacgcctgcgatcgcagcac | 0.09 | 2.04 |
| 15 | 3146 | taatcccagcactttgggaggcc | cgatcgcagcactttgggaggcc | -12.58 | -12.63 |
| 16 | 3335 | ggaagttgcagtgagctgagatt | / | -18.65 | / |
| 16 | 3352 | gagattgcgcccttgcatagaat | gcgatcgcgcccttgcatagaat | 3.46 | 3.85 |
| 16 | 3357 | tgcgcccttgcatagaatagtga | cgcgcccttgcatagaatagtga | -2.02 | -1.89 |
| 17 | 3553 | ttagagaaaattcttttaagaga | ttagagaaaattcttttaagcga | -2.07 | -1.16 |
| 17 | 3555 | agagaaaattcttttaagagatg | / | -9.79 | / |
| 17 | 3563 | ttcttttaagagatgccaaggca | attcttttaagcgatcgcaaggc | -12.15 | -9.63 |
| 17 | 3578 | ccaaggcacatggggtgaagtat | gcaaggcacatggggtgaagtat | -7.79 | -8.67 |
| 18 | 3750 | atcaatcaatatgaattaagtcg | atcaatcaatatgaattaaggcg | -3.54 | -1.34 |
| 18 | 3768 | agtcgattctgtaattgtagaac | aggcgatcgcgtaattgtagaac | 2.76 | 1.23 |
| 18 | 3777 | tgtaattgtagaactcacagcaa | cgtaattgtagaactcacagcaa | -3.24 | -4.36 |
| 19 | 3930 | attaaactatctatagacagatg | attaaactatctatagacagcga | 0.95 | -0.9 |
| 19 | 3936 | ctatctatagacagatgtagcaa | / | -9.4 | / |

The slash symbol "/" stand for disruption of AG dinucleotide
